# Taxonomic, genomic, and functional variation in the gut microbiomes of wild spotted hyenas across two decades of study

**DOI:** 10.1101/2022.08.02.502164

**Authors:** Connie A. Rojas, Kay E. Holekamp, Mariette Viladomat Jasso, Valeria Souza, Jonathan A. Eisen, Kevin R. Theis

**Author notes:** **Name and email addresses of authors (in order of appearance):** Kay E. Holekamp, Mariette Viladomat Jasso, Valeria Souza, Jonathan A. Eisen, Kevin R. Theis. Connie Rojas, (corresponding author). **Full postal address of corresponding author:** Genome and Biomedical Sciences Facility (GBSF) 5212, 451 Health Sciences Drive, University of California, Davis, Davis, CA 95616.

## Abstract

The gut microbiome provides vital functions for mammalian hosts, yet research on the variability and function of the microbiome across adult lifespans and multiple generations is limited in large mammalian carnivores. Here we use 16S rRNA gene and metagenomic sequencing to profile the taxonomic composition, genomic diversity, and metabolic function of the gut microbiome of 12 wild spotted hyenas (*Crocuta crocuta*) residing in the Masai Mara National Reserve, Kenya over a 23-year period spanning three generations. We determined the extent to which host factors predict variation in the gut microbiome and identify the core microbes present in the guts of hyenas. We also investigate novel genomic diversity in the mammalian gut by reporting the first metagenome-assembled genomes (MAGs) for hyenas. We found that gut microbiome taxonomic composition was highly variable across the two decades of sampling, but despite this, a core set of 14 bacterial genera and 19 amplicon sequence variants were identified. The strongest predictors of microbiome alpha and beta-diversity were host identity and age, suggesting that hyenas possess individualized microbiomes, and that these may change with age during adulthood. Gut microbiome functional profiles were also individual-specific, and were moderately correlated with antelope prey abundance, indicating that the functions of the gut microbiome vary with host diet. We recovered 149 high-quality MAGs from the hyena gut, spanning 25 bacterial orders and 51 genera. Some MAGs were classified as taxa previously reported for other carnivores, but many were novel and lacked species level matches to genomes in existing reference databases.

**Importance:** There is a gap in knowledge regarding the genomic diversity and variation of the gut microbiome across a host’s lifespan and across multiple generations of hosts in wild mammals. Using two types of sequencing approaches, we demonstrate that although gut microbiomes are individualized and temporally variable among hyenas, they correlate similarly to large-scale changes in their host’s ecological environment. We also recovered 149 high-quality MAGs from the hyena gut, greatly expanding the microbial genome repertoire known for hyenas, carnivores and wild mammals in general. Some MAGs came from genera abundant in the gastrointestinal tracts of canid species and other carnivores but over 80% of MAGs were novel and from species previously not represented in genome databases. Collectively, our novel body of work illustrates the importance of surveying the gut microbiome of non-model wild hosts, using multiple sequencing methods and computational approaches, and at distinct scales of analysis.

## Introduction

Across mammals, the taxonomic composition of the gut microbiome, specifically the bacterial portion, varies widely among individuals due to variation in diet, reproductive state, age, disease status, habitat, and social interactions of the host (1–10). The gut microbiome can also vary rapidly within hosts over short-time scales on the order of days or weeks, which may be due to fluctuations in body temperature, circadian rhythms, a recent meal, or a stressful event (11, 12). Thus, to accurately capture the dynamism of these intestinal communities and their responses to environmental perturbations, longitudinal studies are required. Yet, longitudinal studies of the gut microbiome across host lifespan are limited, particularly in wild, long-lived mammals. Little is known regarding microbiome trajectories within hosts over time and across hosts within a social group, and about the relative influences of temporal, ecological, and host factors on these microbiome dynamics. A recent study sampled the gut microbiomes of a population of meerkats (*Suricata suricatta*) from 1997-2019 and found that across the meerkat lifespan, annual variation was the strongest predictor of gut microbiome composition (13). Over shorter time scales however, host identity outweighed the effects of the other host factors, particularly when samples were collected within two months of each other (13). The gut microbiomes of meerkats also showed daily diurnal oscillations, in part due to their temperature-constrained foraging schedules. Another recent study examined the gut microbiomes of baboons (*Papio* spp.) over a 14-year period and concluded that although all gut microbiomes exhibited cyclical seasonal shifts in composition, microbiome dynamics across baboons were only weakly synchronized (14). Instead, baboons exhibited largely individualized gut microbiomes over their lifespans, despite their shared diets, environments, and opportunities for between-host microbial dispersal (14). Findings from the two studies suggest that gut microbiome dynamics may be species-specific and influenced by host behavior, temporal factors, and habitat characteristics.

Here, we expand upon this work by investigating gut microbiome dynamics across host lifespan and across multiple generations among wild spotted hyenas (*Crocuta crocuta*) over a 23-year period. Hyenas are socially complex, long-lived carnivores (15), whose groups contain multiple overlapping generations of adult females and their young, along with breeding males that generally immigrated from other clans (15–19). They live in matrilineal societies that are structured by linear dominance hierarchies wherein an individual’s social rank, which is not dependent on body size or fighting ability, determines access to resources and fitness (15, 20–23). Hyena societies are also characterized by strong fission-fusion dynamics. Individuals mainly travel, rest and forage alone or in small subgroups (“fission”) that can ‘fuse’ into larger groups (20, 24, 25). The spotted hyenas from our study population reside in the Masai Mara National Reserve, Kenya (MMNR) (Table 1, Table S1), a savanna habitat that supports high densities of herbivores and carnivores (26). This habitat has undergone significant changes to its landscape and wildlife over the 23-year study period. A consistent decline over time in the densities of wild herbivores and lions has typically been accompanied by an exponential increase in anthropogenic disturbance and livestock grazing within the Reserve (19, 27, 28). These changes have also coincided with increases in hyena population size (29). Given these complex social and ecological dynamics, we expected large variability in gut microbiome taxonomic composition at one or more temporal scales: across the 23-year study period, across host lifespan, and across generations.

**Table 1.**
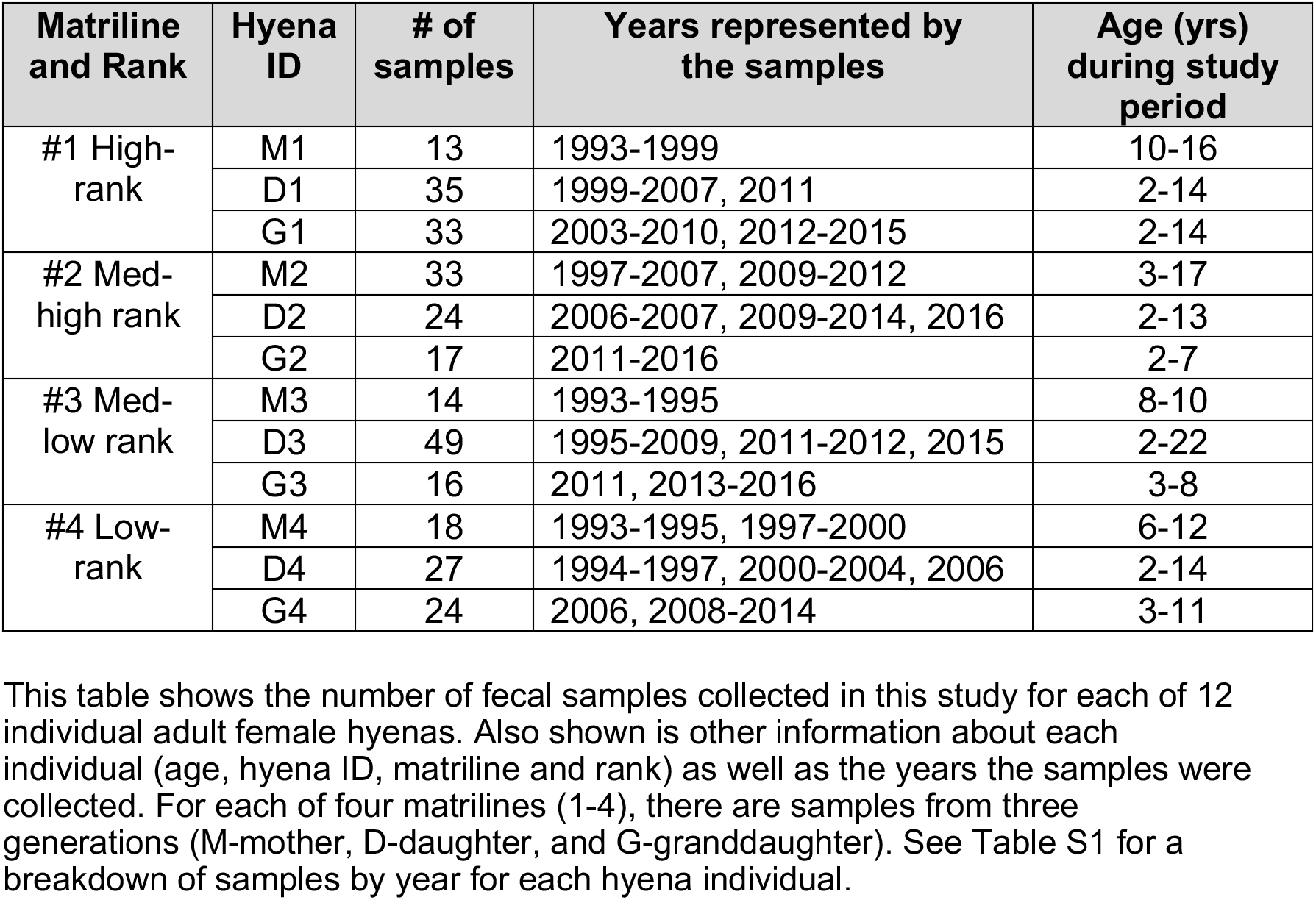
Longitudinal fecal samples collected from 12 adult wild spotted hyenas during 1993-2016.

Given that the metabolic functions of resident gut microbes can directly affect an animal’s development, physiology, behavior, and interactions with other organisms (30, 31), we also assayed gut microbiome functional variation among individuals and matrilines over a shorter time scale of two years. Hyenas are generalist carnivores, and can eat all parts of their prey, including the skin, muscle, bone, and viscera and an array of prey species, ranging from insects and antelope to elephants and hippos (15, 32, 33). In the MMNR, herbivore densities fluctuate throughout the year, which impacts the hyena’s diet and possibly their microbiome. Specifically, during July through October of every year, hyenas feed on large migratory herds of wildebeest (*Connochaetes taurinus*) and zebra (*Equus quagga*) (32, 34) arriving from the Serengeti. The rest of the year, hyenas mainly prey on topi (*Damaliscus lunatus*) and Thomson’s gazelles (*Eudorcas thomsonii*). During the period when ungulate numbers are low, hyenas may also scavenge more (32). Thus, gut microbiome functions may fluctuate with prey densities in this species, echoing prior studies that have found direct links between host diet and microbiome functional potential. One previous study examined the functional repertoire of the gut microbiome in 77 mammalian species and found that herbivores were enriched in plant carbohydrate degradation pathways compared to carnivores and piscivores (35). In contrast, carnivore gut microbiomes were enriched in pathways related to the degradation of choline, an amine principally found in meats. In hyenas, their diet may also be reflected in their gut microbiome functional repertoire.

We used 16S rRNA gene sequencing and shotgun metagenomic sequencing to profile gut microbiome composition and function in 12 adult female hyenas over a 23-year study period. Sampling collectively spanned three generations of female hyenas: mothers, daughters, and granddaughters (Table 1, Table S1). All sampled individuals were members of a single social group but belonged to distinct maternal lineages (matrilines) that varied in their social rank. Individuals ranged in age from 2.4-22 years over the study period, spanning these animal’s natural adult lifespans. On average we collected 25 samples from each of these females across 9.8 years (range = 13 to 48 samples per hyena) with a median of 3.4 yrs between consecutive samples (Table 1, Table S1). With this dataset, we assay taxonomic variation in the gut microbiome over the two decades of sampling, and examine whether any of the variation can be explained by host predictors including individual identity, matriline, age, prey availability, and calendar year. We also identify the gut microbial taxa that constitute the ‘core’ gut microbiome in wild spotted hyenas, which consistently persist over host lifespans and across maternal generations. These taxa may be functionally important to gut health and animal function (36, 37). Third, we profile gut microbiome metagenomes and predicted functions over two years, and investigate whether these vary with host prey densities. Lastly, we report the first metagenome-assembled genomes (MAGs) recovered from hyenas; expanding on what is known about the taxonomic and genomic diversity of the mammalian gut, particularly in less studied species like hyenas. Collectively, our findings provide a novel perspective on the variability, genomic diversity, and function of the gut microbiome in a wild African carnivore using multiple sequencing types over long and short temporal scales.

## Materials and Methods

### Sample and metadata collection

The Masai Mara National Reserve (MMNR; 1,530 km^2^) in southwestern Kenya (1°40’S, 35°50’E) is a rolling grassland habitat that constitutes the northernmost portion of the Mara-Serengeti ecosystem (38–42). The Reserve has two dry seasons (late December-March, and late June-mid November) and two rainy seasons (late November-early December, and April-early June) (28, 43).

Fecal samples from female members of a single social group were collected during the mornings and evenings as they were encountered. Our dataset was restricted to longitudinally collected fecal samples (N=303) from 12 adult (> 2 years old) female spotted hyenas inhabiting the MMNR between 1993 and 2016 (Table 1, Table S1). Upon collection, fecal samples were stored in cryogenic vials in liquid nitrogen until being transported on dry ice to Michigan State University, wherein they were stored at −80 °C until genomic DNA extractions were performed.

In the field, hyenas were identified as individuals by their unique spot patterns, sexed based on the dimorphic morphology of their erect phallus (44), and their birthdates were calculated to ± 7 days based on their appearance as cubs when first observed (45). Each hyena was assigned a dominance rank based on its position in a matrix ordered by submissive behaviors displayed during dyadic agonistic encounters (21) (Table S2). In hyena societies, each new offspring inherits the rank immediately below that of its mother but above those of its older siblings. The four hyena lineages that were sampled in our study varied in their rank with individuals from matriline 1 (M1) occupying the highest ranks in the clan’s hierarchy, and individuals from matriline 4 (M4) occupying some of the lowest ranks in the hierarchy. Individuals from matriline 2 (M2) were high-ranking hyenas, but below all individuals from M1, and individuals from matriline 3 (M3) were low-ranking but not as low-ranking as hyenas from M4. About 77% of all samples were collected from nursing females, 10% of samples came from pregnant females, and the rest came from either nulliparous (e.g., has never given birth) or non-pregnant/non-lactating females (Table S2).

To assay prey abundance, three 4-km line-transects in the clan’s territory were sampled biweekly and all mammalian herbivores were counted within 100 m of each transect centerline. The number of herbivores was summed across the three transects, as detailed in Holekamp *et al*. 1999 (46). These values were averaged to calculate the mean number of herbivores counted during the 30 days prior to each fecal sample being collected (Table S2).

### DNA extractions

Genomic DNA was extracted from the fecal samples using QIAGEN DNeasy PowerSoil Kits (QIAGEN, Valencia, CA), following the manufacturers’ recommended protocol. The order of extractions was randomized by assigning each sample a random number without replacement and then conducting DNA extractions based on this order. Blank extraction kit controls (e.g., sterile swabs) were included to account for any background DNA contamination. The ability to PCR amplify 16S rRNA genes from samples was tested using bacterial-specific primers (8F: 5’ – AGAGTTTGATCCTGGCTCAG – 3’; 1492R: 5’ – ACGGCTACCTTGTTACGACTT – 3’) and gel electrophoresis. The PCR conditions were as follows: an initial denaturation step at 95 °C for 3 minutes, followed by 30 cycles of 95 °C for 45 seconds, 50 °C for 60 seconds and 72 °C for 90 seconds. A final extension occurred at 72 °C for 10 minutes, and a final hold at 15 °C. DNA concentrations were quantified using QUBIT (Invitrogen), which ranged from 1.5-27.8 ng/μL for our samples (mean: 6.3 ng/μL).

### Sequencing and processing of 16S rRNA reads

DNA from all fecal samples (N=303) was sent for multiplexed paired-end 16S rRNA gene sequencing on the Illumina MiSeq v2 platform to the Michigan State University Genomics Core. The V4 hypervariable region of the 16S rRNA gene (250 bp) was amplified with dual indexed, Illumina compatible primers (515f/806r). Sequencing, library preparation, and preliminary quality filtering were completed according to Caporaso *et al*. 2012 (47) and Kozich *et al*. 2013 (48). Base calling was done by Illumina Real Time Analysis (RTA) (v1.18.54) and output of RTA was demultiplexed and converted to FastQ format with Illumina Bcl2fastq (v2.19.0).

Raw Illumina amplicon sequence reads were processed, filtered for quality, and classified into amplicon sequence variants (ASVs) using the Divisive Amplicon Denoising Algorithm (DADA2 v1.14.1) pipeline in R (v3.6.2) (49, 50). Briefly, reads were filtered for quality, allowing for two and three errors per forward and reverse read, respectively. To remove the low-quality portions of the sequences, forward reads were trimmed to 250 bp while reverse reads were trimmed to 220 bp. After calculating error rates, ASVs were inferred using DADA2’s core denoising algorithm. Forward and reverse reads were then merged to calculate ASV relative abundances. It is important to note that the DADA2 pipeline performs merging of paired-end reads after denoising to achieve greater accuracy (50). After this, chimeric sequences were removed, leaving an average of 13,411 ± 5431 sequences per sample (Fig S1A). The resulting ASVs were assigned a taxonomy using the SILVA rRNA gene reference database (v132) (51) and those classified as Eukarya, Chloroplasts, or Mitochondria, were removed from the dataset. Not all sequences were classified to genera or species-level and in those scenarios, their last known classification (e.g., Family) was used. We exported the final ASV relative abundance table, table of ASV taxonomic designations, and sample metadata into R for statistical analysis and visualizations. These files are provided as supplementary materials (Table S2-S4) and stored in the GitHub repository for this project (see *Data availability*).

Prior to further analysis, we removed two samples from the dataset that had <100 sequences after processing, which left 301 samples for subsequent analyses. We used the R decontam package (v1.6.0) (52) to identify and remove contaminant ASVs based on their prevalence in control samples (DNA extracted from sterile swabs) compared to biological samples. A total of four bacterial ASVs (ASV276 Micrococcaceae, ASV1412 Planococcaceae, ASV1797 Delftia, ASV1979 Stenotrophomonas) were present in at least 50% of control samples at relative abundances >1%. The 4 ASVs had decontamination scores below our specified threshold (0.5) (Fig S2) and were thus filtered from our dataset. This left a total of 1,974 unique ASVs for analysis.

### Statistical analysis of 16S rRNA gene profiles: taxonomic composition, and alpha- and beta-diversity

In this study, we examined variation in the taxonomic composition, alpha-diversity, and beta-diversity of the hyena’s gut microbiome. Unless otherwise stated, all statistical analyses and figures were made in R (v3.6.2) (49). We first visualized taxonomic variation through stacked bar plots using the ggplot2 (v3.3.3) package (53). The plots showed the relative abundances of dominant bacterial phyla, orders, and genera across samples over the study period (N=301; 1993-2016). Phyla with mean relative abundances >1%, orders with mean relative abundances > 0.6%, and genera with mean relative abundances >3% across samples were represented and all others were clumped into an ‘Other’ category.

For microbiome alpha-diversity analyses, samples were first subsampled to 2900 reads per sample using mothur (v1.42.3) (54) to control for uneven sequencing depths. Two samples did not meet this read number cutoff and were excluded from alpha-diversity analyses (N=301 to N=299). Rarefaction curves of ASV richness reached saturation, indicating that sequencing depth was sufficient for analyzing these communities (Fig S1B). Microbiome alpha-diversity was estimated using the Chao1 richness, Shannon diversity, and Faith’s phylogenetic diversity (PD) (55–57) indices, which respectively captured microbiome richness, evenness, and phylogenetic taxonomic representation. The phyloseq package (v1.30.0) (58) calculated the values for the first two metrics. The picante package (v1.8.2) (59) estimated the latter metric, after supplying it with a phylogenetic tree of bacterial ASVs constructed with DECIPHER (v2.14.0) (60) and phangorn (v2.5.5) (61).

To evaluate whether host individual identity, age, matriline, average monthly prey abundance, and calendar year predicted gut microbiome alpha-diversity (Chao 1, Shannon, or PD indices on the log scale), we ran generalized linear models using the glm function from the stats package (v3.6.2) (49). After assessing model fit from residuals, we tested for statistical significance (α=0.05) by conducting likelihood ratio tests on all linear models using the car package (v3.0-10) (62). A second set of linear mixed models specified hyena identity and sample year as random factors and evaluated the influences of the remaining variables on gut microbiome alpha-diversity. The linear models were made with the lme4 package (v1.1-26) (63) and statistical significance was calculated as described above. Significant associations between a host factor and gut microbiome alpha-diversity were visualized via scatterplots and boxplots in ggplot2 (v3.3.3).

Microbiome beta-diversity was quantified using Jaccard distances calculated from bacterial ASV presence/absence data, and Bray-Curtis distances calculated from bacterial ASV relative abundance data, after excluding ASVs with ≤ 2 total reads in the dataset. We also estimated weighted Unifrac distances which consider the phylogenetic relationships among bacterial ASVs. Jaccard and Bray-Curtis distances were estimated using the vegan package (v2.5.7) (64), while Unifrac distances were generated using phyloseq (v1.30.0). To determine whether gut microbiome beta-diversity was predicted by five host factors (individual identity, age, matriline, average monthly prey abundance, and year), we performed permutational multivariate analyses of variance (PERMANOVA) tests with vegan (N=299). The tests specified one type of distance matrix as the dependent variable, the five host predictors as the independent variables, and 999 permutations. PCoA ordinations based on Bray-Curtis distances were constructed in ggplot2 and were color-coded by each of the host predictors.

### Statistical analysis of 16S rRNA gene profiles: core gut microbiome

We used the 16S rRNA gene data to identify taxa that constituted the core gut microbiome in wild spotted hyenas. For this, we identified the bacterial genera and ASVs that were present in >85% of samples at mean relative abundances of at least 0.5%. For ASVs when genera was unknown, the next most refined level of known taxonomic classification was used (e.g., Family). Heatmaps made in ggplot2 showcased the relative abundances of core bacterial genera or ASVs for each sample. A bar graph illustrated the proportion of the gut microbiome community that was represented by the core bacterial taxa in each sample.

To determine whether the relative abundances of core bacterial genera or ASVs varied with host age, matriline, average monthly prey abundance, or calendar year, we constructed linear mixed models with the lme4 package. Host identity was set as a random effect, and only bacterial taxa with mean relative abundances of at least 1% were tested. Statistical significance of each model term was assessed by calculating p-values using the Satterthwaite approximation with lmerTest (v.3.1-3) (65) and applying a false discovery rate (FDR) correction. The beta coefficients of statistically significant terms were plotted in ggplot2.

### Sequencing and processing of metagenomic reads

To gain insight into the genomic diversity and functional potential of the gut microbiome in wild hyenas we submitted a subset of fecal samples (N=32) from two mother-daughter pairs (eight samples/hyena) for paired-end shotgun metagenomic sequencing (Table S2). These four hyenas belonged to matrilines 1 (high rank) and 3 (med-low rank), and their samples spanned two two-year periods: 2000-2001 for samples from the mothers and 2013-2015 for samples from the daughters. The samples were sequenced on the Illumina HiSeq 4000 platform at the Michigan State University Genomics Core (150+150 bp). Libraries were prepared using the Rubicon ThruPLEX DNA-Seq Library Preparation Kit following manufacturer’s recommendations. Base calling was done by Illumina Real Time Analysis (RTA; v2.7.7) and output of RTA was demultiplexed and converted to FastQ format with Illumina Bcl2fastq (v2.19.1).

On average, samples yielded ~20 million paired-end reads (range: 14-23 million) with high-quality phred scores (28–30). Trimmomatic (v0.38) (66) was used to remove sequence adapters and low-quality bases from raw reads using the program’s default parameters. After this filtering, samples had an average of 16,374,385 sequences (± 3,049,668). Host DNA was removed by mapping sample reads to the hyena genome (67) using the graph-based aligner HISAT2 (68). Next, the forward and reverse reads for each sample were interleaved using the interleave-fastq script from the Ray assembler (v2.3.1) (69). Kraken2 (v2.1.0) was used to assign taxonomic labels to interleaved reads for each sample (70).

Interleaved reads from all samples were concatenated into a single file and assembled into contigs using Megahit (v1.2.9) (71) with default parameters. A total of 2,742,876 contigs were generated and the quality of the assembly was evaluated using the Quality Assessment Tool for Genome Assemblies (QUAST) (v5.0.0) (72) (Table S5). To functionally annotate metagenomes, contigs were imported into Anvi’o (v.6.2) (73). Anvi’o predicted a total of 7,775,878 gene open-reading frames (ORFs) using Prodigal (74). The program assigned functional annotation to genes using the Clusters of Orthologous Groups (COGs) (75), and the Kyoto Encyclopedia of Genes and Genomes (KEGG) (76) databases. To obtain an estimate of the relative abundance of each gene in a sample, quality-filtered sequences from each sample were mapped to ORFs using Salmon (v1.8.0) (77). Salmon calculated the relative abundances of ORFs in units of Transcripts Per Million (TPM), which normalizes for gene length and sample sequencing depth. Tables of the relative abundances of COG pathways and KEGG proteins appear in the supplementary materials (Table S6-S8).

Contigs with a minimum length of 1000 bp were binned into metagenome-assembled genomes (MAGs) using MetaBat2 (v2.15) (78). Of the MAGs generated, 149 high-quality MAGs were obtained with completeness scores > 80% and contamination scores <5% as assessed by CheckM (v1.1.3) (79) (Table S9). MAGs were assigned a taxonomy using the Genome Taxonomy Database Toolkit (GTDB-Tk) (v1.3.0) (80) with the GTDB taxonomy release 95 (Table S9). A phylogenetic tree of MAGs was built using the multiple sequence alignment file generated by GTDB-Tk and used the taxonomic assignments as input to RAxML (81) for refining the phylogeny. The final tree was visualized using the interactive Tree Of Life (iTOL; v6) (82). Individual trees of each MAG were also constructed in R to visualize the evolutionary distances between each MAG and genomes in GTDBr-95. The R package treeio (83) subsetted the large phylogeny outputted by GTDB-Tk and ggtree visualized the tree (83).

Lastly, the relative abundance of each MAG in a sample was estimated using CoverM (v0.6.1) (https://github.com/wwood/CoverM) by mapping quality-filtered reads to each MAG. For every sample, CoverM output the percentage relative abundance of each MAG, as well as the percentage of unmapped reads (Table S10). On average, ~13.20% ± 7.68% of reads in any given sample (after filtering out host DNA) mapped to the MAGs (range= 1.19% - 27.42% of reads).

### Statistical analysis of metagenomic data

Gut metagenome taxonomic profiles were visualized via stacked bar plots using ggplot2, which showed the relative abundance of metagenomic reads assigned to each bacterial phylum, order, and genus as determined by Kraken2.

The predicted genes that were annotated in Anvi’o coded for 25 broad COG Categories, 67 more specific COG Pathways, and 7313 unique KEGG proteins. The relative abundances (in Transcripts Per Million - TPM) of these functions are in Tables S6-S8. TPM abundances were converted to proportions (e.g., relative abundances). We constructed Bray-Curtis distances from the relative abundances of COG and KEGG functions and ran PERMANOVA tests to examine whether these were associated with three host predictors: individual identity, matriline, or average monthly prey abundance. Sample year and hyena age were not included since the samples only spanned a total of two two-year periods. We followed the methods described previously in the section “*Statistical analysis of 16S rRNA gene profiles: alpha- and beta-diversity*.” The functional categories with the highest relative abundances across samples were visualized via stacked bar plots in ggplot2.

Lastly, we conducted PERMANOVA statistics on the relative abundances of 149 high-quality MAGs to determine whether these were associated with host individual identity, matriline, or mean monthly prey abundance. Tests were based on Bray-Curtis distances and used 999 permutations. The clustering of samples based on their MAG relative abundances was visualized as a PCoA ordination.

### Data availability

Raw sequence files were deposited in NCBI’s Sequence Read Archive, under BioProject PRJNA733503 and accession numbers SAMN19468262 - SAMN19468578 (for 16S rRNA amplicon reads) and BioProject PRJNA734005 and accession numbers SAMN19814909 - SAMN19814940 (for shotgun metagenomic reads). All data files and R scripts for the statistical analyses and figures included in this manuscript are available on the GitHub repository for this project (https://github.com/rojascon/HyenaGutMicrobiome_AcrossGenerations).

### Ethical approval

Our research procedures were approved by the MSU IACUC on January 8, 2020 (approval no. PROTO201900126) and comply with the ethical standards set by Michigan State University, the American Society of Mammalogists (70), the Kenya Wildlife Service, the Kenyan National Commission on Science, Technology and Innovation, and the Mara Conservancy.

## Results

### Global shifts in the composition of the gut microbiome in wild hyenas sampled across 23 years

We first investigated variation in 16S rRNA gene profiles over the 23-year study period and evaluated whether any of the variation present could be explained by host endogenous and/or exogenous factors. Across individuals, we found that gut microbiomes showed high interindividual, interannual, and population-wide shifts in composition. The relative abundances of all five dominant bacterial phyla (*Firmicutes, Actinobacteria, Bacteroidetes*, and *Fusobacteria*) varied over the two decades of sampling (Fig 1A). Additionally, in 2008-2009, there was a marked shift in the gut microbiome compositions of all sampled hyenas. Gut microbiomes which were previously dominated by *Firmicutes* (mainly *Clostridiales*) and *Actinobacteria* (mainly *Bacillales*), came to be dominated by *Firmicutes*, *Bacteroidetes* (*Bacteroidales*) and *Fusobacteria (Fusobacteriales*) (Fig 1A-B). *Bacteroidales* and *Fusobacteriales*, which had a combined mean relative abundance of <4% before 2008, became much higher in relative abundance after 2008 and constituted on average 17.27% and 11.24% of the microbiome, respectively (Figure 1B). In turn, *Bacillales* mean relative abundance decreased from 10.66% prior to 2008 to 1.37% over the remaining years. At the genus level, the shift appeared to be driven by increases in the relative abundances of *Fusobacterium, Bacteroides*, and *Peptoclostridium*, and by decreases in the relative abundances of *Enterococccus*, unclassified *Planococcaceae* and unclassified *Clostridiales* during the years following 2008 (Fig 1C). Thus, it is evident that the composition of the gut microbiome in wild adult hyenas varies greatly across the two decades of sampling and marked shifts occurred in 2008-2009 that were apparent in all sampled hyenas.

**Figure 1.**
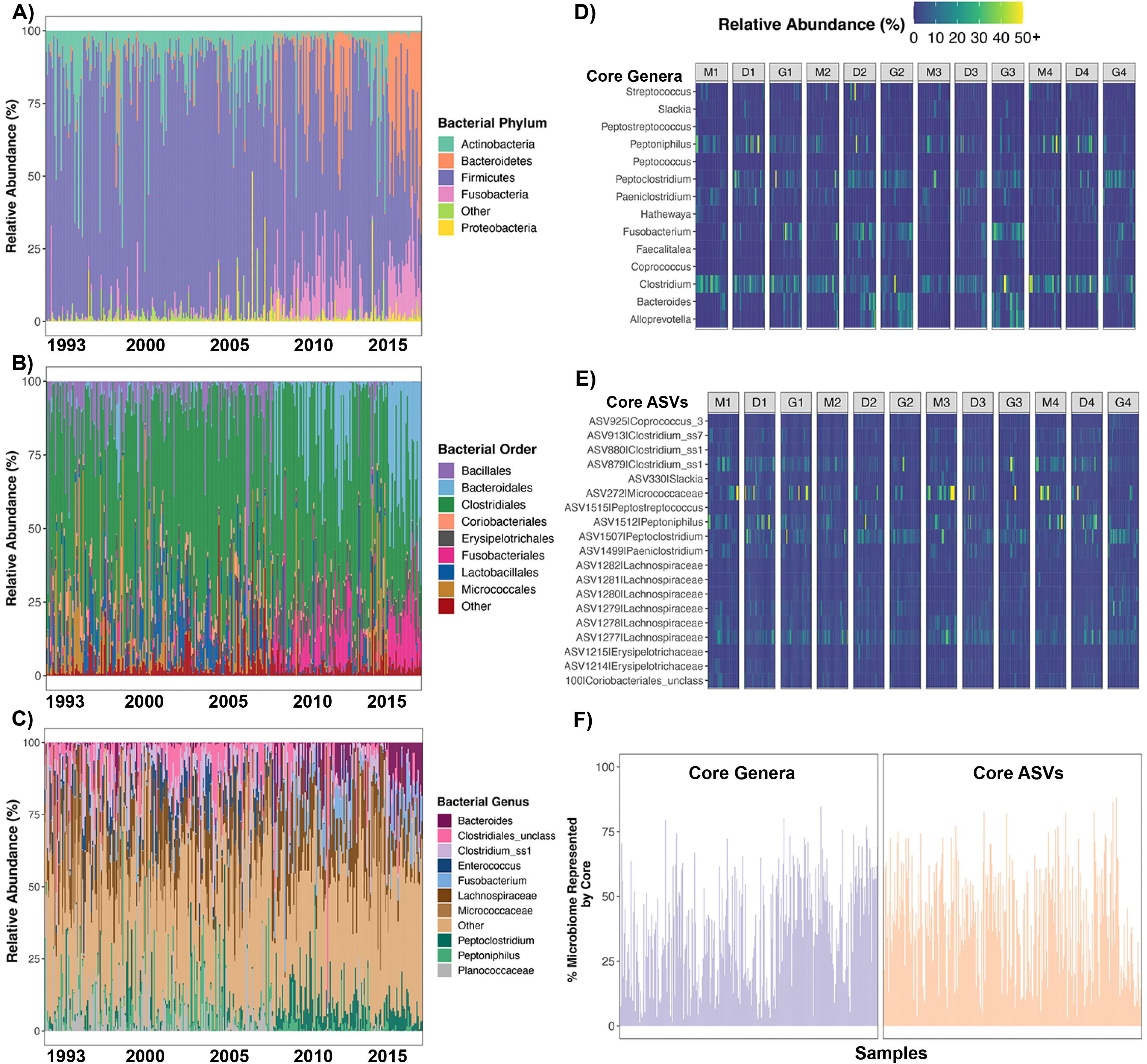
Amidst global and temporal shifts in gut microbiome composition, a taxonomic core is present in the guts of all studied hyenas. Stacked bar plots showing the relative frequency of 16S rRNA gene sequences assigned to each bacterial phylum (**A**), order (**B**), and genus (**C**) across samples. Samples are ordered by sampling date, and each color represents a bacterial phylum, order, or genus. **D-E**) Heatmap of the relative abundances of 14 core bacterial genera (**D**) or 19 core bacterial ASVs (**E**) across samples. These bacterial taxa were found in 85% of samples. Not all sequences were classified to genera or species-level and in those scenarios, their last known classification (e.g., Family) was used. **F)** Proportion of the microbiome that is represented by core genera (purple) or ASVs (orange) in each sample.

Despite the high degree of temporal variability observed in 16S rRNA gene profiles, there were 14 bacterial genera out of 326 genera that represented the ‘core’ and were present in at least 85% of gut microbiome samples (Table 2, Fig 1D). Core bacterial genera included: *Alloprevotella, Bacteroides, Clostridium, Fusobacterium, Paeniclostridium, Peptoclostridium, Peptoniphilus* and *Streptococcus*, among others (Table 2, Fig 1D). All but five of those genera were found at mean relative abundances > 1% across samples. These 14 core genera collectively constituted ~36% (± 20.2%) of the gut bacterial community in any given sample (Table 2, Fig 1F), although there were several samples throughout the study period that lacked most of the core taxa. Furthermore, 19 out of 1689 total bacterial ASVs were present in over 85% of gut microbiome samples (Table 2, Fig 1E). Eight of the 19 core ASVs were assigned a genus and for the remaining ASVs, no information beyond family was known (Table 2, Fig 1E). Ten of the 19 core ASVs were found at mean relative abundances >1% across samples. These 19 core ASVs collectively constituted ~40% (± 20.1%) of the gut microbiome (Fig 1F). Again, there were several samples from certain years that did not contain many of the core ASVs. Interestingly, although the collective relative abundance of core microbial taxa was very low in some samples (2-4%), it was never zero. Overall, these results indicate that a core microbiome is present in hyenas, but as was observed in the broader 16S rRNA gene profiles, the relative abundance of this core microbiome is labile.

**Table 2.**
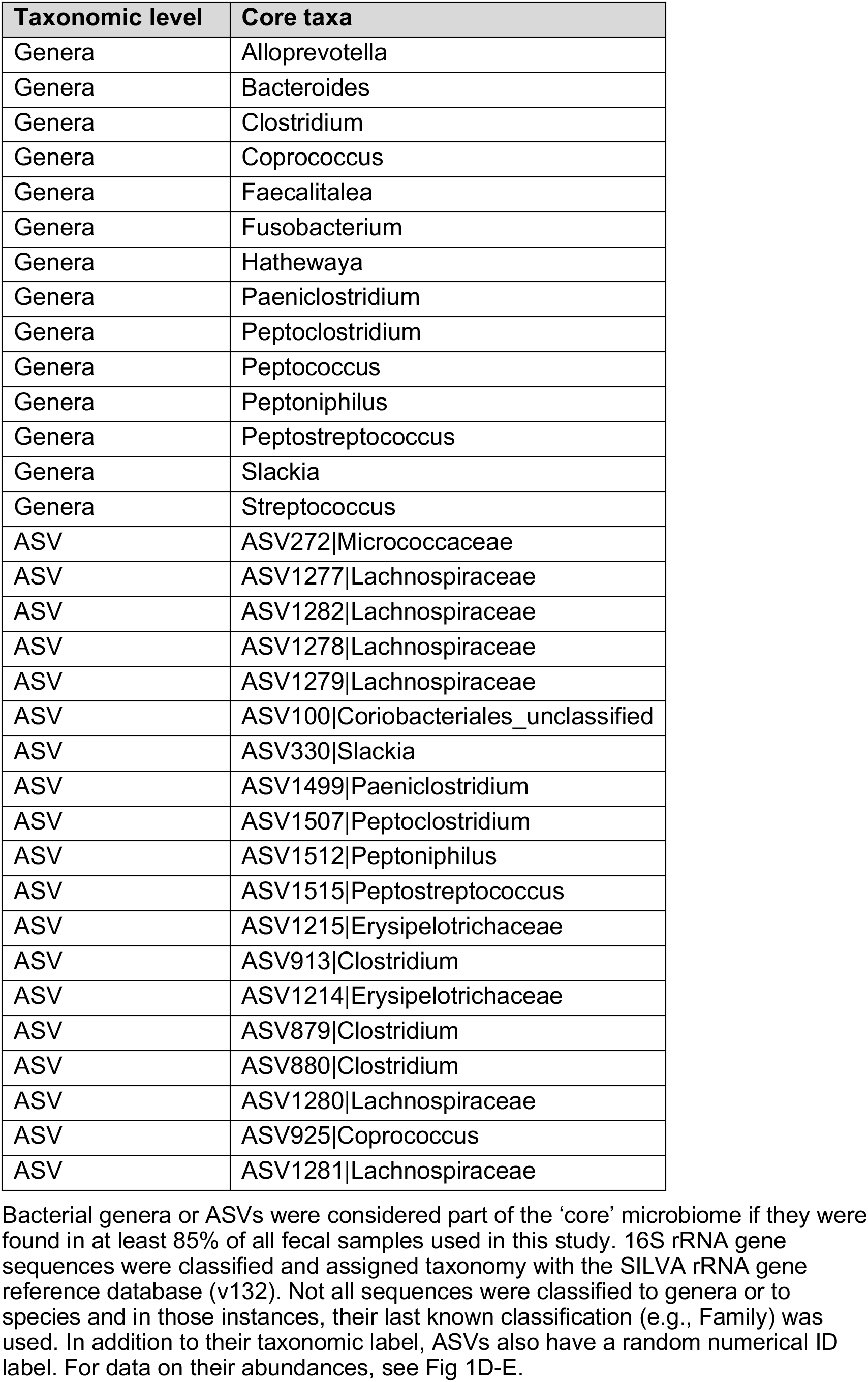
Core bacterial genera and ASVs that are present in >85% of fecal samples from spotted hyenas.

### Host socioecological factors predict distinct aspects of the gut microbiome

We also examined whether the alpha- and beta-diversity based on 16S rRNA gene profiles were associated with host endogenous and exogenous factors including individual identity, matriline, age, mean monthly prey abundance, and sample year. Individual identity most strongly correlated with gut microbiome alpha-diversity, including richness, evenness and phylogenetic diversity (GLM LRT p<0.05; Fig 2A, Table S11). Additionally, mean monthly prey relative abundance was negatively correlated with gut microbiome richness; gut microbiomes were modestly less diverse during months of high prey availability than during periods of prey scarcity (GLM LRT p=0.019; Fig 2C, Table S11). Host maternal lineage and calendar year were not significantly associated with gut microbiome alpha-diversity (GLM LRT p>0.05; Table S11). We also re-evaluated the effects of the host predictors after accounting for the repeated sampling of individuals (i.e., hyena identity) and temporal variation (i.e., sample year). In these models, gut microbiome richness and phylogenetic diversity varied with host age; gut microbiome diversity tended to be lower in older than in younger adult hyenas (GLM LRT p<0.05; Fig 2B, Table S12)). Furthermore, as was observed in the earlier models, mean monthly prey abundance was negatively correlated with gut microbiome richness (Table S12).

**Figure 2.**
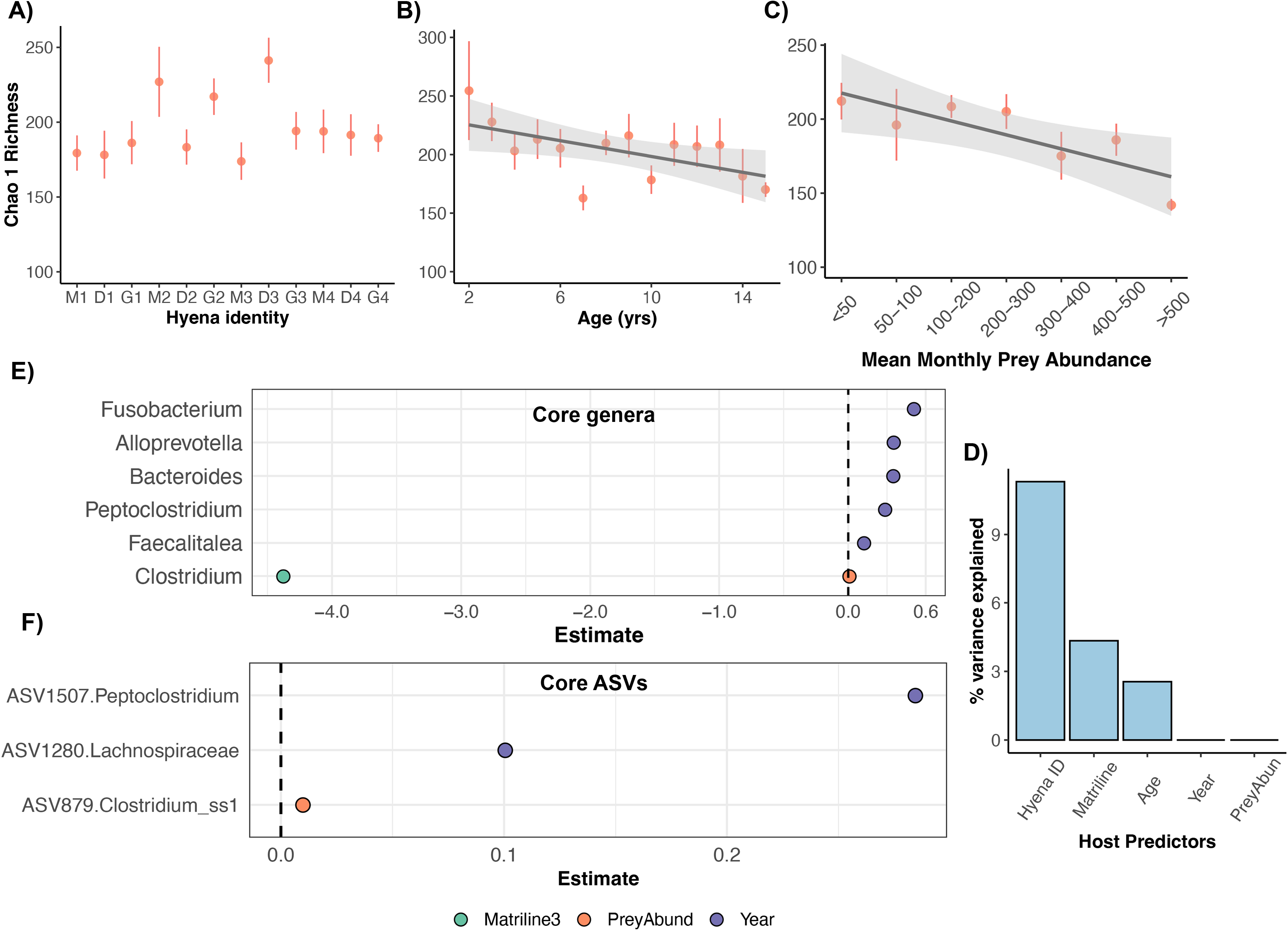
Socioecological predictors of the gut microbiome in wild spotted hyenas. Plots of microbiome Chao 1 Richness (mean ± SE) for each hyena individual (**A**), age category (**B**), or mean monthly prey abundance (**C**) for 16S rRNA gene data. The shaded lines with a 95% CI represent the relationship between x and y estimated as a linear regression. See Table S11-S12 for statistical output from the linear models. **D)** Plots of R^2^ values from PERMANOVA models testing whether host individual identity, matriline, age, mean monthly prey abundance and sample year predicted gut microbiome beta-diversity (for 16S rRNA gene data). See Table 2 for exact model output. **E-F**) Plots of beta-coefficients from linear mixed models regressing the abundance of core genera (**E**) or ASVs (**F**) against the five host predictors listed above. Each point represents a beta-coefficient and is color coded by host predictor. If a beta-coefficient is positive, the abundances of that bacterial taxon were positively correlated with the host predictor. Only beta-coefficients with p-values <0.05 after correcting for multiple comparisons are displayed.

When examining gut microbiome beta-diversity, results showed that hyena identity accounted for up to 11.3% of the variance (Weighted Unifrac PERMANOVA; Table 3, Fig 2D), suggesting that gut microbiomes are individualized to some extent, and may be consistent over an adult’s lifespan. Hyena age and matriline accounted for an additional 4.3% and 2.5% of the variation, respectively (Table 3, Fig 2D). Mean monthly prey abundance, and sample year explained little to none of the variation in microbiome beta-diversity (Table 3, Fig 2D). In PCoA ordination plots, gut microbiome profiles cluster moderately by sample year and do not appear to cluster by the remaining predictors that were evaluated (Fig S3).

**Table 3.** Predictors of gut microbiome beta-diversity in adult female hyenas (N=301).

| Predictor | Jaccard<br>(% variance<br>explained) | Bray-Curtis<br>(% variance<br>explained) | Weighted Unifrac<br>(% variance<br>explained) | Average<br>(% variance) |
| --- | --- | --- | --- | --- |
| hyena identity<br>(categorical) | 6.13 (p=0.001) | 8.75 (p=0.001) | 11.31 (p=0.001) | 8.73 |
| matriline<br>(categorical) | 2.01 (p=0.001) | 2.23 (p=0.001) | 2.55 (p=0.001) | 2.26 |
| age (yrs) | 1.64 (p=0.001) | 2.41 (p=0.001) | 4.34 (p=0.002) | 2.8 |
| prey monthly | 0.57 (p=0.006) | 0.44 (p=0.10) | 0.27 (p=0.40) | <1% |
| year | 0.64 (p=0.001) | 0.52 (p=0.026) | 0.81 (p=0.11) | <1% |
| Residuals | 88 | 85 | 81 |  |

Shown are the $R^2$ values (% variance explained) and p-values for marginal PERMANOVA tests that evaluated whether hyena endogenous and exogenous factors predicted gut microbiome similarity (based on ASV abundances calculated from 16S rRNA gene data). The models utilized 3 types of distance matrices and included five host factors (hyena identity, matriline, age, mean monthly prey abundance, year).

We also ran linear mixed models to determine whether any of the host factors correlated with the relative abundances of core bacterial genera or ASVs. The models accounted for variation attributable to host individual identity and tested core taxa that had mean relative abundances of at least 1%. The relative abundances of five core genera varied with sample year and increased over time (LMM p.adj <0.05; Table S13). These genera were *Alloprevotella, Bacteroides, Peptoclostridium, Fusobacterium*, and *Faecalitalea* (Fig 2E). *Clostridium* relative abundances were marginally positively correlated with mean monthly prey availability (Fig 2E). These genera appeared to have higher relative abundances in the hyena gut during periods of high prey availability than during periods of prey scarcity. No bacterial genera varied with host age and only *Clostridium* abundances varied with maternal lineage (Table S13). Hyenas belonging to matriline 3 harbored lesser abundances of this genera compared to hyenas from the highest-ranking matriline. Lastly, when conducting a similar analysis on the relative abundances of core ASVs, we found that two ASVs were temporally variable, one ASVs was weakly predicted by prey abundance, and none were associated with host age or maternal lineage (Fig 2F, Table S13).

Collectively, our results indicate that individual signatures in gut microbiome alpha- and beta-diversity are observed in hyenas. Gut microbiome profiles also varied temporally, suggesting that they are variable within hosts across lifespan and across generations. Prey abundance was not significantly associated with much variation in 16S rRNA profiles, yet differences may exist in microbiome functional profiles which we analyze below.

### Gut metagenome taxonomic profiles across individuals

We characterized the gut metagenomes of two mother-daughter hyena pairs using shotgun metagenomic sequencing (Table S2). The pairs belonged to matrilines 1 and 3, respectively, and samples from each individual hyena spanned two years (N=8 samples per individual). Putative sequences were classified and assigned taxonomy using Kraken2. Metagenomic sequences were overwhelmingly classified as Bacteria (88% mean relative abundance), and a smaller fraction to Viruses and Archaea (combined mean relative abundance of 0.56%) (Fig 3C).

**Figure 3.**
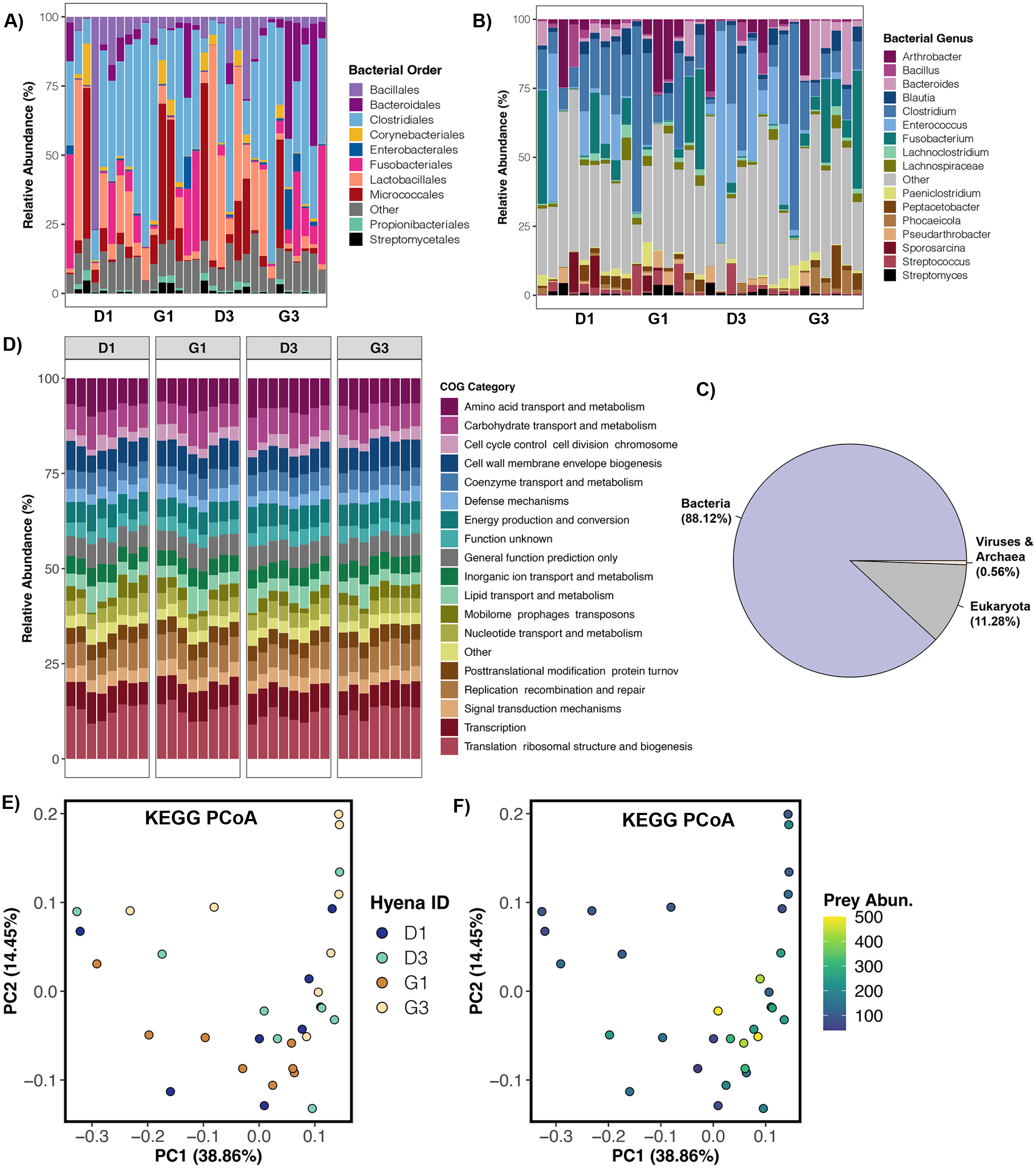
Composition and predicted function of gut metagenomes obtained from shotgun metagenomic data. Stacked bar plots showing the relative frequency of shotgun metagenome sequences assigned to each bacterial order (**A**) and genera (**B**) as determined by Kraken2. Samples are organized by hyena individual. **C)** Proportion of metagenome sequences assigned to Bacteria, Eukaryota, and Archaea & Viruses as calculated by Kraken2. **D)** Relative abundances of the most represented COG functional categories across samples. **E-F)** PCoA ordinations based on KEGG relative abundances color-coded by host individual identity (**E**) or mean monthly prey abundance (**F**). Contigs were assembled from metagenomic data and imported into Anvi’o for gene prediction and functional annotation. Salmon calculated the relative abundance of genes in each sample (in TPM), and these values were converted to proportions (e.g., relative abundances).

We examined the Kraken2 results for the classifications within Bacteria in more detail. When focusing solely on Bacteria, *Firmicutes* (57.9% mean relative abundance across samples) was the dominant phylum, followed by *Actinobacteria* (19.4%), *Fusobacteria* (8.4%), *Bacteroidetes* (8.01%), and *Proteobacteria* (5.09%) (Fig S4). All other phyla appeared at mean relative abundances <1%. The bacterial orders with the highest relative abundances were *Clostridiales* (34.41% mean relative abundance), *Lactobacillales* (13.92%), *Micrococcales* (11.68%), *Fusobacteriales* (8.41%), *Bacillales* (8%) and *Bacteroidales* (6.77%), although the relative abundances of these bacterial orders varied among host matrilines (Fig 3A). Hyenas from the lower-ranking matriline (M3) appeared to contain greater relative abundances of *Bacteroidales, Lactobacillales*, and *Enterobacteriales* than hyenas from the higher-ranking matriline (M1), whose gut microbiomes instead contained more *Clostridiales* and *Fusobacteriales* (Fig 3A). At the genus level, these gut metagenomes mostly contained *Clostridium* (17.88% mean relative abundance), *Enterococcus* (10.64%), *Fusobacterium* (8.29%), *Arthrobacter* (5.28%), and *Bacteroides* (3.38%) (Fig 3B).

### Housekeeping bacterial functions are the most represented in the gut microbiomes of wild spotted hyenas

The third aim of this study was to examine the gut microbiome functions in hyenas and determine whether these were correlated with host socioecology (individual identity, matriline, or prey density). The functional databases used to annotate predicted genes in the metagenomic data were the Clusters of Orthologous Groups (COGs), and the Kyoto Encyclopedia of Genes and Genomes (KEGG).

Not surprisingly, the COG Categories with the highest relative abundances were housekeeping functions that are essential for bacterial growth and replication. The categories were ribosomal structure and biogenesis, amino acid transport and metabolism, carbohydrate transport and metabolism, transcription, cell membrane biogenesis, and DNA recombination and repair (Fig 3D, Table S6). COG pathways, which are less broad than COG categories, were involved in the synthesis of the bacterial ribosome (50S or 30S subunits), aminoacyl tRNA synthetases (which attach amino acids to tRNA), fatty acids, purines, the amino acid lysine (for protein and cell wall synthesis), and peptidoglycan (cell wall component) (Table S7). Other COG pathways were involved in tRNA modification, glycolysis, and pyrimidine salvage. Similarly, in KEGG profiles, housekeeping functions were the most abundant functions across samples (Table S8).

We tested whether the relative abundances of COG categories, COG pathways, or KEGG proteins were associated with host identity, matriline, or prey densities. Overall, gut microbiome functions were individual-specific, as host identity explained ~13% of the variation (Fig 3E, Table S14; KEGG protein relative abundances only). Gut microbiome functional profiles were moderately correlated with prey abundance, and this factor explained ~6% of the variation (Fig 3F, Table S14; KEGG protein relative abundances only). Host matriline explained none of the variation in metabolic functions. These findings suggest gut microbiome functions as a collective may be influenced by host prey densities and habitat, and exhibit individual-specific signatures. No functional pathways or enzymes directly related to the digestion of bone (e.g., phosphatases, collagenases) were identified, although several COG pathways were involved in the synthesis of fatty acids from the fermentation of protein.

### Metagenome-assembled genomes shed light on previously uncharacterized bacterial genomic diversity in the hyena gut

Lastly, to further characterize the bacterial genomic diversity present in the hyena gut, we reconstructed a total of 149 high-quality, low-contamination metagenome-assembled genomes (MAGs) from metagenomic sequences (Fig 4A, Table S9). The average MAG completeness was 91.76% (± 5.37%) and the average contamination was 1.60% (± 1.24%) (Table S9, Fig S5A). Sizes of the MAGs ranged from 560 Kilobases (Kb) to 3.93 Megabases (Mb), with a median of 1.97 Mb (Fig S5B). The MAGs spanned 9 bacterial phyla, 12 classes, 25 orders, 47 families, and 51 genera.

**Figure 4.**
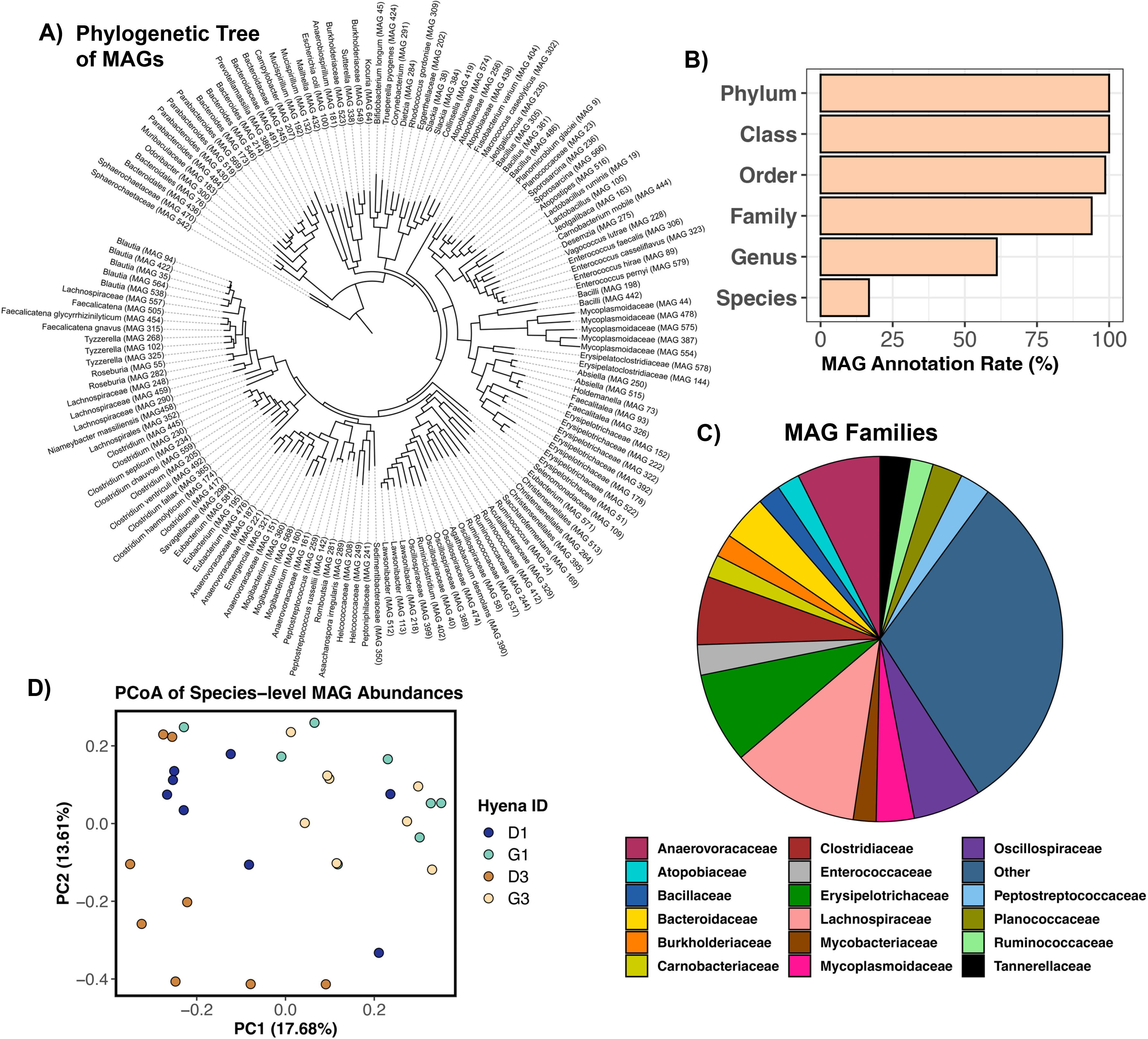
A total of 149 high-quality MAGs were recovered from the hyena gut, expanding the genomic diversity currently known for carnivores. Assembled contigs were binned into MAGs with MetaBat2 and assigned taxonomy with GTDB - release 95. Only high-quality MAGs (>80% completeness, <5% contamination) were retained. **A)** Phylogenetic tree of MAGs constructed from GTDB sequence alignments and labeled with GTDB taxonomy. **B)** % MAGs annotated at different taxonomic levels. **C)** Relative abundances of MAG families. **D)** PCoA showing the clustering of samples based on the abundances of MAGs that were classified to species (31 MAGs). See Fig S7 for PCoA based on the abundances of all 149 MAGs.

All high-quality MAGs were classified to Phylum and Class taxonomic levels and all but two MAGs were assigned an Order (Fig 4B). Seventy-three percent of MAGs were assigned to *Firmicutes*, 9.39% to *Bacteroidota* (formerly *Bacteroidetes*), and 8.72% to *Actinobacteriota* (formerly *Actinobacteria*). The most represented bacterial classes were *Clostridia* (45.6% of MAGs), *Bacilli* (27.5%), *Bacteroidia* (9.39%), and *Coriobacteriia* (4.69%) (Table S9). Dominant families represented among the MAGs included: *Lachnospiraceae, Erysipelotrichaceae, Anaerovoracaceae, Clostridiaceae, Oscillospiraceae* (Fig 4C). About 65% of our high-quality MAGs were assigned a Genus, and a total of 51 genera were identified (Fig 4B, Table S9). Five of the genera represented in the MAGs came from genera that are part of the core gut microbiome in wild spotted hyenas (this study). These genera were *Bacteroides, Clostridium, Faecalitalea, Fusobacterium*, and *Peptostreptococcus*. In addition, 12 of the MAGs came from genera that are very abundant in the gut microbiomes of captive hyenas from zoos in China (84), specifically the genera *Bacteroides, Blautia, Collinsella, Fusobacterium*, and *Peptostreptococcus*. Lastly, 20% of MAGs (31 / 149) were classified at the species level (Fig 4B, Table S9). Among the MAGs classified to species were ones assigned to *Ruminococcus gnavus* (# 315), *Lactobacillus ruminis* (# 19), *Enterococcus faecalis* (# 306), *Enterococcus casseliflavus* (# 323)*, Rhodococcus gordoniae* (# 309), *Planomicrobium glaciei* (# 9), and *Bifidobacterium longum* (# 45) (Fig 4A). Although some of the MAGs could be assigned to putative species, 80% of them could not because they are evolutionarily distant from the genomes in the GTDB database (Fig S6). This suggests that there is a significant amount of genomic novelty in these MAGs (and in the hyena gut microbiome) that has not been characterized anywhere.

Of the high-quality MAGs in our dataset, the one with the highest relative abundance across samples was classified as *Prevotellamassilia* sp. (# 386) (Table S10), a taxon that is closely related to an *Alloprevotella* isolated from jackal guts (*Canis mesomelas*) (85) and a *Prevotella* isolated from the canine GI tract (86) (Fig S6). A lactating, 8 year-old hyena from matriline 3 (G3) harbored the highest relative abundance of this MAG, with over 10% of their metagenomic reads mapping to this particular MAG (Table S10). This same hyena a year later, harbored the 3rd highest relative abundance of this MAG (7.6%). This hyena’s grandmother - pregnant at the time - harbored the 2nd highest relative abundance of this MAG (6.4%). The abundance of this MAG in the remaining samples was 2% or less (Table S10).

Several MAGs were assigned to families (e.g., *Clostridiaceae* and *Peptostreptococcaceae*) that have been reported to be more abundant in carnivores such as lions (*Panthera leo*), cheetahs (*Acinonyx jubatus*), and African wild dogs (*Lycaon picus*) compared to ruminants or other herbivores (87). In addition, several MAGs came from families (e.g., *Clostridiaceae, Erysipelotrichaceae* and *Bacteroidaceae*) reported to be highly correlated with protein and fat digestibility in domestic dogs (*Canis familiarus domesticus*) (88). One MAG (# 142), taxonomically assigned as *Peptostreptococcus russellii* came from a genus of bacteria that formed part of the core microbiome in our surveyed hyenas. This MAG was closely related to a *P. russellii* isolated from a swine manure storage pit (89)(Fig S6). Interestingly, *P. russellii* sequenced from human GI tract appears to suppress inflammation in mice (*Mus musculus*) (90).

There was one high-quality MAG, # 306 (taxonomically assigned as from *Enterococcus faecalis*), that was rare in the dataset (0-0.3% relative abundance in 96% of samples) but constituted 10% of metagenomic reads in a single sample from a low-ranking, pregnant, 7-yr old hyena (Table S10). The fecal sample was taken during a time when prey was scarce in the Reserve. The genome most closely related to this MAG was that of a pathogenic *Streptococcus* isolated from humans (Fig S6) (91). Perhaps this hyena was suffering from a GI infection at the time of sampling. This same hyena also had the highest relative abundance of an *Enterococcus hirae* MAG (# 89) and a *Vagococcus lutrae* MAG (#228), both of which were rare in all other samples (Table S10). MAGs assigned to *E. hirae* bacterial species were also recovered from the GI microbiome of healthy dogs (86).

Other MAGs we would like to call attention to include two that were assigned to the *Spirochetes* phylum (#470 and #542). *Spirochetes* have attracted attention recently as they are found in the gut of a wide diversity of mammalian species but absent or in low abundance in urbanized human populations (92). Another two interesting MAGs are #132 and #192 which were both assigned to the genus *Mucispirillum* from the *Deferribacteraceae* group; a group that is generally known to be capable of reducing iron, manganese, and nitrate during anaerobic respiration (93). The MAGs were found at low relative abundances in all samples, but the *Mucispirillum* genera has been previously documented in the gut microbiomes of swine, rodents, birds, and canids (94–97), although its specific relevance to host functioning need to be investigated.

Lastly, we also determined whether the relative abundances of the high-quality MAGs were associated with host socioecological factors. MAG relative abundances were significantly correlated with host identity (PERMANOVA R^2^=0.12, p=0.004) but not with host matriline, or mean monthly prey availability (PERMANOVA matriline R^2^=0.024, p=0.46; prey R^2^=0.038, p=0.15). In a PCoA ordination, samples do not cluster in a meaningful manner (Fig S7). If we only examined the relative abundances of MAGs classified at the species level, host identity accounted for 14.68% of the variation (PERMANOVA R^2^=0.146, p=0.001), while the remaining predictors had no explanatory power (PERMANOVA matriline R^2^=0.029, P=0.44; prey R^2^=0.027, p=0.51). In this ordination plot, samples appear to cluster by hyena identity along PC Axis 1 (Fig 4D).

## Discussion

### Large scale changes in the hyena’s habitat are reflected in the gut microbiome

Our data showed that the gut microbiome of wild spotted hyenas was highly variable across host adult lifespan, multiple generations of hyenas, and the 23-year study period. The most extreme shift was observed in 2008-2009, when the gut microbiomes of all sampled hyenas changed fundamentally and this changed composition was retained for the remainder of the study period. What was observed were 5-fold increases in the relative abundances of *Bacteroidales (Bacteroides*) and *Fusobacteriales (Fusobacterium*), and 10-fold decreases in the relative abundances of *Bacillales* (*Clostridiaceae* and *Peptoclostridium*), compared to earlier years. This 2-year period happened to encompass one of the most severe droughts in recent Kenyan history. Other ecosystem-wide ecological changes also occurred to the hyena’s habitat during this time. Specifically, over the 23 years of sampling, the MMNR experienced significant declines in its herbivore and lion densities, and increases in livestock grazing and anthropogenic activity related to tourism (19, 27, 28). At the same time, given that hyenas are adept at adjusting to change, hyena group densities doubled. The size of the specific social group studied here went from 60 individuals in 1990 to 120 individuals in 2013 (19). Interestingly, the sharpest increases in livestock grazing and tourism occurred during the severe drought in 2008-2009 (27), suggesting that anthropogenic and ecological disturbance may have a lasting effect on the microbiomes of wild animals (98–103). These changes collectively may have altered the ecology of the region and affected hyena diet, physiology, and behavior, and subsequently their gut microbiome compositions. Thus, our findings show that the gut microbiome of individual hyenas may respond similarly to large-scale ecological changes. Nonetheless, the gut microbiomes of hyenas were also largely individualized outside of the synchronized large-scale shifts in composition.

### Gut microbiomes are individualized in wild spotted hyenas

In line with results from a recently published study on the gut microbiomes of wild baboons (*Papio* spp.) sampled over a 14-year period in Amboseli National Park (14), our findings show that each hyena exhibited largely individualized gut microbiome compositions over its adult lifespan. Gut microbiome alpha-diversity varied with host identity. This same factor accounted for 11% of the variance in gut microbiome beta-diversity, more than was accounted for by host age, matriline, year, or prey abundance. Similarly, gut microbiome functional profiles – the relative abundances of predicted genes and functional pathways - were individual-specific, and so were the relative abundances of 149 MAGs. These findings suggest that individual hyenas possess unique microbiome signatures. In many mammals, including rodents, canids, cervids, and primates, host identity is one of the primary predictors of the composition of the gut microbiome (9, 104, 105). In spotted hyenas and other mammals, individualized microbiomes could arise from individual differences in social rank, immune function, reproductive state, social interactions, quality of diet, stress responses, injury, and genetics (23, 106, 107). All of these variables may act individually or in concert to structure mammalian microbiomes.

We found that gut microbiome alpha-diversity moderately declined with adult age, and that host age predicted 4.3% of the variance in gut microbiome beta-diversity. This is consistent with prior studies conducted in chimpanzees (*Pan troglodytes*) (108), and spotted seals (*Phoca largha*) (109), which report a general decline in gut microbiome diversity with host aging, but is not consistent with studies conducted in humans (110), or yaks (*Bos grunniens*) (111). In spotted hyenas from the Serengeti (112), juveniles have slightly less diverse gut microbiomes than adults, which could be true of the population we studied, but we did not sample juveniles. Individuals undergo significant alterations to their lifestyle, diet, immune system, and overall physiology with aging, which likely impacts the gut and its microbiome diversity (113, 114). Older individuals may have reduced mobility, weakened immune strength, altered gut morphology, reduced metabolic health, or infections (115, 116). Nonetheless, it is unclear what specifically contributes to reducing microbiome diversity as the hyena ages. This necessitates further study given that in mammals, maintaining high gut microbiome alpha-diversities may be beneficial (117).

It was surprising that host matriline was not significantly associated with gut microbiome composition, alpha-diversity, beta-diversity, or metabolic function. Hyenas from the same matrilines did not have more similar microbiomes than hyenas from two different matrilines, which is unexpected given that hyenas from the same matriline occupy similar ranks in the clan’s hierarchy and are more genetically related than hyenas from other matrilines (20, 22). Furthermore, closely related hyenas typically spend more time together than do unrelated hyenas, and occupy similar physical spaces (20, 22). Thus, we expected greater convergence in the gut microbiomes of maternally related hyenas. In this species, it appears that gut microbiome changes are more closely tied to ecological changes than to host social and genetic factors. This plasticity and fluidity in the microbiome might be a consequence of the tremendous dietary and behavioral flexibility that hyenas exhibit; characteristics that allow them to efficiently adapt to change. Firstly, they are opportunistic foragers with a generalist diet, can consume different species and sizes of prey, and can eat multiple if not all parts of their prey (33). They can eat insects or scavenge when ungulate prey numbers are low. They live in fission-fusion societies (20, 24, 25), where the compositions of sub-groups of hyenas change throughout the day, allowing hyenas opportunities to interact with different members of the clan throughout the day (118). Furthermore, the dispersal of immigrant males among neighboring clans diversifies the gene pool in this population (119)–immigrant males sire the majority of cubs in the clans and because of this, overall clan relatedness is very low (120). Hyenas also exhibit great behavioral flexibility and can adjust their behavior to improve their fitness. When anthropogenic disturbance was increasing within and outside Reserve boundaries, hyenas modified their behavior to become more nocturnal and vigilant to avoid conflicts with humans (121). Perhaps the microbiome’s highly variable composition in this species is a consequence of hyenas’ behaviors and traits that allow them to thrive in any environment.

### Host diet is correlated with gut microbiome variation

Our data showed that 16S rRNA gene microbiome alpha-diversity were negatively correlated with monthly prey densities, and that metagenome functional profiles were moderately correlated with this variable. It is known from earlier work (34) that the diets of MMNR hyenas change with the arrival of enormous herds of migratory wildebeest (*Connochaetes taurinus*) and zebras (*Equus quagga*) from Serengeti; this represents a period of very high prey abundance lasting roughly four months each year. Hyenas of all social positions enjoy a food surplus during these months, as they have access to migratory prey in addition to the resident prey that are available year round (e.g., Thompsons’s gazelle (*Eudorcas thomsonii*), impala (*Aepyceros melampus*), warthogs (*Phacochoerus africanus*), and topi (*Damaliscus lunatus*) (32). Conversely, during periods of low prey-availability, antelope prey are more scarce, and more challenging to hunt, requiring larger energetic effort from hyenas (34). Because of this, hyenas tend to scavenge more during this time and are more likely to eat the less desirable parts of carcasses (e.g., bones, viscera). Thus, the prey species and quality of diet changes for hyenas throughout the year, which in theory could directly impact the gut microbes that are present, and the metabolic pathways that are upregulated or downregulated.

Functional shifts in the microbiome in response to host dietary changes are not new and have been documented for other mammals. In goat kids (*Capra aegagrus hircus*), a transition from milk to a solid diet regime is associated with linear increases in volatile fatty acid production and the upregulation of pathways involved in carbohydrate and protein metabolism (122). In captive sifakas (*Propithecus coquereli*), supplementing standard diets with diverse foliage blends versus with single plant species, was associated with greater concentrations of short-chain fatty acids (SCFAs) in the gut (123). At a larger scale across wild mammals, from rodents to carnivores and primates, several studies have found associations between specific microbial functions and diet type (herbivore, omnivore, piscivore, carnivore) (35, 124, 125).

### Core, abundant, or important microbes of the hyena gut

The core gut microbiome in hyenas was composed of 14 bacterial genera and 19 ASVs that represented 36% and 40% of the microbial community, respectively. Thus, sampled hyenas contained some of the same types of microbes, albeit at varying relative abundances. Core bacterial genera included: *Alloprevotella, Bacteroides, Clostridium, Fusobacterium, Paeniclostridium, Peptoclostridium, Peptoniphilus* and *Streptococcus*, among others. Several of these taxa include known commensals of the mammalian GI tract and all but three genera (*Clostridium, Paeniclostridium, and Streptococcus*) were among the top 10 most relatively abundant genera in the GI tract of captive spotted hyenas residing in zoos from the Shandong province in China (84). Interestingly, the gut microbiomes of the captive hyenas had elevated relative abundances of *Fusobacterium* (19.5% mean relative abundance - their study; 4.10% our study), and *Bacteroides* (8.4% their study; 3.1% our study), and harbored significantly less *Clostridium* (<1% mean relative abundance - their study; 8.6% our study) compared to the wild population of spotted hyenas in this study here. This could potentially be attributed to the differences in diet and habitat between the hyenas in the two studies: the captive hyenas were all fed the same diet of rabbit meat, chicken, and beef and resided in a captive environment rather than a wild one.

More detailed studies are required to determine if or how the core genera are potentially interacting with a hyena’s metabolism. We do know that *Bacteroides* forms part of the core microbiome or is highly abundant in the guts of wild cats (*Felis silvestris catus*), coyotes (*Canis latrans*), raccoon dogs (*Nyctereutes procyonoides*), black-backed jackals (*Canis mesomelas*), and myrmecophagous pangolins (*Manis javanica*) (126–129). Members of this bacterial genus are known to be functionally involved in the degradation of protein and are related to a high fat and protein-based diet (130). *Peptoclostridium* is abundant in the gut microbiomes of captive Amur tigers (*Panthera tigris altaica*) fed a raw meat diet (131). Not much is known about *Peptoniphilus* and its role in the guts of wild mammals, but members of this genus use peptone (protein hydrolysate) as a major energy source and produce butyric acid in the process (132). We suggest that future studies examine these genera to see if they are directly or indirectly involved in helping hyenas digest their prey.

A total of 149 high-quality metagenome-assembled genomes were recovered from hyena GI tracts; some of which were assigned genera that formed part of the core microbiome in our studied individuals. Other MAGs were classified to bacterial families or genera found in the GI tracts of other carnivores, including those of wild cats, wild dogs, and domestic dogs (87). Interestingly, 80% of our MAGs were novel, as they were not classified to species-level and were evolutionarily distant from the genomes in GTDB-r95. Similarly, a larger study found a great deal of genomic diversity in the gut microbiomes of animals from five vertebrate classes, Mammalia, Aves, Reptilia, Amphibia, and Actinopterygii (133). The authors produced 5,596 nonredundant MAGs, of which 26% were novel and lacked a species-level match in GTDB-r89 (133). Metagenomic studies are expanding the currently known genomic diversity of the mammalian gut.

### Relatively abundant functions of the gut microbiome

Our study analyses did not elucidate which gut microbiome metabolisms or genes might be functionally important for hyenas. We showed that, not surprisingly, the most abundant bacterial functions were housekeeping functions present in virtually all of bacteria. To identify the core microbial functions that characterize spotted hyenas, comparative studies are required, where the gut metagenomes of hyenas are analyzed within the context of the gut microbiome of other carnivores, herbivores, and omnivores. Only then will it become apparent which metabolic functions are up- or down-regulated in spotted hyenas compared to animals consuming a different diet (e.g., herbivores), or animals with a similar diet but with distinct lifestyles and societies (e.g., less gregarious carnivores). We encourage that follow-up studies shed light on the uniqueness of the functional repertoire of the gut microbiome in hyenas.

## Conclusions

Using longitudinal sampling across two decades, and multiple sequencing approaches, we found that the gut microbiomes of wild spotted hyenas were individualized and correlated with large scale changes in the hosts’ ecological environment. Gut microbiome 16S rRNA profiles and metagenomic functional profiles also varied with host prey density and likely, with host diet. We also recovered 149 high-quality MAGs from the hyena gut, greatly expanding the microbial genome diversity known for hyenas, and wild mammals in general. We encourage that future microbiome studies of wild, captive, or domestic mammals employ longitudinal sampling, metagenomic-based functional analyses, and reconstruct metagenome-assembled genomes from their data. Inclusion of these techniques will capture different aspects of gut microbiome variability and improve our understanding of host-microbe interactions in wild mammals.

## Supplementary material

**Supplementary Figures,** Fig S1-S7, PDF file, 3.9M.

**Supplementary Tables,** Table S1-S14, XLSX, 4.8M.

## Acknowledgements

We thank Mara Hyena Project research assistants for collecting behavioral data and fecal samples in the field and completing data entry and management in the laboratory. We furthermore thank Christina Koehler for extracting genomic DNA from fecal samples and Andrew Winters for providing guidance on lab work. We are indebted to the Kenya Wildlife Service, the Kenyan National Commission on Science, Technology and Innovation, the Kenyan National Environmental Management Authority, the Narok County Government, the Naboisho Conservancy, the Mara Conservancy, the senior warden of the Masai Mara, Brian Heath, and Christine Koshal for allowing us to conduct this research in our field site in the Masai Mara National Reserve. We are also grateful to the UNAM Institute of Ecology labs of Valeria Souza and Luis Eguiarte labs for hosting C.A.R. during a semester and guiding her throughout the bioinformatics processing and analysis of her metagenomes. We thank Dr. Vanja Klepac-Ceraj for providing valuable feedback on the manuscript. Lastly, we also thank the BEACON Center for the Study of Evolution in Action, the MSU Institute for Cyber Enabled Research (iCER) and their High Power Computer Center (HPCC) for providing technical support and the computational resources needed to complete this project.

## Author Contributions

K.R.T., K.E.H., and C.A.R. designed the study, Mara Hyena Project research technicians collected the samples and behavioral data. Our international collaborators, V.S. and M.V., along with J.A.E. assisted with the sequence processing and analysis of shotgun metagenomic data. C.A.R., K.R.T., and J.A.E analyzed and interpreted all data. C.A.R. wrote the manuscript. All authors approved its final version.

## References

1. Trosvik P, Muinck EJ De, Rueness EK, Fashing PJ, Beierschmitt EC, Callingham KR, Kraus JB, Trew TH, Moges A, Mekonnen A, Venkataraman V V, Nguyen N. 2018. Multilevel social structure and diet shape the gut microbiota of the gelada monkey, the only grazing primate. Microbiome 6:1–18.

2. Zhu L, Wu Q, Deng C, Zhang M, Zhang C, Chen H, Lu G, Wei F. 2018. Adaptive evolution to a high purine and fat diet of carnivorans revealed by gut microbiomes and host genomes. Environ Microbiol 20:1711–1722.

3. Daniel H, Gholami AM, Berry D, Desmarchelier C, Hahne H, Loh G, Mondot S, Lepage P, Rothballer M, Walker A, Böhm C, Wenning M, Wagner M, Blaut M, Schmitt-Kopplin P, Kuster B, Haller D, Clavel T. 2014. High-fat diet alters gut microbiota physiology in mice. ISME J 8:295–308.

4. Miyazaki T, Nishimura T, Yamashita T, Miyazaki M. 2018. Olfactory discrimination of anal sac secretions in the domestic cat and the chemical profiles of the volatile compounds. J Ethol 36:99–105.

5. Theis KR, Venkataraman A, Dycus JA, Koonter KD, Schmitt-Matzen EN, Wagner AP, Holekamp KE, Schmidt TM, Greenberg EP. 2013. Symbiotic bacteria appear to mediate hyena social odors. PNAS 110:19832–19837.

6. Gaona O, Cerqueda-García D, Moya A, Neri-Barrios X, Falcón LI. 2020. Geographical separation and physiology drive differentiation of microbial communities of two discrete populations of the bat Leptonycteris yerbabuenae. Microbiol Open 00:1–15.

7. Langille MGI, Meehan CJ, Koenig JE, Dhanani AS, Rose RA, Howlett SE, Beiko RG. 2014. Microbial shifts in the aging mouse gut. Microbiome 2:1–12.

8. Sabey K, Song S, Jolles A, Knight R, Ezenwa V. 2020. Coinfection and infection duration shape how pathogens affect the African buffalo gut microbiota. ISME J.

9. Baniel A, Amato KR, Beehner JC, Bergman TJ, Mercer A, Perlman RF, Petrullo L, Reitsema L, Sams S, Lu A, Snyder-Mackler N. 2020. Seasonal shifts in the gut microbiome indicate plastic responses to diet in wild geladas. Microbiome 1–20.

10. Perofsky AC, Lewis RJ, Abondano LA, Di Fiore A, Meyers LA. 2017. Hierarchical social networks shape gut microbial composition in wild Verreaux’s sifaka. Proceedings Biol Sci 284.

11. Tian J, Guo W, Yan H, Zhang J, Tian S, Lu J. 2021. Defecating during Morning versus Afternoon: The Gut Microbiota of Zoo Rhesus Macaques. Pakistan J Zool 1–8.

12. Huus KE, Ley RE. 2021. Blowing Hot and Cold: Body Temperature and the Microbiome. mSystems 1–9.

13. Risely A, Wilhelm K, Clutton-Brock T, Al. E. 2022. Gut microbiota repeatability is contigent on temporal scale and age in wild meerkats. Preprint 1–32.

14. Björk AJR, Dasari MR, Roche K, Grieneisen L, Trevor J. 2021. Synchrony and idiosyncrasy in the gut microbiome of wild primates. Nat Ecol Evol.

15. Kruuk H. 1972. The spotted hyena: a study of predation and social behavior. University of Chicago Press, Chicago.

16. Drea CM, Frank LG. 2003. The social complexity of spotted hyenas, p. 121–148. In Waal, FBM de, Tyack, PL (eds.), Animal Social Complexity: Intelligence, Culture, and Individualized Societies. Harvard University Press.

17. Holekamp KE, Sakai ST, Lundrigan BL. 2007. Social intelligence in the spotted hyena (Crocuta crocuta). Phil Trans R Soc B 362:523–538.

18. Frank LG, Holekamp KE SL. 1995. Dominance, demography, and reproductive success of female spotted hyenas, p. 364–384. In Sinclair ARE, AP (ed.), Serengeti II: dynamics, management, and conservation of an ecosystem. University of Chicago Press, Chicago.

19. Green DS, Johnson-Ulrich L, Couraud HE, Holekamp KE. 2018. Anthropogenic disturbance induces opposing population trends in spotted hyenas and African lions. Biodivers Conserv 27:871–889.

20. Frank LG. 1986. Social organization of the spotted hyaena Crocuta crocuta. II. Dominance and reproduction. Anim Behav 34:1510–1527.

21. Strauss ED, Holekamp KE. 2019. Inferring longitudinal hierarchies: Framework and methods for studying the dynamics of dominance. J Anim Ecol 88:521–536.

22. Frank LG. 1986. Social organization of the spotted hyaena (Crocuta crocuta). I. Demography. Anim Behav 34:1500–1509.

23. Holekamp KE, Smith JE, Strelioff CC, Van Horn RC, Watts HE. 2012. Society, demography and genetic structure in the spotted hyena. Mol Ecol 21:613–632.

24. Höner OP, Wachter B, East ML, Streich WJ, Wilhelm K, Burke T, Hofer H. 2007. Female mate-choice drives the evolution of male-biased dispersal in a social mammal. Nature 448:798–801.

25. Smith JE, Kolowski JM, Graham KE, Dawes SE, Holekamp KE. 2008. Social and ecological determinants of fission–fusion dynamics in the spotted hyaena. Anim Behav 76:619–636.

26. Broten MD, Said M. 1995. Population Trends of Ungulates in and around Kenya’s Masai Mara Reserve, p. 169–193. In Serengeti II: Dynamics, Management, and Conservation of an Ecosystem.

27. Green DS. 2015. ANTHROPOGENIC DISTURBANCE, ECOLOGICAL CHANGE, AND WILDLIFE CONSERVATION AT THE EDGE OF THE MARA-SERENGETI ECOSYSTEM. Dissertation.

28. Green DS, Zipkin EF, Incorvaia DC, Holekamp KE. 2019. Long-term ecological changes influence herbivore diversity and abundance inside a protected area in the Mara-Serengeti ecosystem. Glob Ecol Conserv 20:e00697.

29. Pangle WM, Holekamp KE. 2010. Lethal and nonlethal anthropogenic effects on spotted hyenas in the Masai Mara National Reserve. J Mammal 91:154–164.

30. Kolodny O, Schulenburg H. 2020. Microbiome-mediated plasticity directs host evolution along several distinct time-scales. Philos Trans R Soc B2 375:20190589.

31. Suzuki TA. 2017. Links between Natural Variation in the Microbiome and Host Fitness in Wild Mammals. Integr Comp Biol 57:1–14.

32. Cooper SM, Holekamp KE, Smale L. 1999. A seasonal feast: long-term analysis of feeding behaviour in the spotted hyaena (*Crocuta crocuta*). Afr J Ecol 37:149–160.

33. Werdelin L. 1989. Constraint and adaptation in the bone-cracking canid Osteoborus (Mammalia: Canidae). Paleobiology 15.

34. Holekamp KE, Smale L, Berg R, Cooper SM. 1997. Hunting rates and hunting success in the spotted hyena (Crocuta crocuta). J Zool 242:1–15.

35. Milani C, Alessandri G, Mancabelli L, Mangifesta M, Lugli GA, Viappiani A, Longhi G, Anzalone R, Duranti S, Turroni F, Ossiprandi MC, Sinderen D van, Ventura M. 2020. Multi-omics Approaches To Decipher the Impact of Diet and Host Physiology on the Mammalian Gut Microbiome. Appl Environ Microbiol 86:1–21.

36. Perlman D, Martínez-álvaro M, Moraïs S, Altshuler I, Hagen LH, Jami E, Roehe R, Pope PB, Mizrahi I. 2022. Concepts and Consequences of a Core Gut Microbiota for Animal Growth and Development 1–25.

37. Salzman S, Whitaker M, Pierce NE. 2018. Cycad-feeding insects share a core gut microbiome. Biol J Linn Soc 1–11.

38. Bell RHV. 1971. A Grazing Ecosystem in the Serengeti. Sci Am 225:86–93.

39. Maddock L. 1979. The “Migration” and Grazing Succession, p. 104–129. In Sinclair, ARE, Norton-Griffiths, M (eds.), Serengeti: Dynamics of an Ecosystem. The University of Chicago Press, Chicago.

40. Riggio J, Jacobson A, Dollar L, Bauer H, Becker M, Dickman A, Funston P, Groom R, Henschel P, de Iongh H, Lichtenfeld L, Pimm S. 2013. The size of savannah Africa: A lion’s (Panthera leo) view. Biodivers Conserv 22:17–35.

41. Spagnuolo OSB, Jarvey JC, Battaglia MJ, Laubach ZM, Miller ME, Holekamp KE, Bourgeau-chavez LL. 2020. Mapping Kenyan Grassland Heights Across Large Spatial Scales with Combined Optical and Radar Satellite Imagery. Remote Sens 12.

42. Western D, Russell S, Cuthill I. 2009. The Status of Wildlife in Protected Areas Compared to Non-Protected Areas of Kenya. PLoS One 4:e6140.

43. Ogutu JO, Piepho HP, Dublin HT, Bhola N, Reid RS. 2008. Rainfall influences on ungulate population abundance in the Mara-Serengeti ecosystem. J Anim Ecol 77:814–829.

44. Frank LG, Glickman SE, Powch I. 1990. Sexual dimorphism in the spotted hyaena (Crocuta crocuta). J Zool 221:308–313.

45. Holekamp KE, Smale L, Szykman M. 1996. Rank and reproduction in the female spotted hyaena. J Reprod Fertil 108:229–237.

46. Holekamp KE, Szykman M, Boydston EE, Smale L. 1999. Association of seasonal reproductive patterns with changing food availability in an equatorial carnivore, the spotted hyaena (Crocuta crocuta). J Reprod Fertil 116:87–93.

47. Caporaso JG, Lauber CL, Walters WA, Berg-Lyons D, Huntley J, Fierer N, Owens SM, Betley J, Fraser L, Bauer M, Gormley N, Gilbert JA, Smith G, Knight R. 2012. Ultra-high-throughput microbial community analysis on the Illumina HiSeq and MiSeq platforms. ISME J 6:1621–1624.

48. Kozich JJ, Westcott SL, Baxter NT, Highlander SK, Schloss PD. 2013. Development of a Dual-Index Sequencing Strategy and Curation Pipeline for Analyzing Amplicon Sequence Data on the MiSeq Illumina Sequencing Platform. Appl Environ Microbiol 79:5112–5120.

49. R Core Team. 2019. R: a language and environment for statistical computingR Foundation for Statistical Computing. Vienna, Austria.

50. Callahan BJ, McMurdie PJ, Rosen MJ, Han AW, Johnson AJA, Holmes SP. 2016. DADA2: High-resolution sample inference from Illumina amplicon data. Nat Methods 13:581–583.

51. Quast C, Pruesse E, Yilmaz P, Gerken J, Schweer T, Yarza P, Peplies J, Glöckner FO. 2013. The SILVA ribosomal RNA gene database project: improved data processing and web-based tools. Nucleic Acids Res 41:D590–6.

52. Davis NM, Proctor DiM, Holmes SP, Relman DA, Callahan BJ. 2018. Simple statistical identification and removal of contaminant sequences in marker-gene and metagenomics data. Microbiome 6:226.

53. Wickham H. 2009. ggplot2: Elegant Graphics for Dat Analysis. Springer-Verlag New York, New York, NY.

54. Schloss PD, Westcott SL, Ryabin T, Hall JR, Hartmann M, Hollister EB, Lesniewski RA, Oakley BB, Parks DH, Robinson CJ, Sahl JW, Stres B, Thallinger GG, Van Horn DJ, Weber CF. 2009. Introducing mothur: open-source, platform-independent, community-supported software for describing and comparing microbial communities. Appl Environ Microbiol 75:7537–41.

55. Chao, Anne. 1984. Nonparametric Estimation of the Number of Classes in a Population. Scand J Stat 11:265–270.

56. Faith DP. 1992. Conservation evaluation and phylogenetic diversity. Biol Conserv 61.

57. Shannon CE. 1948. A Mathematical Theory of Communication. Bell Syst Tech J 27:379–423.

58. McMurdie PJ, Holmes S. 2013. Phyloseq: An R Package for Reproducible Interactive Analysis and Graphics of Microbiome Census Data. PLoS One 8:1–11.

59. Kembel SW, Cowan PD, Helmus MR, Cornwell WK, Morlon H, Ackerly DD, Blomberg SP, Webb CO. 2010. Picante: R tools for integrating phylogenies and ecology. Bioinformatics 26:1463–1464.

60. Wright ES. 2016. Using DECIPHER v2.0 to analyze big biological sequence data in R. R J 8:352–359.

61. Schliep KP. 2011. phangorn: Phylogenetic analysis in R. Bioinformatics 27:592–593.

62. Fox J, Weisberg S, Fox J. 2011. An R companion to applied regression. Sage, Thousand Oaks, CA.

63. Bates D, Mächler M, Bolker B, Walker S. 2015. Fitting Linear Mixed-Effects Models Using lme4. J Stat Softw 67:1–48.

64. Oksanen J, Blanchet F, Friendly M, Kindt R, Legendre P, McGlinn D, Minchin P, R. B. O’Hara, Gavin L. Simpson, Peter Solymos M, Henry H. Stevens, Eduard Szoecs, Helene Wagner. 2018. vegan: Community Ecology Package. R Packag version 24–6.

65. Kuznetsova A, Brockhoff PB, Christensen RHB. 2017. lmerTest Package: Tests in Linear Mixed Effects Models. J Stat Softw 82:1–26.

66. Bolger AM, Lohse M, Usadel B. 2014. Trimmomatic: a flexible trimmer for Illumina sequence data. Bioinformatics 30:2114–2120.

67. Yang C, Li F, Xiong Z, Koepfli KP, Ryder O, Perelman P, Li Q, Zhang G. 2020. A draft genome assembly of spotted hyena, Crocuta crocuta. Sci Data 7:1–10.

68. Kim D, Paggi JM, Park C, Bennett C, Salzberg SL. 2019. Graph-based genome alignment and genotyping with HISAT2 and HISAT-genotype. Nat Biotechnol 37:907–915.

69. Boisvert S, Raymond F, Godzaridis É, Laviolette F, Corbeil J. 2012. Ray Meta: Scalable de novo metagenome assembly and profiling. Genome Biol 13.

70. Wood DE, Lu J, Langmead B. 2019. Improved metagenomic analysis with Kraken 2. Genome Biol 20:1–13.

71. Li D, Liu CM, Luo R, Sadakane K, Lam TW. 2015. MEGAHIT: An ultra-fast single-node solution for large and complex metagenomics assembly via succinct de Bruijn graph. Bioinformatics 31:1674–1676.

72. Gurevich A, Saveliev V, Vyahhi N, Tesler G. 2013. QUAST: quality assessment tool for genome assemblies. Bioinformatics 29:1072–1075.

73. Eren AM, Esen OC, Quince C, Vineis JH, Morrison HG, Sogin ML, Delmont TO. 2015. Anvi’o: An advanced analysis and visualization platformfor ‘omics data. PeerJ 2015:1–29.

74. Doug Hyatt, Gwo-Liang Chen, Philip F LoCascio, Miriam L Land, Frank W Larimer LJH. 2010. Prodigal: prokaryotic gene recognition and translation initiation site identification. BMC Bioinformatics 11:1–8.

75. Tatusov RL, Galperin MY, Natale DA, Koonin E V. 2000. The COG database: A tool for genome-scale analysis of protein functions and evolution. Nucleic Acids Res 28:33–36.

76. Kanehisa M, Sato Y, Kawashima M, Furumichi M, Tanabe M. 2016. KEGG as a reference resource for gene and protein annotation. Nucleic Acids Res 44:D457–D462.

77. Patro R, Duggal G, Love MI, Irizarry RA, Kingsford C. 2017. Salmon provides fast and bias-aware quantification of transcript expression. Nat Methods 14:417–419.

78. Kang DD, Li F, Kirton E, Thomas A, Egan R, An H, Wang Z. 2019. MetaBAT 2: An adaptive binning algorithm for robust and efficient genome reconstruction from metagenome assemblies. PeerJ 2019:1–13.

79. Parks DH, Imelfort M, Skennerton CT, Hugenholtz P, Tyson GW. 2015. CheckM: Assessing the quality of microbial genomes recovered from isolates, single cells, and metagenomes. Genome Res 25:1043–1055.

80. Chaumeil PA, Mussig AJ, Hugenholtz P, Parks DH. 2020. GTDB-Tk: A toolkit to classify genomes with the genome taxonomy database. Bioinformatics 36:1925–1927.

81. Stamatakis A. 2014. RAxML version 8: A tool for phylogenetic analysis and post-analysis of large phylogenies. Bioinformatics 30:1312–1313.

82. Letunic I, Bork P. 2007. Interactive Tree Of Life (iTOL): An online tool for phylogenetic tree display and annotation. Bioinformatics 23:127–128.

83. Yu G. 2020. Using ggtree to Visualize Data on Tree-Like Structures. Curr Protoc Bioinforma 69:1–18.

84. Chen L, Liu M, Zhu J, Gao Y, Sha W, Ding H, Jiang W, Wu S. 2020. Age, Gender, and Feeding Environment Influence Fecal Microbial Diversity in Spotted Hyenas (Crocuta crocuta). Curr Microbiol 77:1139–1149.

85. Menke S, Meier M, Mfune JKE, Melzheimer J, Wachter B, Sommer S. 2017. Effects of host traits and land-use changes on the gut microbiota of the Namibian black-backed jackal (Canis mesomelas). FEMS Microbiol Ecol 93:1–16.

86. Cuscó A, Pérez D, Viñes J, Fàbregas N, Francino O. 2021. Long-read metagenomics retrieves complete single-contig bacterial genomes from canine feces 1–15.

87. de Jonge N, Carlsen B, Christensen MH, Pertoldi C, Nielsen JL. 2022. The Gut Microbiome of 54 Mammalian Species. Front Microbiol 13:1–11.

88. Bermingham EN, Maclean P, Thomas DG, Cave NJ, Young W. 2017. Key bacterial families (Clostridiaceae, Erysipelotrichaceae and Bacteroidaceae) are related to the digestion of protein and energy in dogs. PeerJ 5:e3019.

89. Whitehead TR, Cotta MA, Falsen E, Moore E, Lawson PA. 2011. Peptostreptococcus russellii sp. nov., isolated from a swine-manure storage pit. Int J Syst Evol Microbiol 61:1875–1879.

90. Wlodarska M, Luo C, Kolde R, d’Hennezel E, Annand JW, Heim CE, Krastel P, Schmitt EK, Omar AS, Creasey EA, Garner AL, Mohammadi S, O’Connell DJ, Abubucker S, Arthur TD, Franzosa EA, Huttenhower C, Murphy LO, Haiser HJ, Vlamakis H, Porter JA, Xavier RJ. 2017. Indoleacrylic Acid Produced by Commensal Peptostreptococcus Species Suppresses Inflammation. Cell Host Microbe 22:25–37.e6.

91. Andrewes FW, Horder TJ. 1906. a Study of the Streptococci Pathogenic for Man. Lancet 168:708–713.

92. Thingholm LB, Bang C, Rühlemann MC, Starke A, Sicks F, Kaspari V, Jandowsky A, Frölich K, Ismer G, Bernhard A, Bombis C, Struve B, Rausch P, Franke A. 2021. Ecology impacts the decrease of Spirochaetes and Prevotella in the fecal gut microbiota of urban humans. BMC Microbiol 21:1–20.

93. Alauzet C, Jumas-Bilak E. 2014. The phylum deferribacteres and the genus caldithrix. Prokaryotes Other Major Lineages Bact Archaea 9783642389:595–611.

94. Hildebrand F, Nguyen TLA, Brinkman B, Yunta RG, Cauwe B, Vandenabeele. P, Liston A, Raes J. 2013. Inflammation-associated enterotypes, host genotype, cage and inter-individual effects drive gut microbiota variation in common laboratory mice. Genome Biol 14:1–9.

95. Scupham AJ, Patton TG, Bent E, Bayles DO. 2008. Comparison of the cecal microbiota of domestic and wild turkeys. Microb Ecol 56:322–331.

96. Li RW, Wu S, Li W, Navarro K, Couch RD, Hill D, Urban JF. 2012. Alterations in the porcine colon microbiota induced by the gastrointestinal nematode Trichuris suis. Infect Immun 80:2150–2157.

97. Suchodolski JS, Markel ME, Garcia-Mazcorro JF, Unterer S, Heilmann RM, Dowd SE, Kachroo P, Ivanov I, Minamoto Y, Dillman EM, Steiner JM, Cook AK, Toresson L. 2012. The Fecal Microbiome in Dogs with Acute Diarrhea and Idiopathic Inflammatory Bowel Disease. PLoS One 7.

98. Bendová B, Piálek J, Ď L’, Schmiedová L, Dagmar Č, Martin J, Kreisinger J. 2020. How being synanthropic affects the gut bacteriome and mycobiome: comparison of two mouse species with contrasting ecologies. BMC Microbiol 20:1–13.

99. Barelli C, Albanese D, Stumpf RM, Asangba AE, Donati C, Rovero F, Hauffe HC. 2020. The Gut Microbiota Communities of Wild Arboreal and Ground-Feeding Tropical Primates Are Affected Differently by Habitat Disturbance. mSystems 5:1–18.

100. Watson SE, Hauffe HC, Bull M, McKinney MA, Atwood TA, Perkins SE. 2019. Global change-driven use of onshore habitat impacts polar bear faecal microbiota. ISME J.

101. Sugden S, Sanderson D, Ford K, Stein LY, St. Clair CC. 2020. An altered microbiome in urban coyotes mediates relationships between anthropogenic diet and poor health. Sci Rep 10:1–14.

102. Donohue ME, Weisrock DW, Asangba AE, Ralainirina J, Stumpf RM, Wright PC. 2019. Extensive variability in the gut microbiome of a highly - specialized and critically endangered lemur species across sites. Am J Primatol e23046:1–12.

103. Richardson JB, Dancy BCR, Horton CL, Lee YS, Madejczyk MS, Xu ZZ, Ackermann G, Humphrey G, Palacios G, Knight R, Lewis JA. 2018. Exposure to toxic metals triggers unique responses from the rat gut microbiota. Sci Rep 8:1–12.

104. Sugden S, St. Clair CC, Stein LY. 2020. Individual and Site-Specific Variation in a Biogeographical Profile of the Coyote Gastrointestinal Microbiota. Microb Ecol.

105. Li J, Zhan S, Liu X, Lin Q, Jiang J, Li X. 2018. Divergence of Fecal Microbiota and Their Associations With Host Phylogeny in Cervinae. Front Microbiol 9:1–11.

106. Holekamp KE, Strauss ED. 2016. Aggression and dominance: an interdisciplinary overview. Curr Opin Behav Sci 12:44–51.

107. Flies AS, Mans LS, Flies EJ, Grant CK, Holekamp KE. 2016. Socioecological predictors of immune defences in wild spotted hyenas 1549–1557.

108. Life E, Reese AT, Phillips SR, Owens LA, Goldberg TL, Thompson ME, Carmody RN. 2021. Age Patterning in Wild Chimpanzee Gut Microbiota Diversity Reveals Differences from Humans in Age Patterning in Wild Chimpanzee Gut Microbiota Diversity Reveals Differences from Humans in Early Life. Curr Biol 1–8.

109. Tian J, Du J, Han J, Song X, Lu Z. 2020. Age-related differences in gut microbial community composition of captive spotted seals (Phoca largha). Mar Mammal Sci 1–10.

110. Yatsunenko T, Rey FE, Manary MJ, Trehan I, Dominguez-Bello MG, Contreras M, Magris M, Hidalgo G, Baldassano RN, Anokhin AP, Heath AC, Warner B, Reeder J, Kuczynski J, Caporaso JG, Lozupone CA, Lauber C, Clemente JC, Knights D, Knight R, Gordon JI. 2012. Human gut microbiome viewed across age and geography. Nature 486:222–7.

111. Guo W, Zhou M, Ma T, Bi S, Wang W, Zhang Y, Huang X, Guan LL. 2020. Survey of rumen microbiota of domestic grazing yak during different growth stages revealed novel maturation patterns of four key microbial groups and their dynamic interactions. Anim Microbiome 2.

112. Heitlinger E, Ferreira SCM, Thierer D, Hofer H, East ML. 2017. The Intestinal Eukaryotic and Bacterial Biome of Spotted Hyenas: The Impact of Social Status and Age on Diversity and Composition. Front Cell Infect Microbiol 7.

113. Nagpal R, Mainali R, Ahmadi S, Wang S, Singh R, Kavanagh K, Kitzman DW, Kushugulova A, Marotta F, Yadav H. 2018. Gut microbiome and aging: Physiological and mechanistic insights. Nutr Heal Aging 4:267–285.

114. Jeste D V, Nguyen TT. 2020. The Gut Microbiome, Aging, and Longevity: A Systematic Review. Nutrients 12:1–25.

115. O’Toole PW, Jeffery IB. 2015. Gut microbiota and aging. Science (80-). American Association for the Advancement of Science.

116. Mitchell EL, Davis AT, Brass K, Dendinger M, Barner R, Gharaibeh R, Fodor AA, Kavanagh K. 2017. Reduced intestinal motility, mucosal barrier function, and inflammation in aged monkeys. J Nutr Heal Aging 21:354–361.

117. Kriss M, Hazleton KZ, Nusbacher NM, Martin CG, Lozupone CA. 2018. Low diversity gut microbiota dysbiosis: drivers, functional implications and recovery. Curr Opin Microbiol 44:34–40.

118. Smith JE, Kolowski JM, Graham KE, Dawes SE, Holekamp KE. 2008. Social and ecological determinants of fission-fusion dynamics in the spotted hyaena. Anim Behav 76:619–636.

119. Watts HE, Scribner KT, Garcia HA, Holekamp KE, Heather Watts CE, Volff J-N. 2011. Genetic diversity and structure in two spotted hyena populations reflects social organization and male dispersal. J Zool 285:281–291.

120. Engh AL, Funk SM, Horn RC Van, Scribner KT, Bruford MW, Libants S, Szykman M, Smale L, Holekamp KE. 2002. Reproductive skew among males in a female-dominated mammalian society. Behav Ecol 13.

121. Kolowski JM, Katan D, Theis KR, Holekamp KE. DAILY PATTERNS OF ACTIVITY IN THE SPOTTED HYENA.

122. Lv X, Chai J, Diao Q, Huang W, Zhuang Y, Zhang N. 2019. The signature microbiota drive rumen function shifts in goat kids introduced to solid diet regimes. Microorganisms 7.

123. Greene LK, McKenney EA, O’Connell TM, Drea CM. 2018. The critical role of dietary foliage in maintaining the gut microbiome and metabolome of folivorous sifakas. Sci Rep 8:1–13.

124. Levin D, Raab N, Pinto Y, Rothschild D, Zanir G, Godneva A, Mellul N, Futorian D, Gal D, Leviatan S, Zeevi D, Bachelet I, Segal E. 2021. Diversity and functional landscapes in the microbiota of animals in the wild. Science (80-) 372.

125. Cabral L, Persinoti GF, Paixão DAA, Martins MP, Morais MAB, Chinaglia M, Domingues MN, Sforca ML, Pirolla RAS, Generoso WC, Santos CA, Maciel LF, Terrapon N, Lombard V, Henrissat B, Murakami MT. 2022. Gut microbiome of the largest living rodent harbors unprecedented enzymatic systems to degrade plant polysaccharides. Nat Commun 13:1–16.

126. Wu X, Wei Q, Wang X, Shang Y, Zhang H. 2021. Evolutionary and dietary relationships of wild mammals based on the gut microbiome. Gene 145999.

127. Colborn AS, Kuntze CC, Gadsden GI, Harris NC. 2020. Spatial variation in diet-microbe associations across populations of a generalist North American carnivore. J Anim Ecol 1–9.

128. Alessandri G, Milani C, Mancabelli L, Longhi G, Anzalone R, Lugli GA, Duranti S, Turroni F, Ossiprandi MC, van Sinderen D, Ventura M. 2020. Deciphering the bifidobacterial populations within the canine and feline gut microbiota. Appl Environ Microbiol 86.

129. Zhang F, Xu N, Wang W, Yu Y, Wu S. 2021. The gut microbiome of the Sunda pangolin (Manis javanica) reveals its adaptation to specialized myrmecophagy. PeerJ 9:e11490.

130. Li Q, Lauber CL, Czarnecki-Maulden G, Pan Y, Hannah SS. 2017. Effects of the Dietary Protein and Carbohydrate Ratio on Gut Microbiomes in Dogs of Different Body Conditions. MBio 8:1–14.

131. Zhu Y, Han Z, Wang H, Liu C, Si H, Xu C. 2021. Adaptation of the Gut Microbiota of Amur Tigers to a Special Diet. Curr Microbiol 78:1628–1635.

132. Ezaki T, Kawamura Y, Li N, Li ZY, Zhao L, Shu SE. 2001. Proposal of the genera Anaerococcus gen. nov., Peptoniphilus gen. nov. and Gallicola gen. nov for members of the genus Peptostreptococcus. Int J Syst Evol Microbiol 51:1521–1528.

133. Youngblut ND, de la Cuesta-Zuluaga J, Reischer GH, Dauser S, Schuster N, Walzer C, Stalder G, Farnleitner AH, Ley RE. 2020. Large-Scale Metagenome Assembly Reveals Novel Animal-Associated Microbial Genomes, Biosynthetic Gene Clusters, and Other Genetic Diversity. mSystems 5:1–15.

