## SupplementalFigures1-7 for "Taxonomic, genomic, and functional variation in the gut microbiomes of wild spotted hyenas across two decades of study"

**List of Supplementary figures:**

Fig S1: Histogram of number of 16S rRNA sequences in samples; rarefaction curve of ASV Richness
Fig S2: Heatmap of contaminant ASVs in blank control samples (e.g. sterile swabs)
Fig S3: PCoA ordinations based on 16S rRNA data
Fig S4: Phylum-level barplot of metagenome taxonomic composition (Kraken2)
Fig S5: MAG completeness x contamination; histogram of MAG sizes (Kb)
Fig S6: 149 Phylogenetic Trees of each MAGs showing evolutionary distance to GTDB genomes
Fig S7: PCoA ordinations based on MAG abundance data

**Figure S1. A) Number of reads per sample plotted on the normal scale, B) Rarefaction curves of gut microbiome ASV richness**. Plotted are the number of ASVs (ASV Richness) that are recovered given a certain number of sequences. Each curve represents a unique sample and is color-coded by hyena identity and ordered by matriline (1-highest ranking, 4-lowest ranking) and generation (M-mother, D-daughter, G-granddaughter). The y=x black line shows what the curve would look like if a new ASV came with each read.

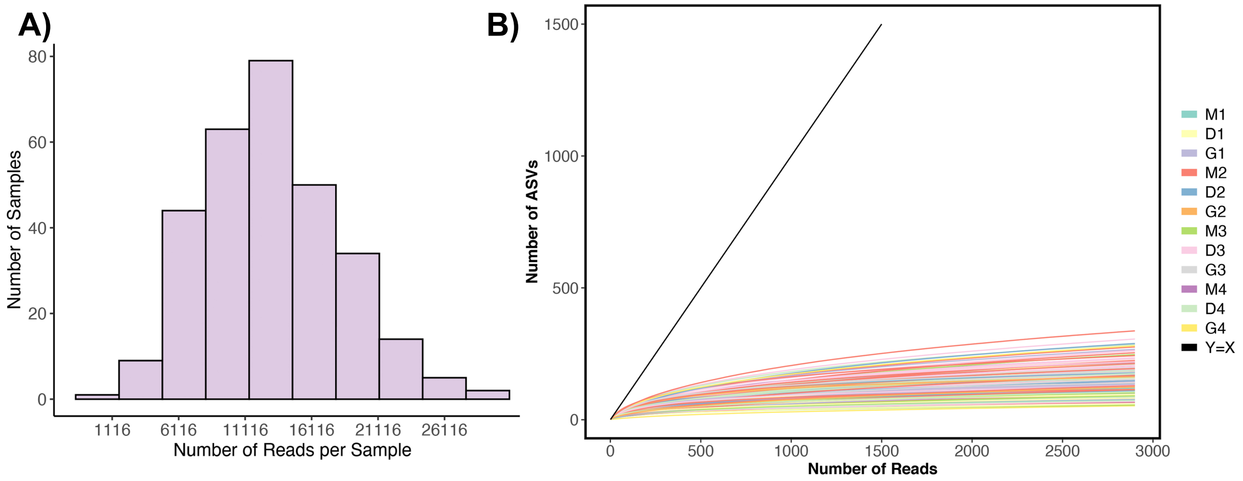

**Figure S2. Relative abundances of ASVs from DNA extraction kit control samples.** Heatmap showing ASVs with average proportional abundances >0.007 (i.e., 0.7%) in control samples (DNA extracted from sterile cotton swabs). Darker colors in the heatmap indicate higher proportional abundances. ASVs in red font were removed from the dataset after being identified as contaminants by the R decontam package and our additional criteria (e.g., must have been present in at least 50% of control samples at proportional abundances >0.01).

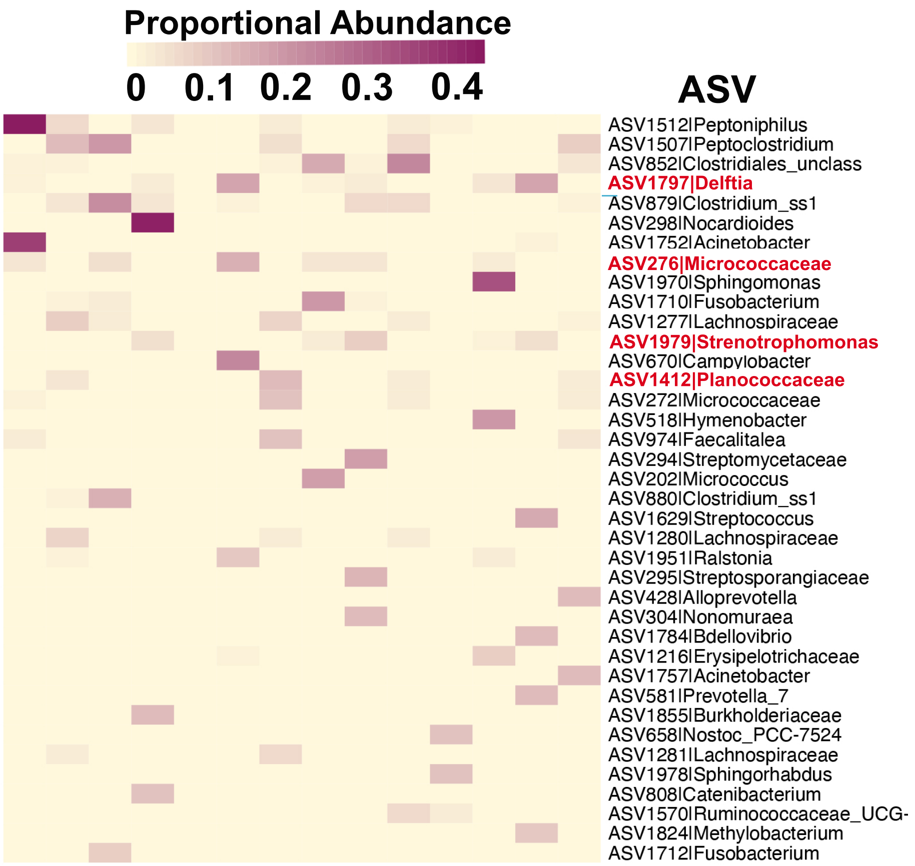

**Figure S3. PCoA ordination plots color-coded by (A) matriline, (B) age, (C) mean monthly prey availability, and (D) sample year**. Plots show all 301 samples spanning 23 years; each point is a sample, and axes represent the first two principal coordinates. Closeness of points indicates high community similarity based on weighted Unifrac distances.

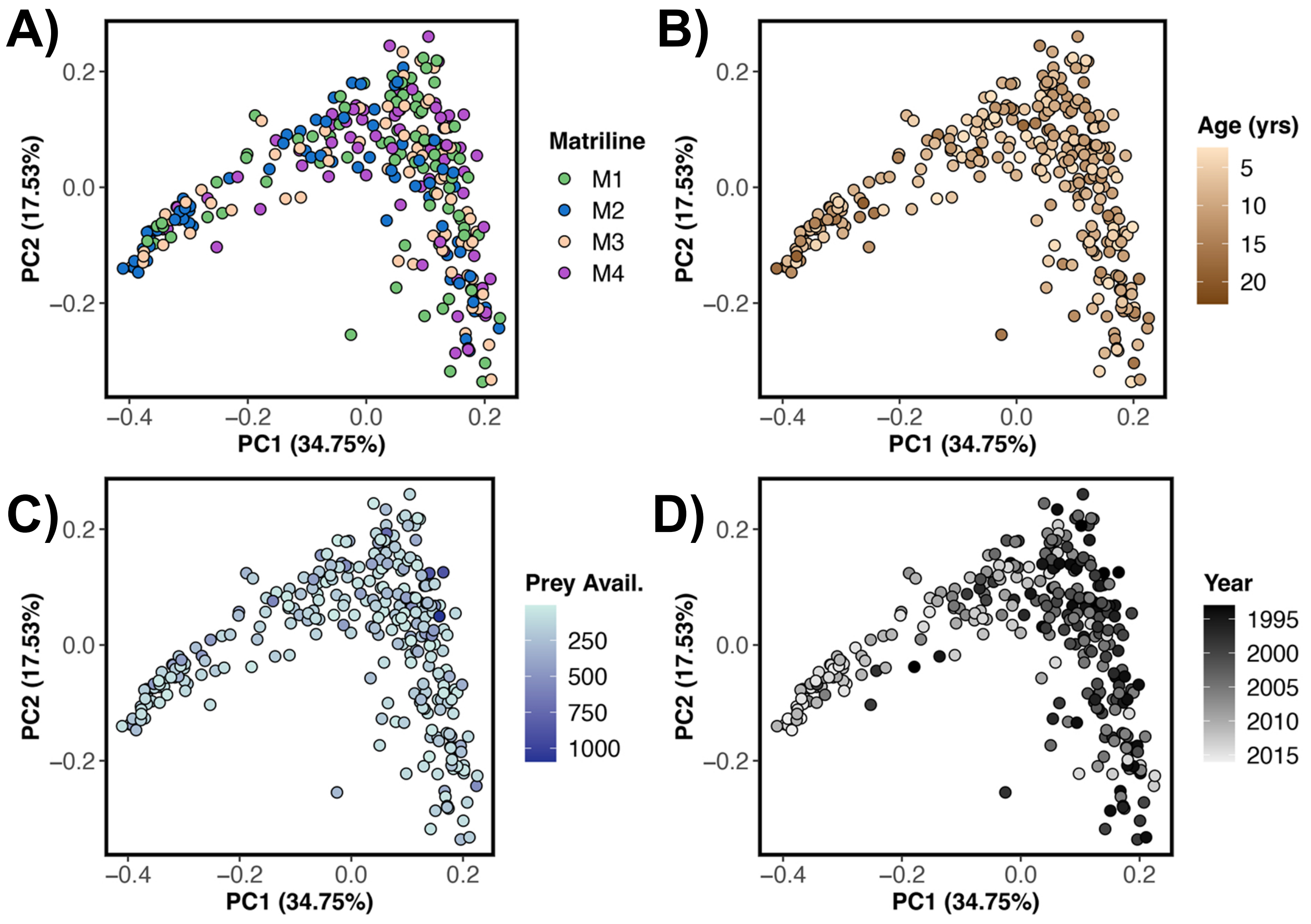

**Figure S4. Taxonomic composition of gut metagenomes at the phylum level.** Kraken2 was used for assigning taxonomic labels to trimmed metagenomic DNA sequences. Stacked bar plots show the relative frequency of sequences assigned to each bacterial phylum across samples. Samples are ordered by individual (D-daughter, G-granddaughter, 1 – matriline #1, 3– matriline #3), and each color represents a bacterial phylum. See Table S2 for the metadata associated with these samples.

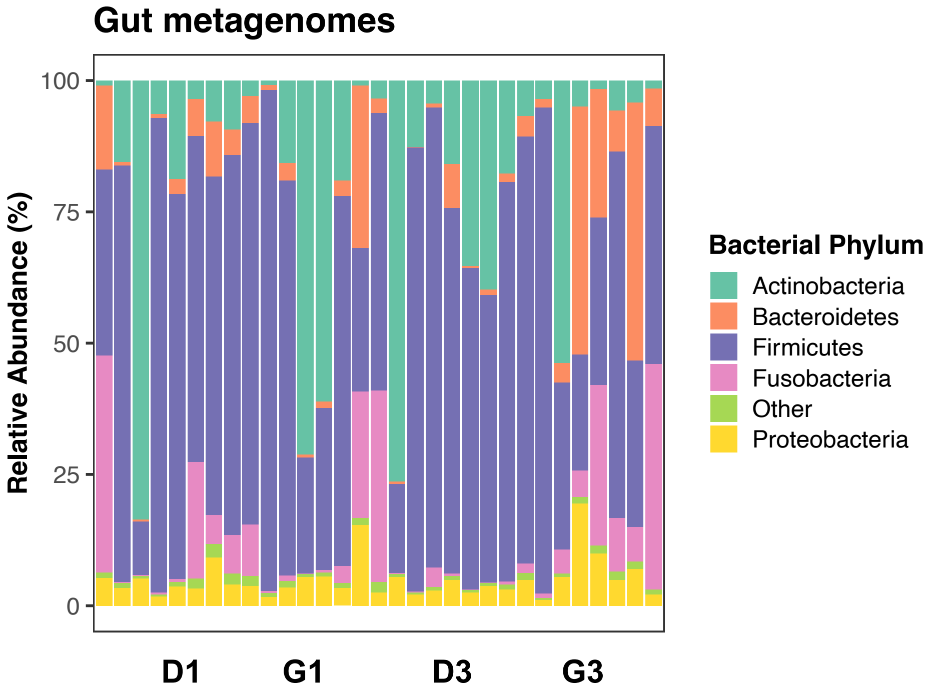


**Figure S5. MAG completeness and contamination scores. A)** plot of MAG completeness vs. MAG contamination, **B)** histogram of MAG genome sizes. MAGs were reconstructed with Metabat2 and assessed for quality with CheckM.

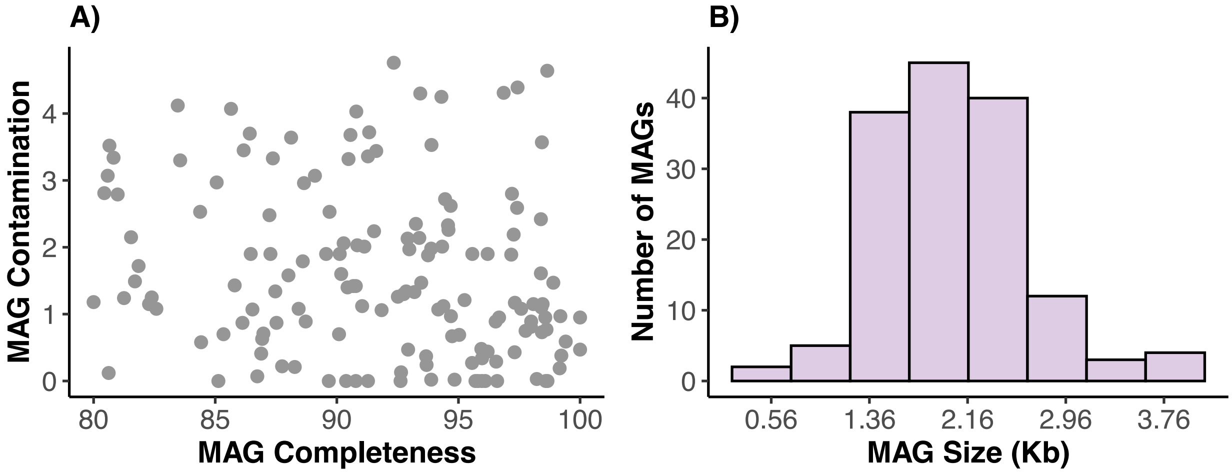

**Fig S6. Individual phylogenetic trees of 149 high-quality MAGs based on the Genome Database Taxonomy (GTDB).** Trees show the evolutionary distance of each MAG to the closest genomes in GTDB - release 95. MAG nodes from this study have an orange label.

*See the 149 trees below.*

**Figure S7. MAG abundances do not cluster by host individual identity.** PCoA ordination based on Bray-Curtis distances calculated from MAG abundances in each sample. The program CoverM calculated the proportion of reads in a sample that mapped to each MAG. For figure key: D-daughter, G-granddaughter, 1 – matriline #1, 3– matriline #3.

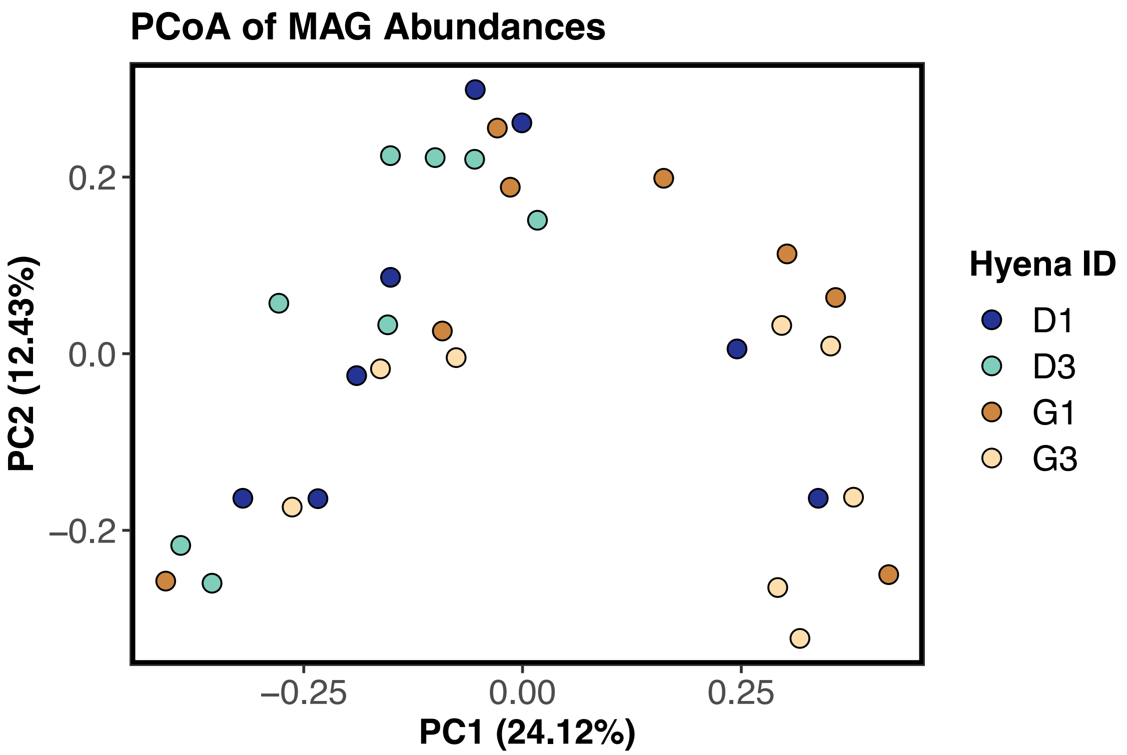

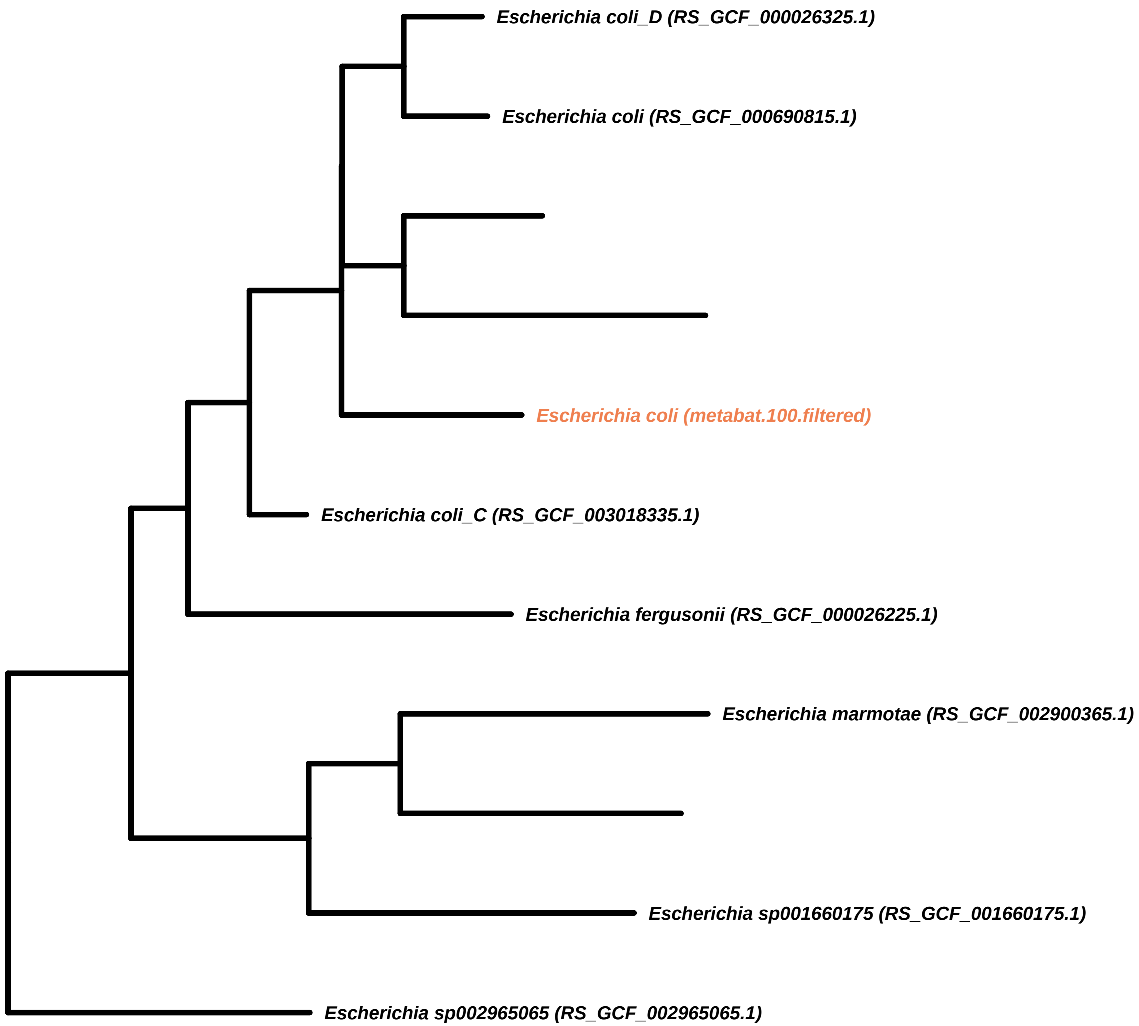

Escherichia coli_D (RS_GCF_000026325.1)
Escherichia coli (RS_GCF_000690815.1)
Escherichia coli (metabat.100.filtered)
Escherichia coli_C (RS_GCF_003018335.1)
Escherichia fergusonii (RS_GCF_000026225.1)
Escherichia marmotae (RS_GCF_002900365.1)
Escherichia sp001660175 (RS_GCF_001660175.1)
Escherichia sp002965065 (RS_GCF_002965065.1)

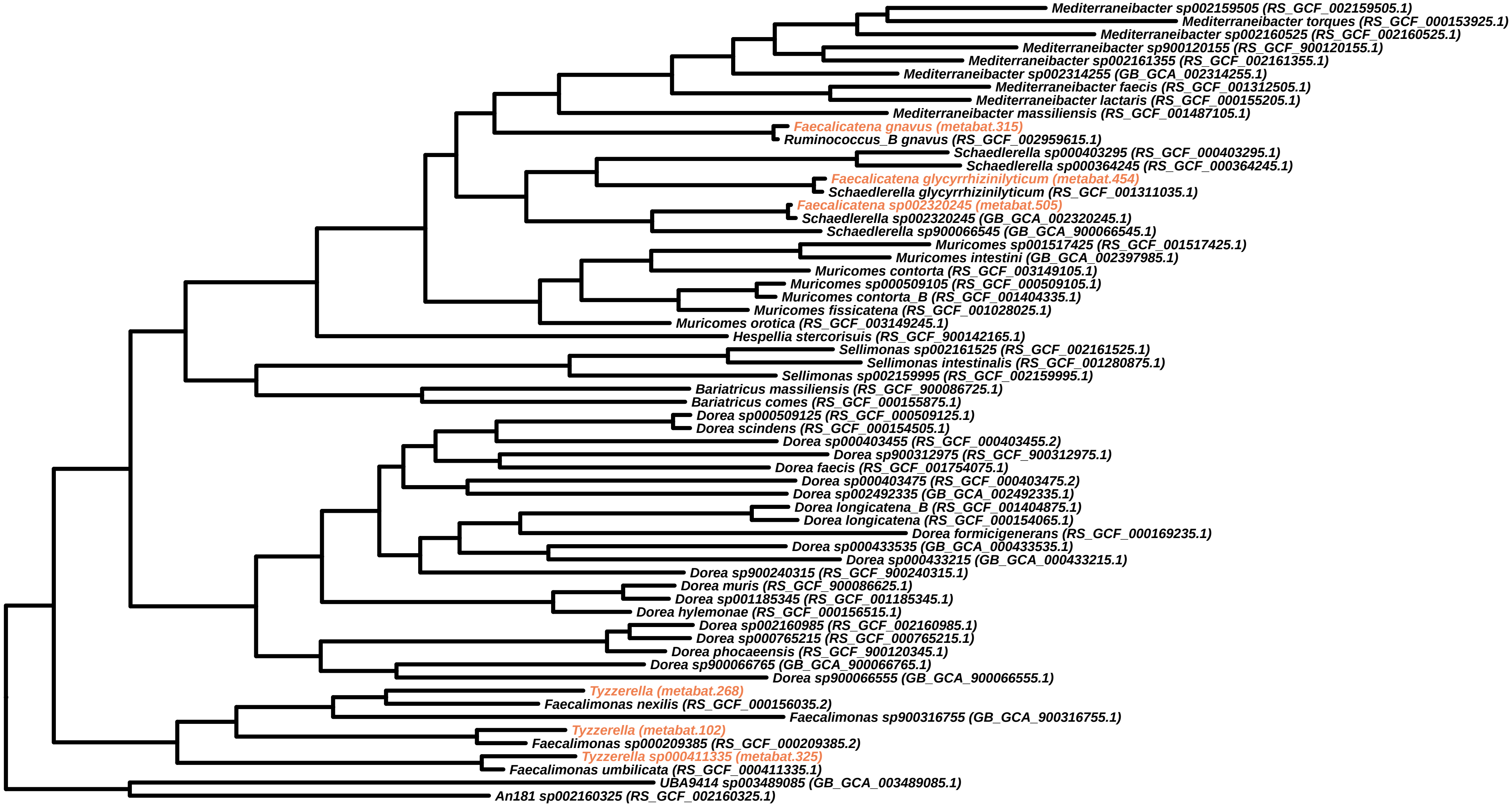
Mediterraneibacter sp002159505 (RS_GCF_002159505.1)
Mediterraneibacter torques (RS_GCF_000153925.1)
Mediterraneibacter sp002160525 (RS_GCF_002160525.1)
Mediterraneibacter sp900120155 (RS_GCF_900120155.1)
Mediterraneibacter sp002161355 (RS_GCF_002161355.1)
Mediterraneibacter sp002314255 (GB_GCA_002314255.1)
Mediterraneibacter faecis (RS_GCF_001312505.1)
Mediterraneibacter lactaris (RS_GCF_000155205.1)
Mediterraneibacter massiliensis (RS_GCF_001487105.1)
Faecalicatena gnavus (metabat.315)
Ruminococcus_B gnavus (RS_GCF_002959615.1)
Schaedlerella sp000403295 (RS_GCF_000403295.1)
Schaedlerella sp000364245 (RS_GCF_000364245.1)
Faecalicatena glycyrrhizinilyticum (metabat.454)
Schaedlerella glycyrrhizinilyticum (RS_GCF_001311035.1)
Faecalicatena sp002320245 (metabat.505)
Schaedlerella sp002320245 (GB_GCA_002320245.1)
Schaedlerella sp900066545 (GB_GCA_900066545.1)
Muricomes sp001517425 (RS_GCF_001517425.1)
Muricomes intestini (GB_GCA_002397985.1)
Muricomes contorta (RS_GCF_003149105.1)
Muricomes sp000509105 (RS_GCF_000509105.1)
Muricomes contorta_B (RS_GCF_001404335.1)
Muricomes fissicatena (RS_GCF_001028025.1)
Muricomes orotica (RS_GCF_003149245.1)
Hespellia stercorisuis (RS_GCF_900142165.1)
Sellimonas sp002161525 (RS_GCF_002161525.1)
Sellimonas intestinalis (RS_GCF_001280875.1)
Sellimonas sp002159995 (RS_GCF_002159995.1)
Bariatricus massiliensis (RS_GCF_900086725.1)
Bariatricus comes (RS_GCF_000155875.1)
Dorea sp000509125 (RS_GCF_000509125.1)
Dorea scindens (RS_GCF_000154505.1)
Dorea sp000403455 (RS_GCF_000403455.2)
Dorea sp900312975 (RS_GCF_900312975.1)
Dorea faecis (RS_GCF_001754075.1)
Dorea sp000403475 (RS_GCF_000403475.2)
Dorea sp002492335 (GB_GCA_002492335.1)
Dorea longicatena_B (RS_GCF_001404875.1)
Dorea longicatena (RS_GCF_000154065.1)
Dorea formicigenerans (RS_GCF_000169235.1)
Dorea sp000433535 (GB_GCA_000433535.1)
Dorea sp000433215 (GB_GCA_000433215.1)
Dorea sp900240315 (RS_GCF_900240315.1)
Dorea muris (RS_GCF_900086625.1)
Dorea sp001185345 (RS_GCF_001185345.1)
Dorea hylemonae (RS_GCF_000156515.1)
Dorea sp002160985 (RS_GCF_002160985.1)
Dorea sp000765215 (RS_GCF_000765215.1)
Dorea phocaeensis (RS_GCF_900120345.1)
Dorea sp900066765 (GB_GCA_900066765.1)
Dorea sp900066555 (GB_GCA_900066555.1)
Tyzzerella (metabat.268)
Faecalimonas nexilis (RS_GCF_000156035.2)
Faecalimonas sp900316755 (GB_GCA_900316755.1)
Tyzzerella (metabat.102)
Faecalimonas sp000209385 (RS_GCF_000209385.2)
Tyzzerella sp000411335 (metabat.325)
Faecalimonas umbilicata (RS_GCF_000411335.1)
UBA9414 sp003489085 (GB_GCA_003489085.1)
An181 sp002160325 (RS_GCF_002160325.1)

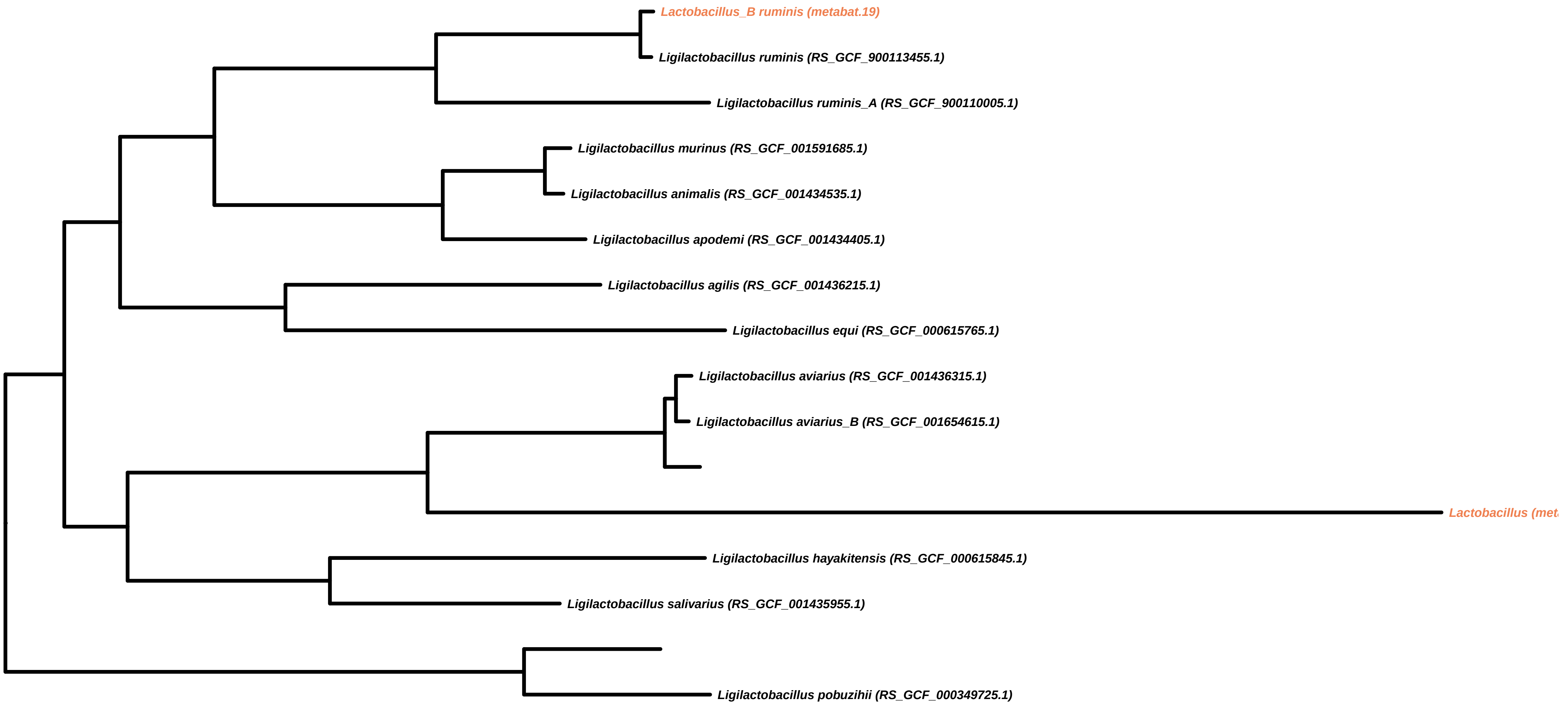

Lactobacillus_B ruminis (metabat.19)
Ligilactobacillus ruminis (RS_GCF_900113455.1)
Ligilactobacillus ruminis_A (RS_GCF_900110005.1)
Ligilactobacillus murinus (RS_GCF_001591685.1)
Ligilactobacillus animalis (RS_GCF_001434535.1)
Ligilactobacillus apodemi (RS_GCF_001434405.1)
Ligilactobacillus agilis (RS_GCF_001436215.1)
Ligilactobacillus equi (RS_GCF_000615765.1)
Ligilactobacillus aviarius (RS_GCF_001436315.1)
Ligilactobacillus aviarius_B (RS_GCF_001654615.1)
Ligilactobacillus hayakitensis (RS_GCF_000615845.1)
Ligilactobacillus salivarius (RS_GCF_001435955.1)
Ligilactobacillus pobuzihii (RS_GCF_000349725.1)

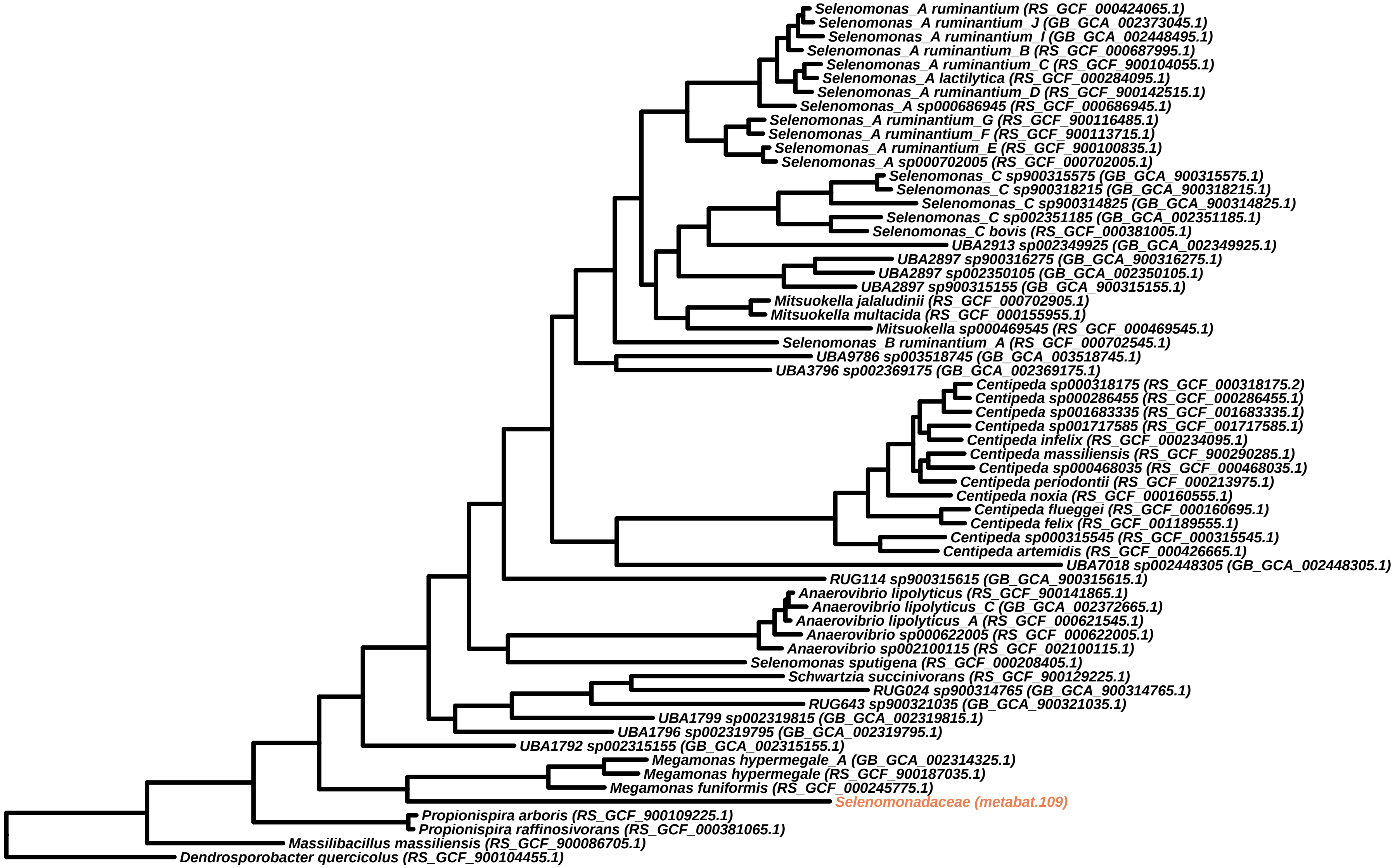
Selenomonas_A ruminantium (RS_GCF_000424065.1)
Selenomonas_A ruminantium_J (GB_GCA_002373045.1)
Selenomonas_A ruminantium_I (GB_GCA_002448495.1)
Selenomonas_A ruminantium_B (RS_GCF_000687995.1)
Selenomonas_A ruminantium_C (RS_GCF_900104055.1)
Selenomonas_A lactilytica (RS_GCF_000284095.1)
Selenomonas_A ruminantium_D (RS_GCF_900142515.1)
Selenomonas_A sp000686945 (RS_GCF_000686945.1)
Selenomonas_A ruminantium_G (RS_GCF_900116485.1)
Selenomonas_A ruminantium_F (RS_GCF_900113715.1)
Selenomonas_A ruminantium_E (RS_GCF_900100835.1)
Selenomonas_A sp000702005 (RS_GCF_000702005.1)
Selenomonas_C sp900315575 (GB_GCA_900315575.1)
Selenomonas_C sp900318215 (GB_GCA_900318215.1)
Selenomonas_C sp900314825 (GB_GCA_900314825.1)
Selenomonas_C sp002351185 (GB_GCA_002351185.1)
Selenomonas_C bovis (RS_GCF_000381005.1)
UBA2913 sp002349925 (GB_GCA_002349925.1)
UBA2897 sp900316275 (GB_GCA_900316275.1)
UBA2897 sp002350105 (GB_GCA_002350105.1)
UBA2897 sp900315155 (GB_GCA_900315155.1)
Mitsuokella jalaludinii (RS_GCF_000702905.1)
Mitsuokella multacida (RS_GCF_000155955.1)
Mitsuokella sp000469545 (RS_GCF_000469545.1)
Selenomonas_B ruminantium_A (RS_GCF_000702545.1)
UBA9786 sp003518745 (GB_GCA_003518745.1)
UBA3796 sp002369175 (GB_GCA_002369175.1)
Centipeda sp000318175 (RS_GCF_000318175.2)
Centipeda sp000286455 (RS_GCF_000286455.1)
Centipeda sp001683335 (RS_GCF_001683335.1)
Centipeda sp001717585 (RS_GCF_001717585.1)
Centipeda infelix (RS_GCF_000234095.1)
Centipeda massiliensis (RS_GCF_900290285.1)
Centipeda sp000468035 (RS_GCF_000468035.1)
Centipeda periodontii (RS_GCF_000213975.1)
Centipeda noxia (RS_GCF_000160555.1)
Centipeda flueggei (RS_GCF_000160695.1)
Centipeda felix (RS_GCF_001189555.1)
Centipeda sp000315545 (RS_GCF_000315545.1)
Centipeda artemidis (RS_GCF_000426665.1)
UBA7018 sp002448305 (GB_GCA_002448305.1)
RUG114 sp900315615 (GB_GCA_900315615.1)
Anaerovibrio lipolyticus (RS_GCF_900141865.1)
Anaerovibrio lipolyticus_C (GB_GCA_002372665.1)
Anaerovibrio lipolyticus_A (RS_GCF_000621545.1)
Anaerovibrio sp000622005 (RS_GCF_000622005.1)
Anaerovibrio sp002100115 (RS_GCF_002100115.1)
Selenomonas sputigena (RS_GCF_000208405.1)
Schwartzia succinivorans (RS_GCF_900129225.1)
RUG024 sp900314765 (GB_GCA_900314765.1)
RUG643 sp900321035 (GB_GCA_900321035.1)
UBA1799 sp002319815 (GB_GCA_002319815.1)
UBA1796 sp002319795 (GB_GCA_002319795.1)
UBA1792 sp002315155 (GB_GCA_002315155.1)
Megamonas hypermegale_A (GB_GCA_002314325.1)
Megamonas hypermegale (RS_GCF_900187035.1)
Megamonas funiformis (RS_GCF_000245775.1)
Selenomonadaceae (metabat.109)
Propionispira arboris (RS_GCF_900109225.1)
Propionispira raffinosivorans (RS_GCF_000381065.1)
Massilibacillus massiliensis (RS_GCF_900086705.1)
Dendrosporobacter quercicolus (RS_GCF_900104455.1)

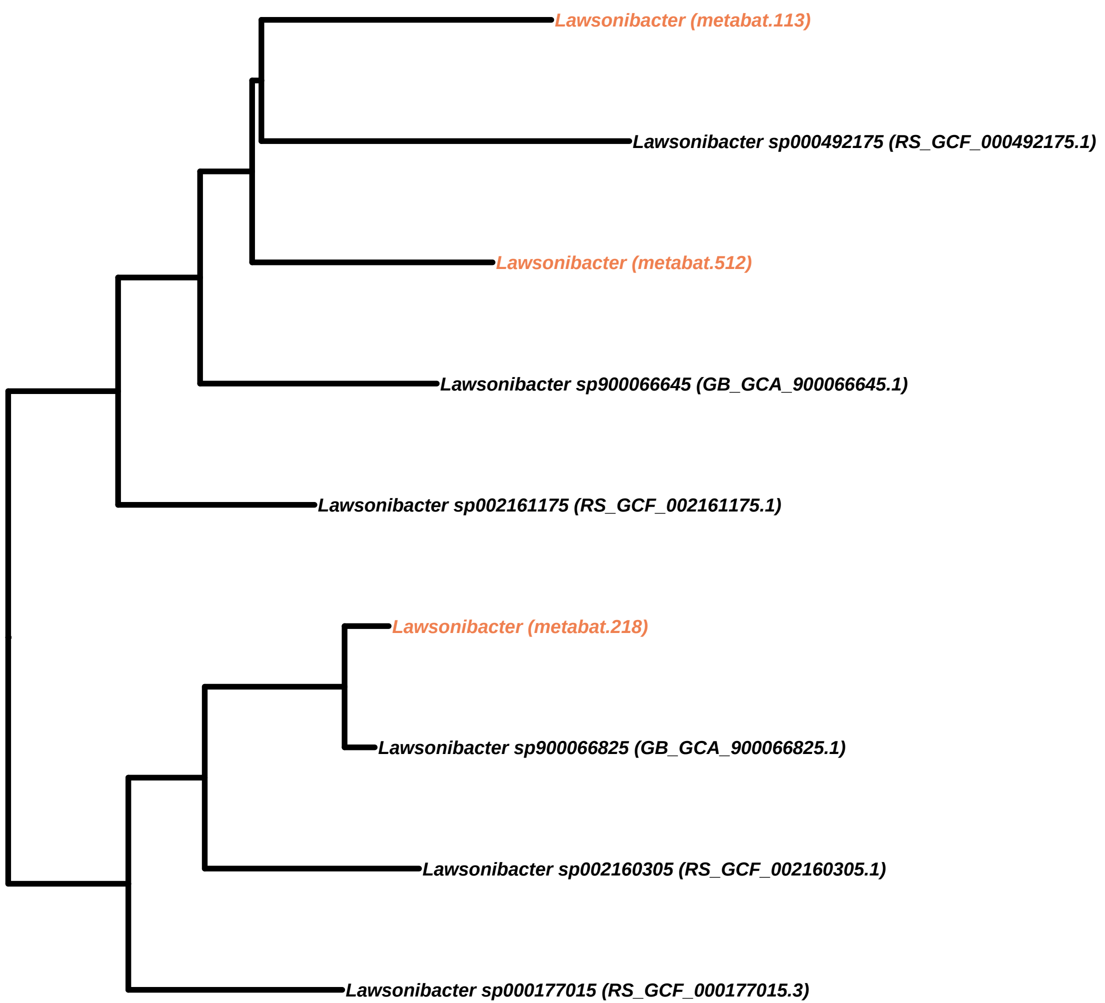
Lawsonibacter (metabat.113)
Lawsonibacter sp000492175 (RS_GCF_000492175.1)
Lawsonibacter (metabat.512)
Lawsonibacter sp900066645 (GB_GCA_900066645.1)
Lawsonibacter sp002161175 (RS_GCF_002161175.1)
Lawsonibacter (metabat.218)
Lawsonibacter sp900066825 (GB_GCA_900066825.1)
Lawsonibacter sp002160305 (RS_GCF_002160305.1)
Lawsonibacter sp000177015 (RS_GCF_000177015.3)

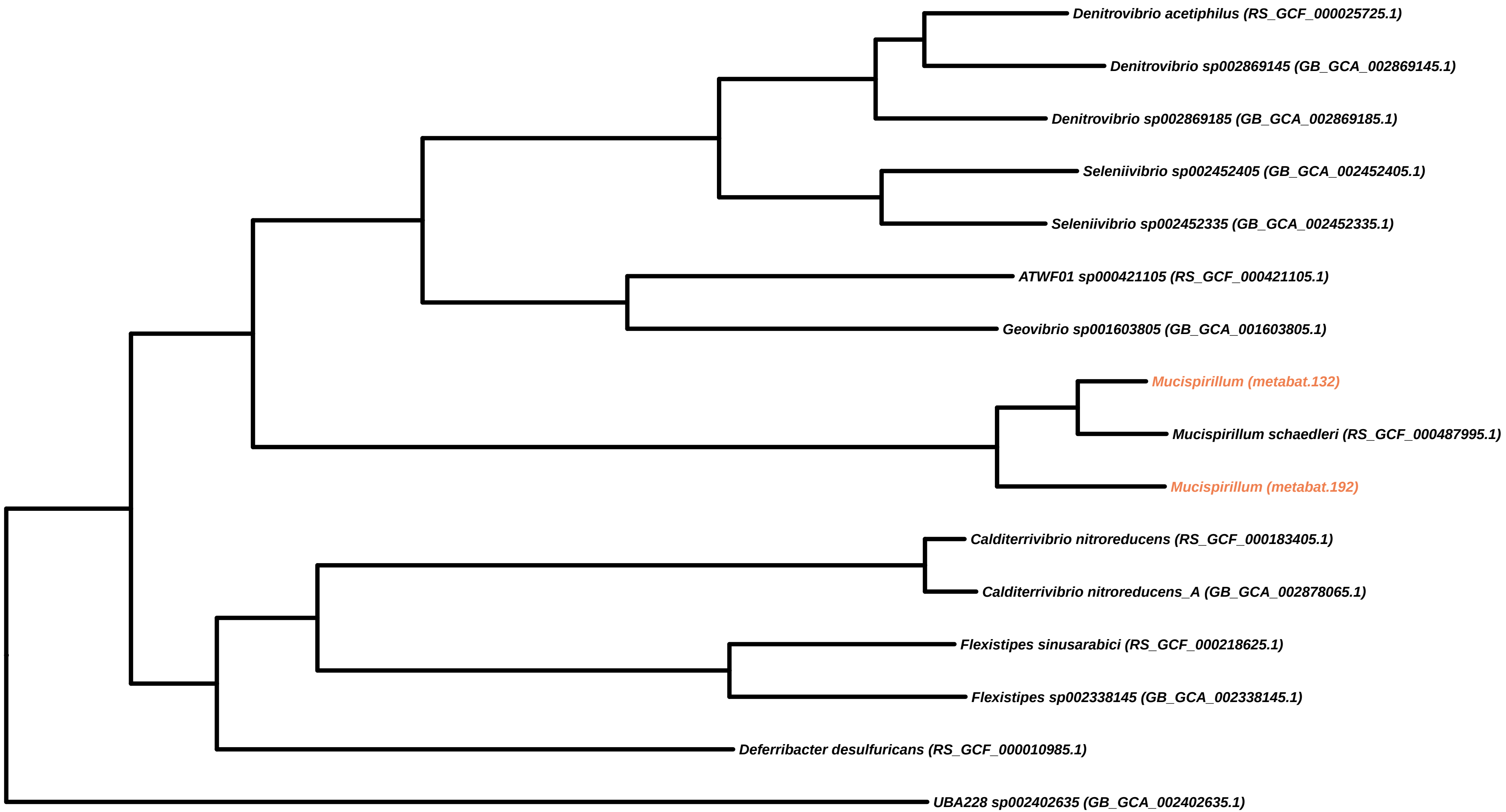

Denitrovibrio acetiphilus (RS_GCF_000025725.1)
Denitrovibrio sp002869145 (GB_GCA_002869145.1)
Denitrovibrio sp002869185 (GB_GCA_002869185.1)
Seleniivibrio sp002452405 (GB_GCA_002452405.1)
Seleniivibrio sp002452335 (GB_GCA_002452335.1)
ATWF01 sp000421105 (RS_GCF_000421105.1)
Geovibrio sp001603805 (GB_GCA_001603805.1)
Mucispirillum (metabat.132)
Mucispirillum schaedleri (RS_GCF_000487995.1)
Mucispirillum (metabat.192)
Calditerrivibrio nitroreducens (RS_GCF_000183405.1)
Calditerrivibrio nitroreducens_A (GB_GCA_002878065.1)
Flexistipes sinusarabici (RS_GCF_000218625.1)
Flexistipes sp002338145 (GB_GCA_002338145.1)
Deferribacter desulfuricans (RS_GCF_000010985.1)
UBA228 sp002402635 (GB_GCA_002402635.1)

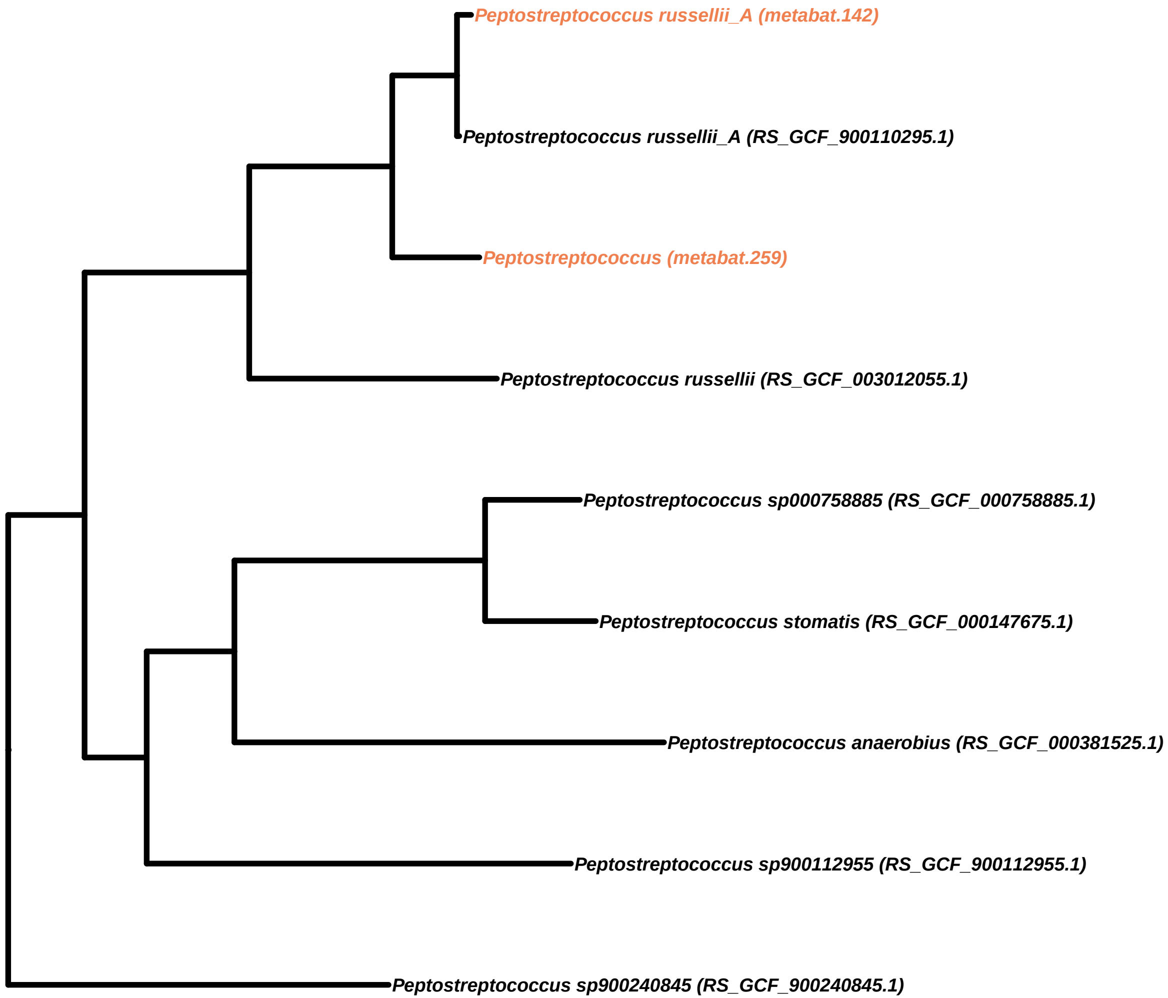
Peptostreptococcus russellii_A (metabat.142)
Peptostreptococcus russellii_A (RS_GCF_900110295.1)
Peptostreptococcus (metabat.259)
Peptostreptococcus russellii (RS_GCF_003012055.1)
Peptostreptococcus sp000758885 (RS_GCF_000758885.1)
Peptostreptococcus stomatis (RS_GCF_000147675.1)
Peptostreptococcus anaerobius (RS_GCF_000381525.1)
Peptostreptococcus sp900112955 (RS_GCF_900112955.1)
Peptostreptococcus sp900240845 (RS_GCF_900240845.1)

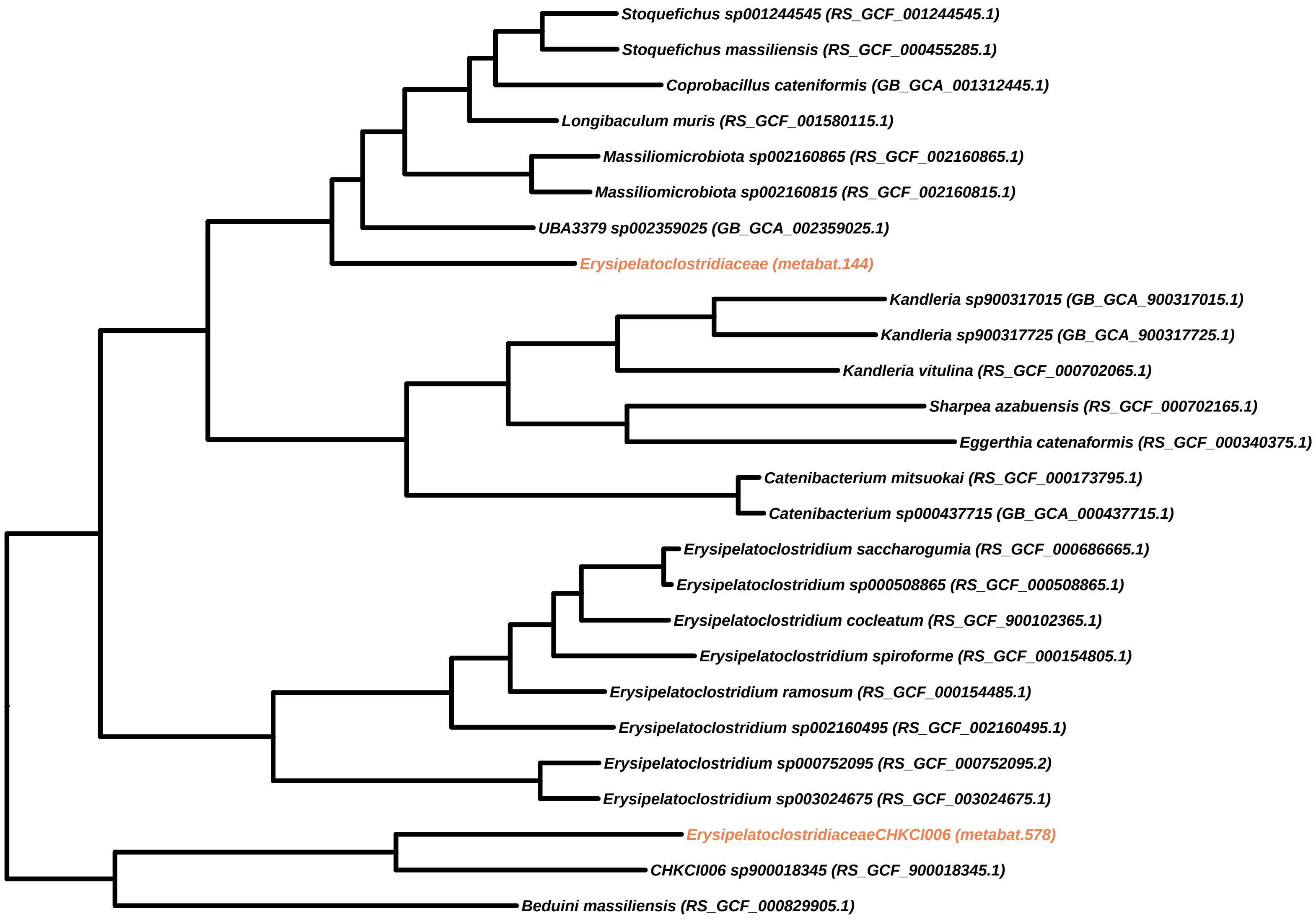

Stoquefichus sp001244545 (RS_GCF_001244545.1)
Stoquefichus massiliensis (RS_GCF_000455285.1)
Coprobacillus cateniformis (GB_GCA_001312445.1)
Longibaculum muris (RS_GCF_001580115.1)
Massiliomicrobiota sp002160865 (RS_GCF_002160865.1)
Massiliomicrobiota sp002160815 (RS_GCF_002160815.1)
UBA3379 sp002359025 (GB_GCA_002359025.1)
Erysipelatoclostridiaceae (metabat.144)
Kandleria sp900317015 (GB_GCA_900317015.1)
Kandleria sp900317725 (GB_GCA_900317725.1)
Kandleria vitulina (RS_GCF_000702065.1)
Sharpea azabuensis (RS_GCF_000702165.1)
Eggerthia catenaformis (RS_GCF_000340375.1)
Catenibacterium mitsuokai (RS_GCF_000173795.1)
Catenibacterium sp000437715 (GB_GCA_000437715.1)
Erysipelatoclostridium saccharogumia (RS_GCF_000686665.1)
Erysipelatoclostridium sp000508865 (RS_GCF_000508865.1)
Erysipelatoclostridium cocleatum (RS_GCF_900102365.1)
Erysipelatoclostridium spiroforme (RS_GCF_000154805.1)
Erysipelatoclostridium ramosum (RS_GCF_000154485.1)
Erysipelatoclostridium sp002160495 (RS_GCF_002160495.1)
Erysipelatoclostridium sp000752095 (RS_GCF_000752095.2)
Erysipelatoclostridium sp003024675 (RS_GCF_003024675.1)
ErysipelatoclostridiaceaeCHKCI006 (metabat.578)
CHKCI006 sp900018345 (RS_GCF_900018345.1)
Beduini massiliensis (RS_GCF_000829905.1)

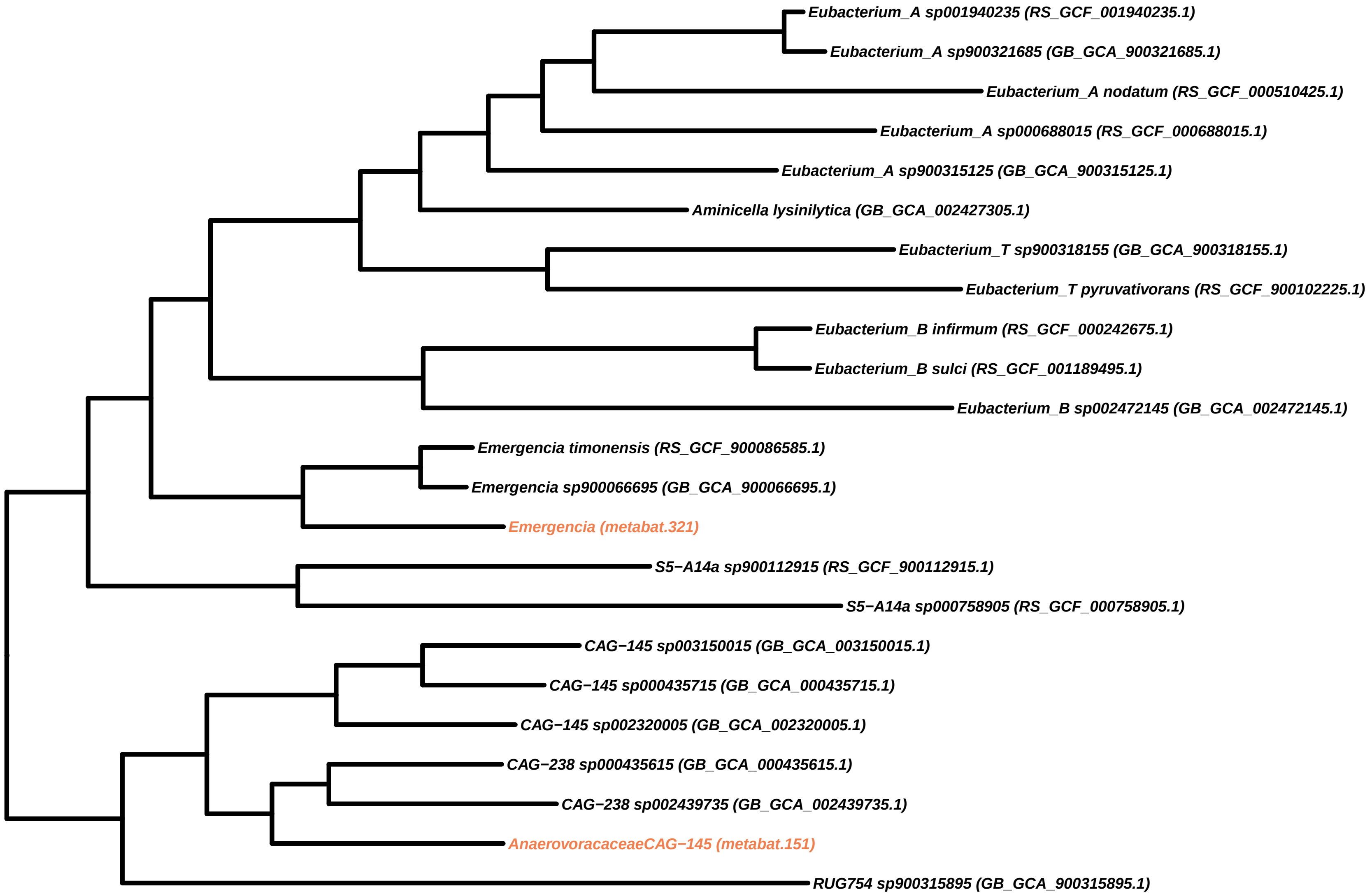

Eubacterium_A sp001940235 (RS_GCF_001940235.1)
Eubacterium_A sp900321685 (GB_GCA_900321685.1)
Eubacterium_A nodatum (RS_GCF_000510425.1)
Eubacterium_A sp000688015 (RS_GCF_000688015.1)
Eubacterium_A sp900315125 (GB_GCA_900315125.1)
Aminicella lysinilytica (GB_GCA_002427305.1)
Eubacterium_T sp900318155 (GB_GCA_900318155.1)
Eubacterium_T pyruvativorans (RS_GCF_900102225.1)
Eubacterium_B infirmum (RS_GCF_000242675.1)
Eubacterium_B sulci (RS_GCF_001189495.1)
Eubacterium_B sp002472145 (GB_GCA_002472145.1)
Emergencia timonensis (RS_GCF_900086585.1)
Emergencia sp900066695 (GB_GCA_900066695.1)
Emergencia (metabat.321)
S5−A14a sp900112915 (RS_GCF_900112915.1)
S5−A14a sp000758905 (RS_GCF_000758905.1)
CAG−145 sp003150015 (GB_GCA_003150015.1)
CAG−145 sp000435715 (GB_GCA_000435715.1)
CAG−145 sp002320005 (GB_GCA_002320005.1)
CAG−238 sp000435615 (GB_GCA_000435615.1)
CAG−238 sp002439735 (GB_GCA_002439735.1)
AnaerovoracaceaeCAG−145 (metabat.151)
RUG754 sp900315895 (GB_GCA_900315895.1)

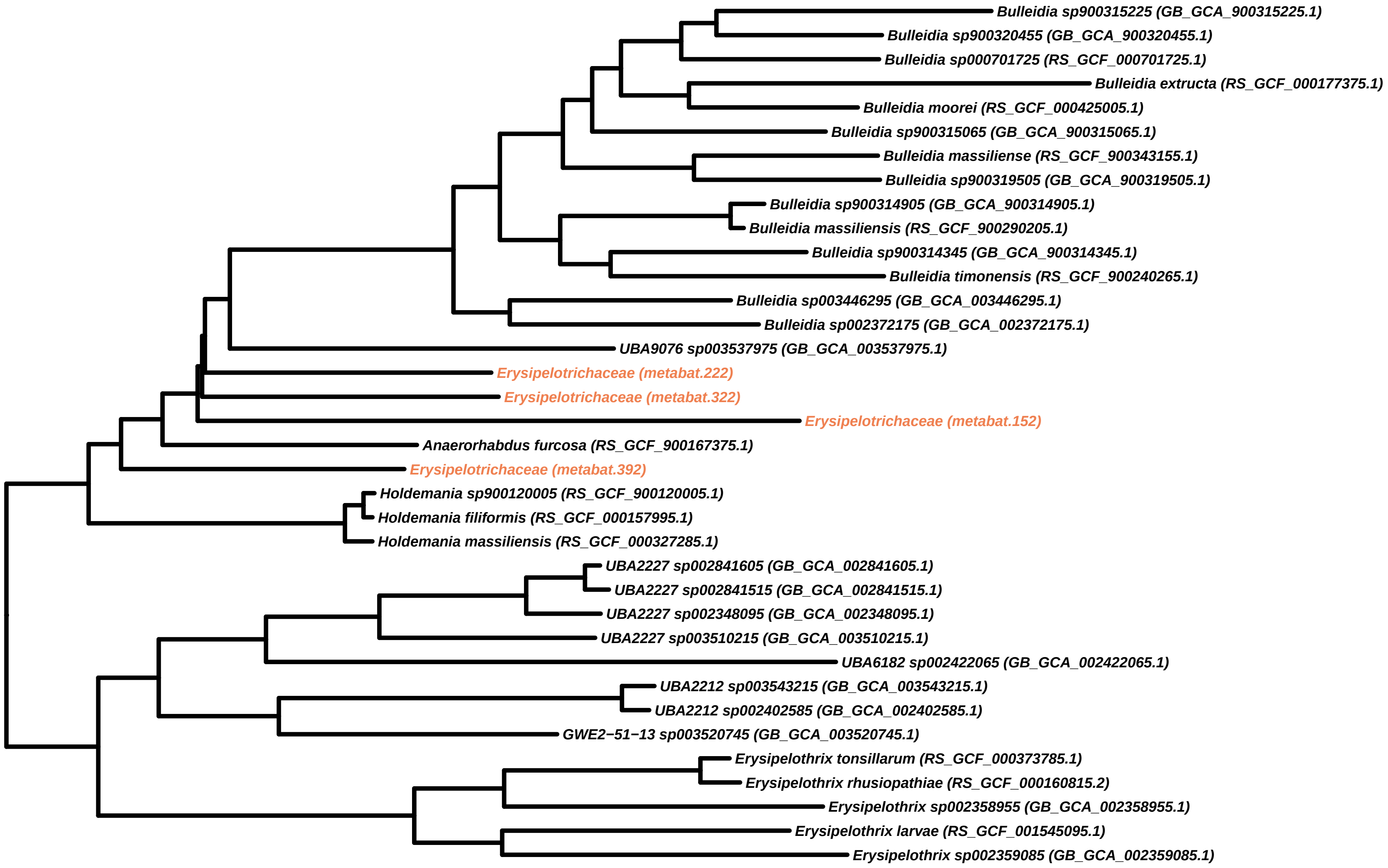
Bulleidia sp900315225 (GB_GCA_900315225.1)
Bulleidia sp900320455 (GB_GCA_900320455.1)
Bulleidia sp000701725 (RS_GCF_000701725.1)
Bulleidia extructa (RS_GCF_000177375.1)
Bulleidia moorei (RS_GCF_000425005.1)
Bulleidia sp900315065 (GB_GCA_900315065.1)
Bulleidia massiliense (RS_GCF_900343155.1)
Bulleidia sp900319505 (GB_GCA_900319505.1)
Bulleidia sp900314905 (GB_GCA_900314905.1)
Bulleidia massiliensis (RS_GCF_900290205.1)
Bulleidia sp900314345 (GB_GCA_900314345.1)
Bulleidia timonensis (RS_GCF_900240265.1)
Bulleidia sp003446295 (GB_GCA_003446295.1)
Bulleidia sp002372175 (GB_GCA_002372175.1)
UBA9076 sp003537975 (GB_GCA_003537975.1)
Erysipelotrichaceae (metabat.222)
Erysipelotrichaceae (metabat.322)
Erysipelotrichaceae (metabat.152)
Anaerorhabdus furcosa (RS_GCF_900167375.1)
Erysipelotrichaceae (metabat.392)
Holdemania sp900120005 (RS_GCF_900120005.1)
Holdemania filiformis (RS_GCF_000157995.1)
Holdemania massiliensis (RS_GCF_000327285.1)
UBA2227 sp002841605 (GB_GCA_002841605.1)
UBA2227 sp002841515 (GB_GCA_002841515.1)
UBA2227 sp002348095 (GB_GCA_002348095.1)
UBA2227 sp003510215 (GB_GCA_003510215.1)
UBA6182 sp002422065 (GB_GCA_002422065.1)
UBA2212 sp003543215 (GB_GCA_003543215.1)
UBA2212 sp002402585 (GB_GCA_002402585.1)
GWE2−51−13 sp003520745 (GB_GCA_003520745.1)
Erysipelothrix tonsillarum (RS_GCF_000373785.1)
Erysipelothrix rhusiopathiae (RS_GCF_000160815.2)
Erysipelothrix sp002358955 (GB_GCA_002358955.1)
Erysipelothrix larvae (RS_GCF_001545095.1)
Erysipelothrix sp002359085 (GB_GCA_002359085.1)

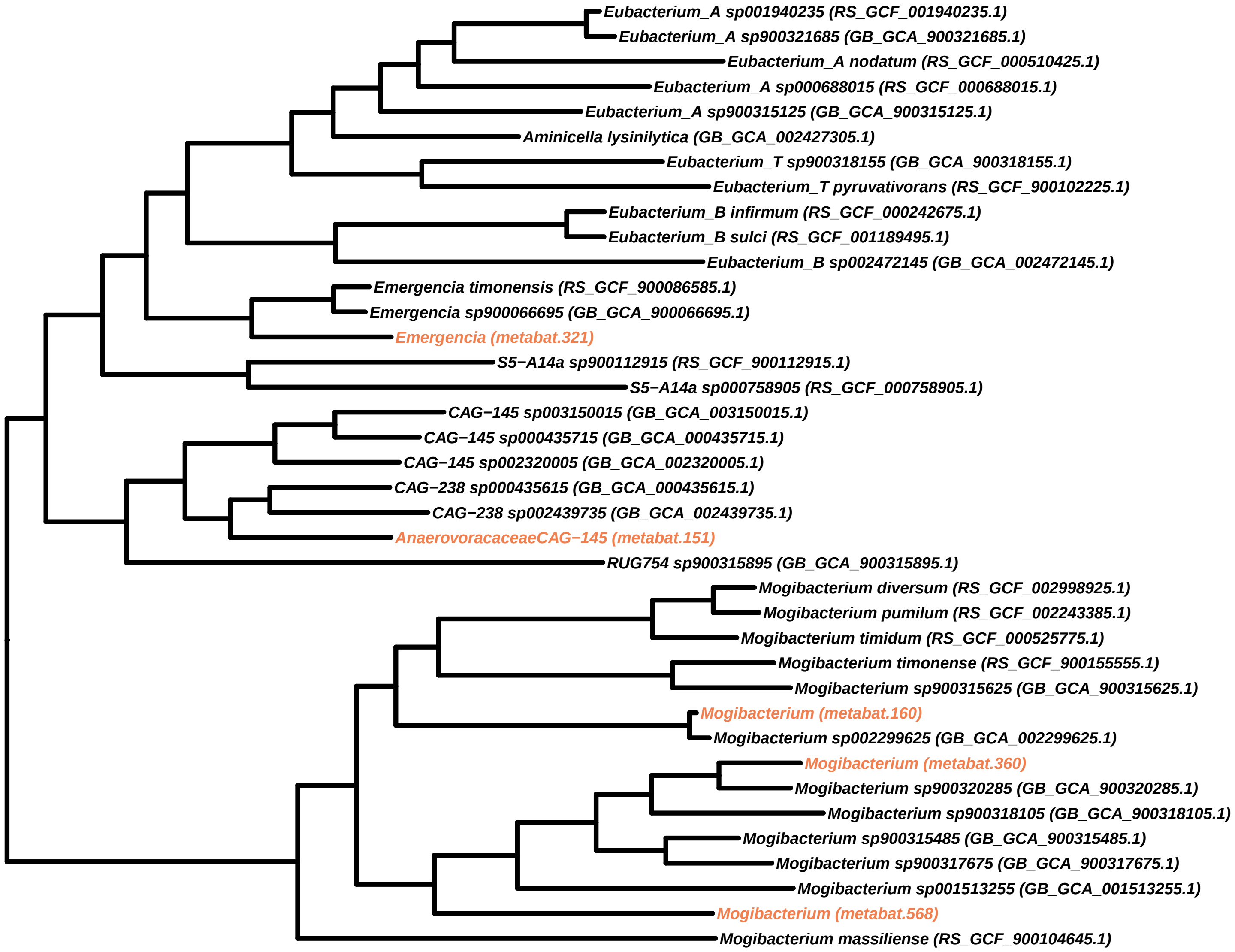
Eubacterium_A sp001940235 (RS_GCF_001940235.1)
Eubacterium_A sp900321685 (GB_GCA_900321685.1)
Eubacterium_A nodatum (RS_GCF_000510425.1)
Eubacterium_A sp000688015 (RS_GCF_000688015.1)
Eubacterium_A sp900315125 (GB_GCA_900315125.1)
Aminicella lysinilytica (GB_GCA_002427305.1)
Eubacterium_T sp900318155 (GB_GCA_900318155.1)
Eubacterium_T pyruvativorans (RS_GCF_900102225.1)
Eubacterium_B infirmum (RS_GCF_000242675.1)
Eubacterium_B sulci (RS_GCF_001189495.1)
Eubacterium_B sp002472145 (GB_GCA_002472145.1)
Emergencia timonensis (RS_GCF_900086585.1)
Emergencia sp900066695 (GB_GCA_900066695.1)
Emergencia (metabat.321)
S5−A14a sp900112915 (RS_GCF_900112915.1)
S5−A14a sp000758905 (RS_GCF_000758905.1)
CAG−145 sp003150015 (GB_GCA_003150015.1)
CAG−145 sp000435715 (GB_GCA_000435715.1)
CAG−145 sp002320005 (GB_GCA_002320005.1)
CAG−238 sp000435615 (GB_GCA_000435615.1)
CAG−238 sp002439735 (GB_GCA_002439735.1)
AnaerovoracaceaeCAG−145 (metabat.151)
RUG754 sp900315895 (GB_GCA_900315895.1)
Mogibacterium diversum (RS_GCF_002998925.1)
Mogibacterium pumilum (RS_GCF_002243385.1)
Mogibacterium timidum (RS_GCF_000525775.1)
Mogibacterium timonense (RS_GCF_900155555.1)
Mogibacterium sp900315625 (GB_GCA_900315625.1)
Mogibacterium (metabat.160)
Mogibacterium sp002299625 (GB_GCA_002299625.1)
Mogibacterium (metabat.360)
Mogibacterium sp900320285 (GB_GCA_900320285.1)
Mogibacterium sp900318105 (GB_GCA_900318105.1)
Mogibacterium sp900315485 (GB_GCA_900315485.1)
Mogibacterium sp900317675 (GB_GCA_900317675.1)
Mogibacterium sp001513255 (GB_GCA_001513255.1)
Mogibacterium (metabat.568)
Mogibacterium massiliense (RS_GCF_900104645.1)

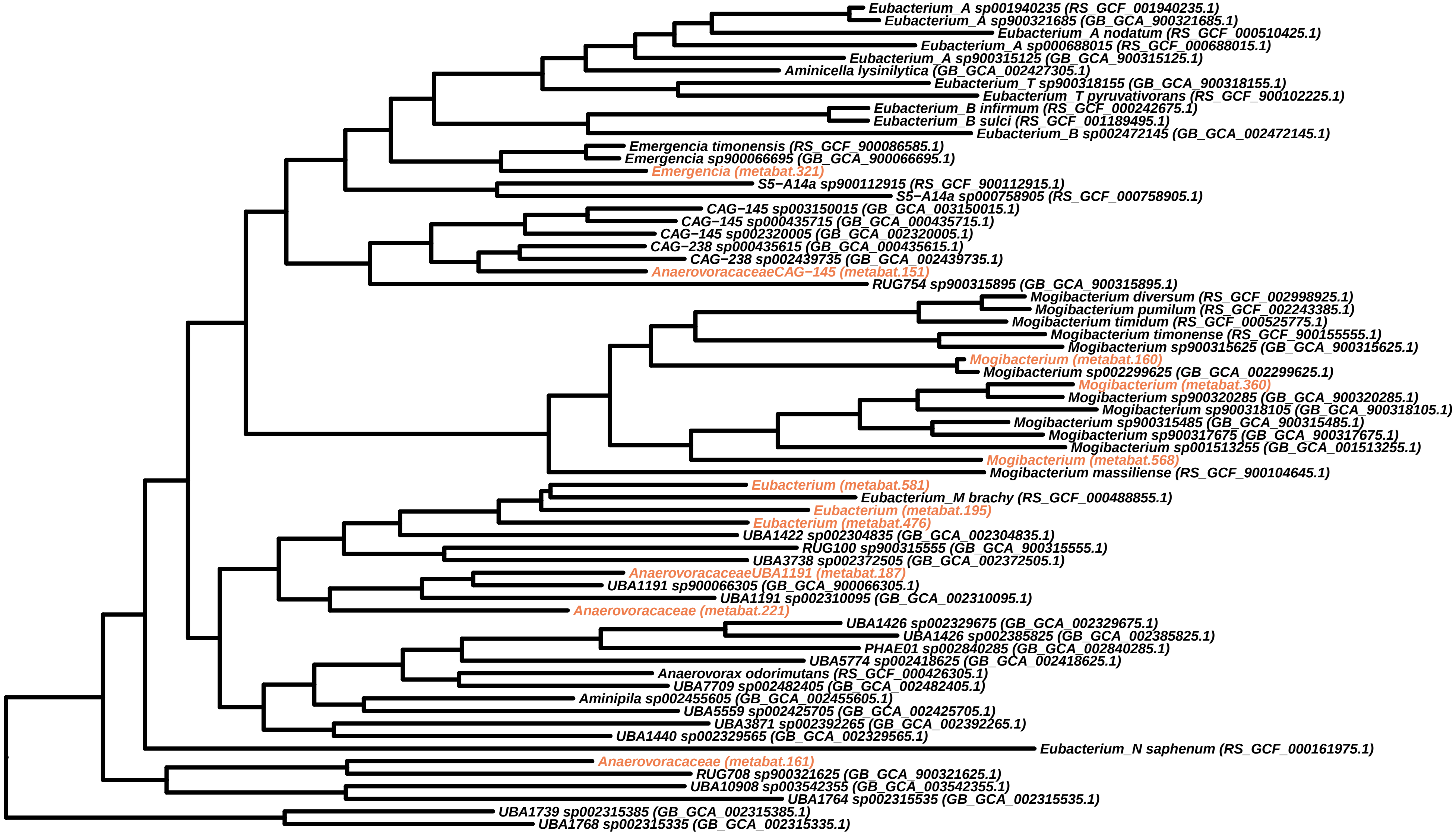
Eubacterium_A sp001940235 (RS_GCF_001940235.1)
Eubacterium_A sp900321685 (GB_GCA_900321685.1)
Eubacterium_A nodatum (RS_GCF_000510425.1)
Eubacterium_A sp000688015 (RS_GCF_000688015.1)
Eubacterium_A sp900315125 (GB_GCA_900315125.1)
Aminicella lysinilytica (GB_GCA_002427305.1)
Eubacterium_T sp900318155 (GB_GCA_900318155.1)
Eubacterium_T pyruvativorans (RS_GCF_900102225.1)
Eubacterium_B infirmum (RS_GCF_000242675.1)
Eubacterium_B sulci (RS_GCF_001189495.1)
Eubacterium_B sp002472145 (GB_GCA_002472145.1)
Emergencia timonensis (RS_GCF_900086585.1)
Emergencia sp900066695 (GB_GCA_900066695.1)
Emergencia (metabat.321)
S5−A14a sp900112915 (RS_GCF_900112915.1)
S5−A14a sp000758905 (RS_GCF_000758905.1)
CAG−145 sp003150015 (GB_GCA_003150015.1)
CAG−145 sp000435715 (GB_GCA_000435715.1)
CAG−145 sp002320005 (GB_GCA_002320005.1)
CAG−238 sp000435615 (GB_GCA_000435615.1)
CAG−238 sp002439735 (GB_GCA_002439735.1)
AnaerovoracaceaeCAG−145 (metabat.151)
RUG754 sp900315895 (GB_GCA_900315895.1)
Mogibacterium diversum (RS_GCF_002998925.1)
Mogibacterium pumilum (RS_GCF_002243385.1)
Mogibacterium timidum (RS_GCF_000525775.1)
Mogibacterium timonense (RS_GCF_900155555.1)
Mogibacterium sp900315625 (GB_GCA_900315625.1)
Mogibacterium (metabat.160)
Mogibacterium sp002299625 (GB_GCA_002299625.1)
Mogibacterium (metabat.360)
Mogibacterium sp900320285 (GB_GCA_900320285.1)
Mogibacterium sp900318105 (GB_GCA_900318105.1)
Mogibacterium sp900315485 (GB_GCA_900315485.1)
Mogibacterium sp900317675 (GB_GCA_900317675.1)
Mogibacterium sp001513255 (GB_GCA_001513255.1)
Mogibacterium (metabat.568)
Mogibacterium massiliense (RS_GCF_900104645.1)
Eubacterium (metabat.581)
Eubacterium_M brachy (RS_GCF_000488855.1)
Eubacterium (metabat.195)
Eubacterium (metabat.476)
UBA1422 sp002304835 (GB_GCA_002304835.1)
RUG100 sp900315555 (GB_GCA_900315555.1)
UBA3738 sp002372505 (GB_GCA_002372505.1)
AnaerovoracaceaeUBA1191 (metabat.187)
UBA1191 sp900066305 (GB_GCA_900066305.1)
UBA1191 sp002310095 (GB_GCA_002310095.1)
Anaerovoracaceae (metabat.221)
UBA1426 sp002329675 (GB_GCA_002329675.1)
UBA1426 sp002385825 (GB_GCA_002385825.1)
PHAE01 sp002840285 (GB_GCA_002840285.1)
UBA5774 sp002418625 (GB_GCA_002418625.1)
Anaerovorax odorimutans (RS_GCF_000426305.1)
UBA7709 sp002482405 (GB_GCA_002482405.1)
Aminipila sp002455605 (GB_GCA_002455605.1)
UBA5559 sp002425705 (GB_GCA_002425705.1)
UBA3871 sp002392265 (GB_GCA_002392265.1)
UBA1440 sp002329565 (GB_GCA_002329565.1)
Eubacterium_N saphenum (RS_GCF_000161975.1)
Anaerovoracaceae (metabat.161)
RUG708 sp900321625 (GB_GCA_900321625.1)
UBA10908 sp003542355 (GB_GCA_003542355.1)
UBA1764 sp002315535 (GB_GCA_002315535.1)
UBA1739 sp002315385 (GB_GCA_002315385.1)
UBA1768 sp002315335 (GB_GCA_002315335.1)

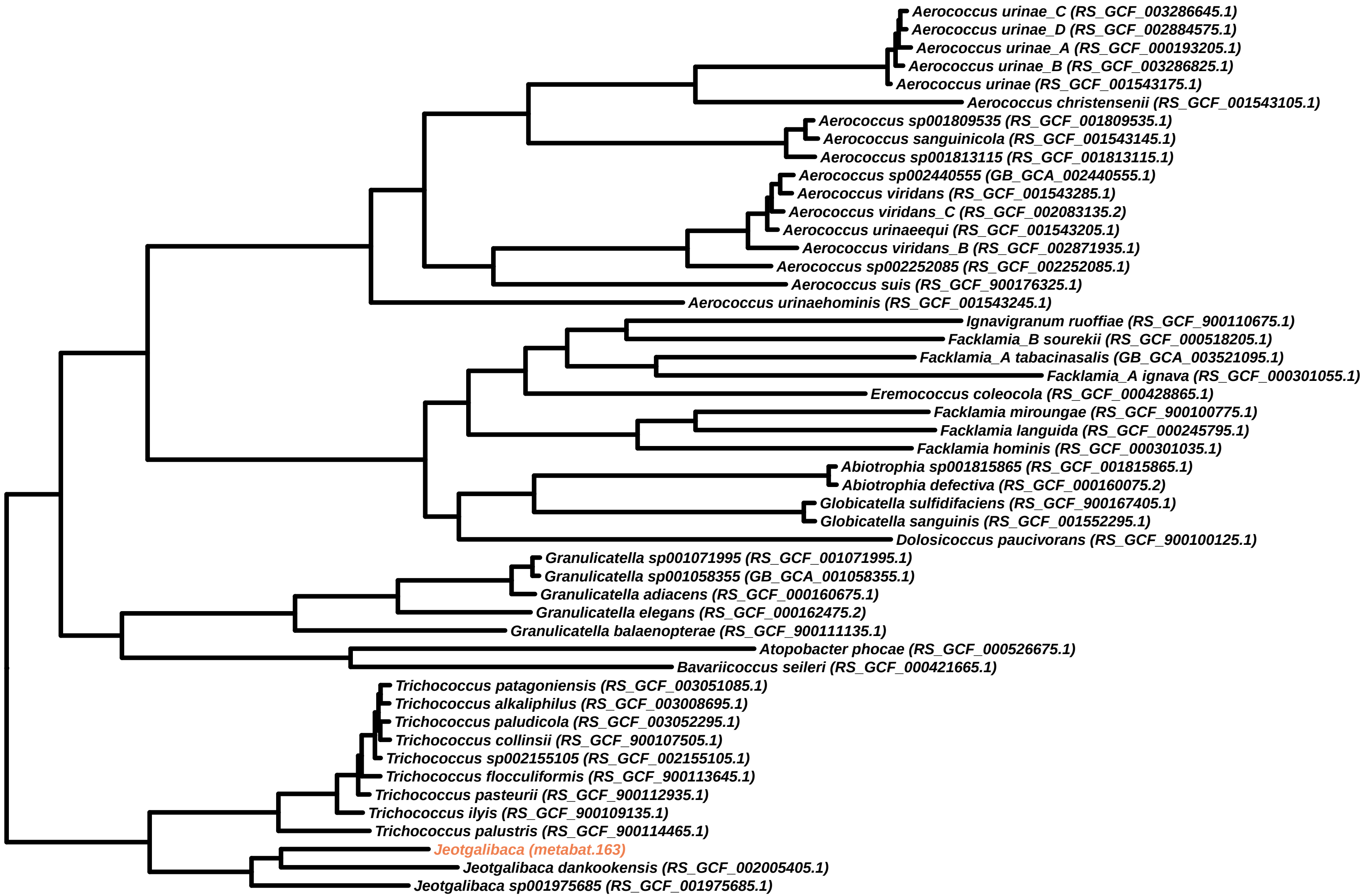

Aerococcus urinae_C (RS_GCF_003286645.1)
Aerococcus urinae_D (RS_GCF_002884575.1)
Aerococcus urinae_A (RS_GCF_000193205.1)
Aerococcus urinae_B (RS_GCF_003286825.1)
Aerococcus urinae (RS_GCF_001543175.1)
Aerococcus christensenii (RS_GCF_001543105.1)
Aerococcus sp001809535 (RS_GCF_001809535.1)
Aerococcus sanguinicola (RS_GCF_001543145.1)
Aerococcus sp001813115 (RS_GCF_001813115.1)
Aerococcus sp002440555 (GB_GCA_002440555.1)
Aerococcus viridans (RS_GCF_001543285.1)
Aerococcus viridans_C (RS_GCF_002083135.2)
Aerococcus urinaeequi (RS_GCF_001543205.1)
Aerococcus viridans_B (RS_GCF_002871935.1)
Aerococcus sp002252085 (RS_GCF_002252085.1)
Aerococcus suis (RS_GCF_900176325.1)
Aerococcus urinaehominis (RS_GCF_001543245.1)
Ignavigranum ruoffiae (RS_GCF_900110675.1)
Facklamia_B sourekii (RS_GCF_000518205.1)
Facklamia_A tabacinasalis (GB_GCA_003521095.1)
Facklamia_A ignava (RS_GCF_000301055.1)
Eremococcus coleocola (RS_GCF_000428865.1)
Facklamia miroungae (RS_GCF_900100775.1)
Facklamia languida (RS_GCF_000245795.1)
Facklamia hominis (RS_GCF_000301035.1)
Abiotrophia sp001815865 (RS_GCF_001815865.1)
Abiotrophia defectiva (RS_GCF_000160075.2)
Globicatella sulfidifaciens (RS_GCF_900167405.1)
Globicatella sanguinis (RS_GCF_001552295.1)
Dolosicoccus paucivorans (RS_GCF_900100125.1)
Granulicatella sp001071995 (RS_GCF_001071995.1)
Granulicatella sp001058355 (GB_GCA_001058355.1)
Granulicatella adiacens (RS_GCF_000160675.1)
Granulicatella elegans (RS_GCF_000162475.2)
Granulicatella balaenopterae (RS_GCF_900111135.1)
Atopobacter phocae (RS_GCF_000526675.1)
Bavariicoccus seileri (RS_GCF_000421665.1)
Trichococcus patagoniensis (RS_GCF_003051085.1)
Trichococcus alkaliphilus (RS_GCF_003008695.1)
Trichococcus paludicola (RS_GCF_003052295.1)
Trichococcus collinsii (RS_GCF_900107505.1)
Trichococcus sp002155105 (RS_GCF_002155105.1)
Trichococcus flocculiformis (RS_GCF_900113645.1)
Trichococcus pasteurii (RS_GCF_900112935.1)
Trichococcus ilyis (RS_GCF_900109135.1)
Trichococcus palustris (RS_GCF_900114465.1)
Jeotgalibaca (metabat.163)
Jeotgalibaca dankookensis (RS_GCF_002005405.1)
Jeotgalibaca sp001975685 (RS_GCF_001975685.1)

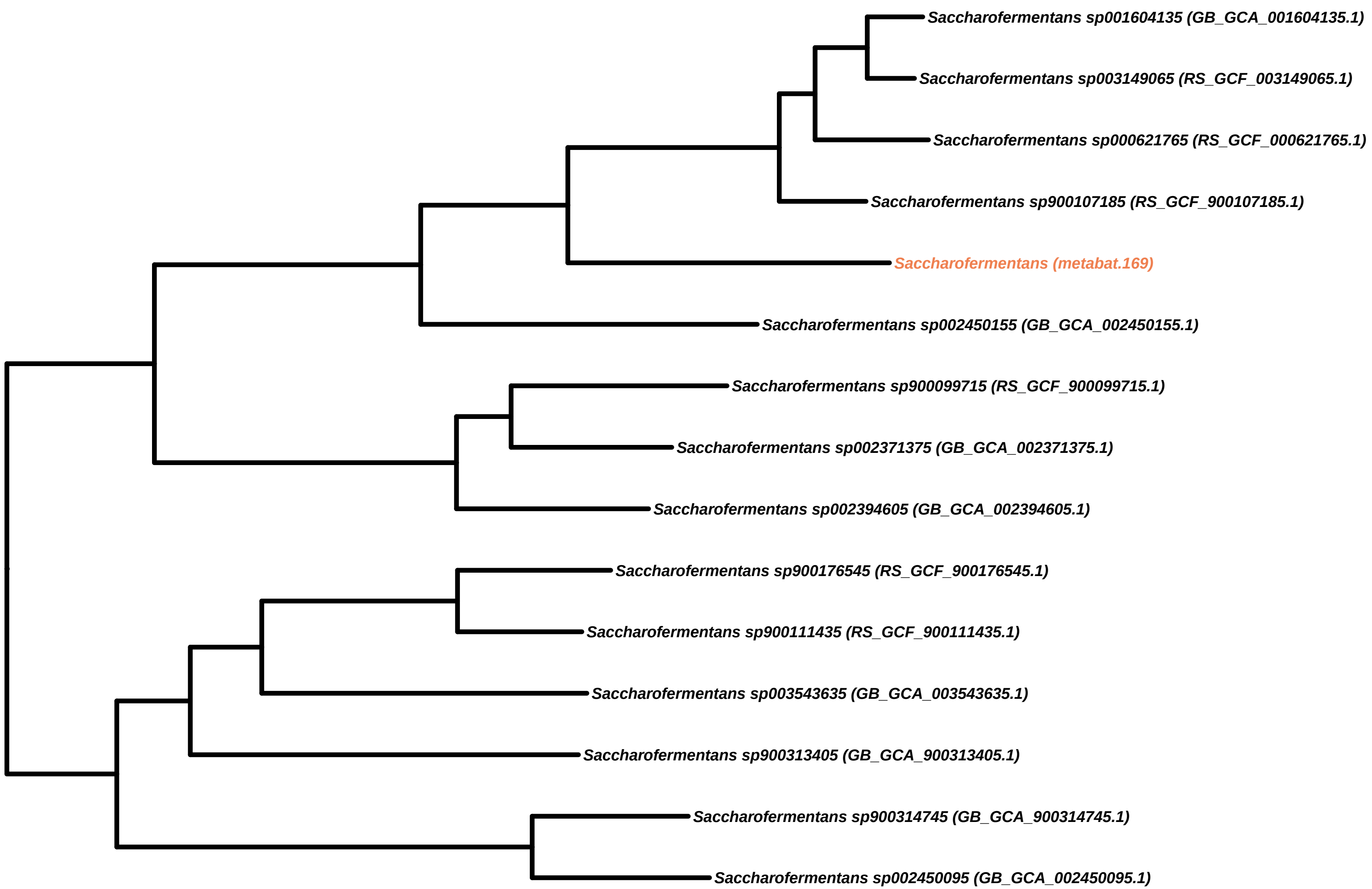
Saccharofermentans sp001604135 (GB_GCA_001604135.1)
Saccharofermentans sp003149065 (RS_GCF_003149065.1)
Saccharofermentans sp000621765 (RS_GCF_000621765.1)
Saccharofermentans sp900107185 (RS_GCF_900107185.1)
Saccharofermentans (metabat.169)
Saccharofermentans sp002450155 (GB_GCA_002450155.1)
Saccharofermentans sp900099715 (RS_GCF_900099715.1)
Saccharofermentans sp002371375 (GB_GCA_002371375.1)
Saccharofermentans sp002394605 (GB_GCA_002394605.1)
Saccharofermentans sp900176545 (RS_GCF_900176545.1)
Saccharofermentans sp900111435 (RS_GCF_900111435.1)
Saccharofermentans sp003543635 (GB_GCA_003543635.1)
Saccharofermentans sp900313405 (GB_GCA_900313405.1)
Saccharofermentans sp900314745 (GB_GCA_900314745.1)
Saccharofermentans sp002450095 (GB_GCA_002450095.1)

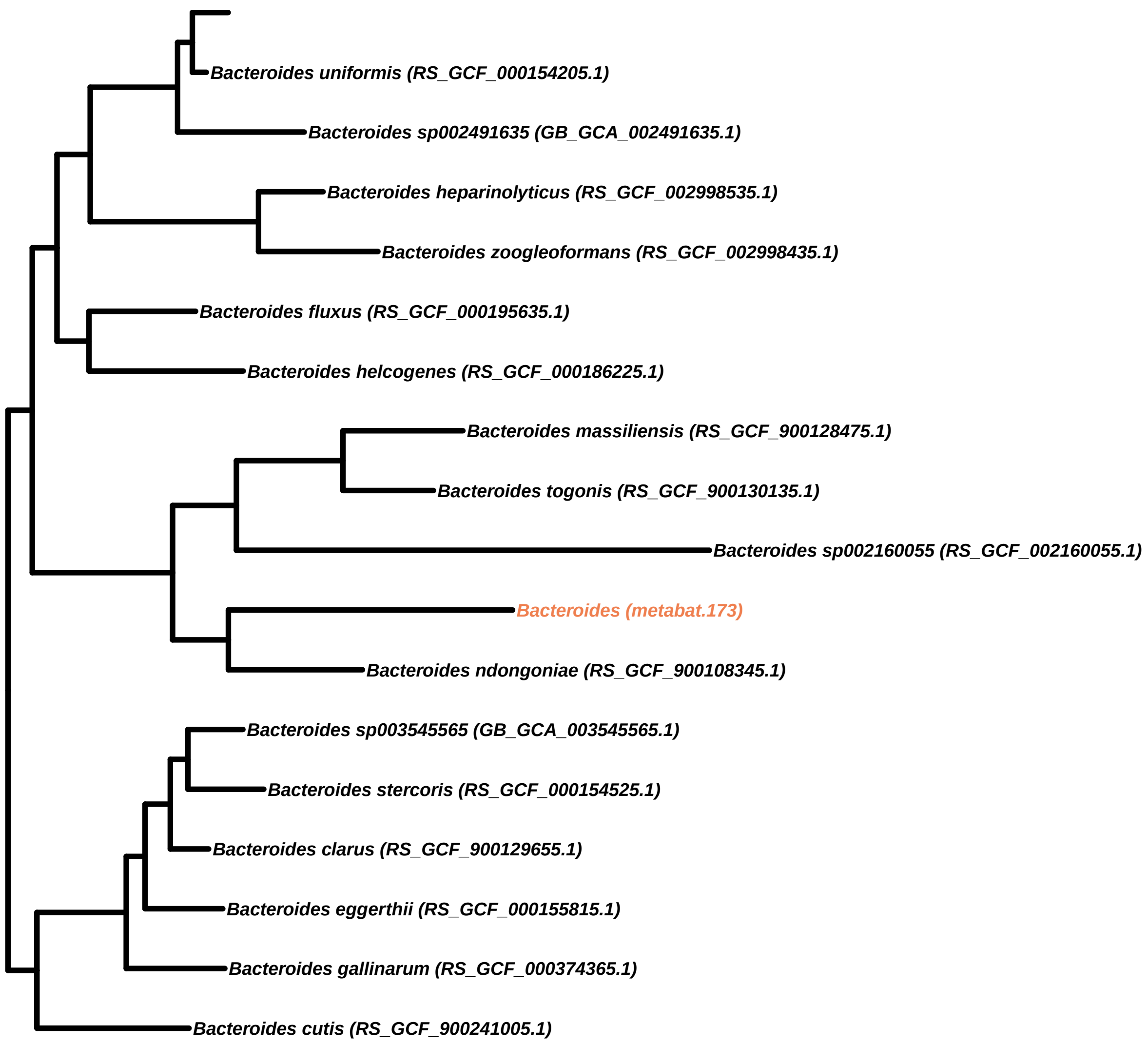
Bacteroides uniformis (RS_GCF_000154205.1)
Bacteroides sp002491635 (GB_GCA_002491635.1)
Bacteroides heparinolyticus (RS_GCF_002998535.1)
Bacteroides zoogleoformans (RS_GCF_002998435.1)
Bacteroides fluxus (RS_GCF_000195635.1)
Bacteroides helcogenes (RS_GCF_000186225.1)
Bacteroides massiliensis (RS_GCF_900128475.1)
Bacteroides togonis (RS_GCF_900130135.1)
Bacteroides sp002160055 (RS_GCF_002160055.1)
Bacteroides (metabat.173)
Bacteroides ndongoniae (RS_GCF_900108345.1)
Bacteroides sp003545565 (GB_GCA_003545565.1)
Bacteroides stercoris (RS_GCF_000154525.1)
Bacteroides clarus (RS_GCF_900129655.1)
Bacteroides eggerthii (RS_GCF_000155815.1)
Bacteroides gallinarum (RS_GCF_000374365.1)
Bacteroides cutis (RS_GCF_900241005.1)

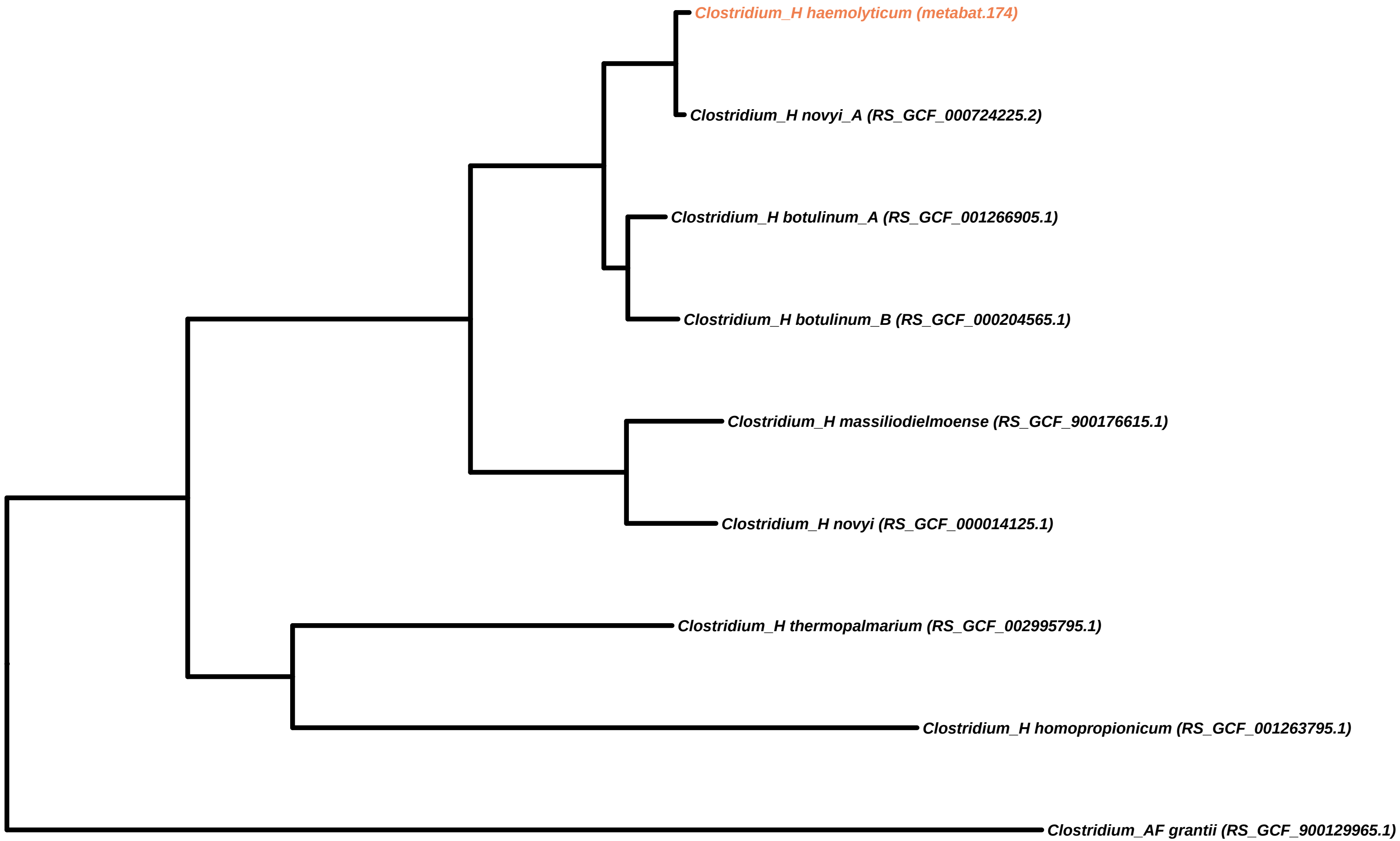

Clostridium_H haemolyticum (metabat.174)
Clostridium_H novyi_A (RS_GCF_000724225.2)
Clostridium_H botulinum_A (RS_GCF_001266905.1)
Clostridium_H botulinum_B (RS_GCF_000204565.1)
Clostridium_H massiliodielmoense (RS_GCF_900176615.1)
Clostridium_H novyi (RS_GCF_000014125.1)
Clostridium_H thermopalmarium (RS_GCF_002995795.1)
Clostridium_H homopropionicum (RS_GCF_001263795.1)
Clostridium_AF grantii (RS_GCF_900129965.1)

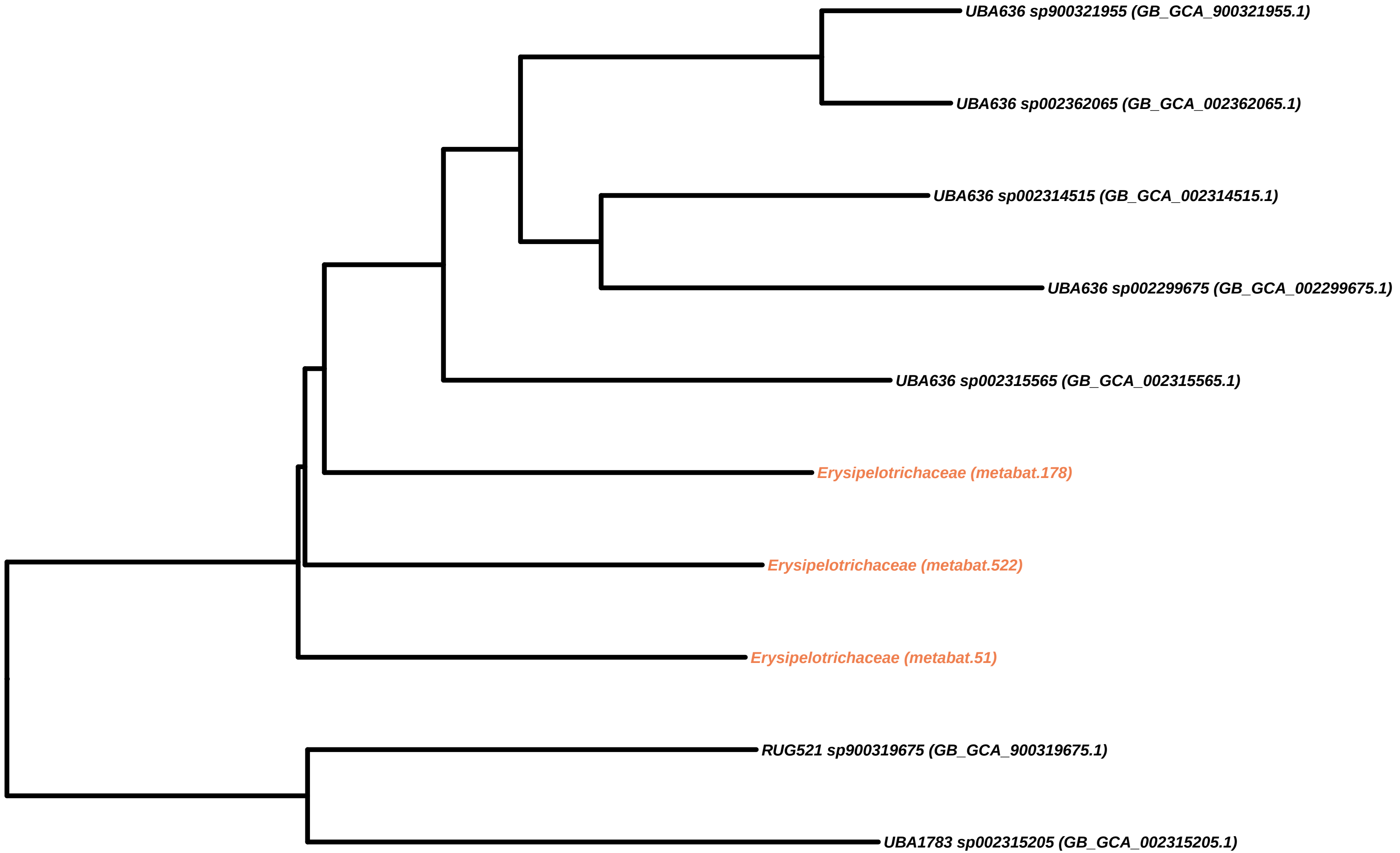
UBA636 sp900321955 (GB_GCA_900321955.1)
UBA636 sp002362065 (GB_GCA_002362065.1)
UBA636 sp002314515 (GB_GCA_002314515.1)
UBA636 sp002299675 (GB_GCA_002299675.1)
UBA636 sp002315565 (GB_GCA_002315565.1)
Erysipelotrichaceae (metabat.178)
Erysipelotrichaceae (metabat.522)
Erysipelotrichaceae (metabat.51)
RUG521 sp900319675 (GB_GCA_900319675.1)
UBA1783 sp002315205 (GB_GCA_002315205.1)

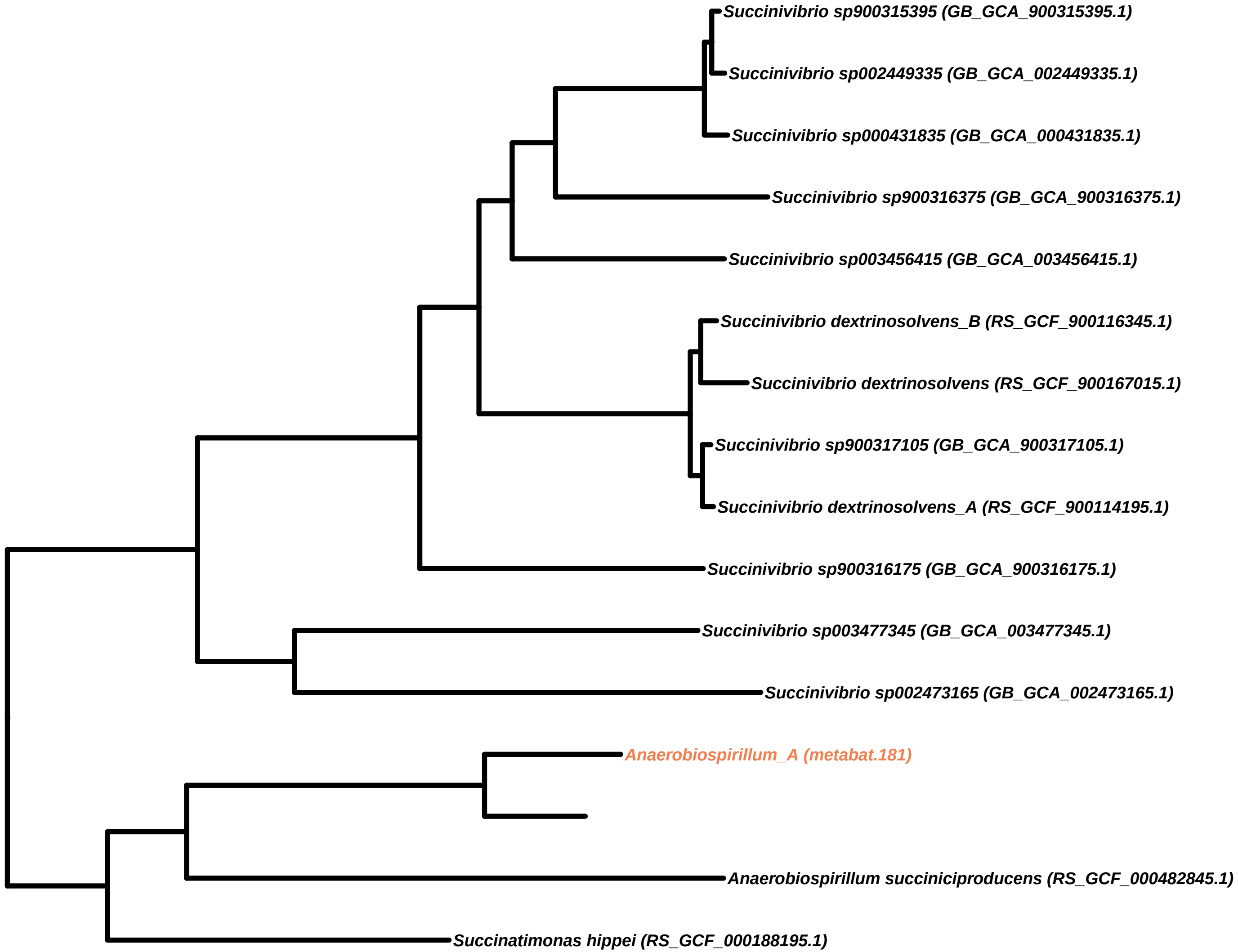

Succinivibrio sp900315395 (GB_GCA_900315395.1)
Succinivibrio sp002449335 (GB_GCA_002449335.1)
Succinivibrio sp000431835 (GB_GCA_000431835.1)
Succinivibrio sp900316375 (GB_GCA_900316375.1)
Succinivibrio sp003456415 (GB_GCA_003456415.1)
Succinivibrio dextrinosolvens_B (RS_GCF_900116345.1)
Succinivibrio dextrinosolvens (RS_GCF_900167015.1)
Succinivibrio sp900317105 (GB_GCA_900317105.1)
Succinivibrio dextrinosolvens_A (RS_GCF_900114195.1)
Succinivibrio sp900316175 (GB_GCA_900316175.1)
Succinivibrio sp003477345 (GB_GCA_003477345.1)
Succinivibrio sp002473165 (GB_GCA_002473165.1)
Anaerobiospirillum_A (metabat.181)
Anaerobiospirillum succiniciproducens (RS_GCF_000482845.1)
Succinatimonas hippei (RS_GCF_000188195.1)

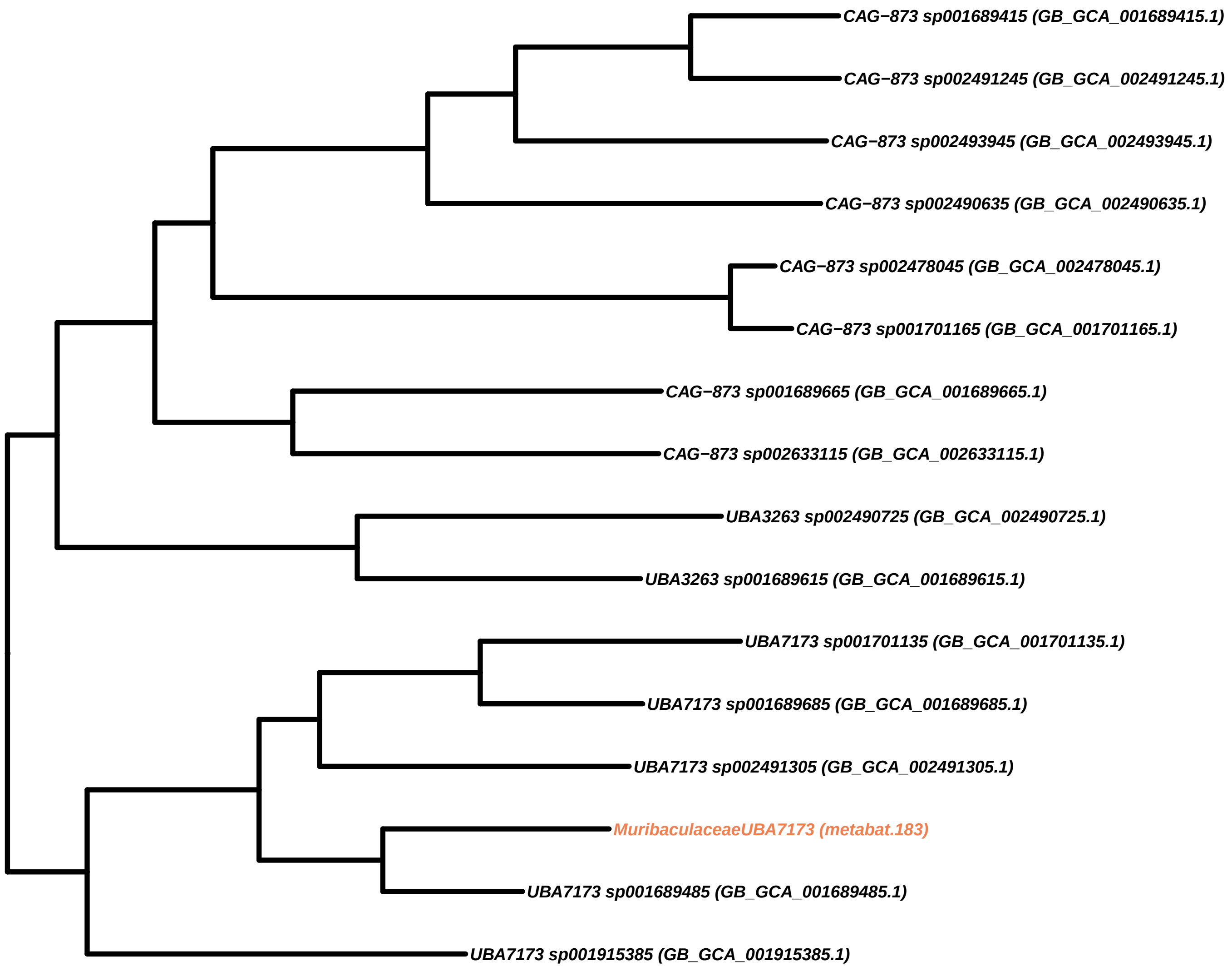
CAG−873 sp001689415 (GB_GCA_001689415.1)
CAG−873 sp002491245 (GB_GCA_002491245.1)
CAG−873 sp002493945 (GB_GCA_002493945.1)
CAG−873 sp002490635 (GB_GCA_002490635.1)
CAG−873 sp002478045 (GB_GCA_002478045.1)
CAG−873 sp001701165 (GB_GCA_001701165.1)
CAG−873 sp001689665 (GB_GCA_001689665.1)
CAG−873 sp002633115 (GB_GCA_002633115.1)
UBA3263 sp002490725 (GB_GCA_002490725.1)
UBA3263 sp001689615 (GB_GCA_001689615.1)
UBA7173 sp001701135 (GB_GCA_001701135.1)
UBA7173 sp001689685 (GB_GCA_001689685.1)
UBA7173 sp002491305 (GB_GCA_002491305.1)
MuribaculaceaeUBA7173 (metabat.183)
UBA7173 sp001689485 (GB_GCA_001689485.1)
UBA7173 sp001915385 (GB_GCA_001915385.1)

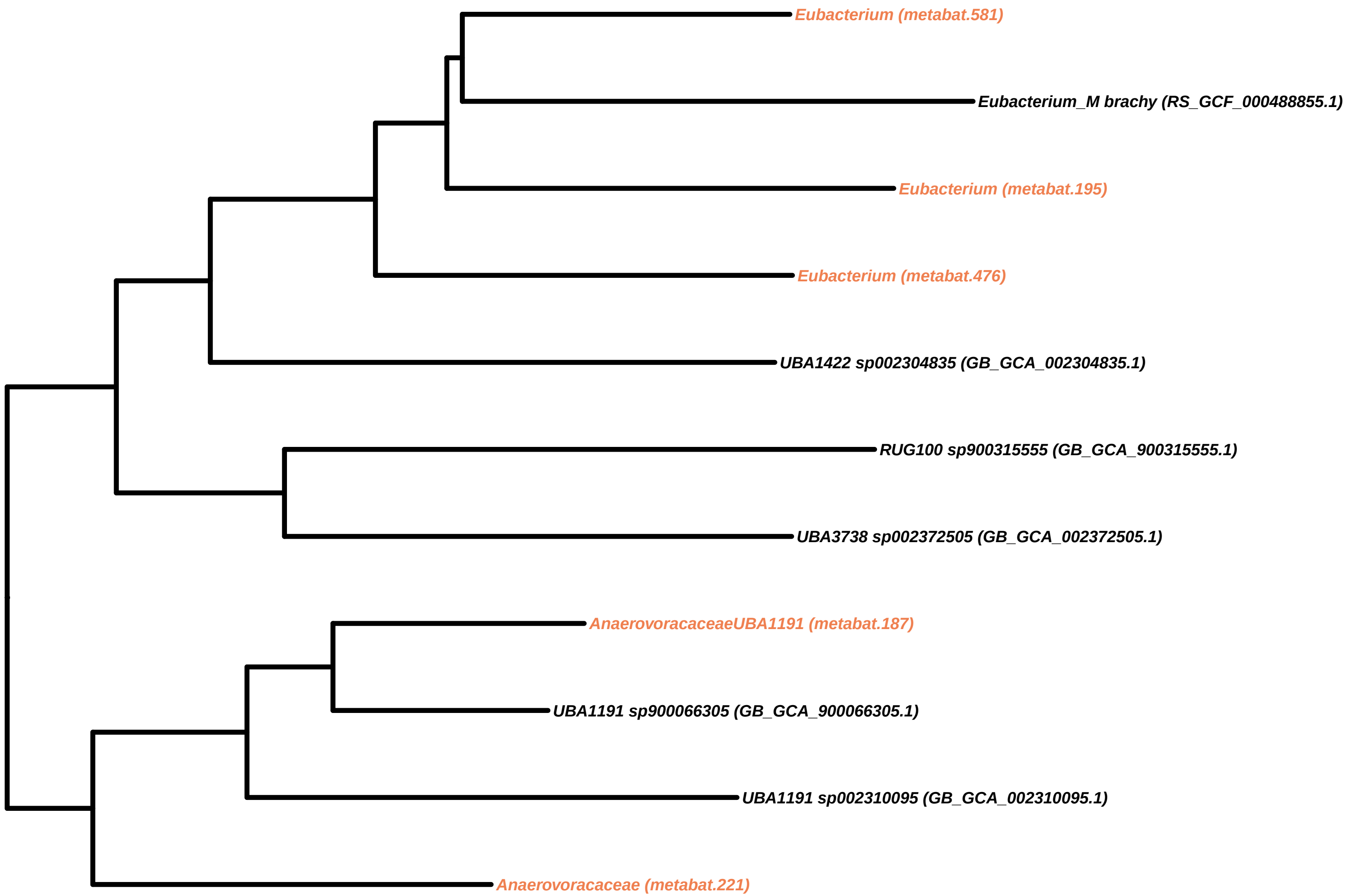
Eubacterium (metabat.581)
Eubacterium_M brachy (RS_GCF_000488855.1)
Eubacterium (metabat.195)
Eubacterium (metabat.476)
UBA1422 sp002304835 (GB_GCA_002304835.1)
RUG100 sp900315555 (GB_GCA_900315555.1)
UBA3738 sp002372505 (GB_GCA_002372505.1)
AnaerovoracaceaeUBA1191 (metabat.187)
UBA1191 sp900066305 (GB_GCA_900066305.1)
UBA1191 sp002310095 (GB_GCA_002310095.1)
Anaerovoracaceae (metabat.221)

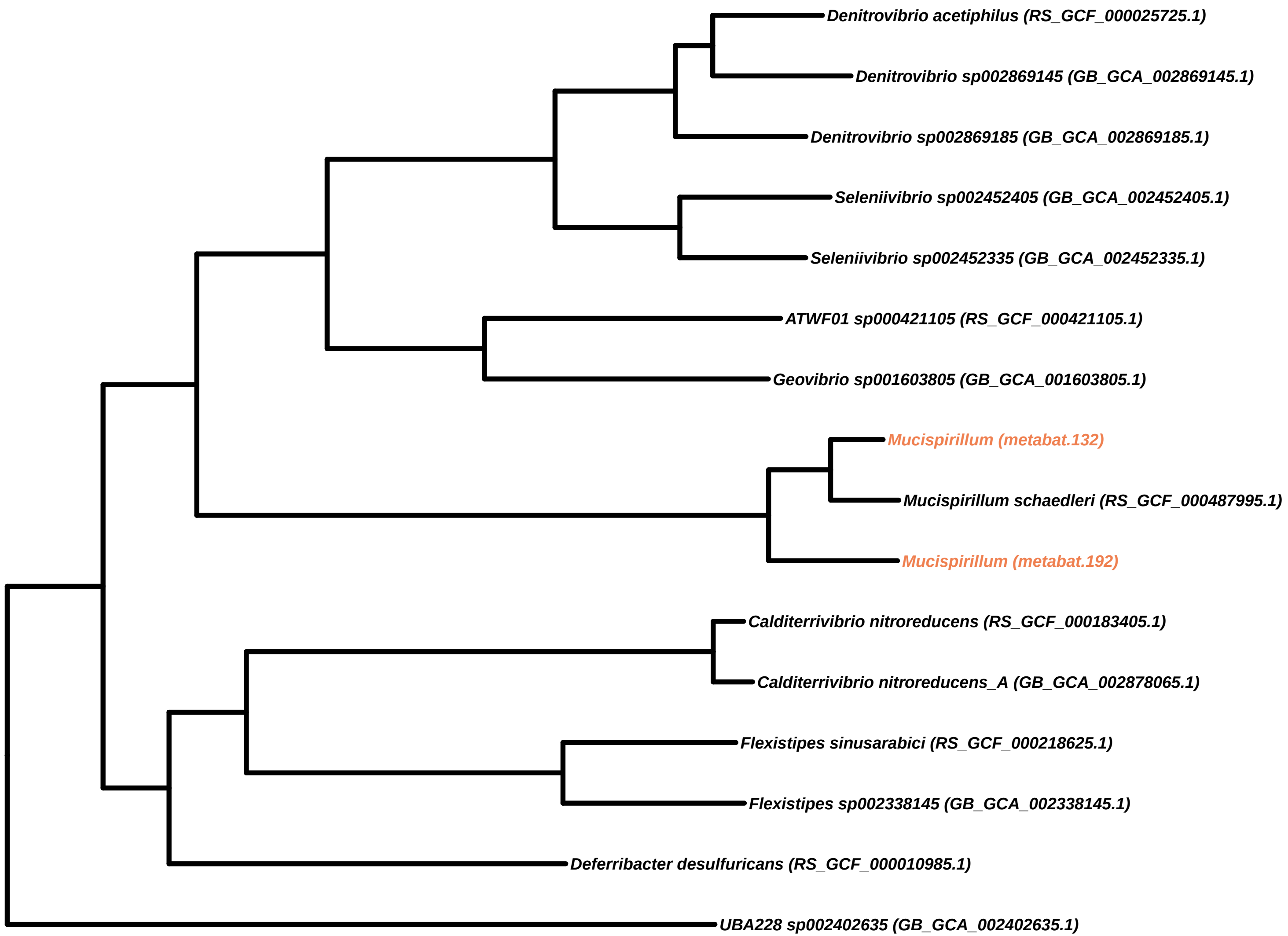
Denitrovibrio acetiphilus (RS_GCF_000025725.1)
Denitrovibrio sp002869145 (GB_GCA_002869145.1)
Denitrovibrio sp002869185 (GB_GCA_002869185.1)
Seleniivibrio sp002452405 (GB_GCA_002452405.1)
Seleniivibrio sp002452335 (GB_GCA_002452335.1)
ATWF01 sp000421105 (RS_GCF_000421105.1)
Geovibrio sp001603805 (GB_GCA_001603805.1)
Mucispirillum (metabat.132)
Mucispirillum schaedleri (RS_GCF_000487995.1)
Mucispirillum (metabat.192)
Calditerrivibrio nitroreducens (RS_GCF_000183405.1)
Calditerrivibrio nitroreducens_A (GB_GCA_002878065.1)
Flexistipes sinusarabici (RS_GCF_000218625.1)
Flexistipes sp002338145 (GB_GCA_002338145.1)
Deferribacter desulfuricans (RS_GCF_000010985.1)
UBA228 sp002402635 (GB_GCA_002402635.1)

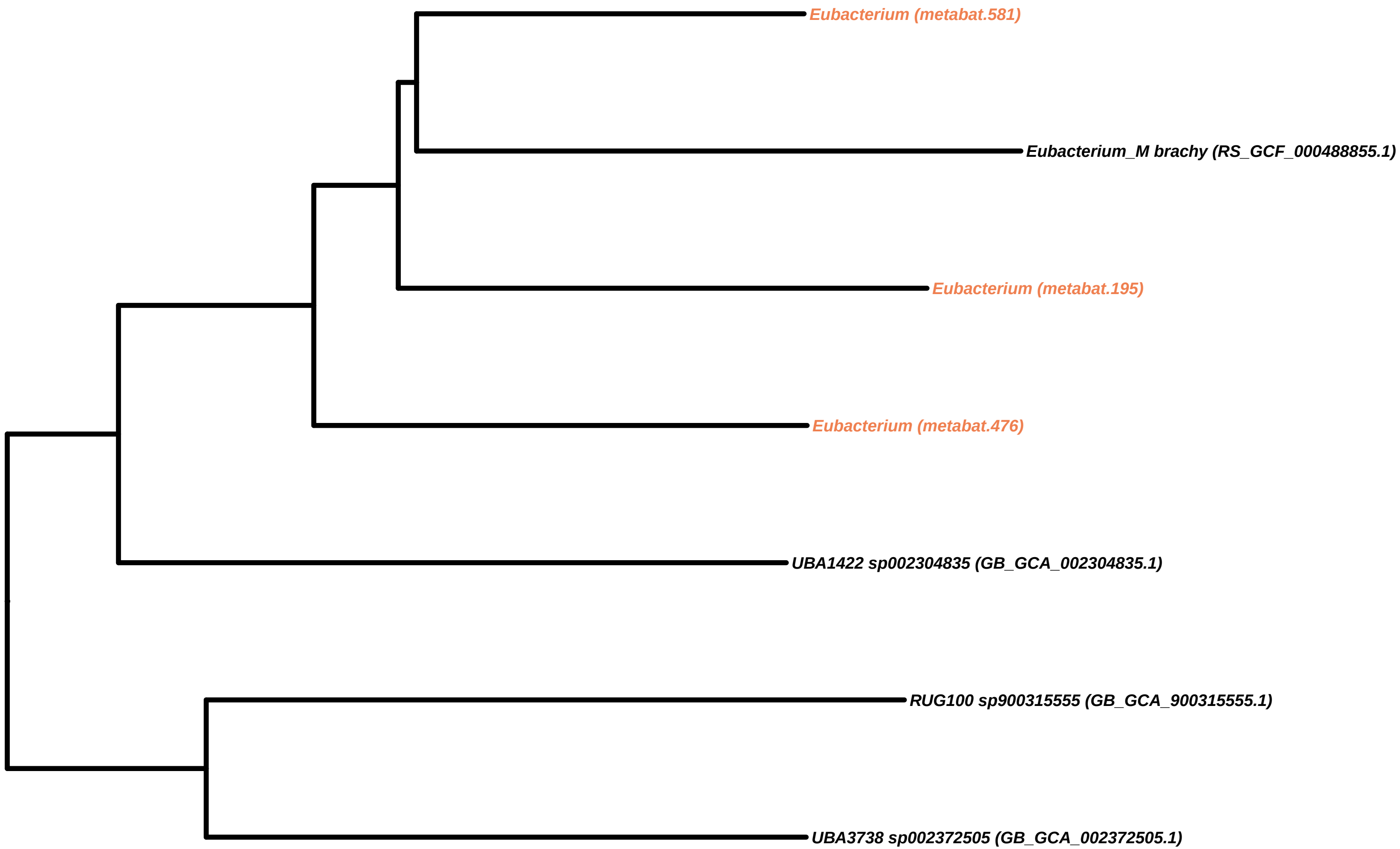

Eubacterium (metabat.581)
Eubacterium_M brachy (RS_GCF_000488855.1)
Eubacterium (metabat.195)
Eubacterium (metabat.476)
UBA1422 sp002304835 (GB_GCA_002304835.1)
RUG100 sp900315555 (GB_GCA_900315555.1)
UBA3738 sp002372505 (GB_GCA_002372505.1)

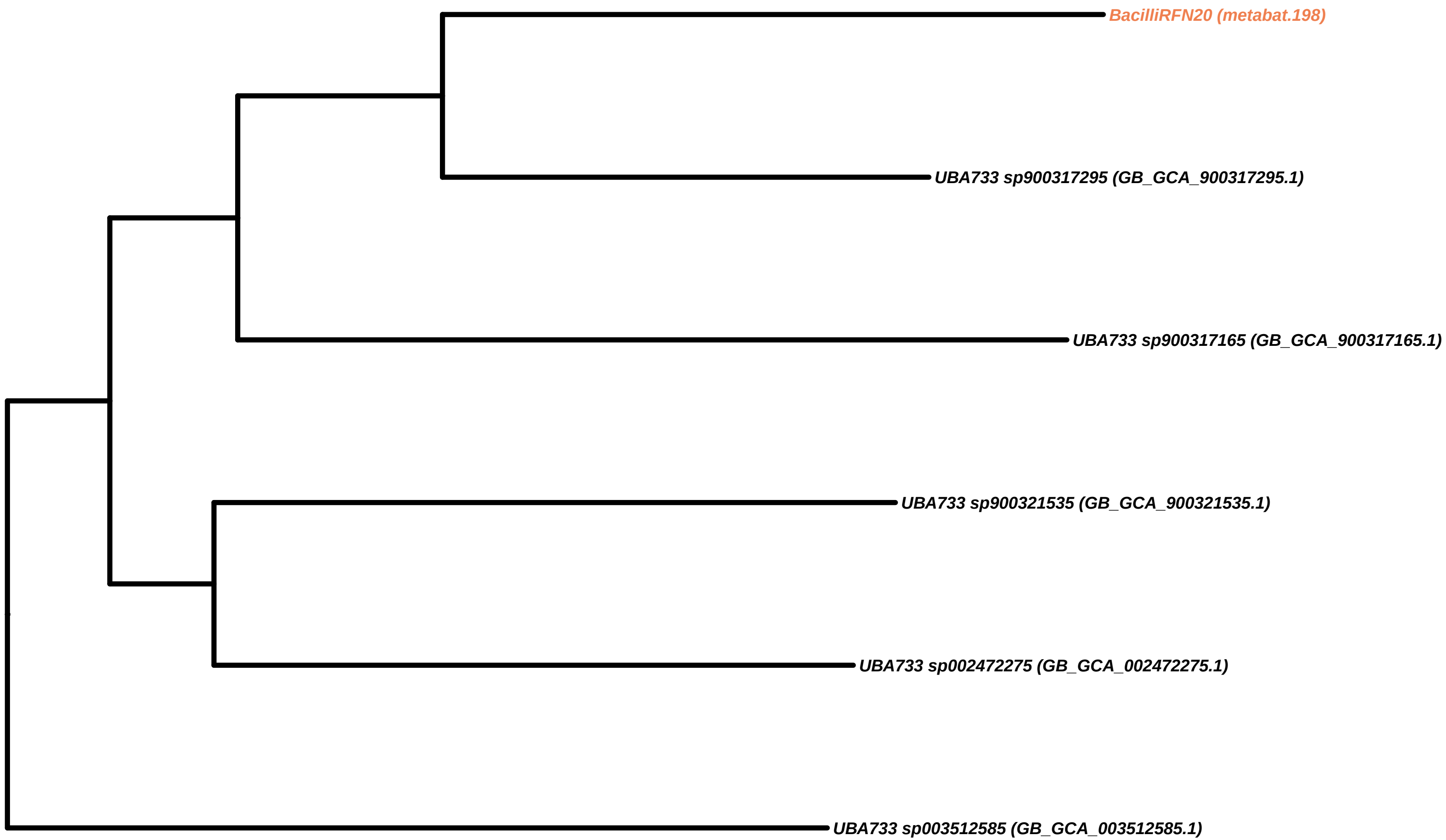
BacilliRFN20 (metabat.198)
UBA733 sp900317295 (GB_GCA_900317295.1)
UBA733 sp900317165 (GB_GCA_900317165.1)
UBA733 sp900321535 (GB_GCA_900321535.1)
UBA733 sp002472275 (GB_GCA_002472275.1)
UBA733 sp003512585 (GB_GCA_003512585.1)

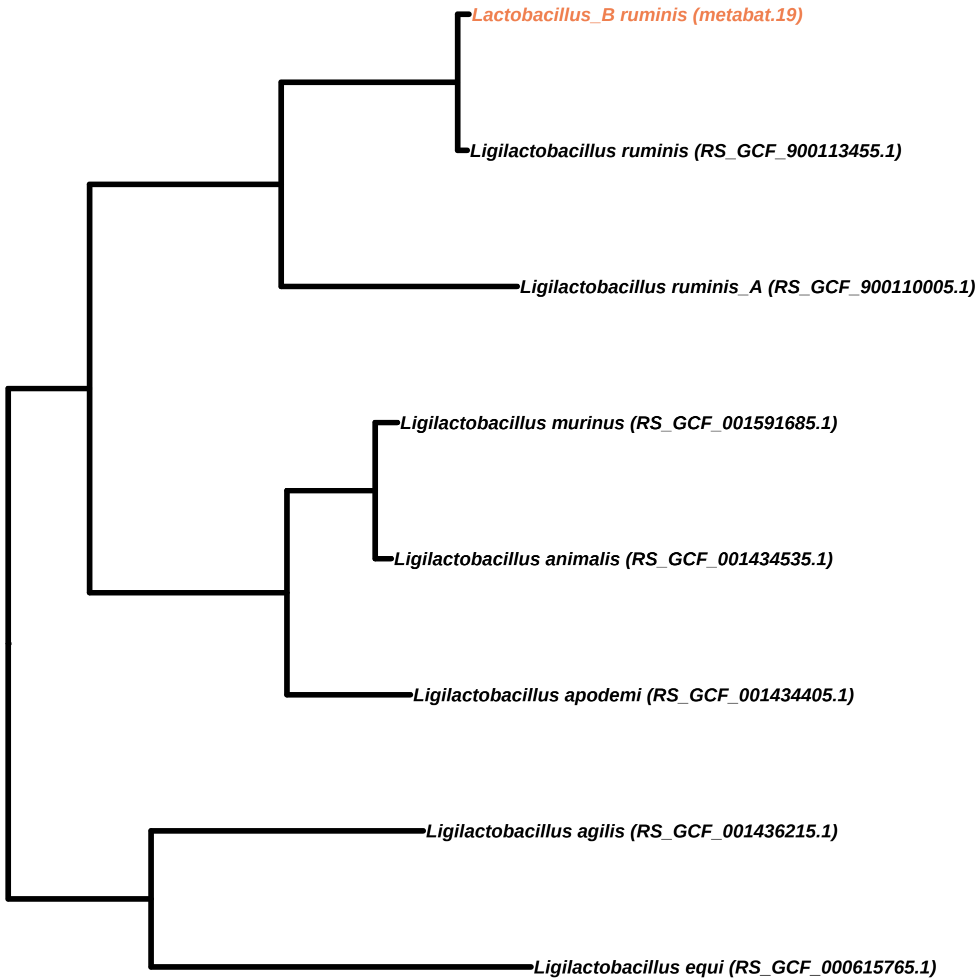
Lactobacillus_B ruminis (metabat.19)
Ligilactobacillus ruminis (RS_GCF_900113455.1)
Ligilactobacillus ruminis_A (RS_GCF_900110005.1)
Ligilactobacillus murinus (RS_GCF_001591685.1)
Ligilactobacillus animalis (RS_GCF_001434535.1)
Ligilactobacillus apodemi (RS_GCF_001434405.1)
Ligilactobacillus agilis (RS_GCF_001436215.1)
Ligilactobacillus equi (RS_GCF_000615765.1)

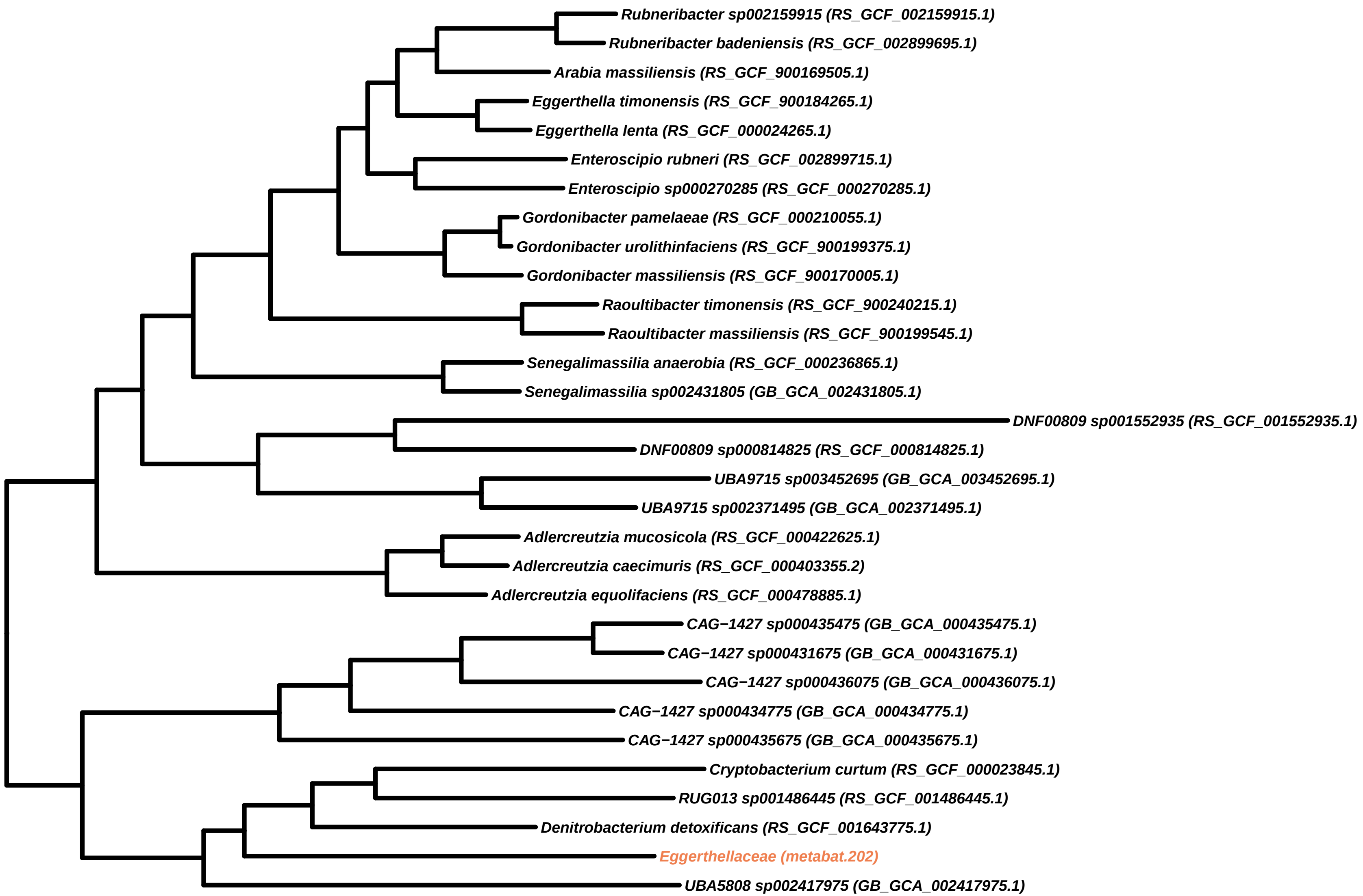

Rubneribacter sp002159915 (RS_GCF_002159915.1)
Rubneribacter badeniensis (RS_GCF_002899695.1)
Arabia massiliensis (RS_GCF_900169505.1)
Eggerthella timonensis (RS_GCF_900184265.1)
Eggerthella lenta (RS_GCF_000024265.1)
Enteroscipio rubneri (RS_GCF_002899715.1)
Enteroscipio sp000270285 (RS_GCF_000270285.1)
Gordonibacter pamelaeae (RS_GCF_000210055.1)
Gordonibacter urolithinfaciens (RS_GCF_900199375.1)
Gordonibacter massiliensis (RS_GCF_900170005.1)
Raoultibacter timonensis (RS_GCF_900240215.1)
Raoultibacter massiliensis (RS_GCF_900199545.1)
Senegalimassilia anaerobia (RS_GCF_000236865.1)
Senegalimassilia sp002431805 (GB_GCA_002431805.1)
DNF00809 sp001552935 (RS_GCF_001552935.1)
DNF00809 sp000814825 (RS_GCF_000814825.1)
UBA9715 sp003452695 (GB_GCA_003452695.1)
UBA9715 sp002371495 (GB_GCA_002371495.1)
Adlercreutzia mucosicola (RS_GCF_000422625.1)
Adlercreutzia caecimuris (RS_GCF_000403355.2)
Adlercreutzia equolifaciens (RS_GCF_000478885.1)
CAG−1427 sp000435475 (GB_GCA_000435475.1)
CAG−1427 sp000431675 (GB_GCA_000431675.1)
CAG−1427 sp000436075 (GB_GCA_000436075.1)
CAG−1427 sp000434775 (GB_GCA_000434775.1)
CAG−1427 sp000435675 (GB_GCA_000435675.1)
Cryptobacterium curtum (RS_GCF_000023845.1)
RUG013 sp001486445 (RS_GCF_001486445.1)
Denitrobacterium detoxificans (RS_GCF_001643775.1)
Eggerthellaceae (metabat.202)
UBA5808 sp002417975 (GB_GCA_002417975.1)

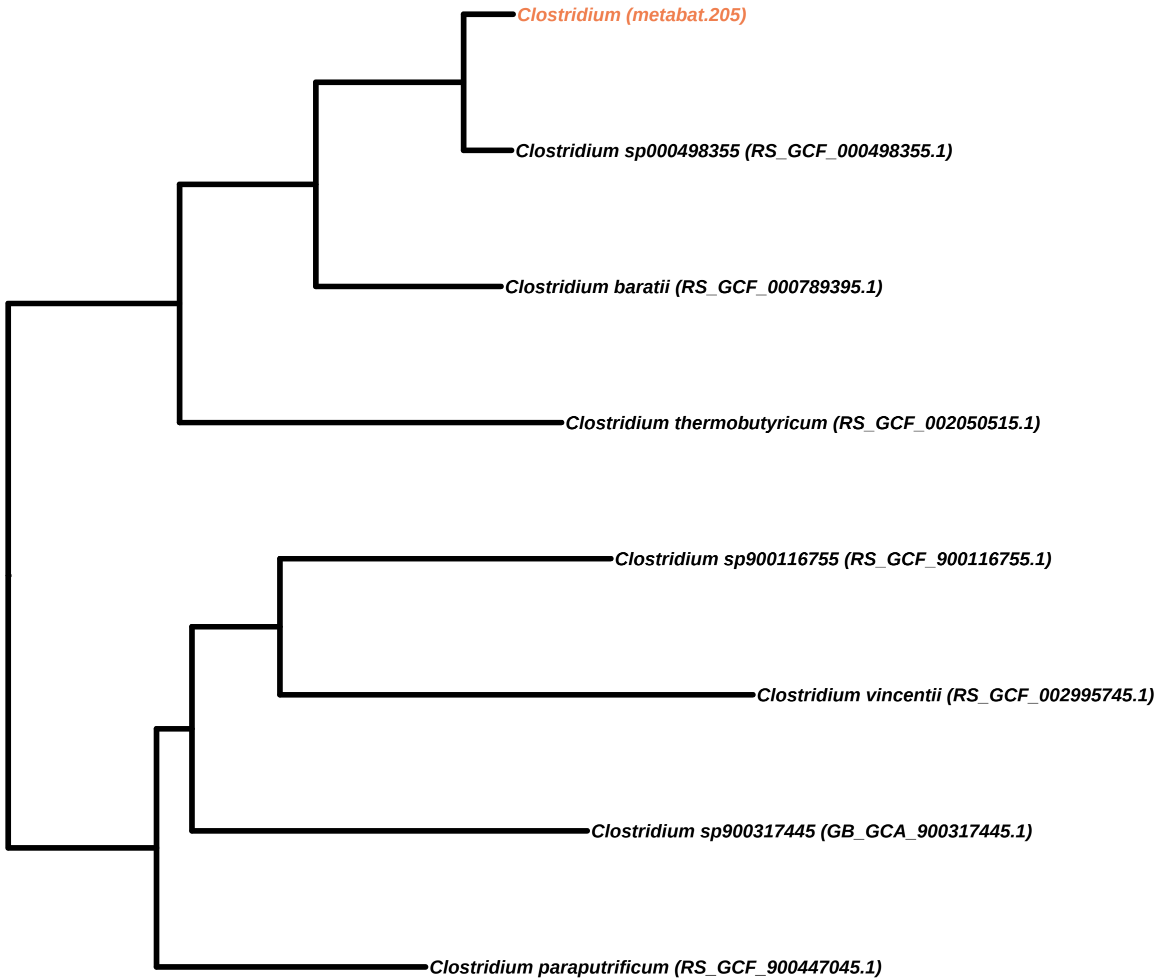

Clostridium (metabat.205)
Clostridium sp000498355 (RS_GCF_000498355.1)
Clostridium baratii (RS_GCF_000789395.1)
Clostridium thermobutyricum (RS_GCF_002050515.1)
Clostridium sp900116755 (RS_GCF_900116755.1)
Clostridium vincentii (RS_GCF_002995745.1)
Clostridium sp900317445 (GB_GCA_900317445.1)
Clostridium paraputrificum (RS_GCF_900447045.1)

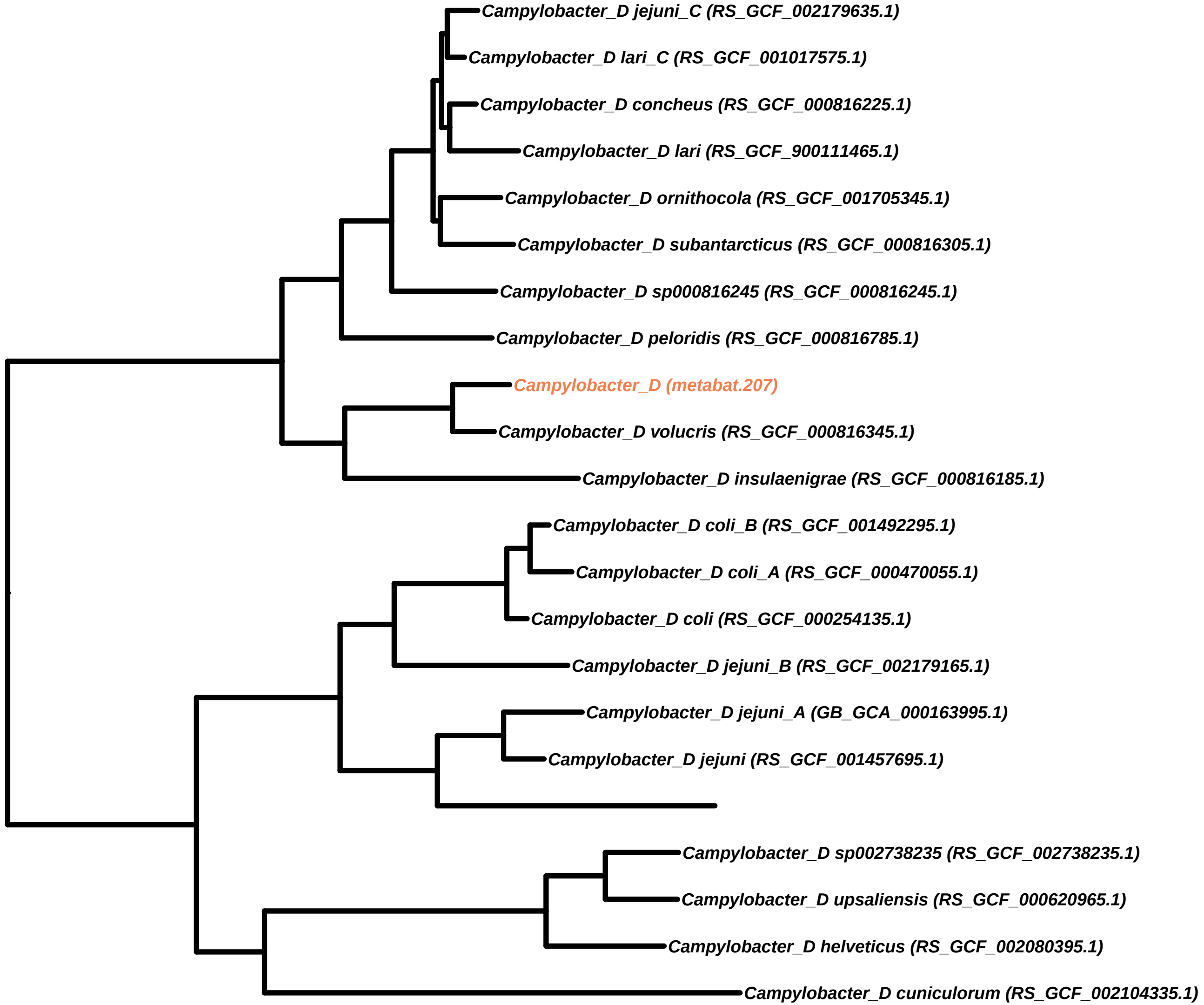
Campylobacter_D jejuni_C (RS_GCF_002179635.1)
Campylobacter_D lari_C (RS_GCF_001017575.1)
Campylobacter_D concheus (RS_GCF_000816225.1)
Campylobacter_D lari (RS_GCF_900111465.1)
Campylobacter_D ornithocola (RS_GCF_001705345.1)
Campylobacter_D subantarcticus (RS_GCF_000816305.1)
Campylobacter_D sp000816245 (RS_GCF_000816245.1)
Campylobacter_D peloridis (RS_GCF_000816785.1)
Campylobacter_D (metabat.207)
Campylobacter_D volucris (RS_GCF_000816345.1)
Campylobacter_D insulaenigrae (RS_GCF_000816185.1)
Campylobacter_D coli_B (RS_GCF_001492295.1)
Campylobacter_D coli_A (RS_GCF_000470055.1)
Campylobacter_D coli (RS_GCF_000254135.1)
Campylobacter_D jejuni_B (RS_GCF_002179165.1)
Campylobacter_D jejuni_A (GB_GCA_000163995.1)
Campylobacter_D jejuni (RS_GCF_001457695.1)
Campylobacter_D sp002738235 (RS_GCF_002738235.1)
Campylobacter_D upsaliensis (RS_GCF_000620965.1)
Campylobacter_D helveticus (RS_GCF_002080395.1)
Campylobacter_D cuniculorum (RS_GCF_002104335.1)

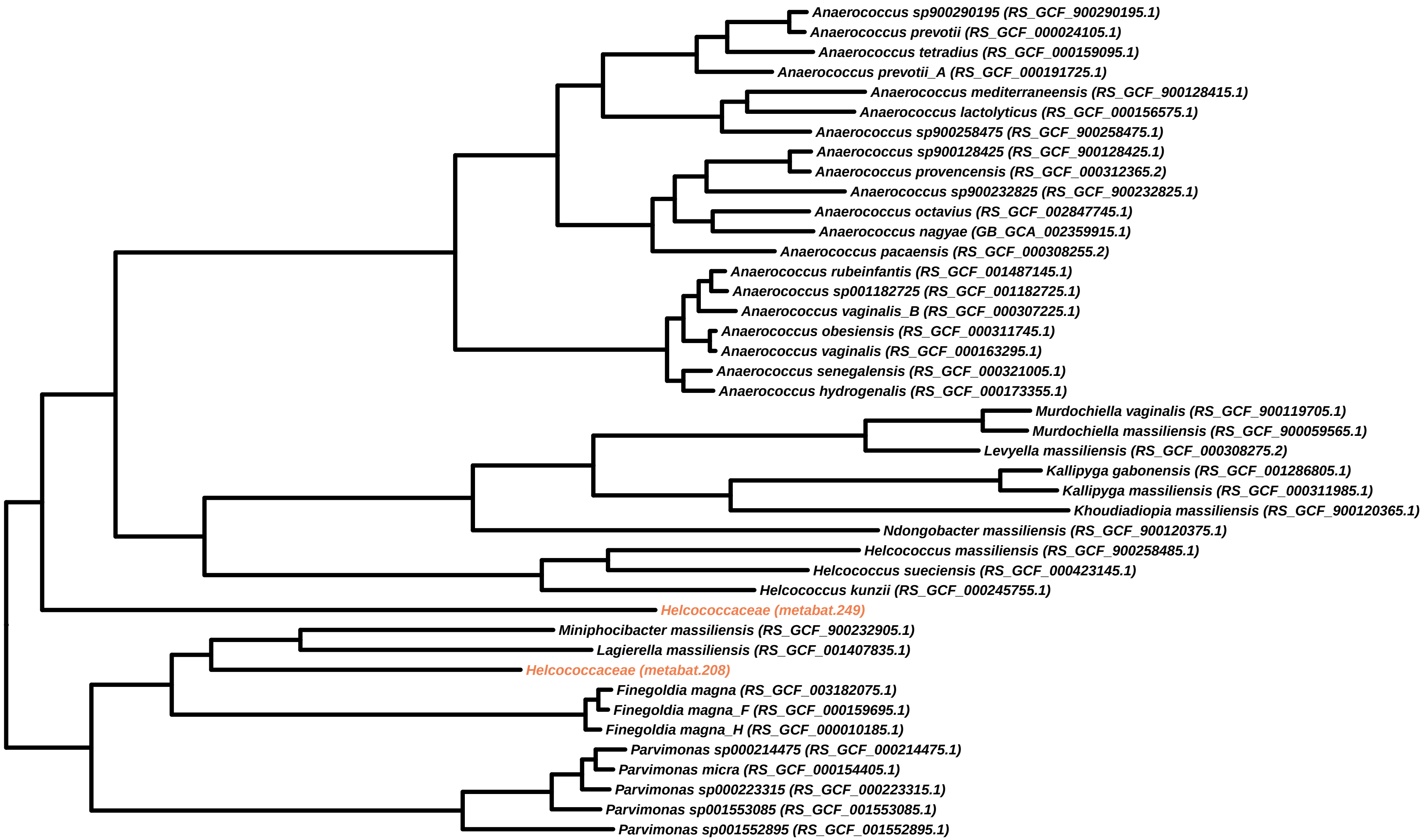

Anaerococcus sp900290195 (RS_GCF_900290195.1)
Anaerococcus prevotii (RS_GCF_000024105.1)
Anaerococcus tetradius (RS_GCF_000159095.1)
Anaerococcus prevotii_A (RS_GCF_000191725.1)
Anaerococcus mediterraneensis (RS_GCF_900128415.1)
Anaerococcus lactolyticus (RS_GCF_000156575.1)
Anaerococcus sp900258475 (RS_GCF_900258475.1)
Anaerococcus sp900128425 (RS_GCF_900128425.1)
Anaerococcus provencensis (RS_GCF_000312365.2)
Anaerococcus sp900232825 (RS_GCF_900232825.1)
Anaerococcus octavius (RS_GCF_002847745.1)
Anaerococcus nagyae (GB_GCA_002359915.1)
Anaerococcus pacaensis (RS_GCF_000308255.2)
Anaerococcus rubeinfantis (RS_GCF_001487145.1)
Anaerococcus sp001182725 (RS_GCF_001182725.1)
Anaerococcus vaginalis_B (RS_GCF_000307225.1)
Anaerococcus obesiensis (RS_GCF_000311745.1)
Anaerococcus vaginalis (RS_GCF_000163295.1)
Anaerococcus senegalensis (RS_GCF_000321005.1)
Anaerococcus hydrogenalis (RS_GCF_000173355.1)
Murdochiella vaginalis (RS_GCF_900119705.1)
Murdochiella massiliensis (RS_GCF_900059565.1)
Levyella massiliensis (RS_GCF_000308275.2)
Kallipyga gabonensis (RS_GCF_001286805.1)
Kallipyga massiliensis (RS_GCF_000311985.1)
Khoudiadiopia massiliensis (RS_GCF_900120365.1)
Ndongobacter massiliensis (RS_GCF_900120375.1)
Helcococcus massiliensis (RS_GCF_900258485.1)
Helcococcus sueciensis (RS_GCF_000423145.1)
Helcococcus kunzii (RS_GCF_000245755.1)
Helcococcaceae (metabat.249)
Miniphocibacter massiliensis (RS_GCF_900232905.1)
Lagierella massiliensis (RS_GCF_001407835.1)
Helcococcaceae (metabat.208)
Finegoldia magna (RS_GCF_003182075.1)
Finegoldia magna_F (RS_GCF_000159695.1)
Finegoldia magna_H (RS_GCF_000010185.1)
Parvimonas sp000214475 (RS_GCF_000214475.1)
Parvimonas micra (RS_GCF_000154405.1)
Parvimonas sp000223315 (RS_GCF_000223315.1)
Parvimonas sp001553085 (RS_GCF_001553085.1)
Parvimonas sp001552895 (RS_GCF_001552895.1)

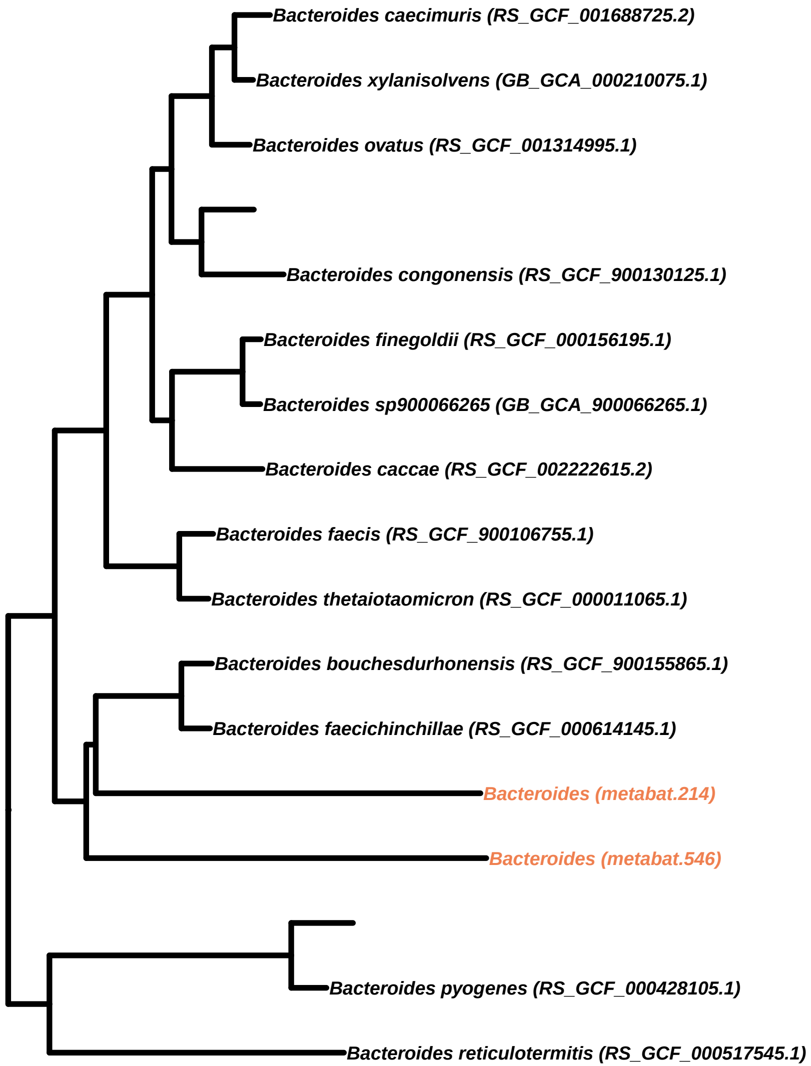
Bacteroides caecimuris (RS_GCF_001688725.2)
Bacteroides xylanisolvens (GB_GCA_000210075.1)
Bacteroides ovatus (RS_GCF_001314995.1)
Bacteroides congonensis (RS_GCF_900130125.1)
Bacteroides finegoldii (RS_GCF_000156195.1)
Bacteroides sp900066265 (GB_GCA_900066265.1)
Bacteroides caccae (RS_GCF_002222615.2)
Bacteroides faecis (RS_GCF_900106755.1)
Bacteroides thetaiotaomicron (RS_GCF_000011065.1)
Bacteroides bouchesdurhonensis (RS_GCF_900155865.1)
Bacteroides faecichinchillae (RS_GCF_000614145.1)
Bacteroides (metabat.214)
Bacteroides (metabat.546)
Bacteroides pyogenes (RS_GCF_000428105.1)
Bacteroides reticulotermitis (RS_GCF_000517545.1)

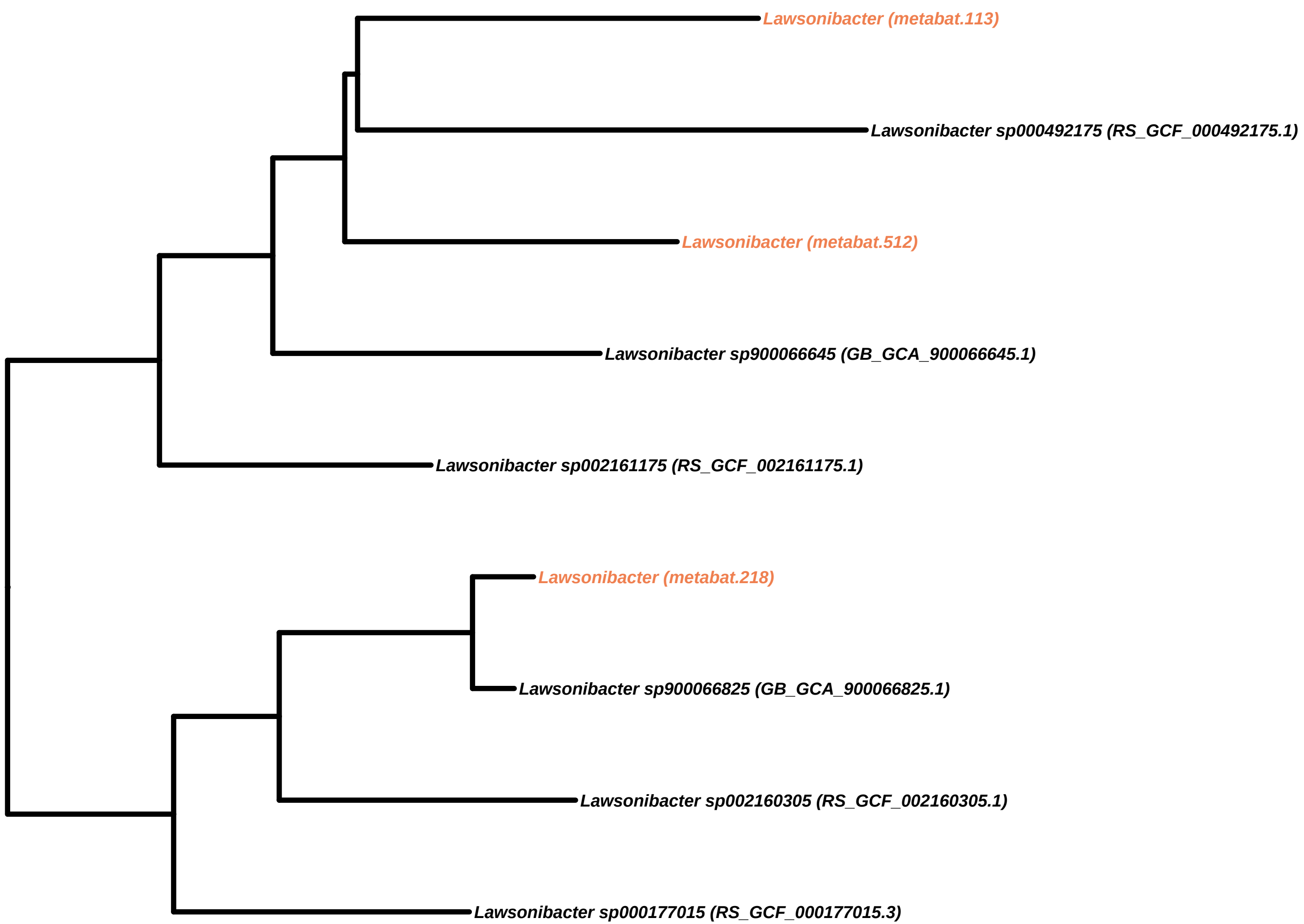
Lawsonibacter (metabat.113)
Lawsonibacter sp000492175 (RS_GCF_000492175.1)
Lawsonibacter (metabat.512)
Lawsonibacter sp900066645 (GB_GCA_900066645.1)
Lawsonibacter sp002161175 (RS_GCF_002161175.1)
Lawsonibacter (metabat.218)
Lawsonibacter sp900066825 (GB_GCA_900066825.1)
Lawsonibacter sp002160305 (RS_GCF_002160305.1)
Lawsonibacter sp000177015 (RS_GCF_000177015.3)

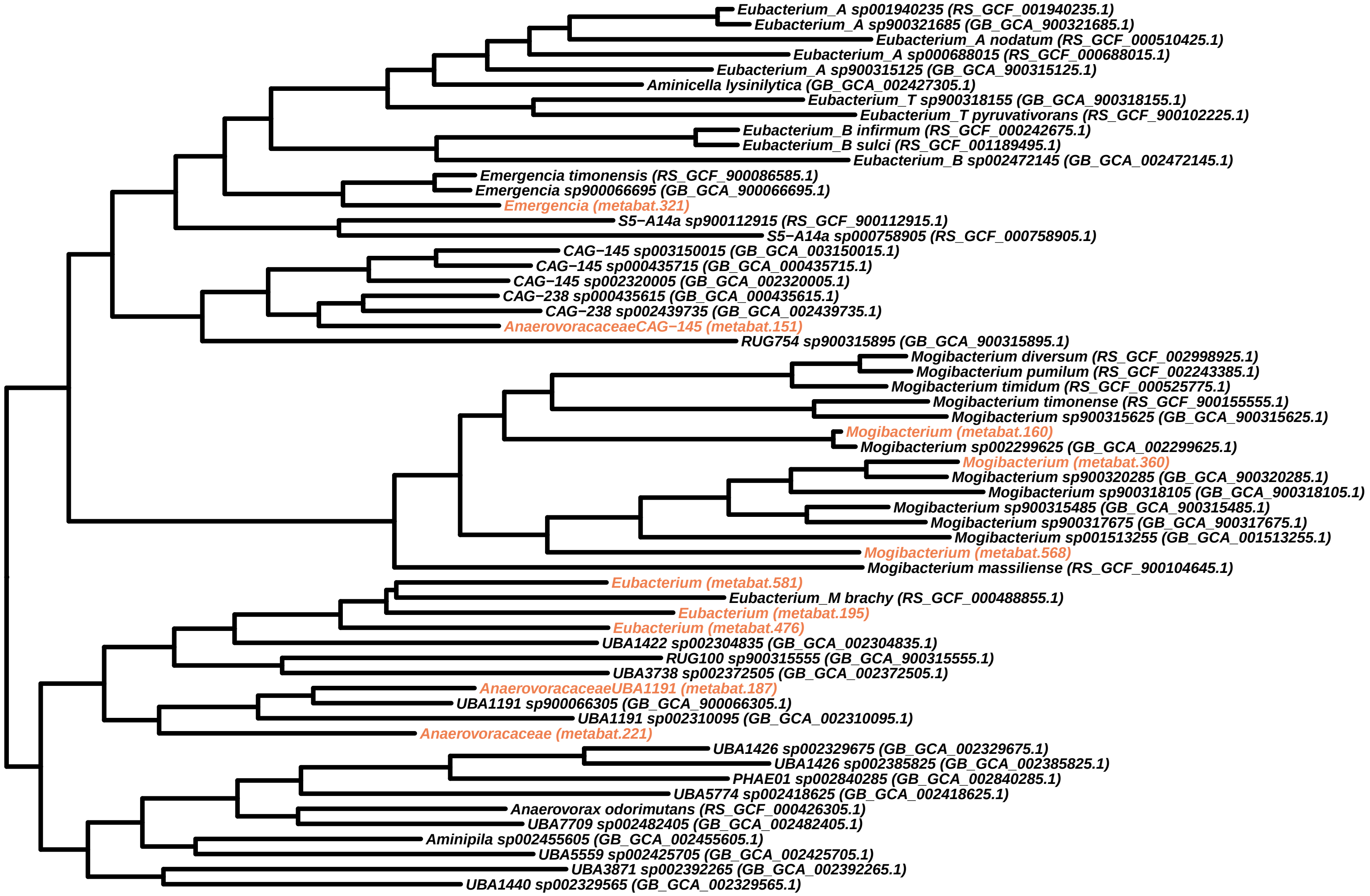

Eubacterium_A sp001940235 (RS_GCF_001940235.1)
Eubacterium_A sp900321685 (GB_GCA_900321685.1)
Eubacterium_A nodatum (RS_GCF_000510425.1)
Eubacterium_A sp000688015 (RS_GCF_000688015.1)
Eubacterium_A sp900315125 (GB_GCA_900315125.1)
Aminicella lysinilytica (GB_GCA_002427305.1)
Eubacterium_T sp900318155 (GB_GCA_900318155.1)
Eubacterium_T pyruvativorans (RS_GCF_900102225.1)
Eubacterium_B infirmum (RS_GCF_000242675.1)
Eubacterium_B sulci (RS_GCF_001189495.1)
Eubacterium_B sp002472145 (GB_GCA_002472145.1)
Emergencia timonensis (RS_GCF_900086585.1)
Emergencia sp900066695 (GB_GCA_900066695.1)
Emergencia (metabat.321)
S5−A14a sp900112915 (RS_GCF_900112915.1)
S5−A14a sp000758905 (RS_GCF_000758905.1)
CAG−145 sp003150015 (GB_GCA_003150015.1)
CAG−145 sp000435715 (GB_GCA_000435715.1)
CAG−145 sp002320005 (GB_GCA_002320005.1)
CAG−238 sp000435615 (GB_GCA_000435615.1)
CAG−238 sp002439735 (GB_GCA_002439735.1)
AnaerovoracaceaeCAG−145 (metabat.151)
RUG754 sp900315895 (GB_GCA_900315895.1)
Mogibacterium diversum (RS_GCF_002998925.1)
Mogibacterium pumilum (RS_GCF_002243385.1)
Mogibacterium timidum (RS_GCF_000525775.1)
Mogibacterium timonense (RS_GCF_900155555.1)
Mogibacterium sp900315625 (GB_GCA_900315625.1)
Mogibacterium (metabat.160)
Mogibacterium sp002299625 (GB_GCA_002299625.1)
Mogibacterium (metabat.360)
Mogibacterium sp900320285 (GB_GCA_900320285.1)
Mogibacterium sp900318105 (GB_GCA_900318105.1)
Mogibacterium sp900315485 (GB_GCA_900315485.1)
Mogibacterium sp900317675 (GB_GCA_900317675.1)
Mogibacterium sp001513255 (GB_GCA_001513255.1)
Mogibacterium (metabat.568)
Mogibacterium massiliense (RS_GCF_900104645.1)
Eubacterium (metabat.581)
Eubacterium_M brachy (RS_GCF_000488855.1)
Eubacterium (metabat.195)
Eubacterium (metabat.476)
UBA1422 sp002304835 (GB_GCA_002304835.1)
RUG100 sp900315555 (GB_GCA_900315555.1)
UBA3738 sp002372505 (GB_GCA_002372505.1)
AnaerovoracaceaeUBA1191 (metabat.187)
UBA1191 sp900066305 (GB_GCA_900066305.1)
UBA1191 sp002310095 (GB_GCA_002310095.1)
Anaerovoracaceae (metabat.221)
UBA1426 sp002329675 (GB_GCA_002329675.1)
UBA1426 sp002385825 (GB_GCA_002385825.1)
PHAE01 sp002840285 (GB_GCA_002840285.1)
UBA5774 sp002418625 (GB_GCA_002418625.1)
Anaerovorax odorimutans (RS_GCF_000426305.1)
UBA7709 sp002482405 (GB_GCA_002482405.1)
Aminipila sp002455605 (GB_GCA_002455605.1)
UBA5559 sp002425705 (GB_GCA_002425705.1)
UBA3871 sp002392265 (GB_GCA_002392265.1)
UBA1440 sp002329565 (GB_GCA_002329565.1)

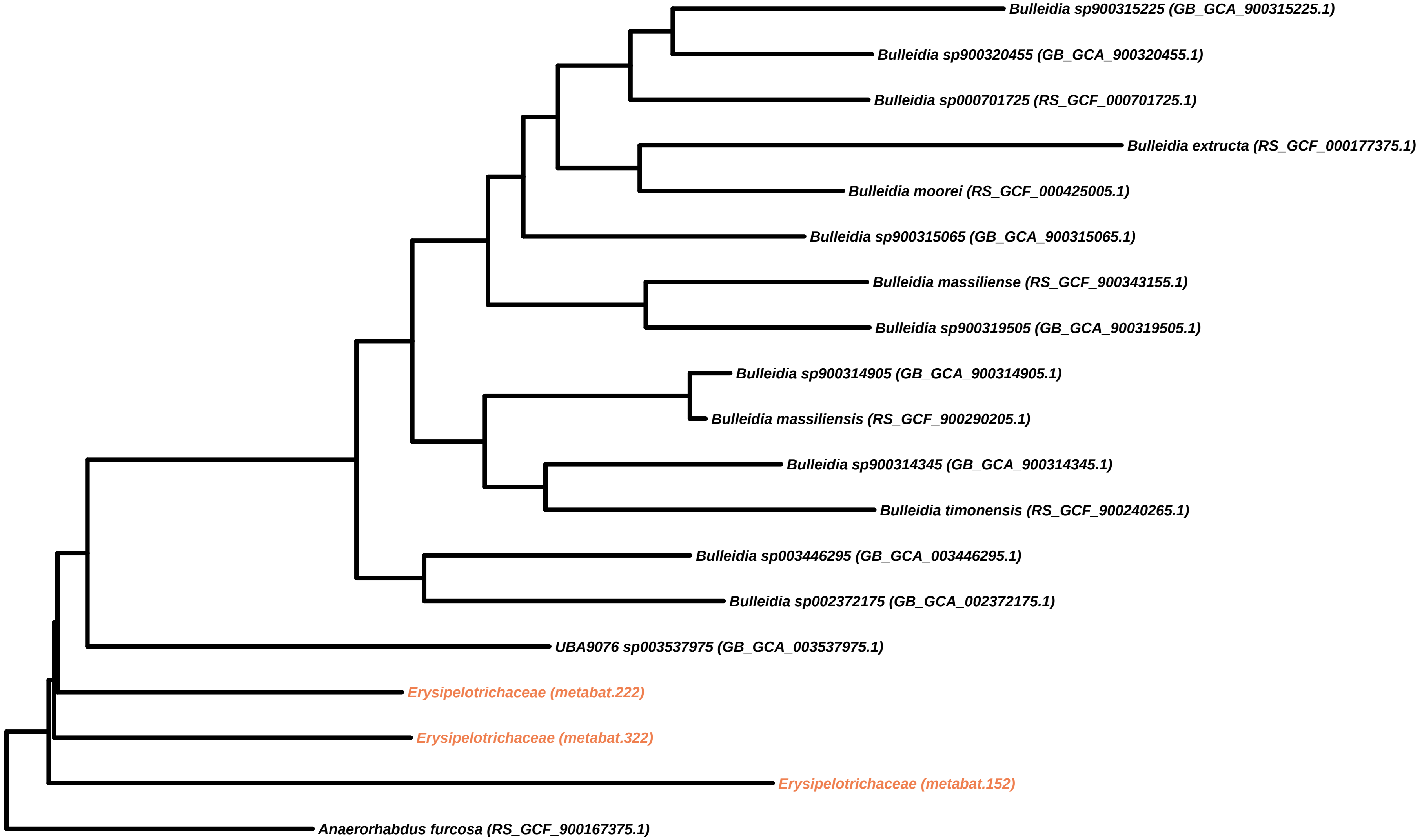
Bulleidia sp900315225 (GB_GCA_900315225.1)
Bulleidia sp900320455 (GB_GCA_900320455.1)
Bulleidia sp000701725 (RS_GCF_000701725.1)
Bulleidia extructa (RS_GCF_000177375.1)
Bulleidia moorei (RS_GCF_000425005.1)
Bulleidia sp900315065 (GB_GCA_900315065.1)
Bulleidia massiliense (RS_GCF_900343155.1)
Bulleidia sp900319505 (GB_GCA_900319505.1)
Bulleidia sp900314905 (GB_GCA_900314905.1)
Bulleidia massiliensis (RS_GCF_900290205.1)
Bulleidia sp900314345 (GB_GCA_900314345.1)
Bulleidia timonensis (RS_GCF_900240265.1)
Bulleidia sp003446295 (GB_GCA_003446295.1)
Bulleidia sp002372175 (GB_GCA_002372175.1)
UBA9076 sp003537975 (GB_GCA_003537975.1)
Erysipelotrichaceae (metabat.222)
Erysipelotrichaceae (metabat.322)
Erysipelotrichaceae (metabat.152)
Anaerorhabdus furcosa (RS_GCF_900167375.1)

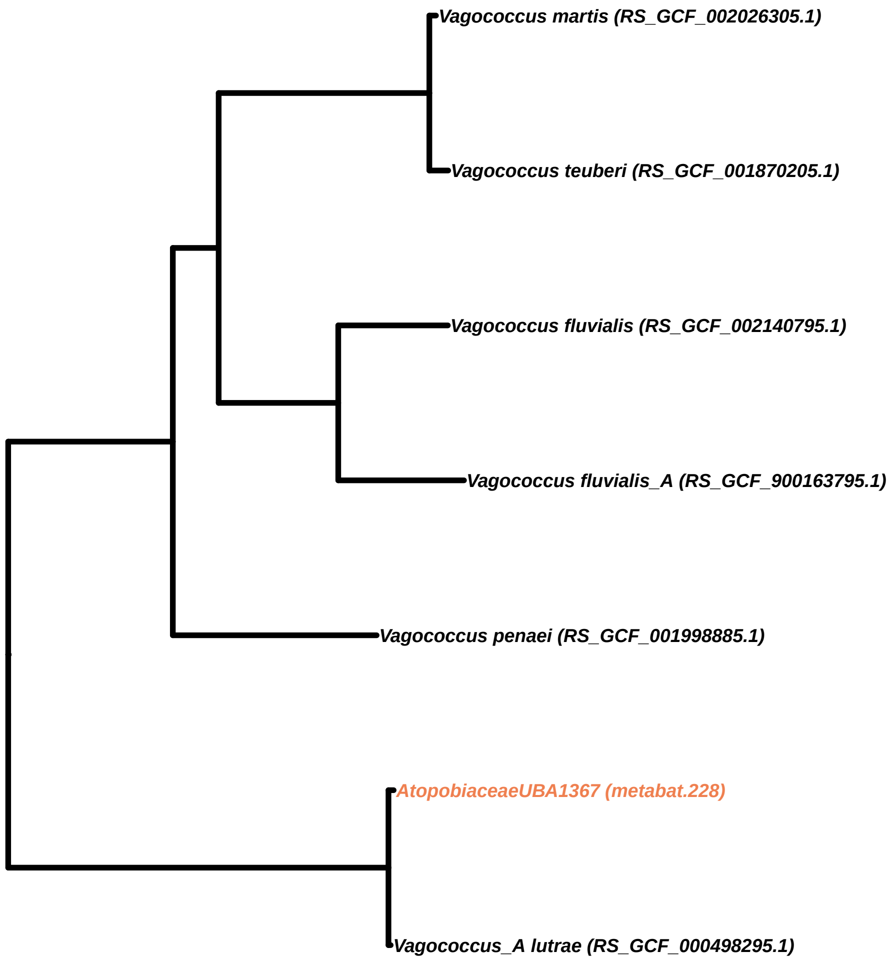
Vagococcus martis (RS_GCF_002026305.1)
Vagococcus teuberi (RS_GCF_001870205.1)
Vagococcus fluvialis (RS_GCF_002140795.1)
Vagococcus fluvialis_A (RS_GCF_900163795.1)
Vagococcus penaei (RS_GCF_001998885.1)
AtopobiaceaeUBA1367 (metabat.228)
Vagococcus_A lutrae (RS_GCF_000498295.1)

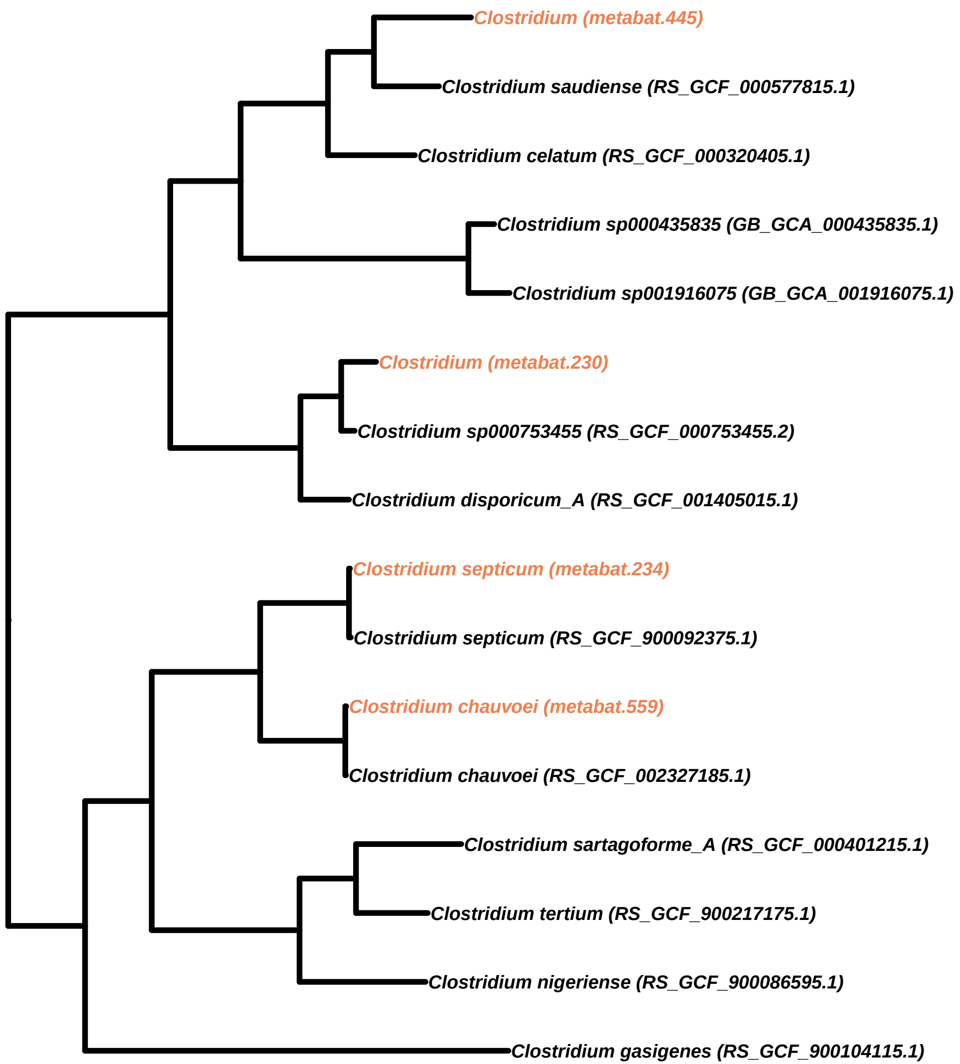
Clostridium (metabat.445)
Clostridium saudiense (RS_GCF_000577815.1)
Clostridium celatum (RS_GCF_000320405.1)
Clostridium sp000435835 (GB_GCA_000435835.1)
Clostridium sp001916075 (GB_GCA_001916075.1)
Clostridium (metabat.230)
Clostridium sp000753455 (RS_GCF_000753455.2)
Clostridium disporicum_A (RS_GCF_001405015.1)
Clostridium septicum (metabat.234)
Clostridium septicum (RS_GCF_900092375.1)
Clostridium chauvoei (metabat.559)
Clostridium chauvoei (RS_GCF_002327185.1)
Clostridium sartagoforme_A (RS_GCF_000401215.1)
Clostridium tertium (RS_GCF_900217175.1)
Clostridium nigeriense (RS_GCF_900086595.1)
Clostridium gasigenes (RS_GCF_900104115.1)

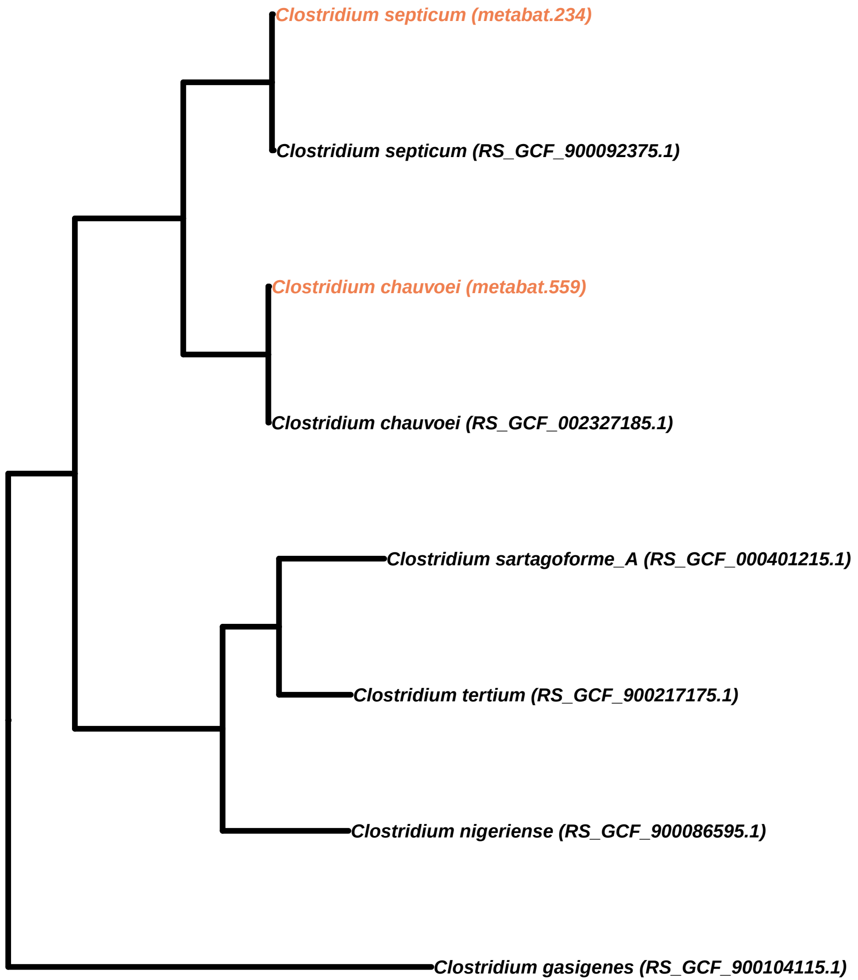
Clostridium septicum (metabat.234)
Clostridium septicum (RS_GCF_900092375.1)
Clostridium chauvoei (metabat.559)
Clostridium chauvoei (RS_GCF_002327185.1)
Clostridium sartagoforme_A (RS_GCF_000401215.1)
Clostridium tertium (RS_GCF_900217175.1)
Clostridium nigeriense (RS_GCF_900086595.1)
Clostridium gasigenes (RS_GCF_900104115.1)

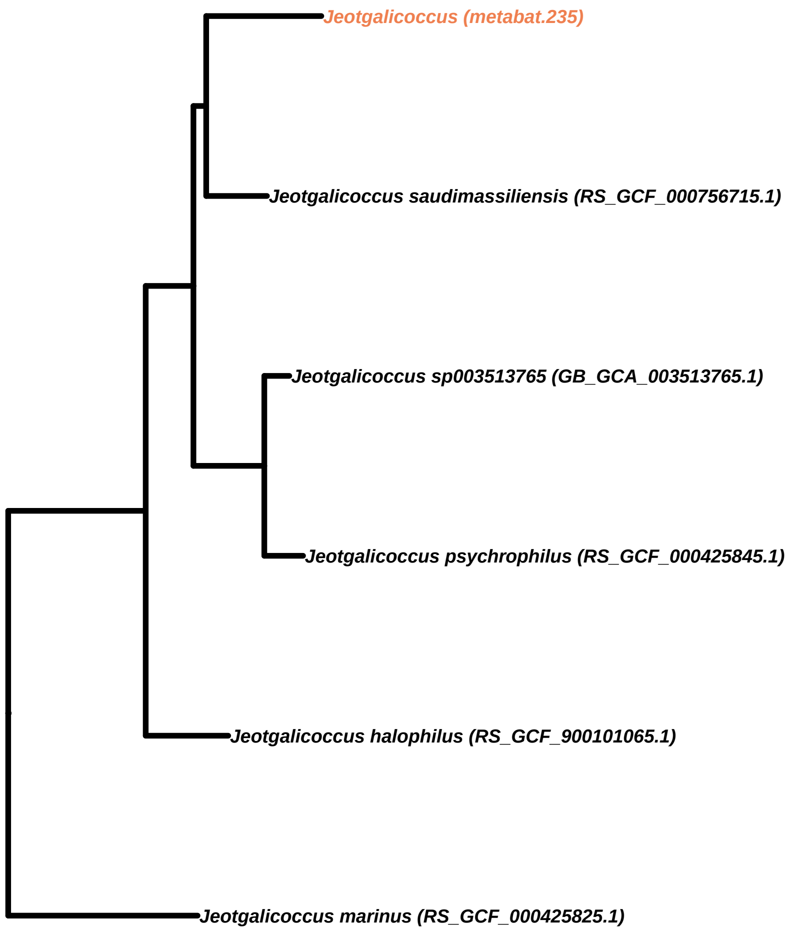
Jeotgalicoccus (metabat.235)
Jeotgalicoccus saudimassiliensis (RS_GCF_000756715.1)
Jeotgalicoccus sp003513765 (GB_GCA_003513765.1)
Jeotgalicoccus psychrophilus (RS_GCF_000425845.1)
Jeotgalicoccus halophilus (RS_GCF_900101065.1)
Jeotgalicoccus marinus (RS_GCF_000425825.1)

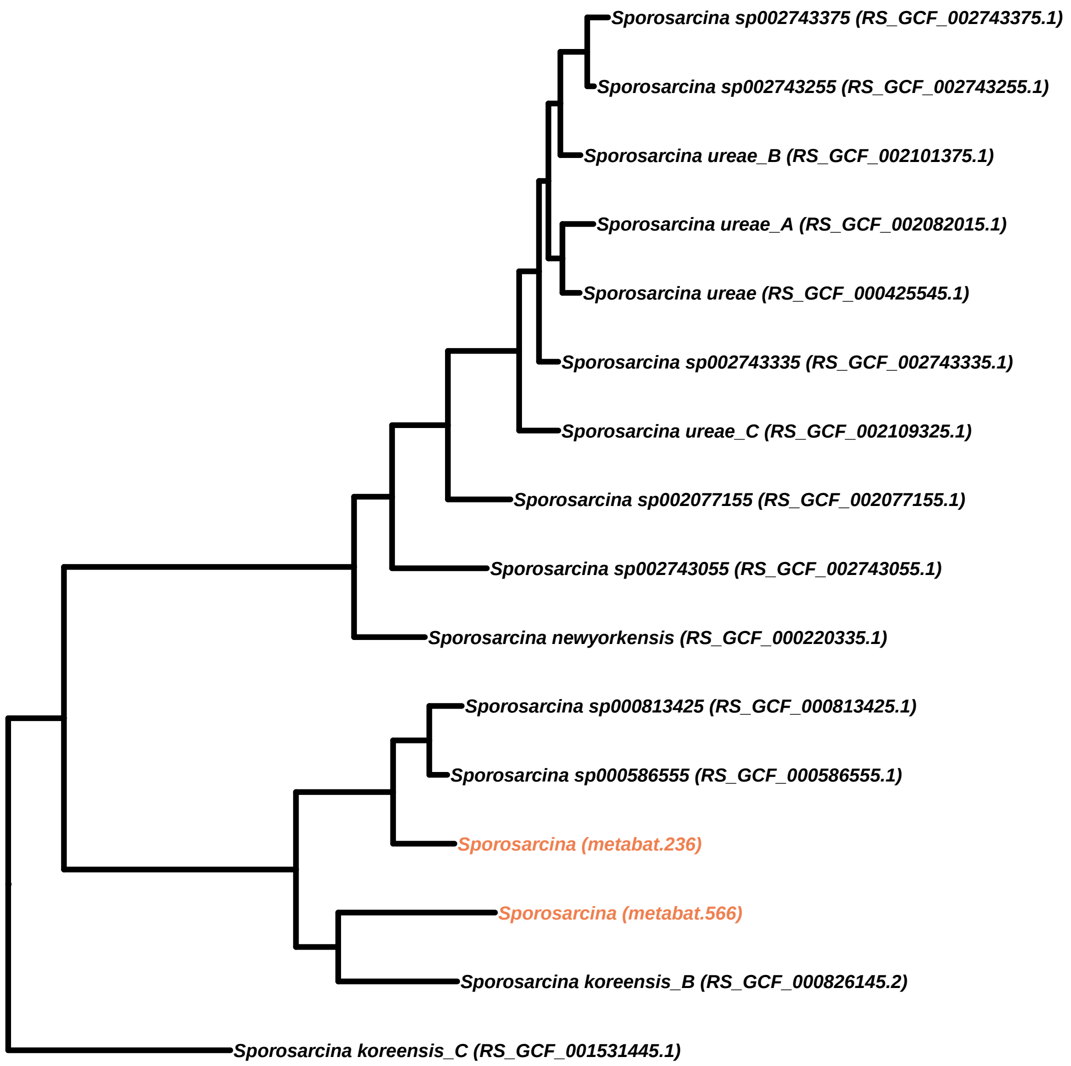

Sporosarcina sp002743375 (RS_GCF_002743375.1)
Sporosarcina sp002743255 (RS_GCF_002743255.1)
Sporosarcina ureae_B (RS_GCF_002101375.1)
Sporosarcina ureae_A (RS_GCF_002082015.1)
Sporosarcina ureae (RS_GCF_000425545.1)
Sporosarcina sp002743335 (RS_GCF_002743335.1)
Sporosarcina ureae_C (RS_GCF_002109325.1)
Sporosarcina sp002077155 (RS_GCF_002077155.1)
Sporosarcina sp002743055 (RS_GCF_002743055.1)
Sporosarcina newyorkensis (RS_GCF_000220335.1)
Sporosarcina sp000813425 (RS_GCF_000813425.1)
Sporosarcina sp000586555 (RS_GCF_000586555.1)
Sporosarcina (metabat.236)
Sporosarcina (metabat.566)
Sporosarcina koreensis_B (RS_GCF_000826145.2)
Sporosarcina koreensis_C (RS_GCF_001531445.1)

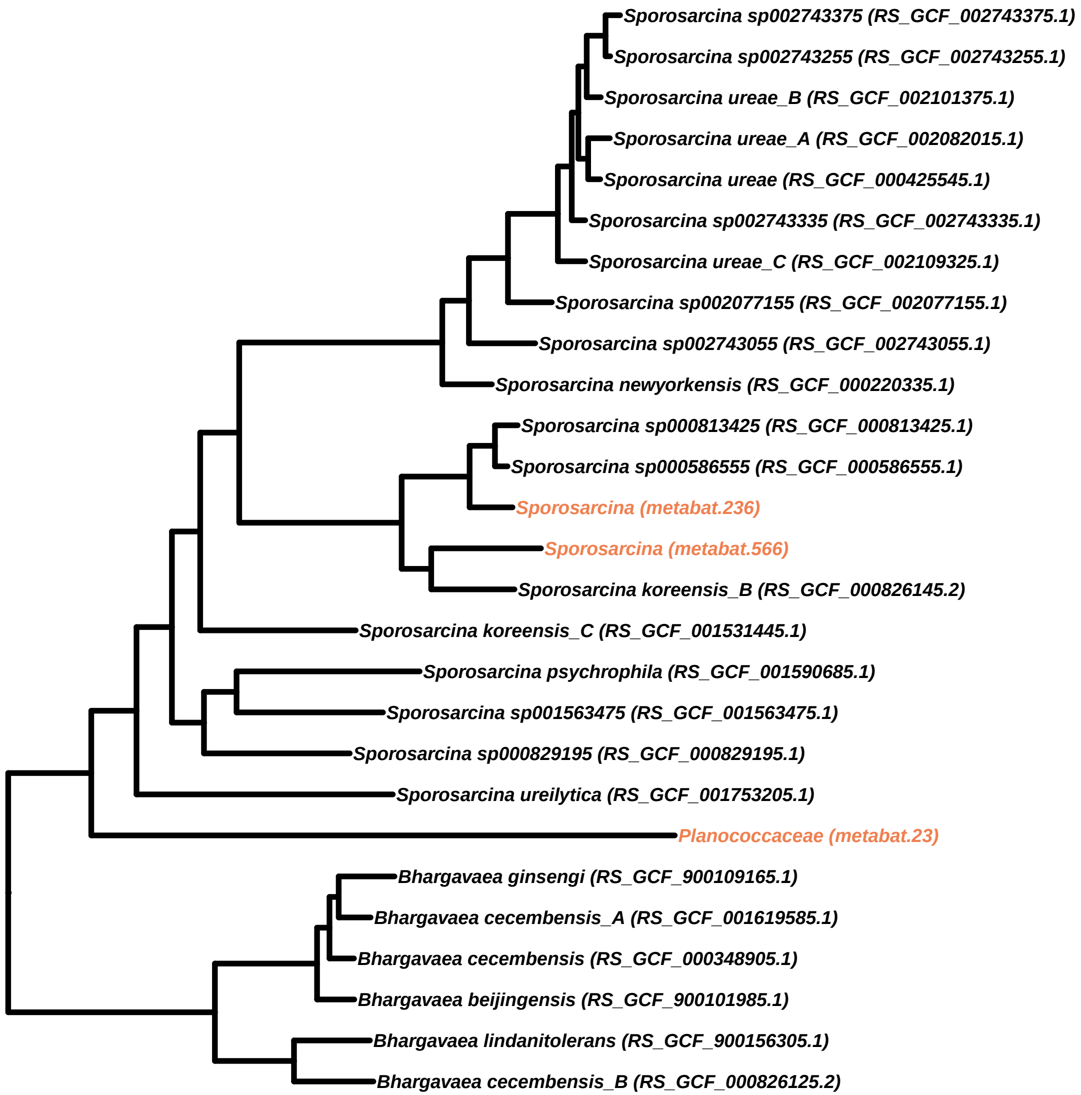
Sporosarcina sp002743375 (RS_GCF_002743375.1)
Sporosarcina sp002743255 (RS_GCF_002743255.1)
Sporosarcina ureae_B (RS_GCF_002101375.1)
Sporosarcina ureae_A (RS_GCF_002082015.1)
Sporosarcina ureae (RS_GCF_000425545.1)
Sporosarcina sp002743335 (RS_GCF_002743335.1)
Sporosarcina ureae_C (RS_GCF_002109325.1)
Sporosarcina sp002077155 (RS_GCF_002077155.1)
Sporosarcina sp002743055 (RS_GCF_002743055.1)
Sporosarcina newyorkensis (RS_GCF_000220335.1)
Sporosarcina sp000813425 (RS_GCF_000813425.1)
Sporosarcina sp000586555 (RS_GCF_000586555.1)
Sporosarcina (metabat.236)
Sporosarcina (metabat.566)
Sporosarcina koreensis_B (RS_GCF_000826145.2)
Sporosarcina koreensis_C (RS_GCF_001531445.1)
Sporosarcina psychrophila (RS_GCF_001590685.1)
Sporosarcina sp001563475 (RS_GCF_001563475.1)
Sporosarcina sp000829195 (RS_GCF_000829195.1)
Sporosarcina ureilytica (RS_GCF_001753205.1)
Planococcaceae (metabat.23)
Bhargavaea ginsengi (RS_GCF_900109165.1)
Bhargavaea cecembensis_A (RS_GCF_001619585.1)
Bhargavaea cecembensis (RS_GCF_000348905.1)
Bhargavaea beijingensis (RS_GCF_900101985.1)
Bhargavaea lindanitolerans (RS_GCF_900156305.1)
Bhargavaea cecembensis_B (RS_GCF_000826125.2)

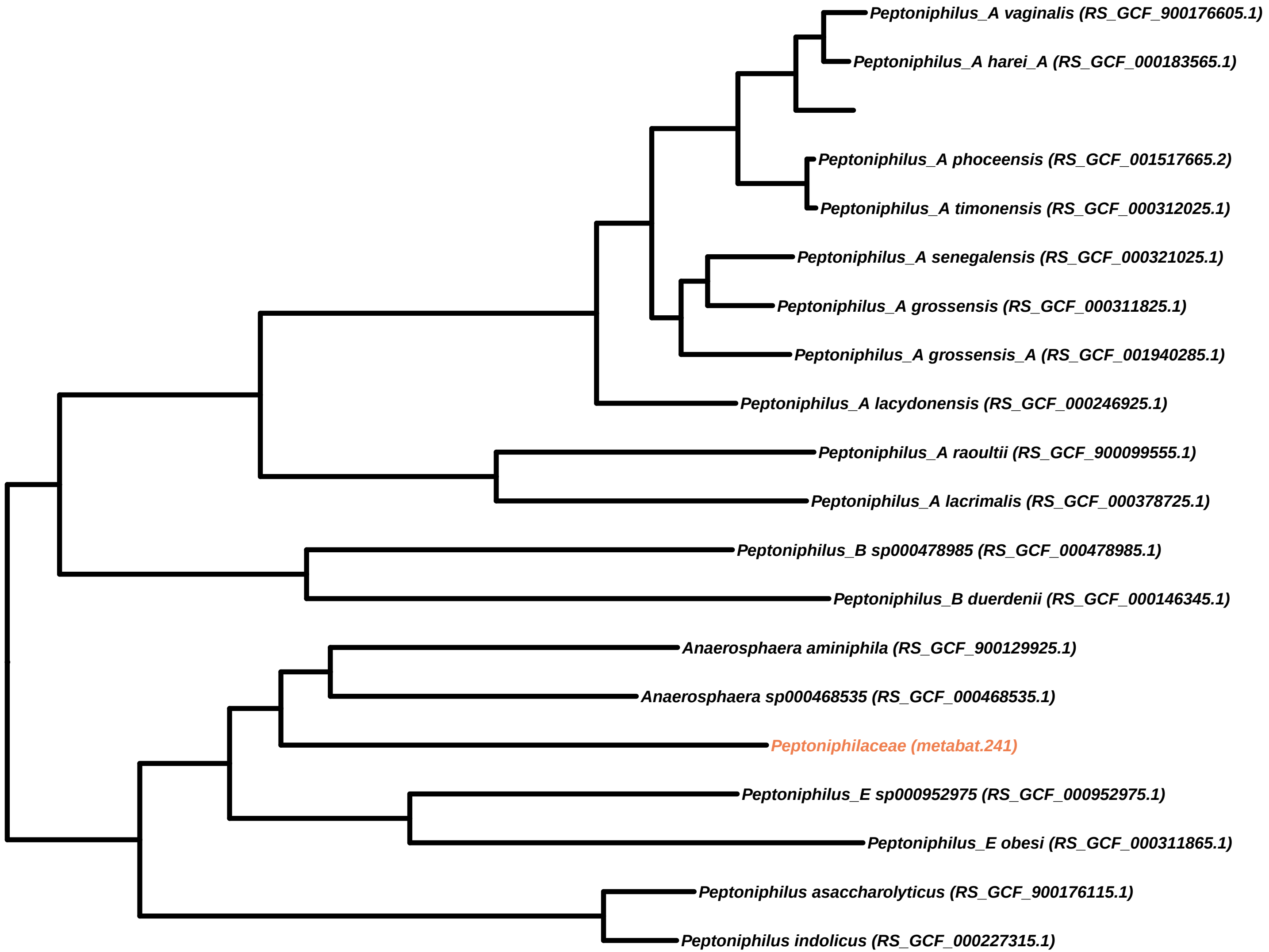
Peptoniphilus_A vaginalis (RS_GCF_900176605.1)
Peptoniphilus_A harei_A (RS_GCF_000183565.1)
Peptoniphilus_A phoceensis (RS_GCF_001517665.2)
Peptoniphilus_A timonensis (RS_GCF_000312025.1)
Peptoniphilus_A senegalensis (RS_GCF_000321025.1)
Peptoniphilus_A grossensis (RS_GCF_000311825.1)
Peptoniphilus_A grossensis_A (RS_GCF_001940285.1)
Peptoniphilus_A lacydonensis (RS_GCF_000246925.1)
Peptoniphilus_A raoultii (RS_GCF_900099555.1)
Peptoniphilus_A lacrimalis (RS_GCF_000378725.1)
Peptoniphilus_B sp000478985 (RS_GCF_000478985.1)
Peptoniphilus_B duerdenii (RS_GCF_000146345.1)
Anaerosphaera aminiphila (RS_GCF_900129925.1)
Anaerosphaera sp000468535 (RS_GCF_000468535.1)
Peptoniphilaceae (metabat.241)
Peptoniphilus_E sp000952975 (RS_GCF_000952975.1)
Peptoniphilus_E obesi (RS_GCF_000311865.1)
Peptoniphilus asaccharolyticus (RS_GCF_900176115.1)
Peptoniphilus indolicus (RS_GCF_000227315.1)

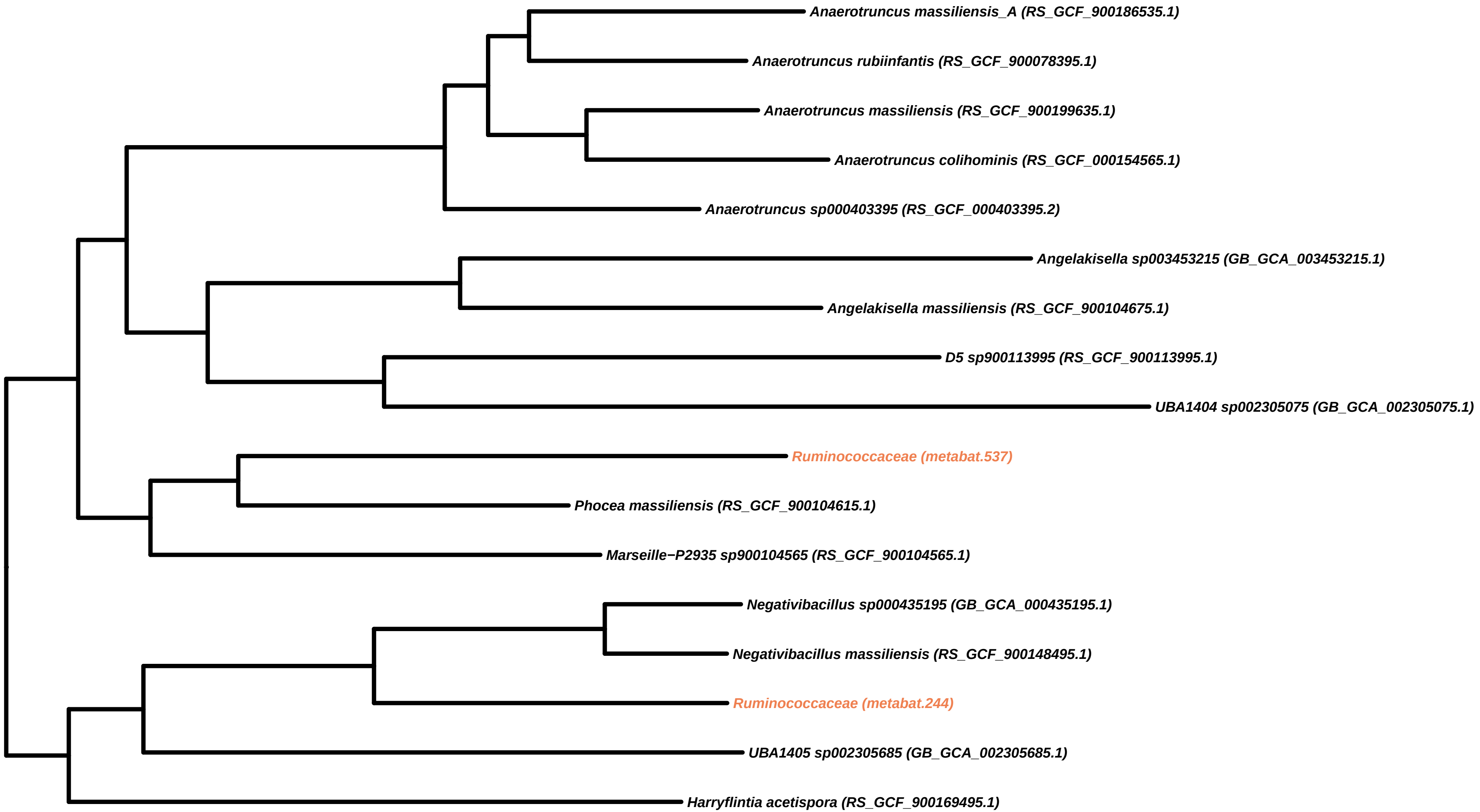

Anaerotruncus massiliensis_A (RS_GCF_900186535.1)
Anaerotruncus rubiinfantis (RS_GCF_900078395.1)
Anaerotruncus massiliensis (RS_GCF_900199635.1)
Anaerotruncus colihominis (RS_GCF_000154565.1)
Anaerotruncus sp000403395 (RS_GCF_000403395.2)
Angelakisella sp003453215 (GB_GCA_003453215.1)
Angelakisella massiliensis (RS_GCF_900104675.1)
D5 sp900113995 (RS_GCF_900113995.1)
UBA1404 sp002305075 (GB_GCA_002305075.1)
Ruminococcaceae (metabat.537)
Phocea massiliensis (RS_GCF_900104615.1)
Marseille−P2935 sp900104565 (RS_GCF_900104565.1)
Negativibacillus sp000435195 (GB_GCA_000435195.1)
Negativibacillus massiliensis (RS_GCF_900148495.1)
Ruminococcaceae (metabat.244)
UBA1405 sp002305685 (GB_GCA_002305685.1)
Harryflintia acetispora (RS_GCF_900169495.1)

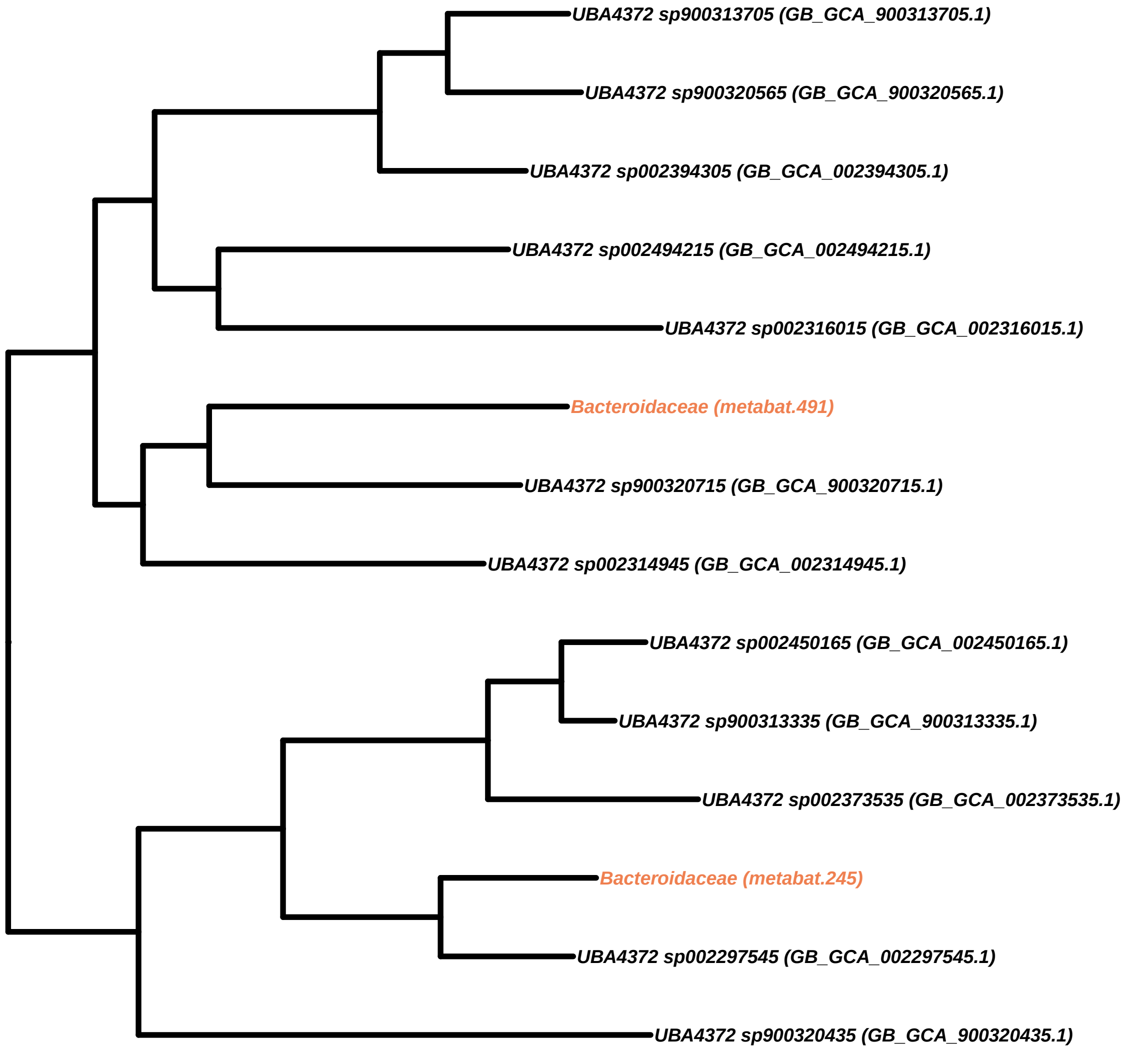
UBA4372 sp900313705 (GB_GCA_900313705.1)
UBA4372 sp900320565 (GB_GCA_900320565.1)
UBA4372 sp002394305 (GB_GCA_002394305.1)
UBA4372 sp002494215 (GB_GCA_002494215.1)
UBA4372 sp002316015 (GB_GCA_002316015.1)
Bacteroidaceae (metabat.491)
UBA4372 sp900320715 (GB_GCA_900320715.1)
UBA4372 sp002314945 (GB_GCA_002314945.1)
UBA4372 sp002450165 (GB_GCA_002450165.1)
UBA4372 sp900313335 (GB_GCA_900313335.1)
UBA4372 sp002373535 (GB_GCA_002373535.1)
Bacteroidaceae (metabat.245)
UBA4372 sp002297545 (GB_GCA_002297545.1)
UBA4372 sp900320435 (GB_GCA_900320435.1)

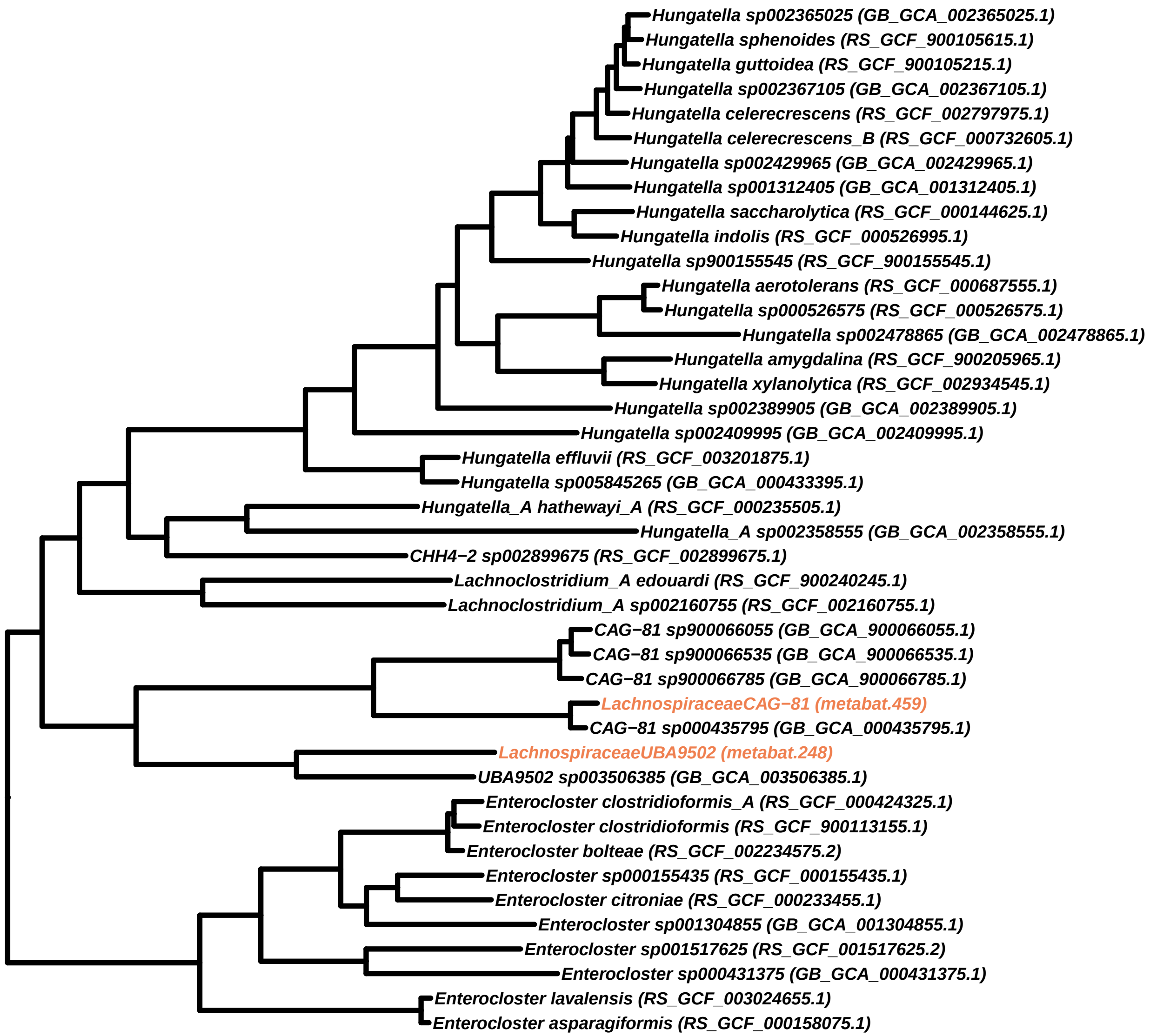
Hungatella sp002365025 (GB_GCA_002365025.1)
Hungatella sphenoides (RS_GCF_900105615.1)
Hungatella guttoidea (RS_GCF_900105215.1)
Hungatella sp002367105 (GB_GCA_002367105.1)
Hungatella celerecrescens (RS_GCF_002797975.1)
Hungatella celerecrescens_B (RS_GCF_000732605.1)
Hungatella sp002429965 (GB_GCA_002429965.1)
Hungatella sp001312405 (GB_GCA_001312405.1)
Hungatella saccharolytica (RS_GCF_000144625.1)
Hungatella indolis (RS_GCF_000526995.1)
Hungatella sp900155545 (RS_GCF_900155545.1)
Hungatella aerotolerans (RS_GCF_000687555.1)
Hungatella sp000526575 (RS_GCF_000526575.1)
Hungatella sp002478865 (GB_GCA_002478865.1)
Hungatella amygdalina (RS_GCF_900205965.1)
Hungatella xylanolytica (RS_GCF_002934545.1)
Hungatella sp002389905 (GB_GCA_002389905.1)
Hungatella sp002409995 (GB_GCA_002409995.1)
Hungatella effluvii (RS_GCF_003201875.1)
Hungatella sp005845265 (GB_GCA_000433395.1)
Hungatella_A hathewayi_A (RS_GCF_000235505.1)
Hungatella_A sp002358555 (GB_GCA_002358555.1)
CHH4−2 sp002899675 (RS_GCF_002899675.1)
Lachnoclostridium_A edouardi (RS_GCF_900240245.1)
Lachnoclostridium_A sp002160755 (RS_GCF_002160755.1)
CAG−81 sp900066055 (GB_GCA_900066055.1)
CAG−81 sp900066535 (GB_GCA_900066535.1)
CAG−81 sp900066785 (GB_GCA_900066785.1)
LachnospiraceaeCAG−81 (metabat.459)
CAG−81 sp000435795 (GB_GCA_000435795.1)
LachnospiraceaeUBA9502 (metabat.248)
UBA9502 sp003506385 (GB_GCA_003506385.1)
Enterocloster clostridioformis_A (RS_GCF_000424325.1)
Enterocloster clostridioformis (RS_GCF_900113155.1)
Enterocloster bolteae (RS_GCF_002234575.2)
Enterocloster sp000155435 (RS_GCF_000155435.1)
Enterocloster citroniae (RS_GCF_000233455.1)
Enterocloster sp001304855 (GB_GCA_001304855.1)
Enterocloster sp001517625 (RS_GCF_001517625.2)
Enterocloster sp000431375 (GB_GCA_000431375.1)
Enterocloster lavalensis (RS_GCF_003024655.1)
Enterocloster asparagiformis (RS_GCF_000158075.1)

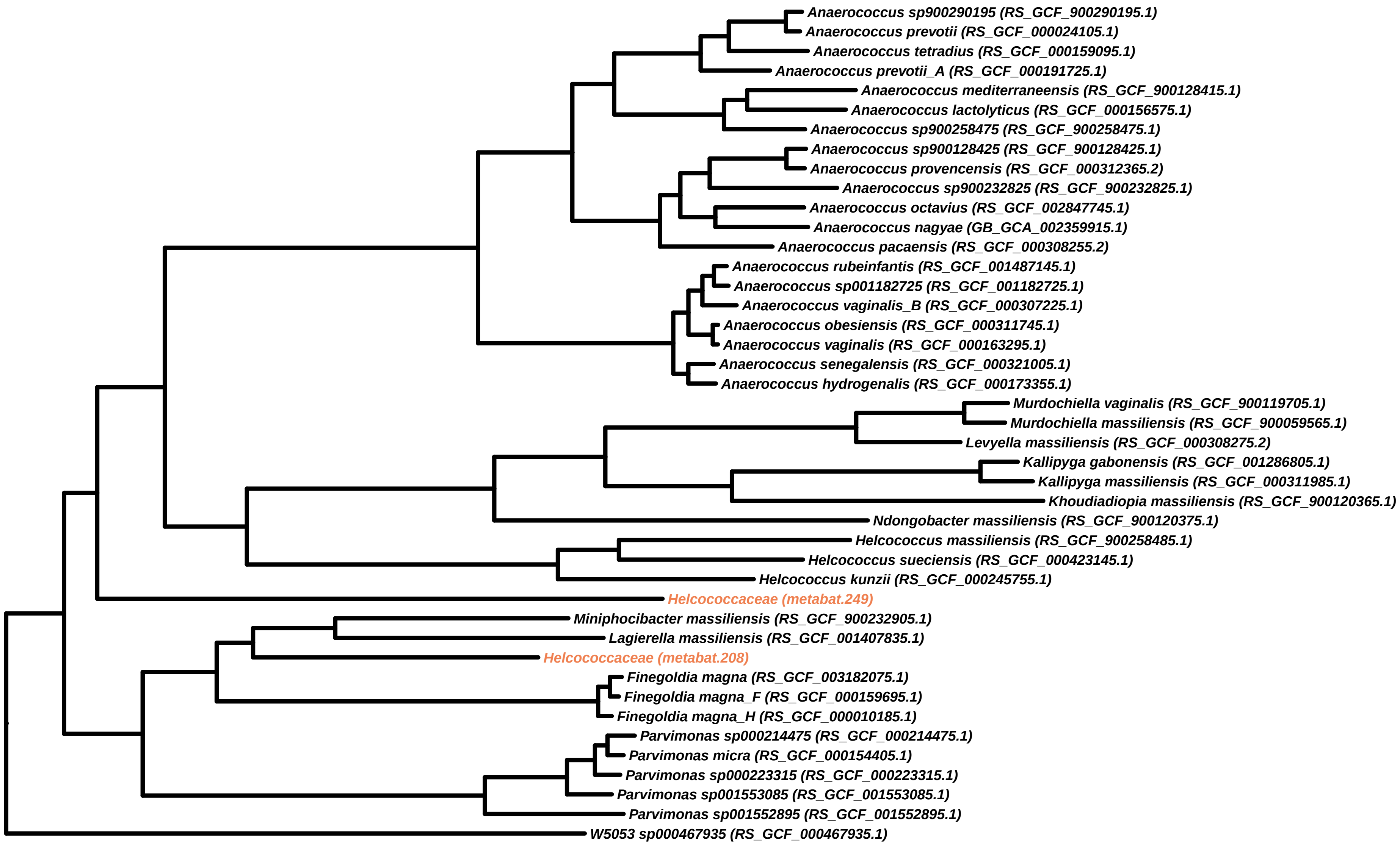
Anaerococcus sp900290195 (RS_GCF_900290195.1)
Anaerococcus prevotii (RS_GCF_000024105.1)
Anaerococcus tetradius (RS_GCF_000159095.1)
Anaerococcus prevotii_A (RS_GCF_000191725.1)
Anaerococcus mediterraneensis (RS_GCF_900128415.1)
Anaerococcus lactolyticus (RS_GCF_000156575.1)
Anaerococcus sp900258475 (RS_GCF_900258475.1)
Anaerococcus sp900128425 (RS_GCF_900128425.1)
Anaerococcus provencensis (RS_GCF_000312365.2)
Anaerococcus sp900232825 (RS_GCF_900232825.1)
Anaerococcus octavius (RS_GCF_002847745.1)
Anaerococcus nagyae (GB_GCA_002359915.1)
Anaerococcus pacaensis (RS_GCF_000308255.2)
Anaerococcus rubeinfantis (RS_GCF_001487145.1)
Anaerococcus sp001182725 (RS_GCF_001182725.1)
Anaerococcus vaginalis_B (RS_GCF_000307225.1)
Anaerococcus obesiensis (RS_GCF_000311745.1)
Anaerococcus vaginalis (RS_GCF_000163295.1)
Anaerococcus senegalensis (RS_GCF_000321005.1)
Anaerococcus hydrogenalis (RS_GCF_000173355.1)
Murdochiella vaginalis (RS_GCF_900119705.1)
Murdochiella massiliensis (RS_GCF_900059565.1)
Levyella massiliensis (RS_GCF_000308275.2)
Kallipyga gabonensis (RS_GCF_001286805.1)
Kallipyga massiliensis (RS_GCF_000311985.1)
Khoudiadiopia massiliensis (RS_GCF_900120365.1)
Ndongobacter massiliensis (RS_GCF_900120375.1)
Helcococcus massiliensis (RS_GCF_900258485.1)
Helcococcus sueciensis (RS_GCF_000423145.1)
Helcococcus kunzii (RS_GCF_000245755.1)
Helcococcaceae (metabat.249)
Miniphocibacter massiliensis (RS_GCF_900232905.1)
Lagierella massiliensis (RS_GCF_001407835.1)
Helcococcaceae (metabat.208)
Finegoldia magna (RS_GCF_003182075.1)
Finegoldia magna_F (RS_GCF_000159695.1)
Finegoldia magna_H (RS_GCF_000010185.1)
Parvimonas sp000214475 (RS_GCF_000214475.1)
Parvimonas micra (RS_GCF_000154405.1)
Parvimonas sp000223315 (RS_GCF_000223315.1)
Parvimonas sp001553085 (RS_GCF_001553085.1)
Parvimonas sp001552895 (RS_GCF_001552895.1)
W5053 sp000467935 (RS_GCF_000467935.1)

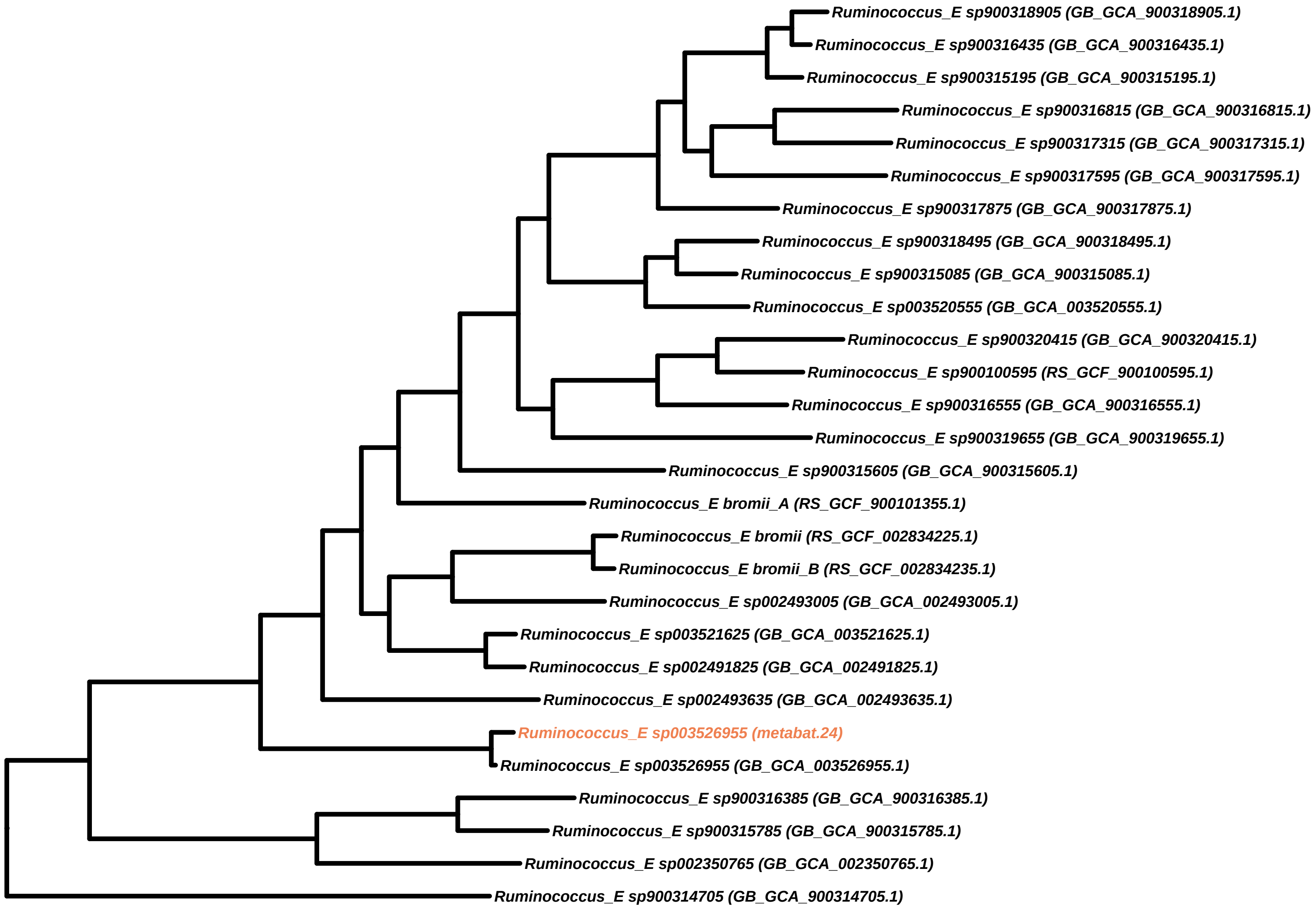

Ruminococcus_E sp900318905 (GB_GCA_900318905.1)
Ruminococcus_E sp900316435 (GB_GCA_900316435.1)
Ruminococcus_E sp900315195 (GB_GCA_900315195.1)
Ruminococcus_E sp900316815 (GB_GCA_900316815.1)
Ruminococcus_E sp900317315 (GB_GCA_900317315.1)
Ruminococcus_E sp900317595 (GB_GCA_900317595.1)
Ruminococcus_E sp900317875 (GB_GCA_900317875.1)
Ruminococcus_E sp900318495 (GB_GCA_900318495.1)
Ruminococcus_E sp900315085 (GB_GCA_900315085.1)
Ruminococcus_E sp003520555 (GB_GCA_003520555.1)
Ruminococcus_E sp900320415 (GB_GCA_900320415.1)
Ruminococcus_E sp900100595 (RS_GCF_900100595.1)
Ruminococcus_E sp900316555 (GB_GCA_900316555.1)
Ruminococcus_E sp900319655 (GB_GCA_900319655.1)
Ruminococcus_E sp900315605 (GB_GCA_900315605.1)
Ruminococcus_E bromii_A (RS_GCF_900101355.1)
Ruminococcus_E bromii (RS_GCF_002834225.1)
Ruminococcus_E bromii_B (RS_GCF_002834235.1)
Ruminococcus_E sp002493005 (GB_GCA_002493005.1)
Ruminococcus_E sp003521625 (GB_GCA_003521625.1)
Ruminococcus_E sp002491825 (GB_GCA_002491825.1)
Ruminococcus_E sp002493635 (GB_GCA_002493635.1)
Ruminococcus_E sp003526955 (metabat.24)
Ruminococcus_E sp003526955 (GB_GCA_003526955.1)
Ruminococcus_E sp900316385 (GB_GCA_900316385.1)
Ruminococcus_E sp900315785 (GB_GCA_900315785.1)
Ruminococcus_E sp002350765 (GB_GCA_002350765.1)
Ruminococcus_E sp900314705 (GB_GCA_900314705.1)

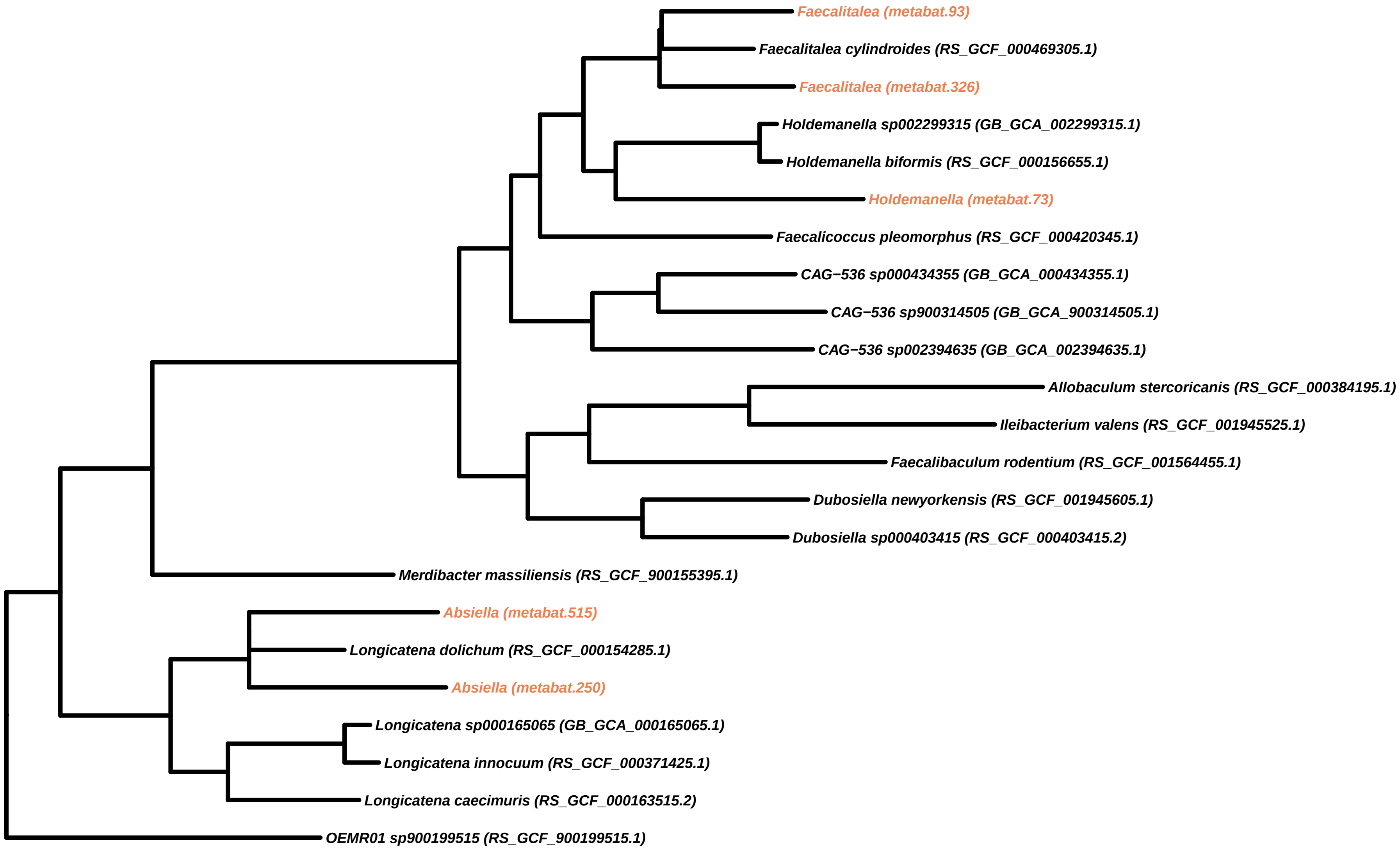

Faecalitalea (metabat.93)
Faecalitalea cylindroides (RS_GCF_000469305.1)
Faecalitalea (metabat.326)
Holdemanella sp002299315 (GB_GCA_002299315.1)
Holdemanella biformis (RS_GCF_000156655.1)
Holdemanella (metabat.73)
Faecalicoccus pleomorphus (RS_GCF_000420345.1)
CAG−536 sp000434355 (GB_GCA_000434355.1)
CAG−536 sp900314505 (GB_GCA_900314505.1)
CAG−536 sp002394635 (GB_GCA_002394635.1)
Allobaculum stercoricanis (RS_GCF_000384195.1)
Ileibacterium valens (RS_GCF_001945525.1)
Faecalibaculum rodentium (RS_GCF_001564455.1)
Dubosiella newyorkensis (RS_GCF_001945605.1)
Dubosiella sp000403415 (RS_GCF_000403415.2)
Merdibacter massiliensis (RS_GCF_900155395.1)
Absiella (metabat.515)
Longicatena dolichum (RS_GCF_000154285.1)
Absiella (metabat.250)
Longicatena sp000165065 (GB_GCA_000165065.1)
Longicatena innocuum (RS_GCF_000371425.1)
Longicatena caecimuris (RS_GCF_000163515.2)
OEMR01 sp900199515 (RS_GCF_900199515.1)

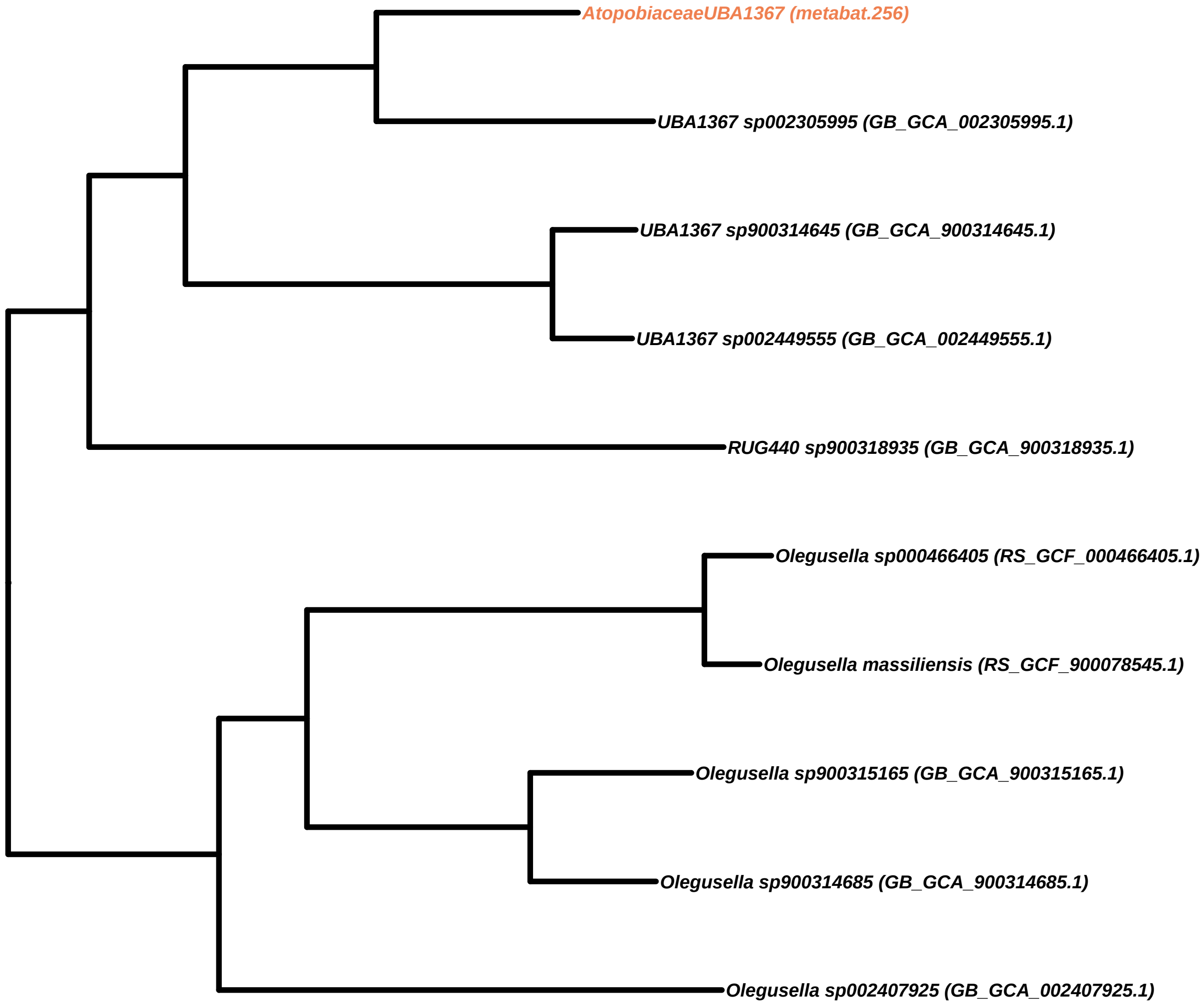
AtopobiaceaeUBA1367 (metabat.256)
UBA1367 sp002305995 (GB_GCA_002305995.1)
UBA1367 sp900314645 (GB_GCA_900314645.1)
UBA1367 sp002449555 (GB_GCA_002449555.1)
RUG440 sp900318935 (GB_GCA_900318935.1)
Olegusella sp000466405 (RS_GCF_000466405.1)
Olegusella massiliensis (RS_GCF_900078545.1)
Olegusella sp900315165 (GB_GCA_900315165.1)
Olegusella sp900314685 (GB_GCA_900314685.1)
Olegusella sp002407925 (GB_GCA_002407925.1)

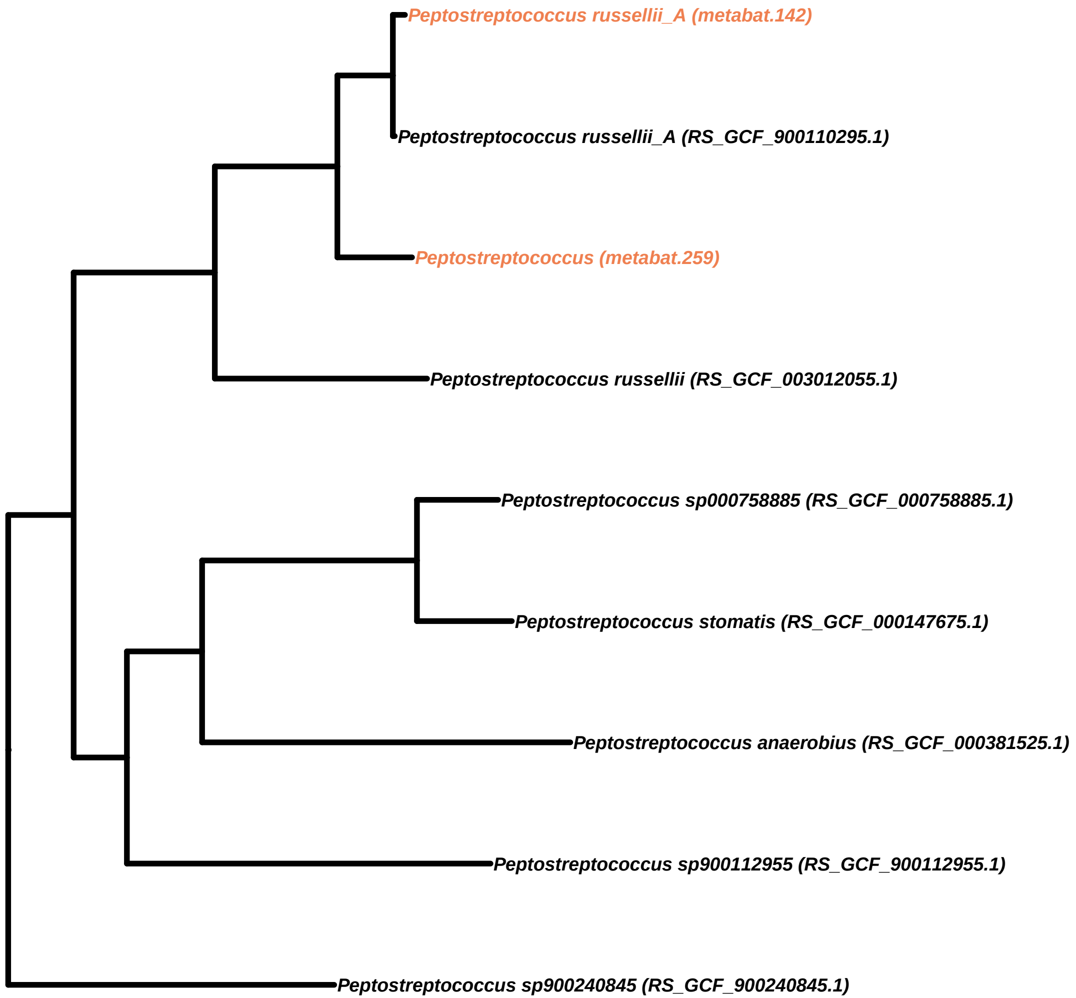

Peptostreptococcus russellii_A (metabat.142)
Peptostreptococcus russellii_A (RS_GCF_900110295.1)
Peptostreptococcus (metabat.259)
Peptostreptococcus russellii (RS_GCF_003012055.1)
Peptostreptococcus sp000758885 (RS_GCF_000758885.1)
Peptostreptococcus stomatis (RS_GCF_000147675.1)
Peptostreptococcus anaerobius (RS_GCF_000381525.1)
Peptostreptococcus sp900112955 (RS_GCF_900112955.1)
Peptostreptococcus sp900240845 (RS_GCF_900240845.1)

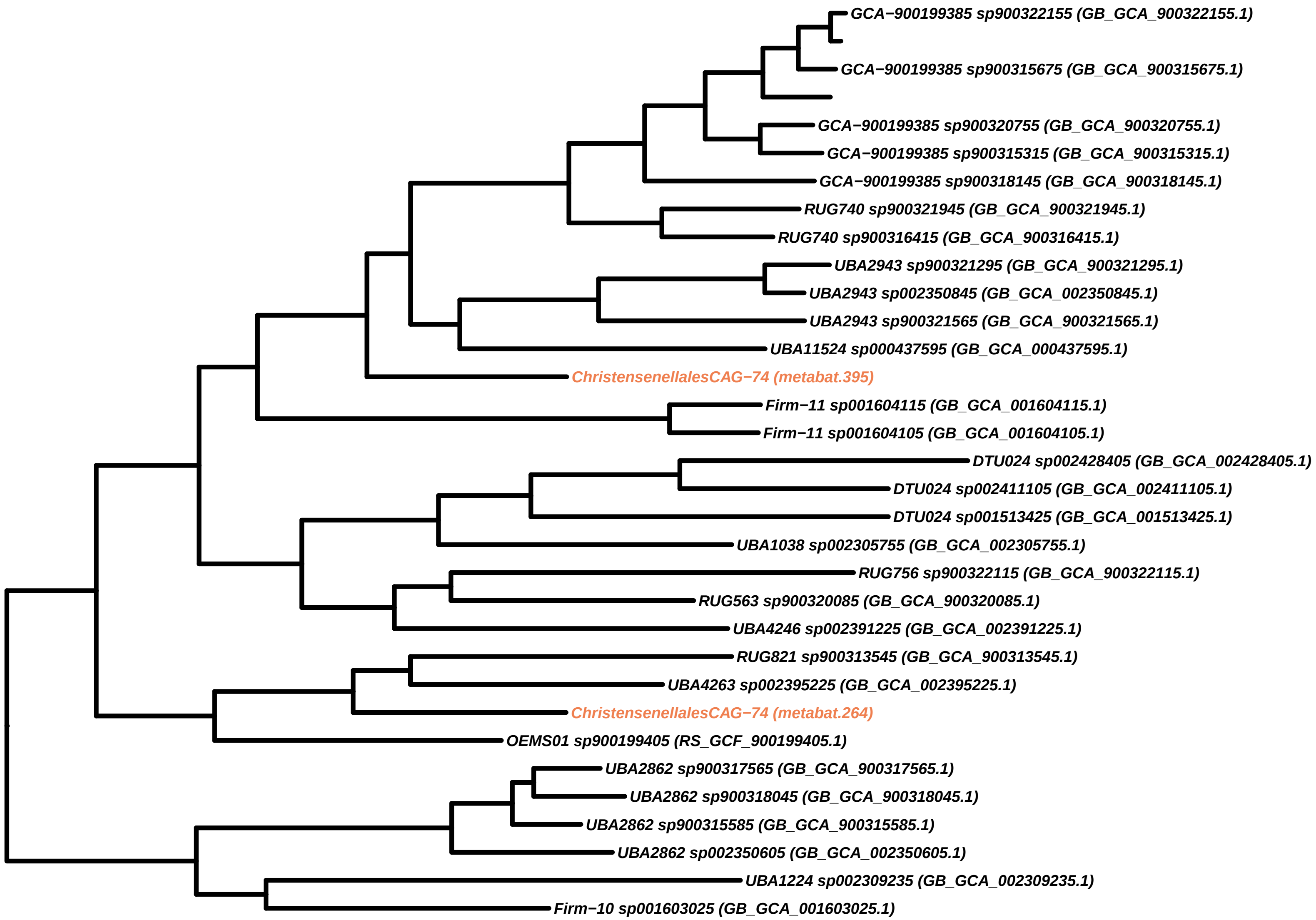
GCA−900199385 sp900322155 (GB_GCA_900322155.1)
GCA−900199385 sp900315675 (GB_GCA_900315675.1)
GCA−900199385 sp900320755 (GB_GCA_900320755.1)
GCA−900199385 sp900315315 (GB_GCA_900315315.1)
GCA−900199385 sp900318145 (GB_GCA_900318145.1)
RUG740 sp900321945 (GB_GCA_900321945.1)
RUG740 sp900316415 (GB_GCA_900316415.1)
UBA2943 sp900321295 (GB_GCA_900321295.1)
UBA2943 sp002350845 (GB_GCA_002350845.1)
UBA2943 sp900321565 (GB_GCA_900321565.1)
UBA11524 sp000437595 (GB_GCA_000437595.1)
ChristensenellalesCAG−74 (metabat.395)
Firm−11 sp001604115 (GB_GCA_001604115.1)
Firm−11 sp001604105 (GB_GCA_001604105.1)
DTU024 sp002428405 (GB_GCA_002428405.1)
DTU024 sp002411105 (GB_GCA_002411105.1)
DTU024 sp001513425 (GB_GCA_001513425.1)
UBA1038 sp002305755 (GB_GCA_002305755.1)
RUG756 sp900322115 (GB_GCA_900322115.1)
RUG563 sp900320085 (GB_GCA_900320085.1)
UBA4246 sp002391225 (GB_GCA_002391225.1)
RUG821 sp900313545 (GB_GCA_900313545.1)
UBA4263 sp002395225 (GB_GCA_002395225.1)
ChristensenellalesCAG−74 (metabat.264)
OEMS01 sp900199405 (RS_GCF_900199405.1)
UBA2862 sp900317565 (GB_GCA_900317565.1)
UBA2862 sp900318045 (GB_GCA_900318045.1)
UBA2862 sp900315585 (GB_GCA_900315585.1)
UBA2862 sp002350605 (GB_GCA_002350605.1)
UBA1224 sp002309235 (GB_GCA_002309235.1)
Firm−10 sp001603025 (GB_GCA_001603025.1)

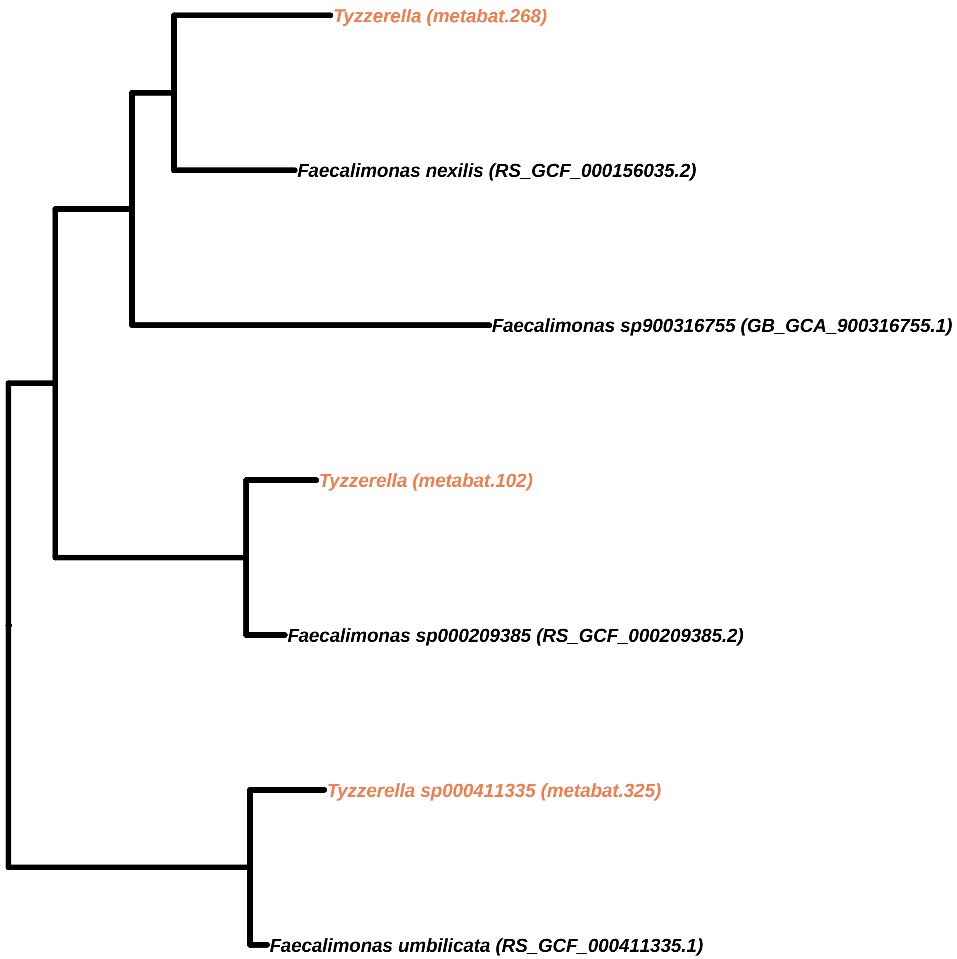
Tyzzerella (metabat.268)
Faecalimonas nexilis (RS_GCF_000156035.2)
Faecalimonas sp900316755 (GB_GCA_900316755.1)
Tyzzerella (metabat.102)
Faecalimonas sp000209385 (RS_GCF_000209385.2)
Tyzzerella sp000411335 (metabat.325)
Faecalimonas umbilicata (RS_GCF_000411335.1)

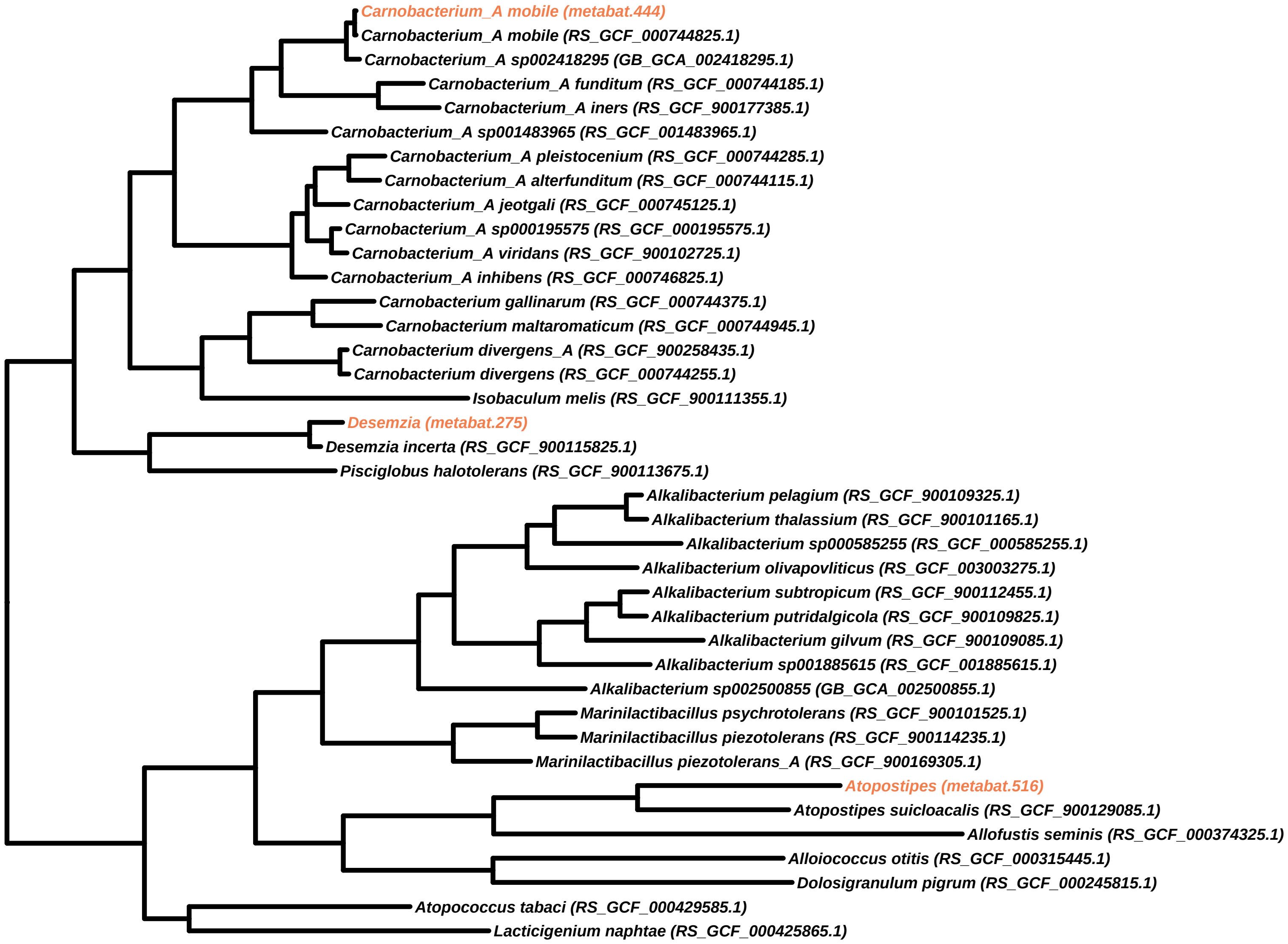
Carnobacterium_A mobile (metabat.444)
Carnobacterium_A mobile (RS_GCF_000744825.1)
Carnobacterium_A sp002418295 (GB_GCA_002418295.1)
Carnobacterium_A funditum (RS_GCF_000744185.1)
Carnobacterium_A iners (RS_GCF_900177385.1)
Carnobacterium_A sp001483965 (RS_GCF_001483965.1)
Carnobacterium_A pleistocenium (RS_GCF_000744285.1)
Carnobacterium_A alterfunditum (RS_GCF_000744115.1)
Carnobacterium_A jeotgali (RS_GCF_000745125.1)
Carnobacterium_A sp000195575 (RS_GCF_000195575.1)
Carnobacterium_A viridans (RS_GCF_900102725.1)
Carnobacterium_A inhibens (RS_GCF_000746825.1)
Carnobacterium gallinarum (RS_GCF_000744375.1)
Carnobacterium maltaromaticum (RS_GCF_000744945.1)
Carnobacterium divergens_A (RS_GCF_900258435.1)
Carnobacterium divergens (RS_GCF_000744255.1)
Isobaculum melis (RS_GCF_900111355.1)
Desemzia (metabat.275)
Desemzia incerta (RS_GCF_900115825.1)
Pisciglobus halotolerans (RS_GCF_900113675.1)
Alkalibacterium pelagium (RS_GCF_900109325.1)
Alkalibacterium thalassium (RS_GCF_900101165.1)
Alkalibacterium sp000585255 (RS_GCF_000585255.1)
Alkalibacterium olivapovliticus (RS_GCF_003003275.1)
Alkalibacterium subtropicum (RS_GCF_900112455.1)
Alkalibacterium putridalgicola (RS_GCF_900109825.1)
Alkalibacterium gilvum (RS_GCF_900109085.1)
Alkalibacterium sp001885615 (RS_GCF_001885615.1)
Alkalibacterium sp002500855 (GB_GCA_002500855.1)
Marinilactibacillus psychrotolerans (RS_GCF_900101525.1)
Marinilactibacillus piezotolerans (RS_GCF_900114235.1)
Marinilactibacillus piezotolerans_A (RS_GCF_900169305.1)
Atopostipes (metabat.516)
Atopostipes suicloacalis (RS_GCF_900129085.1)
Allofustis seminis (RS_GCF_000374325.1)
Alloiococcus otitis (RS_GCF_000315445.1)
Dolosigranulum pigrum (RS_GCF_000245815.1)
Atopococcus tabaci (RS_GCF_000429585.1)
Lacticigenium naphtae (RS_GCF_000425865.1)

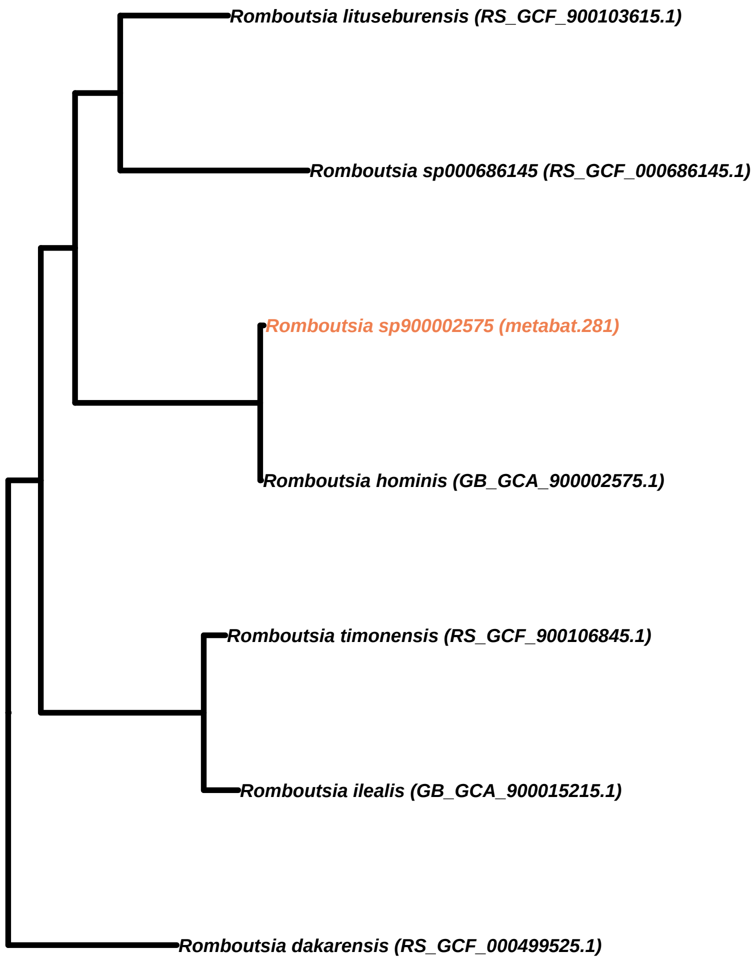
Romboutsia lituseburensis (RS_GCF_900103615.1)
Romboutsia sp000686145 (RS_GCF_000686145.1)
Romboutsia sp900002575 (metabat.281)
Romboutsia hominis (GB_GCA_900002575.1)
Romboutsia timonensis (RS_GCF_900106845.1)
Romboutsia ilealis (GB_GCA_900015215.1)
Romboutsia dakarensis (RS_GCF_000499525.1)

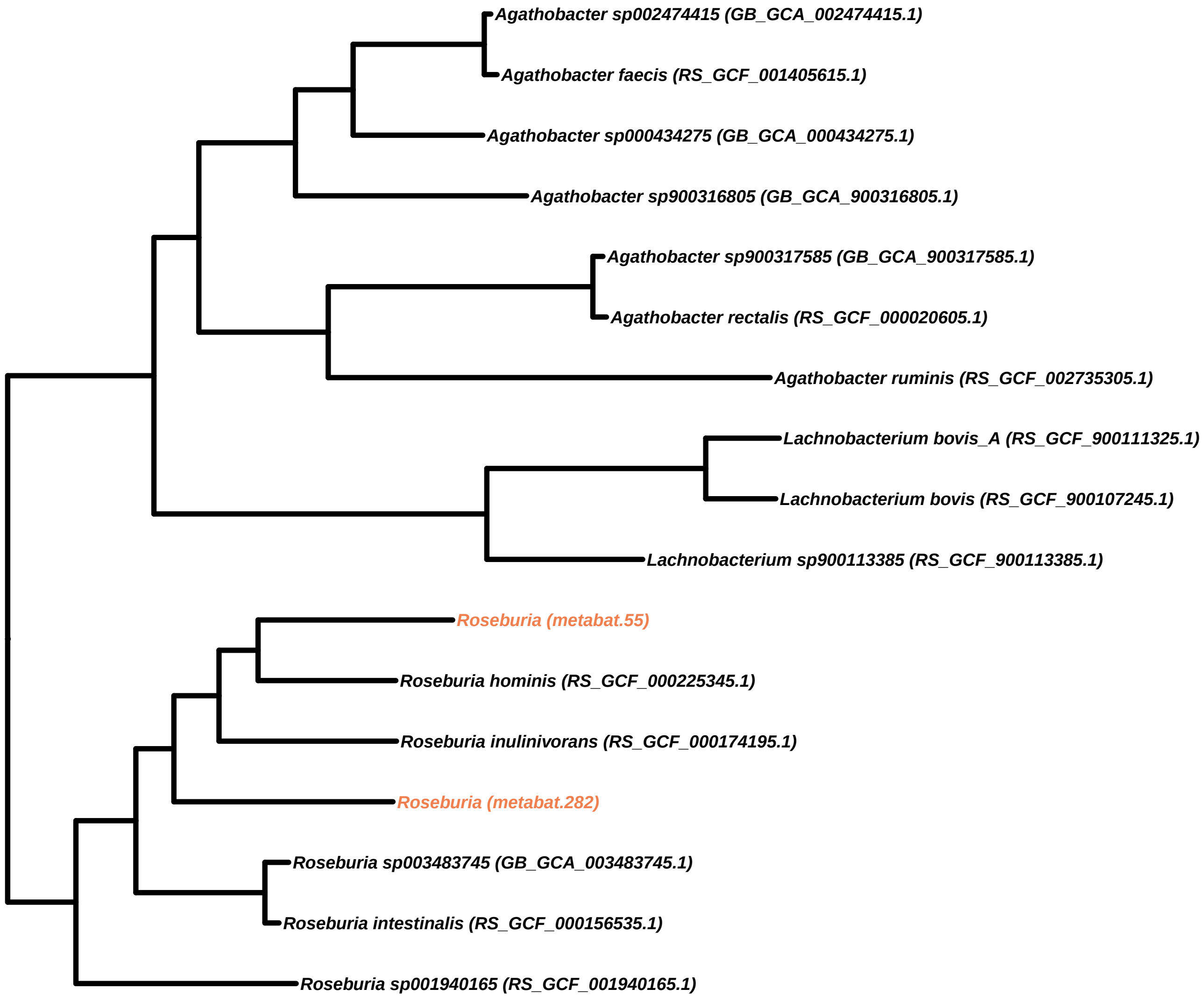
Agathobacter sp002474415 (GB_GCA_002474415.1)
Agathobacter faecis (RS_GCF_001405615.1)
Agathobacter sp000434275 (GB_GCA_000434275.1)
Agathobacter sp900316805 (GB_GCA_900316805.1)
Agathobacter sp900317585 (GB_GCA_900317585.1)
Agathobacter rectalis (RS_GCF_000020605.1)
Agathobacter ruminis (RS_GCF_002735305.1)
Lachnobacterium bovis_A (RS_GCF_900111325.1)
Lachnobacterium bovis (RS_GCF_900107245.1)
Lachnobacterium sp900113385 (RS_GCF_900113385.1)
Roseburia (metabat.55)
Roseburia hominis (RS_GCF_000225345.1)
Roseburia inulinivorans (RS_GCF_000174195.1)
Roseburia (metabat.282)
Roseburia sp003483745 (GB_GCA_003483745.1)
Roseburia intestinalis (RS_GCF_000156535.1)
Roseburia sp001940165 (RS_GCF_001940165.1)

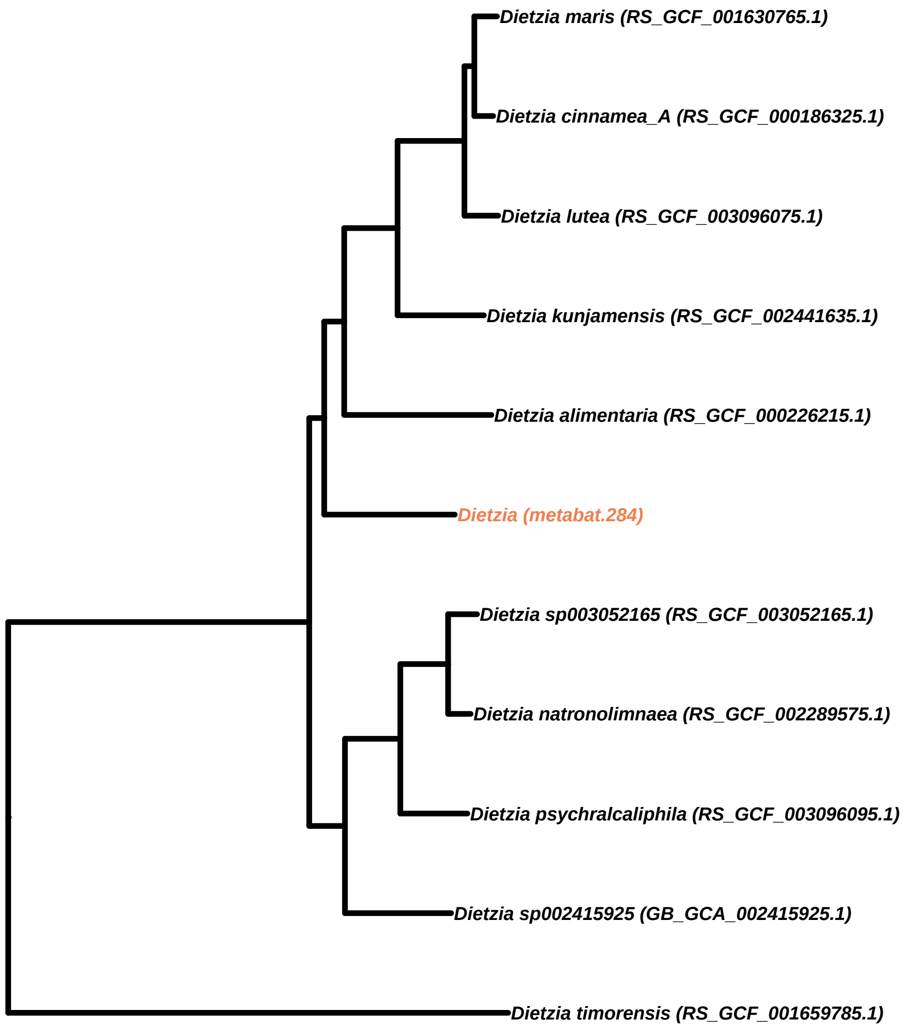
Dietzia maris (RS_GCF_001630765.1)
Dietzia cinnamea_A (RS_GCF_000186325.1)
Dietzia lutea (RS_GCF_003096075.1)
Dietzia kunjamensis (RS_GCF_002441635.1)
Dietzia alimentaria (RS_GCF_000226215.1)
Dietzia (metabat.284)
Dietzia sp003052165 (RS_GCF_003052165.1)
Dietzia natronolimnaea (RS_GCF_002289575.1)
Dietzia psychralcaliphila (RS_GCF_003096095.1)
Dietzia sp002415925 (GB_GCA_002415925.1)
Dietzia timorensis (RS_GCF_001659785.1)

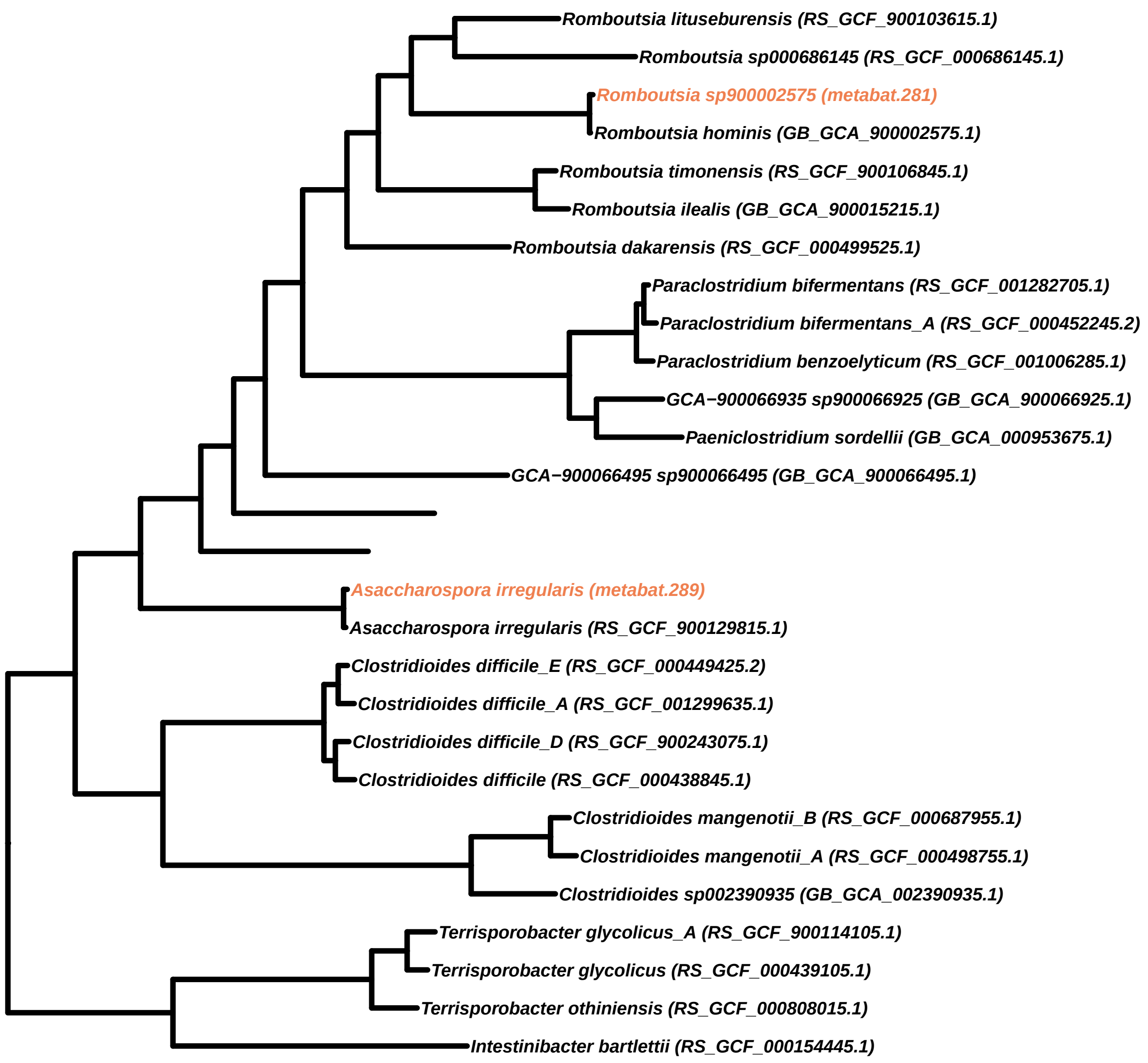
Romboutsia lituseburensis (RS_GCF_900103615.1)
Romboutsia sp000686145 (RS_GCF_000686145.1)
Romboutsia sp900002575 (metabat.281)
Romboutsia hominis (GB_GCA_900002575.1)
Romboutsia timonensis (RS_GCF_900106845.1)
Romboutsia ilealis (GB_GCA_900015215.1)
Romboutsia dakarensis (RS_GCF_000499525.1)
Paraclostridium bifermentans (RS_GCF_001282705.1)
Paraclostridium bifermentans_A (RS_GCF_000452245.2)
Paraclostridium benzoelyticum (RS_GCF_001006285.1)
GCA−900066935 sp900066925 (GB_GCA_900066925.1)
Paeniclostridium sordellii (GB_GCA_000953675.1)
GCA−900066495 sp900066495 (GB_GCA_900066495.1)
Asaccharospora irregularis (metabat.289)
Asaccharospora irregularis (RS_GCF_900129815.1)
Clostridioides difficile_E (RS_GCF_000449425.2)
Clostridioides difficile_A (RS_GCF_001299635.1)
Clostridioides difficile_D (RS_GCF_900243075.1)
Clostridioides difficile (RS_GCF_000438845.1)
Clostridioides mangenotii_B (RS_GCF_000687955.1)
Clostridioides mangenotii_A (RS_GCF_000498755.1)
Clostridioides sp002390935 (GB_GCA_002390935.1)
Terrisporobacter glycolicus_A (RS_GCF_900114105.1)
Terrisporobacter glycolicus (RS_GCF_000439105.1)
Terrisporobacter othiniensis (RS_GCF_000808015.1)
Intestinibacter bartlettii (RS_GCF_000154445.1)

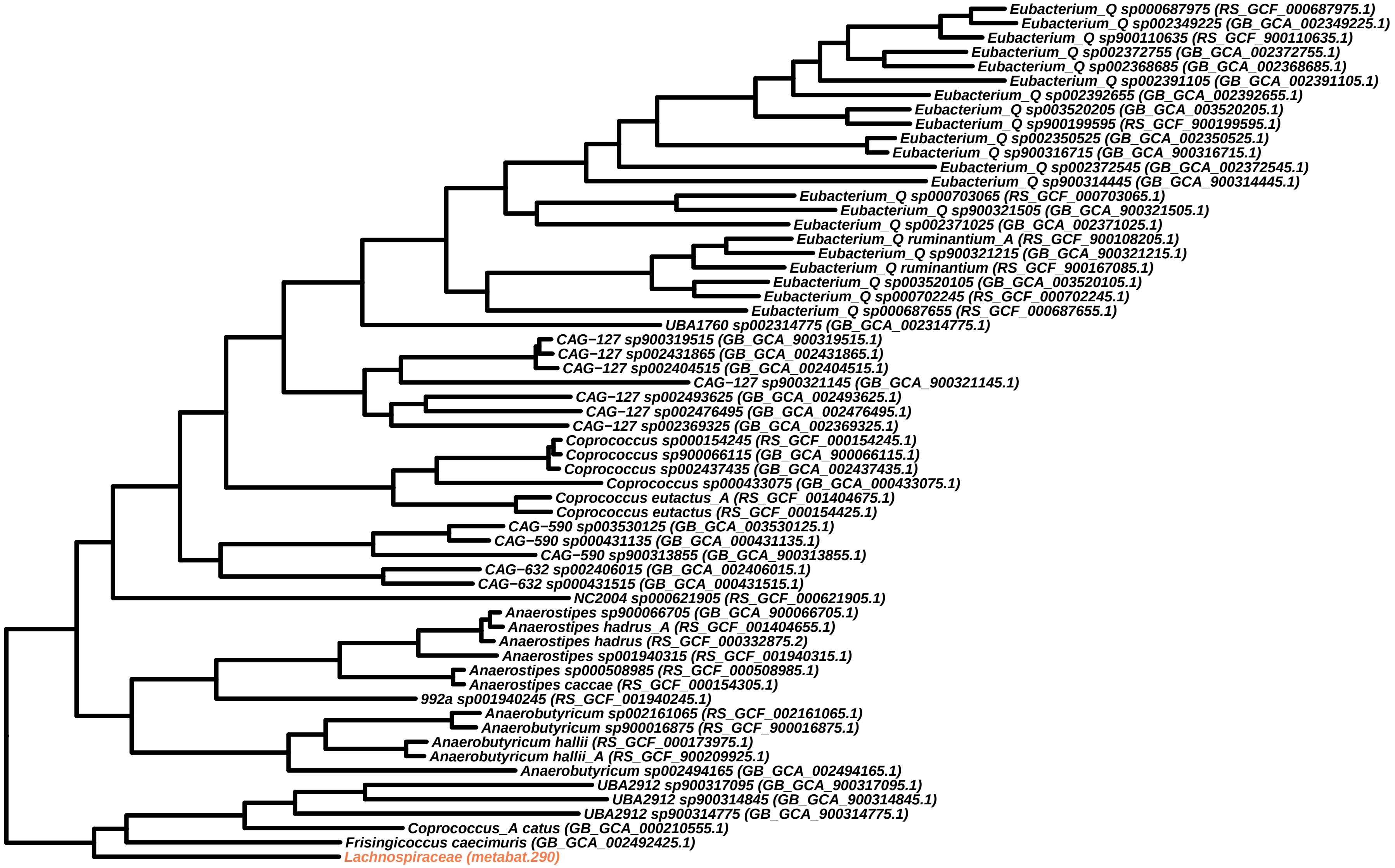
Eubacterium_Q sp000687975 (RS_GCF_000687975.1)
Eubacterium_Q sp002349225 (GB_GCA_002349225.1)
Eubacterium_Q sp900110635 (RS_GCF_900110635.1)
Eubacterium_Q sp002372755 (GB_GCA_002372755.1)
Eubacterium_Q sp002368685 (GB_GCA_002368685.1)
Eubacterium_Q sp002391105 (GB_GCA_002391105.1)
Eubacterium_Q sp002392655 (GB_GCA_002392655.1)
Eubacterium_Q sp003520205 (GB_GCA_003520205.1)
Eubacterium_Q sp900199595 (RS_GCF_900199595.1)
Eubacterium_Q sp002350525 (GB_GCA_002350525.1)
Eubacterium_Q sp900316715 (GB_GCA_900316715.1)
Eubacterium_Q sp002372545 (GB_GCA_002372545.1)
Eubacterium_Q sp900314445 (GB_GCA_900314445.1)
Eubacterium_Q sp000703065 (RS_GCF_000703065.1)
Eubacterium_Q sp900321505 (GB_GCA_900321505.1)
Eubacterium_Q sp002371025 (GB_GCA_002371025.1)
Eubacterium_Q ruminantium_A (RS_GCF_900108205.1)
Eubacterium_Q sp900321215 (GB_GCA_900321215.1)
Eubacterium_Q ruminantium (RS_GCF_900167085.1)
Eubacterium_Q sp003520105 (GB_GCA_003520105.1)
Eubacterium_Q sp000702245 (RS_GCF_000702245.1)
Eubacterium_Q sp000687655 (RS_GCF_000687655.1)
UBA1760 sp002314775 (GB_GCA_002314775.1)
CAG−127 sp900319515 (GB_GCA_900319515.1)
CAG−127 sp002431865 (GB_GCA_002431865.1)
CAG−127 sp002404515 (GB_GCA_002404515.1)
CAG−127 sp900321145 (GB_GCA_900321145.1)
CAG−127 sp002493625 (GB_GCA_002493625.1)
CAG−127 sp002476495 (GB_GCA_002476495.1)
CAG−127 sp002369325 (GB_GCA_002369325.1)
Coprococcus sp000154245 (RS_GCF_000154245.1)
Coprococcus sp900066115 (GB_GCA_900066115.1)
Coprococcus sp002437435 (GB_GCA_002437435.1)
Coprococcus sp000433075 (GB_GCA_000433075.1)
Coprococcus eutactus_A (RS_GCF_001404675.1)
Coprococcus eutactus (RS_GCF_000154425.1)
CAG−590 sp003530125 (GB_GCA_003530125.1)
CAG−590 sp000431135 (GB_GCA_000431135.1)
CAG−590 sp900313855 (GB_GCA_900313855.1)
CAG−632 sp002406015 (GB_GCA_002406015.1)
CAG−632 sp000431515 (GB_GCA_000431515.1)
NC2004 sp000621905 (RS_GCF_000621905.1)
Anaerostipes sp900066705 (GB_GCA_900066705.1)
Anaerostipes hadrus_A (RS_GCF_001404655.1)
Anaerostipes hadrus (RS_GCF_000332875.2)
Anaerostipes sp001940315 (RS_GCF_001940315.1)
Anaerostipes sp000508985 (RS_GCF_000508985.1)
Anaerostipes caccae (RS_GCF_000154305.1)
992a sp001940245 (RS_GCF_001940245.1)
Anaerobutyricum sp002161065 (RS_GCF_002161065.1)
Anaerobutyricum sp900016875 (RS_GCF_900016875.1)
Anaerobutyricum hallii (RS_GCF_000173975.1)
Anaerobutyricum hallii_A (RS_GCF_900209925.1)
Anaerobutyricum sp002494165 (GB_GCA_002494165.1)
UBA2912 sp900317095 (GB_GCA_900317095.1)
UBA2912 sp900314845 (GB_GCA_900314845.1)
UBA2912 sp900314775 (GB_GCA_900314775.1)
Coprococcus_A catus (GB_GCA_000210555.1)
Frisingicoccus caecimuris (GB_GCA_002492425.1)
Lachnospiraceae (metabat.290)

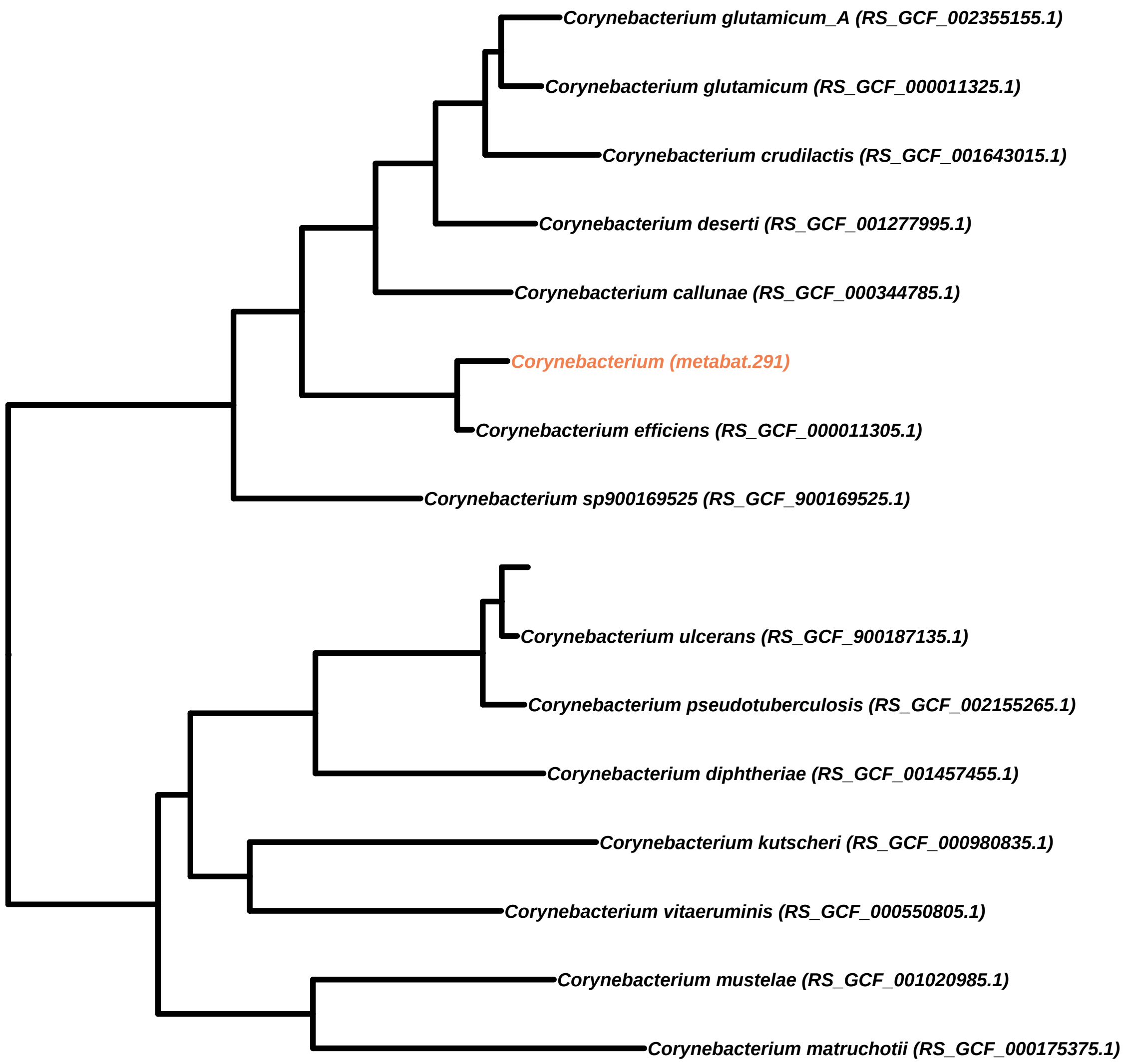
Corynebacterium glutamicum_A (RS_GCF_002355155.1)
Corynebacterium glutamicum (RS_GCF_000011325.1)
Corynebacterium crudilactis (RS_GCF_001643015.1)
Corynebacterium deserti (RS_GCF_001277995.1)
Corynebacterium callunae (RS_GCF_000344785.1)
Corynebacterium (metabat.291)
Corynebacterium efficiens (RS_GCF_000011305.1)
Corynebacterium sp900169525 (RS_GCF_900169525.1)
Corynebacterium ulcerans (RS_GCF_900187135.1)
Corynebacterium pseudotuberculosis (RS_GCF_002155265.1)
Corynebacterium diphtheriae (RS_GCF_001457455.1)
Corynebacterium kutscheri (RS_GCF_000980835.1)
Corynebacterium vitaeruminis (RS_GCF_000550805.1)
Corynebacterium mustelae (RS_GCF_001020985.1)
Corynebacterium matruchotii (RS_GCF_000175375.1)

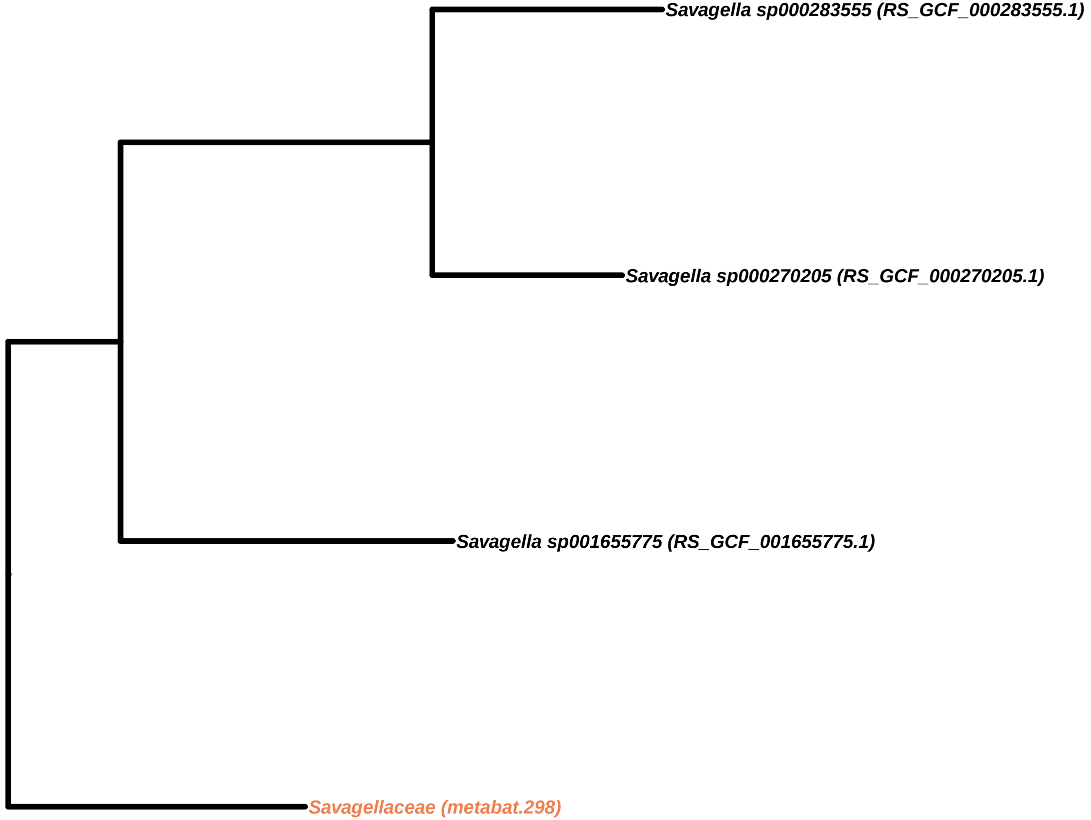
Savagella sp000283555 (RS_GCF_000283555.1)
Savagella sp000270205 (RS_GCF_000270205.1)
Savagella sp001655775 (RS_GCF_001655775.1)
Savagellaceae (metabat.298)

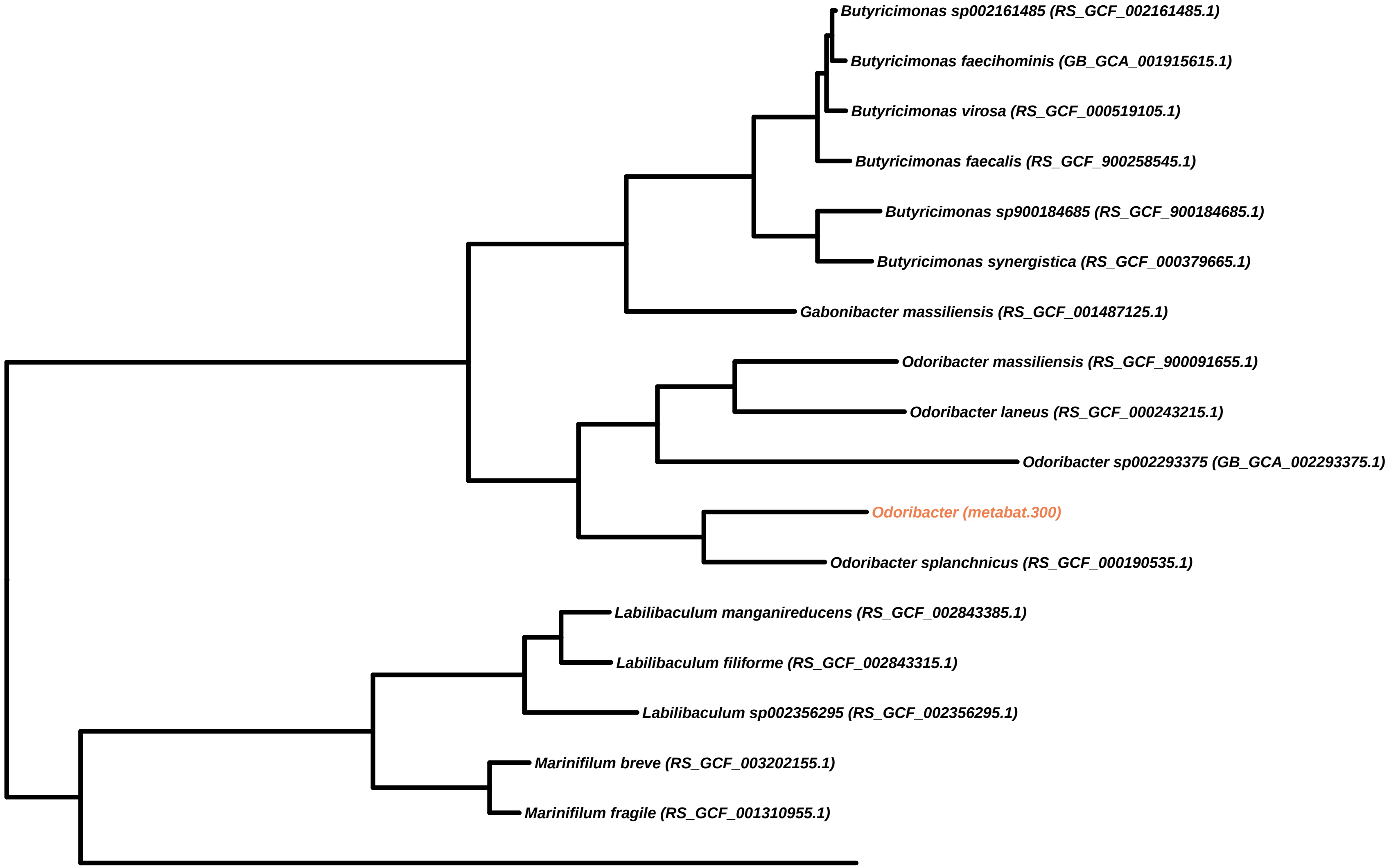

Butyricimonas sp002161485 (RS_GCF_002161485.1)
Butyricimonas faecihominis (GB_GCA_001915615.1)
Butyricimonas virosa (RS_GCF_000519105.1)
Butyricimonas faecalis (RS_GCF_900258545.1)
Butyricimonas sp900184685 (RS_GCF_900184685.1)
Butyricimonas synergistica (RS_GCF_000379665.1)
Gabonibacter massiliensis (RS_GCF_001487125.1)
Odoribacter massiliensis (RS_GCF_900091655.1)
Odoribacter laneus (RS_GCF_000243215.1)
Odoribacter sp002293375 (GB_GCA_002293375.1)
Odoribacter (metabat.300)
Odoribacter splanchnicus (RS_GCF_000190535.1)
Labilibaculum manganireducens (RS_GCF_002843385.1)
Labilibaculum filiforme (RS_GCF_002843315.1)
Labilibaculum sp002356295 (RS_GCF_002356295.1)
Marinifilum breve (RS_GCF_003202155.1)
Marinifilum fragile (RS_GCF_001310955.1)

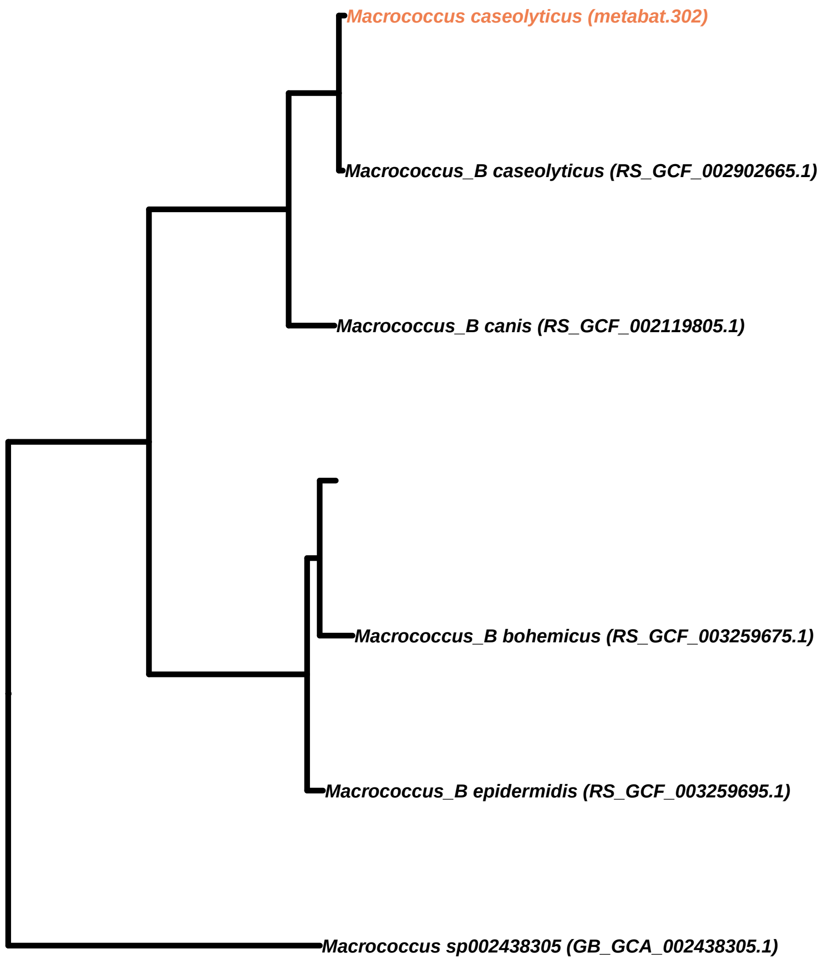

Macrococcus caseolyticus (metabat.302)
Macrococcus_B caseolyticus (RS_GCF_002902665.1)
Macrococcus_B canis (RS_GCF_002119805.1)
Macrococcus_B bohemicus (RS_GCF_003259675.1)
Macrococcus_B epidermidis (RS_GCF_003259695.1)
Macrococcus sp002438305 (GB_GCA_002438305.1)

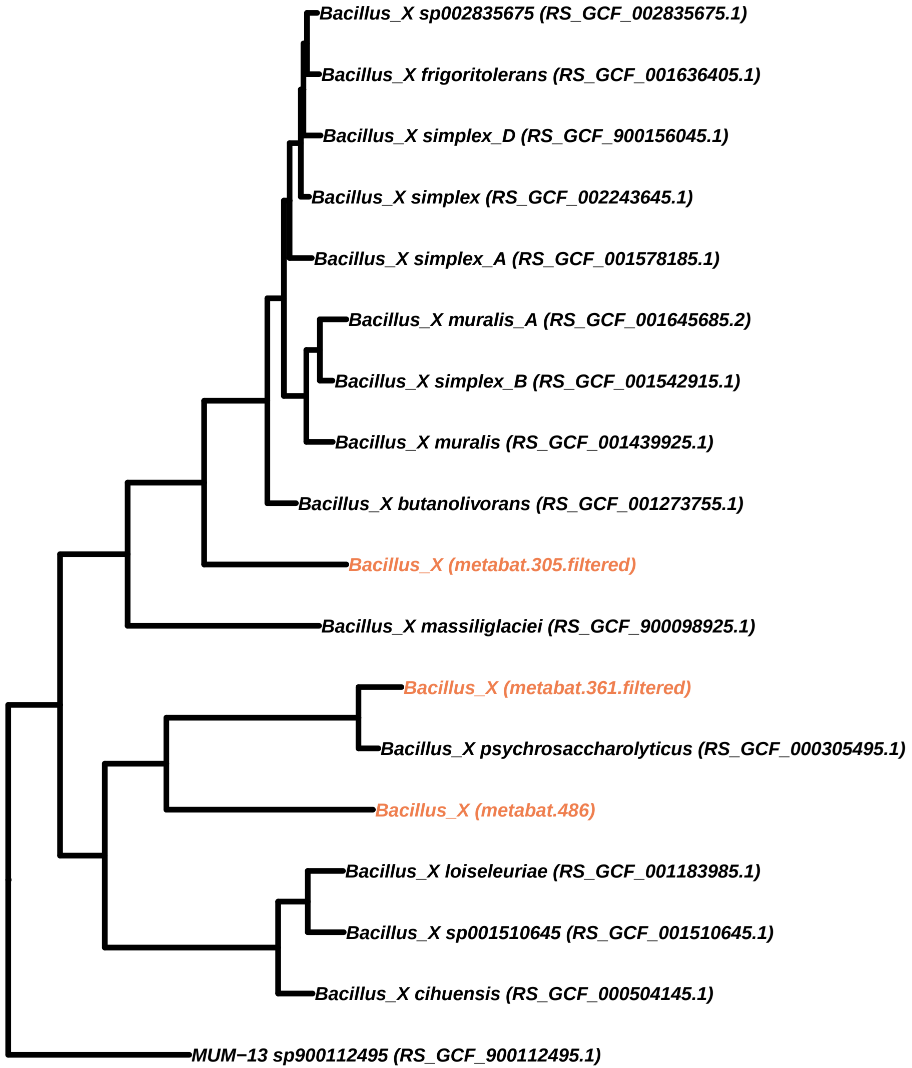
Bacillus_X sp002835675 (RS_GCF_002835675.1)
Bacillus_X frigoritolerans (RS_GCF_001636405.1)
Bacillus_X simplex_D (RS_GCF_900156045.1)
Bacillus_X simplex (RS_GCF_002243645.1)
Bacillus_X simplex_A (RS_GCF_001578185.1)
Bacillus_X muralis_A (RS_GCF_001645685.2)
Bacillus_X simplex_B (RS_GCF_001542915.1)
Bacillus_X muralis (RS_GCF_001439925.1)
Bacillus_X butanolivorans (RS_GCF_001273755.1)
Bacillus_X (metabat.305.filtered)
Bacillus_X massiliglaciei (RS_GCF_900098925.1)
Bacillus_X (metabat.361.filtered)
Bacillus_X psychrosaccharolyticus (RS_GCF_000305495.1)
Bacillus_X (metabat.486)
Bacillus_X loiseleuriae (RS_GCF_001183985.1)
Bacillus_X sp001510645 (RS_GCF_001510645.1)
Bacillus_X cihuensis (RS_GCF_000504145.1)
MUM−13 sp900112495 (RS_GCF_900112495.1)

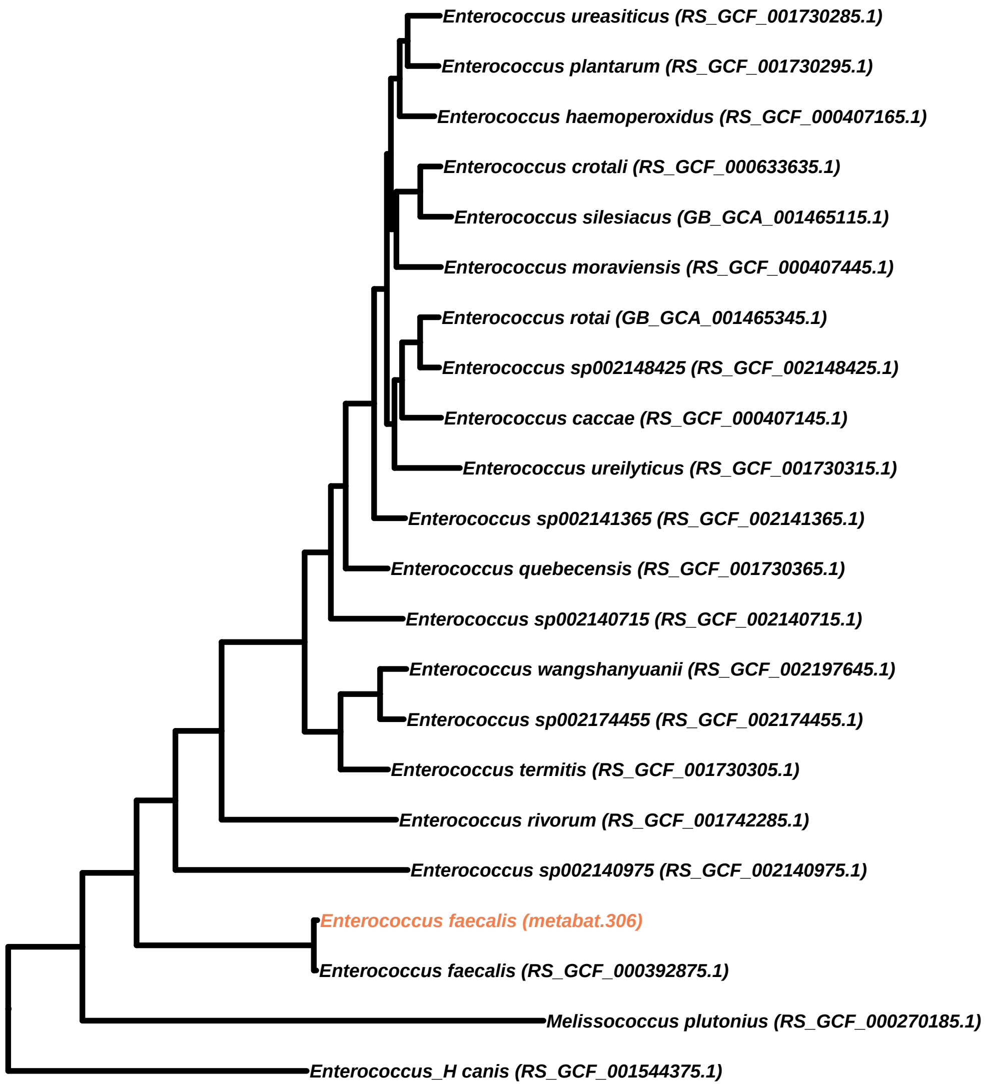

Enterococcus ureasiticus (RS_GCF_001730285.1)
Enterococcus plantarum (RS_GCF_001730295.1)
Enterococcus haemoperoxidus (RS_GCF_000407165.1)
Enterococcus crotali (RS_GCF_000633635.1)
Enterococcus silesiacus (GB_GCA_001465115.1)
Enterococcus moraviensis (RS_GCF_000407445.1)
Enterococcus rotai (GB_GCA_001465345.1)
Enterococcus sp002148425 (RS_GCF_002148425.1)
Enterococcus caccae (RS_GCF_000407145.1)
Enterococcus ureilyticus (RS_GCF_001730315.1)
Enterococcus sp002141365 (RS_GCF_002141365.1)
Enterococcus quebecensis (RS_GCF_001730365.1)
Enterococcus sp002140715 (RS_GCF_002140715.1)
Enterococcus wangshanyuanii (RS_GCF_002197645.1)
Enterococcus sp002174455 (RS_GCF_002174455.1)
Enterococcus termitis (RS_GCF_001730305.1)
Enterococcus rivorum (RS_GCF_001742285.1)
Enterococcus sp002140975 (RS_GCF_002140975.1)
Enterococcus faecalis (metabat.306)
Enterococcus faecalis (RS_GCF_000392875.1)
Melissococcus plutonius (RS_GCF_000270185.1)
Enterococcus_H canis (RS_GCF_001544375.1)

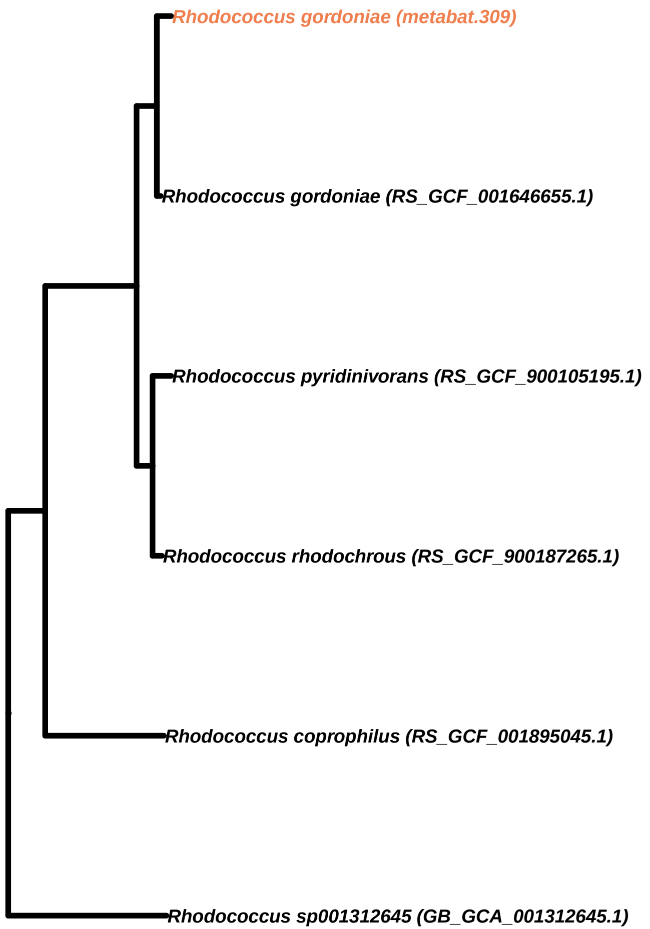
Rhodococcus gordoniae (metabat.309)
Rhodococcus gordoniae (RS_GCF_001646655.1)
Rhodococcus pyridinivorans (RS_GCF_900105195.1)
Rhodococcus rhodochrous (RS_GCF_900187265.1)
Rhodococcus coprophilus (RS_GCF_001895045.1)
Rhodococcus sp001312645 (GB_GCA_001312645.1)

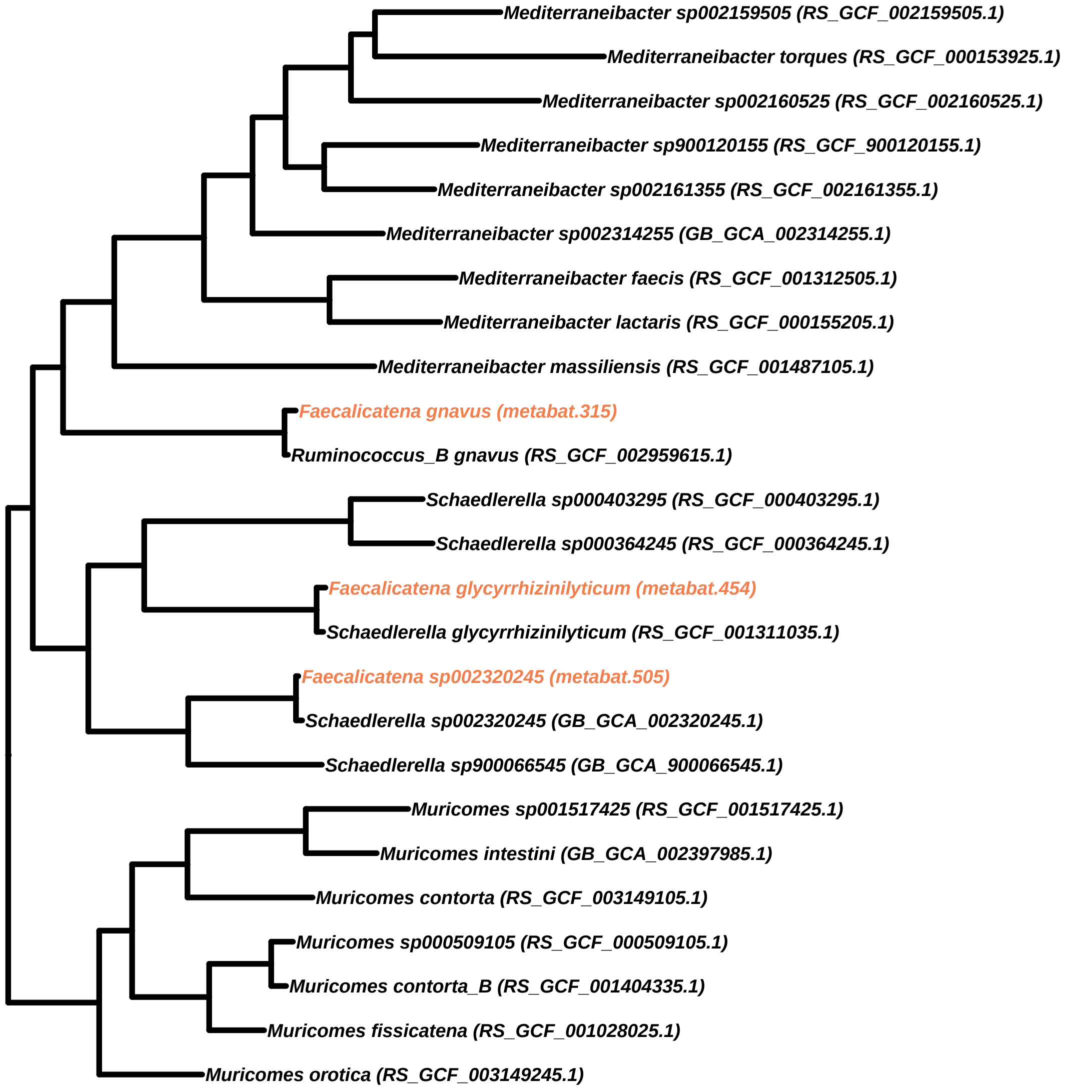
Mediterraneibacter sp002159505 (RS_GCF_002159505.1)
Mediterraneibacter torques (RS_GCF_000153925.1)
Mediterraneibacter sp002160525 (RS_GCF_002160525.1)
Mediterraneibacter sp900120155 (RS_GCF_900120155.1)
Mediterraneibacter sp002161355 (RS_GCF_002161355.1)
Mediterraneibacter sp002314255 (GB_GCA_002314255.1)
Mediterraneibacter faecis (RS_GCF_001312505.1)
Mediterraneibacter lactaris (RS_GCF_000155205.1)
Mediterraneibacter massiliensis (RS_GCF_001487105.1)
Faecalicatena gnavus (metabat.315)
Ruminococcus_B gnavus (RS_GCF_002959615.1)
Schaedlerella sp000403295 (RS_GCF_000403295.1)
Schaedlerella sp000364245 (RS_GCF_000364245.1)
Faecalicatena glycyrrhizinilyticum (metabat.454)
Schaedlerella glycyrrhizinilyticum (RS_GCF_001311035.1)
Faecalicatena sp002320245 (metabat.505)
Schaedlerella sp002320245 (GB_GCA_002320245.1)
Schaedlerella sp900066545 (GB_GCA_900066545.1)
Muricomes sp001517425 (RS_GCF_001517425.1)
Muricomes intestini (GB_GCA_002397985.1)
Muricomes contorta (RS_GCF_003149105.1)
Muricomes sp000509105 (RS_GCF_000509105.1)
Muricomes contorta_B (RS_GCF_001404335.1)
Muricomes fissicatena (RS_GCF_001028025.1)
Muricomes orotica (RS_GCF_003149245.1)

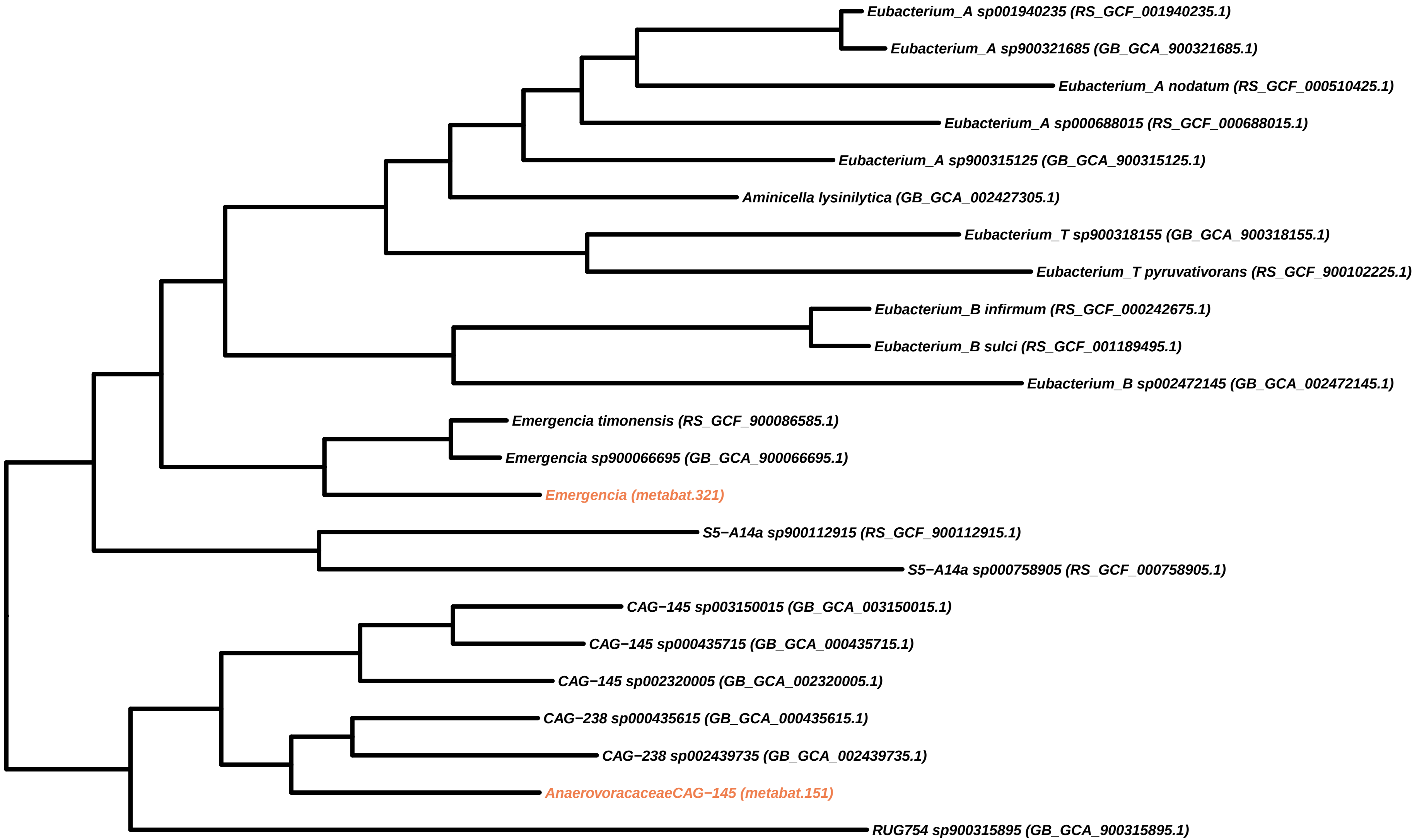

Eubacterium_A sp001940235 (RS_GCF_001940235.1)
Eubacterium_A sp900321685 (GB_GCA_900321685.1)
Eubacterium_A nodatum (RS_GCF_000510425.1)
Eubacterium_A sp000688015 (RS_GCF_000688015.1)
Eubacterium_A sp900315125 (GB_GCA_900315125.1)
Aminicella lysinilytica (GB_GCA_002427305.1)
Eubacterium_T sp900318155 (GB_GCA_900318155.1)
Eubacterium_T pyruvativorans (RS_GCF_900102225.1)
Eubacterium_B infirmum (RS_GCF_000242675.1)
Eubacterium_B sulci (RS_GCF_001189495.1)
Eubacterium_B sp002472145 (GB_GCA_002472145.1)
Emergencia timonensis (RS_GCF_900086585.1)
Emergencia sp900066695 (GB_GCA_900066695.1)
Emergencia (metabat.321)
S5−A14a sp900112915 (RS_GCF_900112915.1)
S5−A14a sp000758905 (RS_GCF_000758905.1)
CAG−145 sp003150015 (GB_GCA_003150015.1)
CAG−145 sp000435715 (GB_GCA_000435715.1)
CAG−145 sp002320005 (GB_GCA_002320005.1)
CAG−238 sp000435615 (GB_GCA_000435615.1)
CAG−238 sp002439735 (GB_GCA_002439735.1)
AnaerovoracaceaeCAG−145 (metabat.151)
RUG754 sp900315895 (GB_GCA_900315895.1)

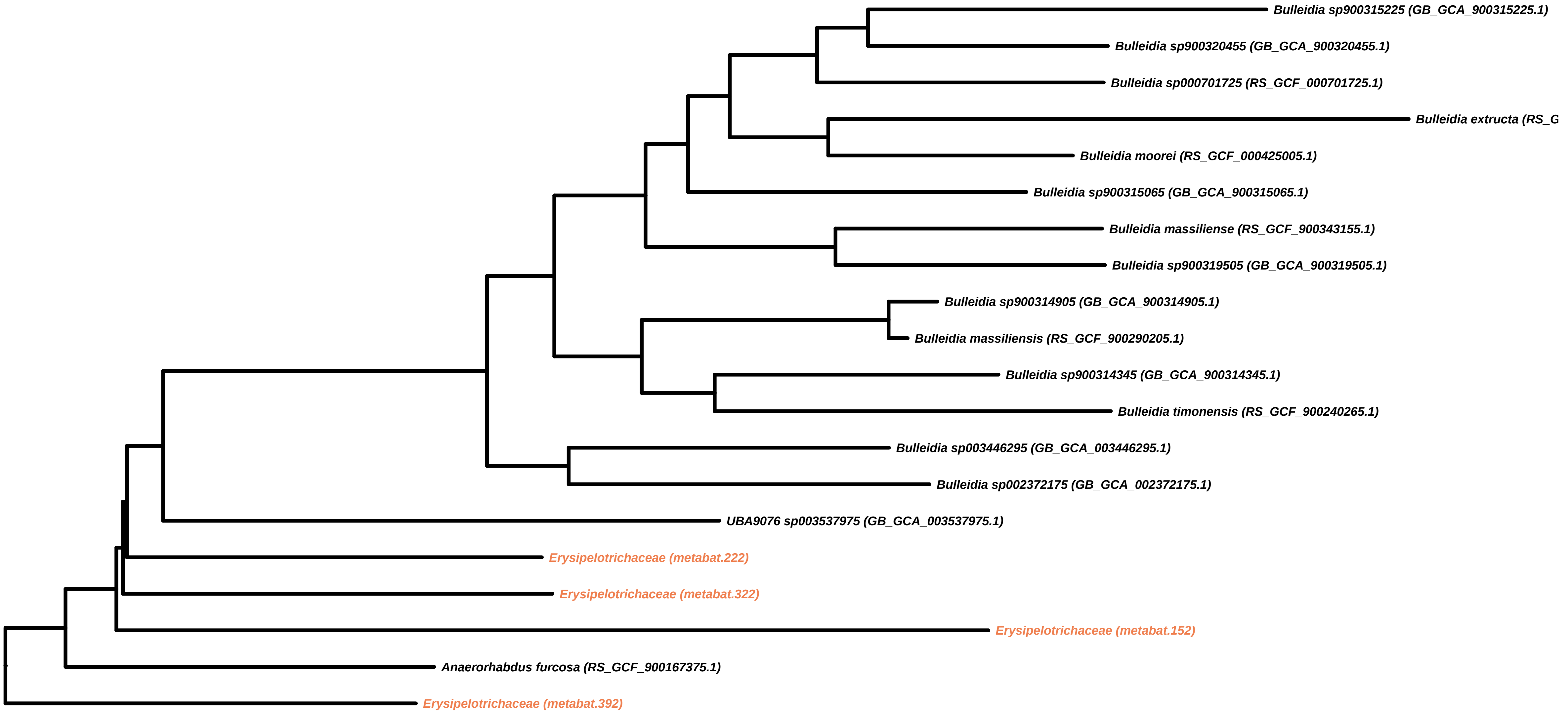

Bulleidia sp900315225 (GB_GCA_900315225.1)
Bulleidia sp900320455 (GB_GCA_900320455.1)
Bulleidia sp000701725 (RS_GCF_000701725.1)
Bulleidia extructa (RS_G
Bulleidia moorei (RS_GCF_000425005.1)
Bulleidia sp900315065 (GB_GCA_900315065.1)
Bulleidia massiliense (RS_GCF_900343155.1)
Bulleidia sp900319505 (GB_GCA_900319505.1)
Bulleidia sp900314905 (GB_GCA_900314905.1)
Bulleidia massiliensis (RS_GCF_900290205.1)
Bulleidia sp900314345 (GB_GCA_900314345.1)
Bulleidia timonensis (RS_GCF_900240265.1)
Bulleidia sp003446295 (GB_GCA_003446295.1)
Bulleidia sp002372175 (GB_GCA_002372175.1)
UBA9076 sp003537975 (GB_GCA_003537975.1)
Erysipelotrichaceae (metabat.222)
Erysipelotrichaceae (metabat.322)
Erysipelotrichaceae (metabat.152)
Anaerorhabdus furcosa (RS_GCF_900167375.1)
Erysipelotrichaceae (metabat.392)

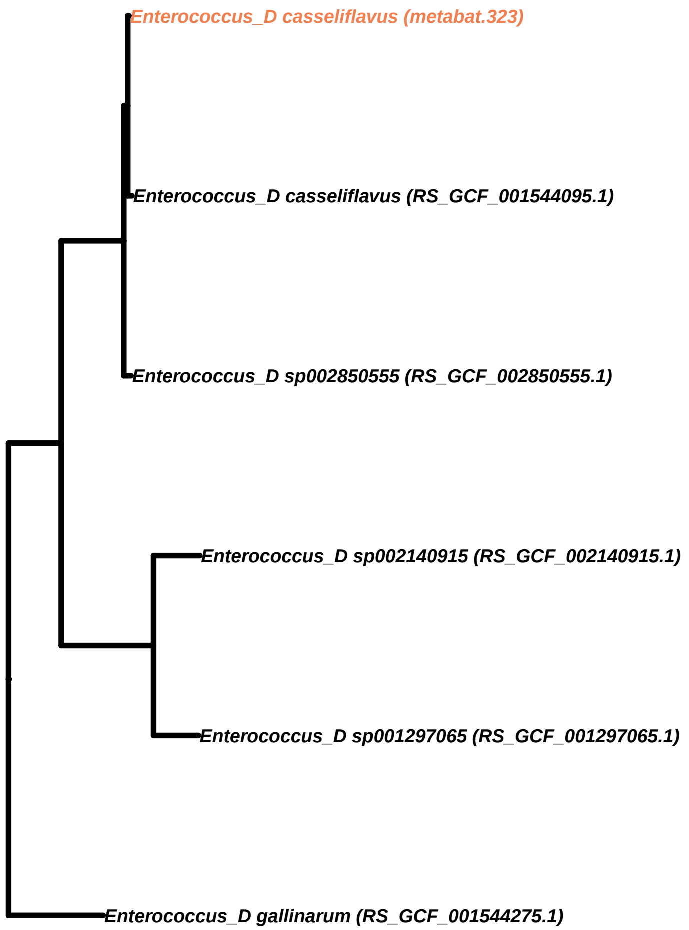
Enterococcus_D casseliflavus (metabat.323)
Enterococcus_D casseliflavus (RS_GCF_001544095.1)
Enterococcus_D sp002850555 (RS_GCF_002850555.1)
Enterococcus_D sp002140915 (RS_GCF_002140915.1)
Enterococcus_D sp001297065 (RS_GCF_001297065.1)
Enterococcus_D gallinarum (RS_GCF_001544275.1)

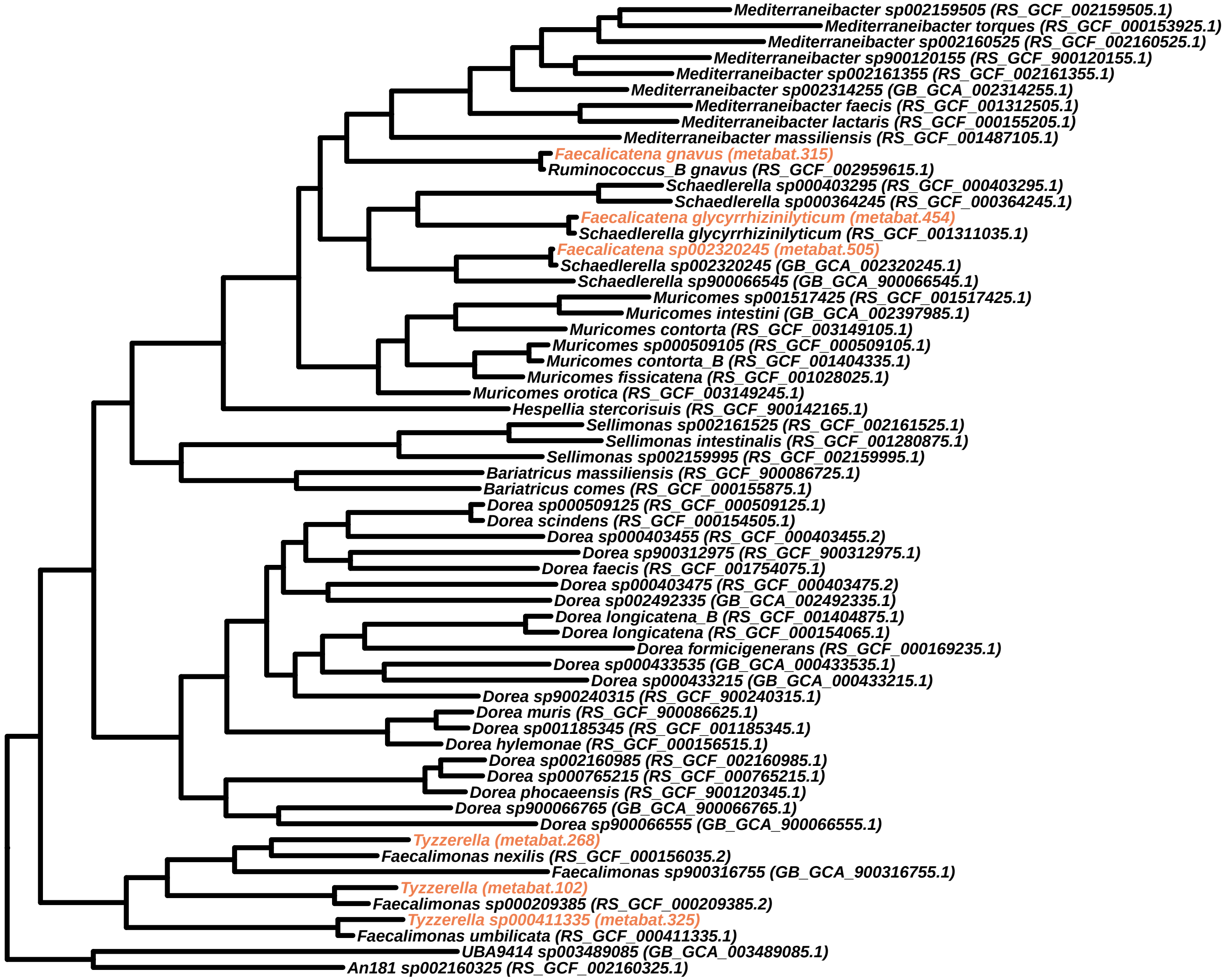
Mediterraneibacter sp002159505 (RS_GCF_002159505.1)
Mediterraneibacter torques (RS_GCF_000153925.1)
Mediterraneibacter sp002160525 (RS_GCF_002160525.1)
Mediterraneibacter sp900120155 (RS_GCF_900120155.1)
Mediterraneibacter sp002161355 (RS_GCF_002161355.1)
Mediterraneibacter sp002314255 (GB_GCA_002314255.1)
Mediterraneibacter faecis (RS_GCF_001312505.1)
Mediterraneibacter lactaris (RS_GCF_000155205.1)
Mediterraneibacter massiliensis (RS_GCF_001487105.1)
Faecalicatena gnavus (metabat.315)
Ruminococcus_B gnavus (RS_GCF_002959615.1)
Schaedlerella sp000403295 (RS_GCF_000403295.1)
Schaedlerella sp000364245 (RS_GCF_000364245.1)
Faecalicatena glycyrrhizinilyticum (metabat.454)
Schaedlerella glycyrrhizinilyticum (RS_GCF_001311035.1)
Faecalicatena sp002320245 (metabat.505)
Schaedlerella sp002320245 (GB_GCA_002320245.1)
Schaedlerella sp900066545 (GB_GCA_900066545.1)
Muricomes sp001517425 (RS_GCF_001517425.1)
Muricomes intestini (GB_GCA_002397985.1)
Muricomes contorta (RS_GCF_003149105.1)
Muricomes sp000509105 (RS_GCF_000509105.1)
Muricomes contorta_B (RS_GCF_001404335.1)
Muricomes fissicatena (RS_GCF_001028025.1)
Muricomes orotica (RS_GCF_003149245.1)
Hespellia stercorisuis (RS_GCF_900142165.1)
Sellimonas sp002161525 (RS_GCF_002161525.1)
Sellimonas intestinalis (RS_GCF_001280875.1)
Sellimonas sp002159995 (RS_GCF_002159995.1)
Bariatricus massiliensis (RS_GCF_900086725.1)
Bariatricus comes (RS_GCF_000155875.1)
Dorea sp000509125 (RS_GCF_000509125.1)
Dorea scindens (RS_GCF_000154505.1)
Dorea sp000403455 (RS_GCF_000403455.2)
Dorea sp900312975 (RS_GCF_900312975.1)
Dorea faecis (RS_GCF_001754075.1)
Dorea sp000403475 (RS_GCF_000403475.2)
Dorea sp002492335 (GB_GCA_002492335.1)
Dorea longicatena_B (RS_GCF_001404875.1)
Dorea longicatena (RS_GCF_000154065.1)
Dorea formicigenerans (RS_GCF_000169235.1)
Dorea sp000433535 (GB_GCA_000433535.1)
Dorea sp000433215 (GB_GCA_000433215.1)
Dorea sp900240315 (RS_GCF_900240315.1)
Dorea muris (RS_GCF_900086625.1)
Dorea sp001185345 (RS_GCF_001185345.1)
Dorea hylemonae (RS_GCF_000156515.1)
Dorea sp002160985 (RS_GCF_002160985.1)
Dorea sp000765215 (RS_GCF_000765215.1)
Dorea phocaeensis (RS_GCF_900120345.1)
Dorea sp900066765 (GB_GCA_900066765.1)
Dorea sp900066555 (GB_GCA_900066555.1)
Tyzzerella (metabat.268)
Faecalimonas nexilis (RS_GCF_000156035.2)
Faecalimonas sp900316755 (GB_GCA_900316755.1)
Tyzzerella (metabat.102)
Faecalimonas sp000209385 (RS_GCF_000209385.2)
Tyzzerella sp000411335 (metabat.325)
Faecalimonas umbilicata (RS_GCF_000411335.1)
UBA9414 sp003489085 (GB_GCA_003489085.1)
An181 sp002160325 (RS_GCF_002160325.1)

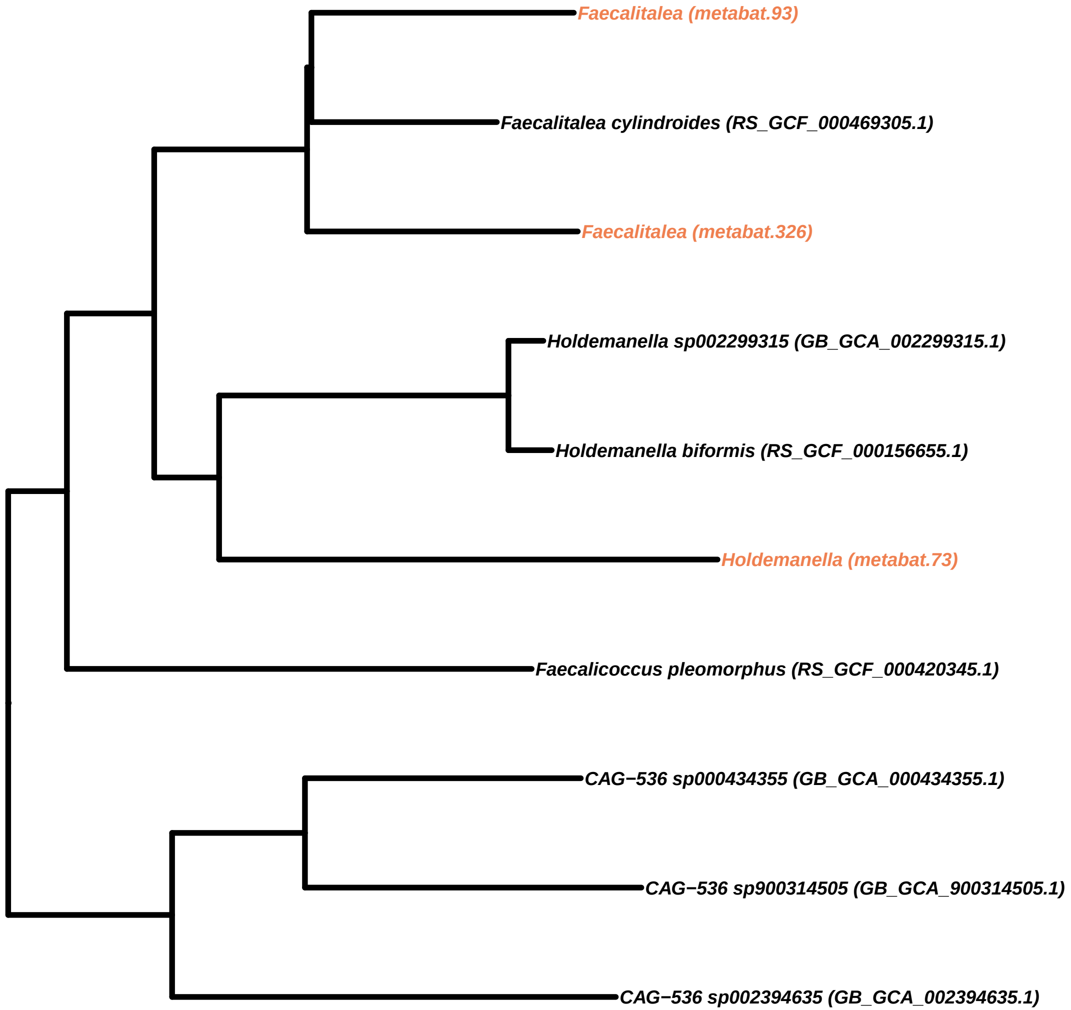
Faecalitalea (metabat.93)
Faecalitalea cylindroides (RS_GCF_000469305.1)
Faecalitalea (metabat.326)
Holdemanella sp002299315 (GB_GCA_002299315.1)
Holdemanella biformis (RS_GCF_000156655.1)
Holdemanella (metabat.73)
Faecalicoccus pleomorphus (RS_GCF_000420345.1)
CAG−536 sp000434355 (GB_GCA_000434355.1)
CAG−536 sp900314505 (GB_GCA_900314505.1)
CAG−536 sp002394635 (GB_GCA_002394635.1)

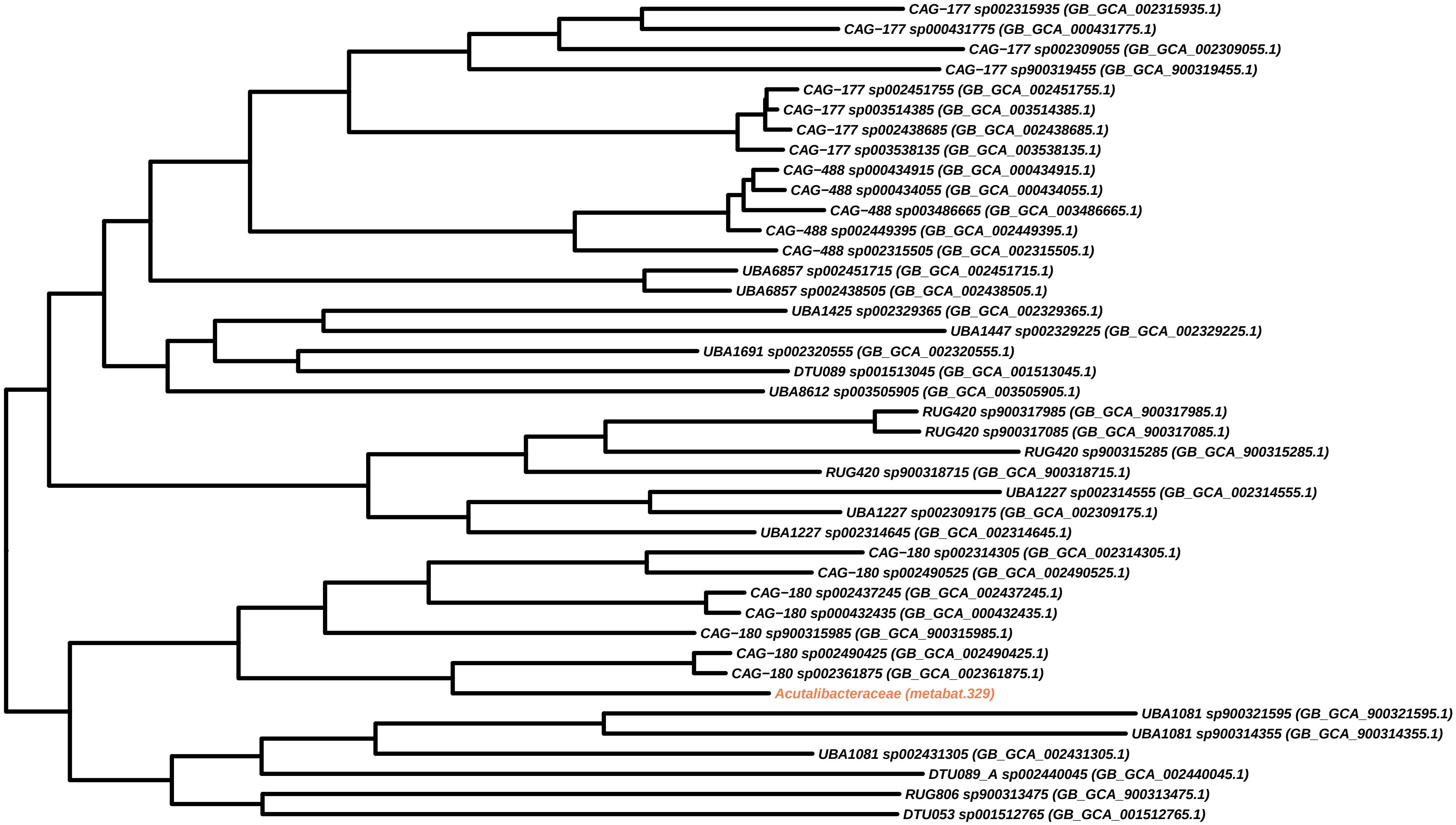

CAG−177 sp002315935 (GB_GCA_002315935.1)
CAG−177 sp000431775 (GB_GCA_000431775.1)
CAG−177 sp002309055 (GB_GCA_002309055.1)
CAG−177 sp900319455 (GB_GCA_900319455.1)
CAG−177 sp002451755 (GB_GCA_002451755.1)
CAG−177 sp003514385 (GB_GCA_003514385.1)
CAG−177 sp002438685 (GB_GCA_002438685.1)
CAG−177 sp003538135 (GB_GCA_003538135.1)
CAG−488 sp000434915 (GB_GCA_000434915.1)
CAG−488 sp000434055 (GB_GCA_000434055.1)
CAG−488 sp003486665 (GB_GCA_003486665.1)
CAG−488 sp002449395 (GB_GCA_002449395.1)
CAG−488 sp002315505 (GB_GCA_002315505.1)
UBA6857 sp002451715 (GB_GCA_002451715.1)
UBA6857 sp002438505 (GB_GCA_002438505.1)
UBA1425 sp002329365 (GB_GCA_002329365.1)
UBA1447 sp002329225 (GB_GCA_002329225.1)
UBA1691 sp002320555 (GB_GCA_002320555.1)
DTU089 sp001513045 (GB_GCA_001513045.1)
UBA8612 sp003505905 (GB_GCA_003505905.1)
RUG420 sp900317985 (GB_GCA_900317985.1)
RUG420 sp900317085 (GB_GCA_900317085.1)
RUG420 sp900315285 (GB_GCA_900315285.1)
RUG420 sp900318715 (GB_GCA_900318715.1)
UBA1227 sp002314555 (GB_GCA_002314555.1)
UBA1227 sp002309175 (GB_GCA_002309175.1)
UBA1227 sp002314645 (GB_GCA_002314645.1)
CAG−180 sp002314305 (GB_GCA_002314305.1)
CAG−180 sp002490525 (GB_GCA_002490525.1)
CAG−180 sp002437245 (GB_GCA_002437245.1)
CAG−180 sp000432435 (GB_GCA_000432435.1)
CAG−180 sp900315985 (GB_GCA_900315985.1)
CAG−180 sp002490425 (GB_GCA_002490425.1)
CAG−180 sp002361875 (GB_GCA_002361875.1)
Acutalibacteraceae (metabat.329)
UBA1081 sp900321595 (GB_GCA_900321595.1)
UBA1081 sp900314355 (GB_GCA_900314355.1)
UBA1081 sp002431305 (GB_GCA_002431305.1)
DTU089_A sp002440045 (GB_GCA_002440045.1)
RUG806 sp900313475 (GB_GCA_900313475.1)
DTU053 sp001512765 (GB_GCA_001512765.1)

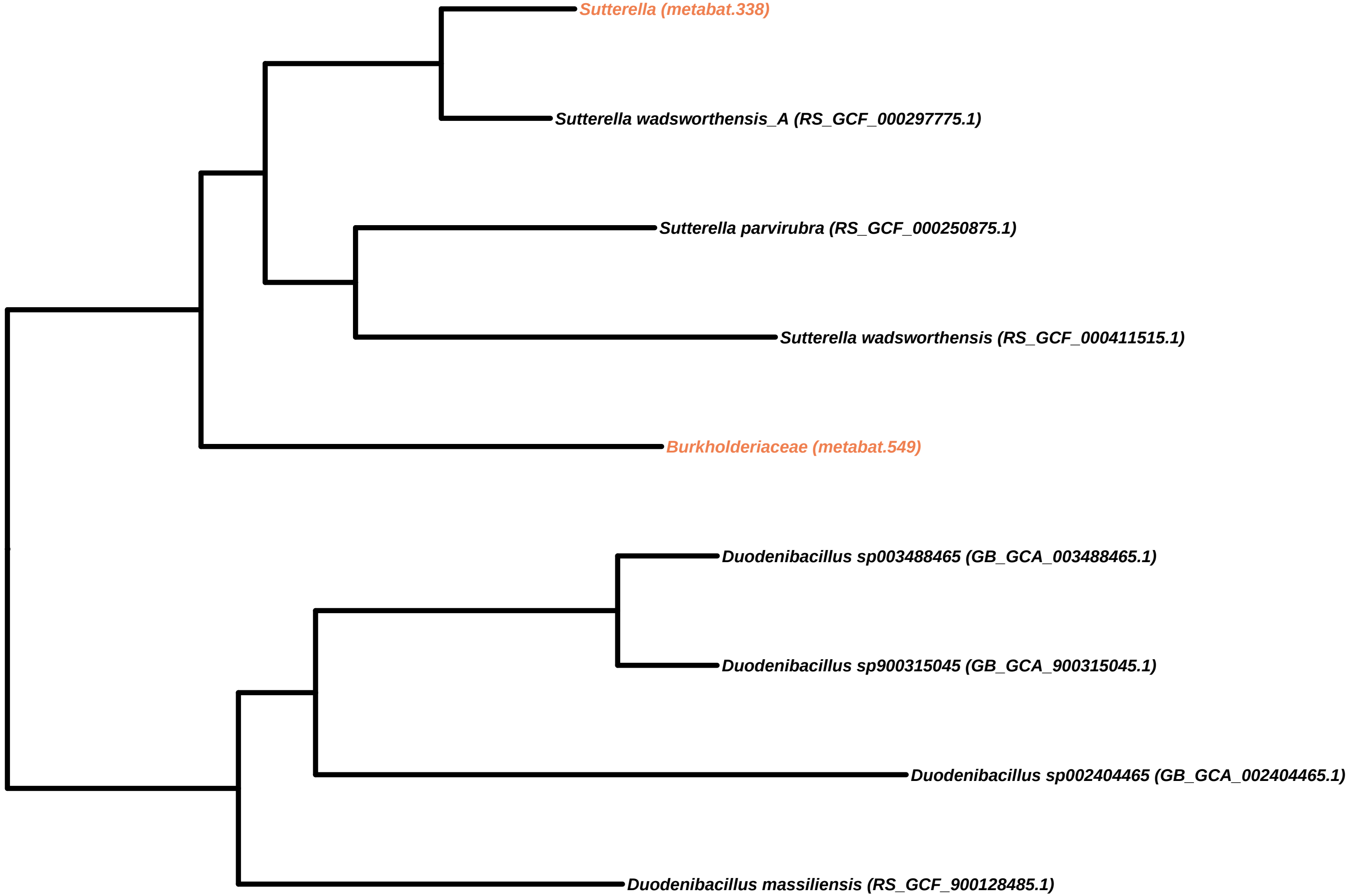
Sutterella (metabat.338)
Sutterella wadsworthensis_A (RS_GCF_000297775.1)
Sutterella parvirubra (RS_GCF_000250875.1)
Sutterella wadsworthensis (RS_GCF_000411515.1)
Burkholderiaceae (metabat.549)
Duodenibacillus sp003488465 (GB_GCA_003488465.1)
Duodenibacillus sp900315045 (GB_GCA_900315045.1)
Duodenibacillus sp002404465 (GB_GCA_002404465.1)
Duodenibacillus massiliensis (RS_GCF_900128485.1)

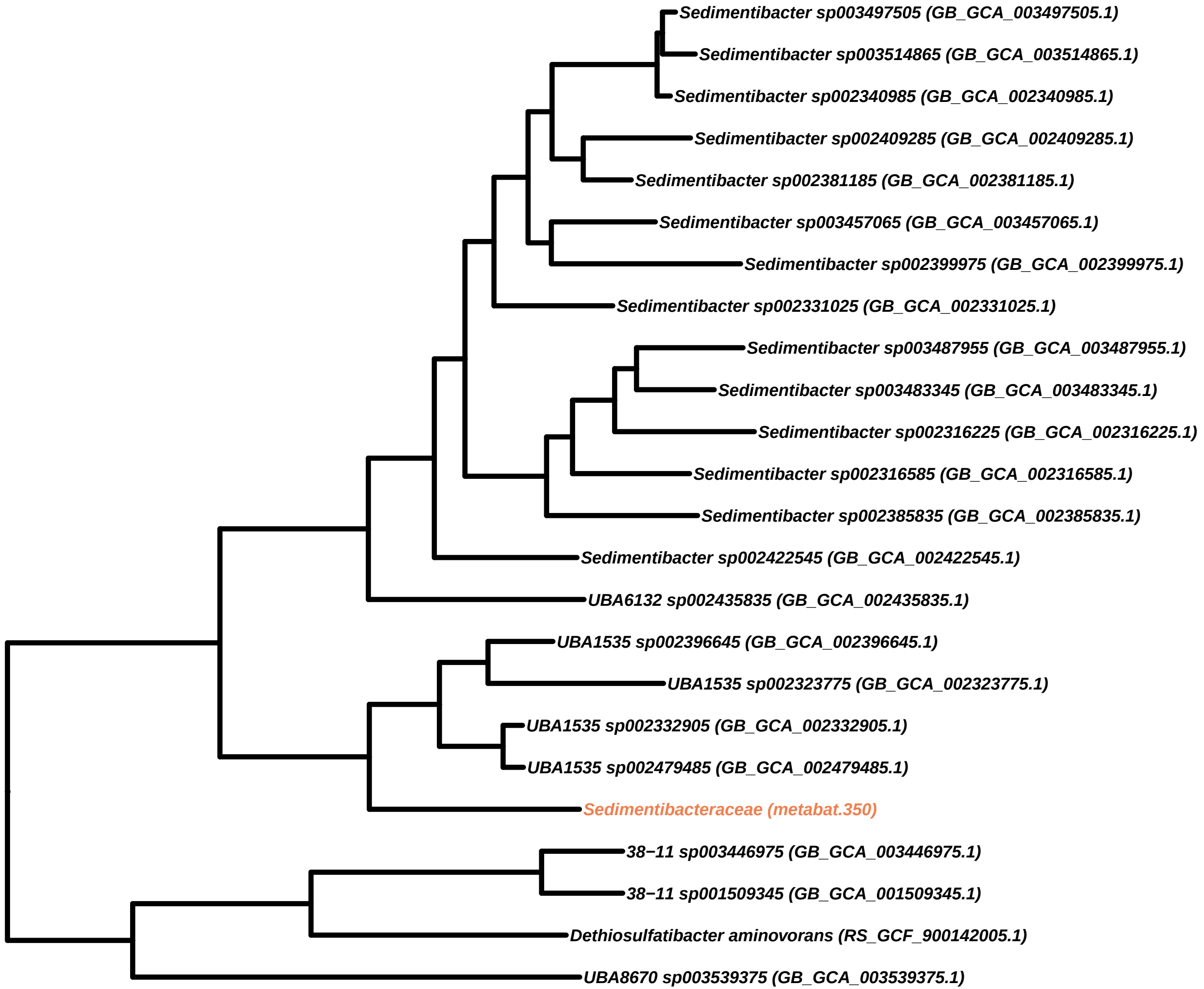
Sedimentibacter sp003497505 (GB_GCA_003497505.1)
Sedimentibacter sp003514865 (GB_GCA_003514865.1)
Sedimentibacter sp002340985 (GB_GCA_002340985.1)
Sedimentibacter sp002409285 (GB_GCA_002409285.1)
Sedimentibacter sp002381185 (GB_GCA_002381185.1)
Sedimentibacter sp003457065 (GB_GCA_003457065.1)
Sedimentibacter sp002399975 (GB_GCA_002399975.1)
Sedimentibacter sp002331025 (GB_GCA_002331025.1)
Sedimentibacter sp003487955 (GB_GCA_003487955.1)
Sedimentibacter sp003483345 (GB_GCA_003483345.1)
Sedimentibacter sp002316225 (GB_GCA_002316225.1)
Sedimentibacter sp002316585 (GB_GCA_002316585.1)
Sedimentibacter sp002385835 (GB_GCA_002385835.1)
Sedimentibacter sp002422545 (GB_GCA_002422545.1)
UBA6132 sp002435835 (GB_GCA_002435835.1)
UBA1535 sp002396645 (GB_GCA_002396645.1)
UBA1535 sp002323775 (GB_GCA_002323775.1)
UBA1535 sp002332905 (GB_GCA_002332905.1)
UBA1535 sp002479485 (GB_GCA_002479485.1)
Sedimentibacteraceae (metabat.350)
38−11 sp003446975 (GB_GCA_003446975.1)
38−11 sp001509345 (GB_GCA_001509345.1)
Dethiosulfatibacter aminovorans (RS_GCF_900142005.1)
UBA8670 sp003539375 (GB_GCA_003539375.1)

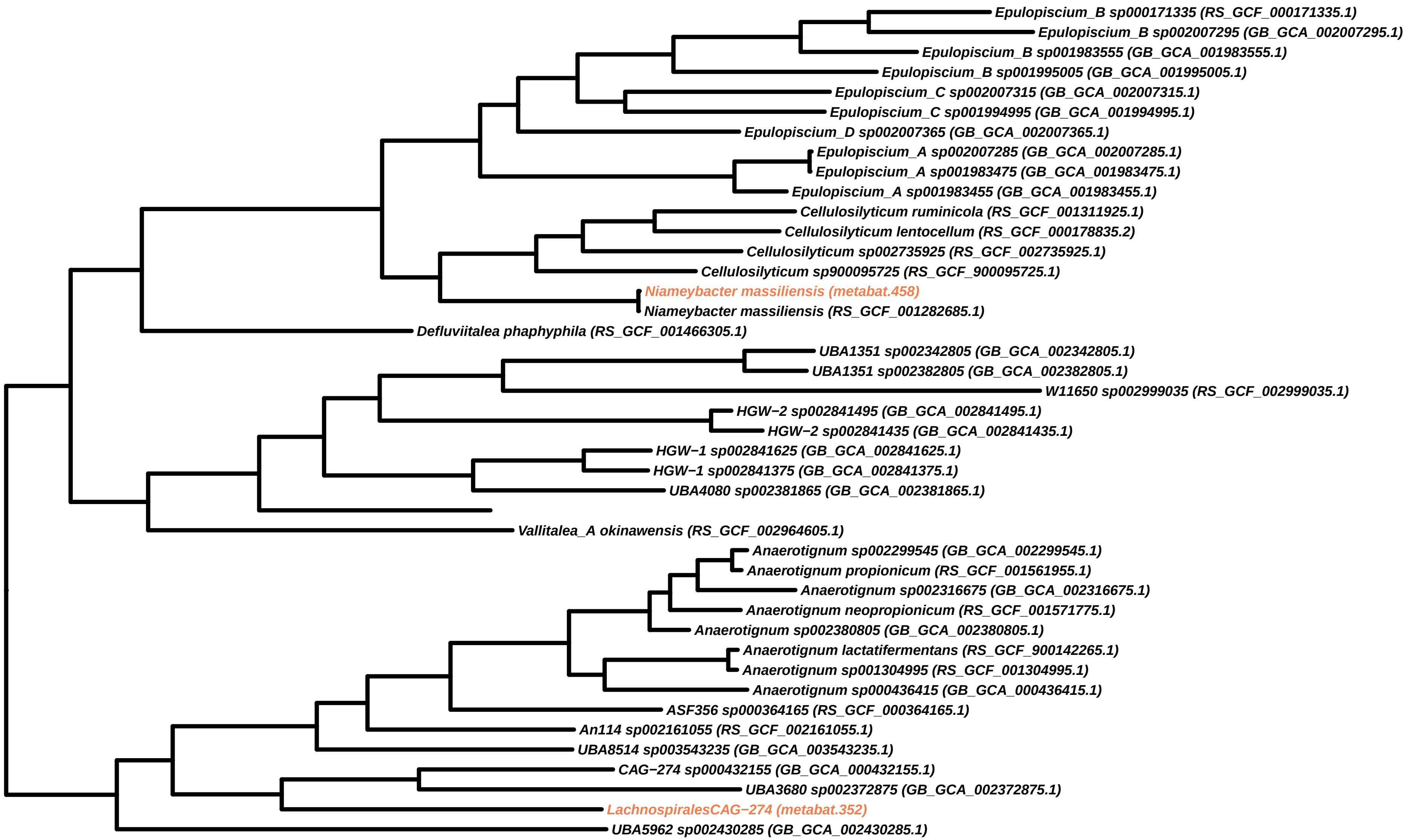
Epulopiscium_B sp000171335 (RS_GCF_000171335.1)
Epulopiscium_B sp002007295 (GB_GCA_002007295.1)
Epulopiscium_B sp001983555 (GB_GCA_001983555.1)
Epulopiscium_B sp001995005 (GB_GCA_001995005.1)
Epulopiscium_C sp002007315 (GB_GCA_002007315.1)
Epulopiscium_C sp001994995 (GB_GCA_001994995.1)
Epulopiscium_D sp002007365 (GB_GCA_002007365.1)
Epulopiscium_A sp002007285 (GB_GCA_002007285.1)
Epulopiscium_A sp001983475 (GB_GCA_001983475.1)
Epulopiscium_A sp001983455 (GB_GCA_001983455.1)
Cellulosilyticum ruminicola (RS_GCF_001311925.1)
Cellulosilyticum lentocellum (RS_GCF_000178835.2)
Cellulosilyticum sp002735925 (RS_GCF_002735925.1)
Cellulosilyticum sp900095725 (RS_GCF_900095725.1)
Niameybacter massiliensis (metabat.458)
Niameybacter massiliensis (RS_GCF_001282685.1)
Defluviitalea phaphyphila (RS_GCF_001466305.1)
UBA1351 sp002342805 (GB_GCA_002342805.1)
UBA1351 sp002382805 (GB_GCA_002382805.1)
W11650 sp002999035 (RS_GCF_002999035.1)
HGW−2 sp002841495 (GB_GCA_002841495.1)
HGW−2 sp002841435 (GB_GCA_002841435.1)
HGW−1 sp002841625 (GB_GCA_002841625.1)
HGW−1 sp002841375 (GB_GCA_002841375.1)
UBA4080 sp002381865 (GB_GCA_002381865.1)
Vallitalea_A okinawensis (RS_GCF_002964605.1)
Anaerotignum sp002299545 (GB_GCA_002299545.1)
Anaerotignum propionicum (RS_GCF_001561955.1)
Anaerotignum sp002316675 (GB_GCA_002316675.1)
Anaerotignum neopropionicum (RS_GCF_001571775.1)
Anaerotignum sp002380805 (GB_GCA_002380805.1)
Anaerotignum lactatifermentans (RS_GCF_900142265.1)
Anaerotignum sp001304995 (RS_GCF_001304995.1)
Anaerotignum sp000436415 (GB_GCA_000436415.1)
ASF356 sp000364165 (RS_GCF_000364165.1)
An114 sp002161055 (RS_GCF_002161055.1)
UBA8514 sp003543235 (GB_GCA_003543235.1)
CAG−274 sp000432155 (GB_GCA_000432155.1)
UBA3680 sp002372875 (GB_GCA_002372875.1)
LachnospiralesCAG−274 (metabat.352)
UBA5962 sp002430285 (GB_GCA_002430285.1)

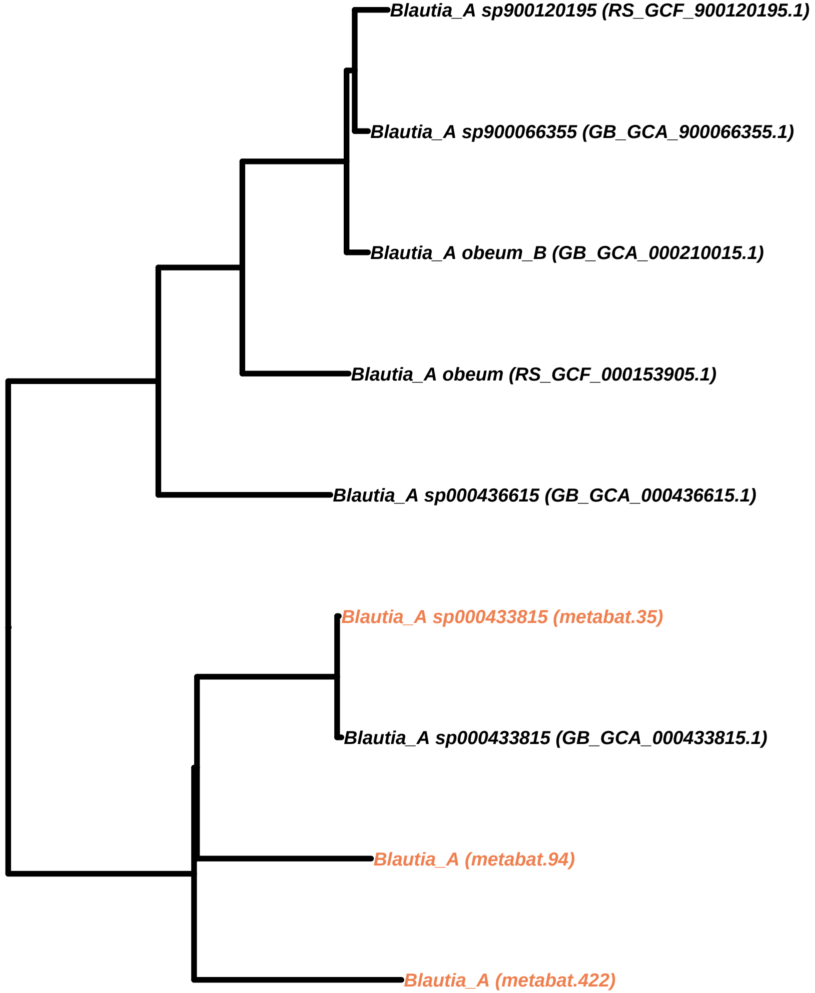
Blautia_A sp900120195 (RS_GCF_900120195.1)
Blautia_A sp900066355 (GB_GCA_900066355.1)
Blautia_A obeum_B (GB_GCA_000210015.1)
Blautia_A obeum (RS_GCF_000153905.1)
Blautia_A sp000436615 (GB_GCA_000436615.1)
Blautia_A sp000433815 (metabat.35)
Blautia_A sp000433815 (GB_GCA_000433815.1)
Blautia_A (metabat.94)
Blautia_A (metabat.422)

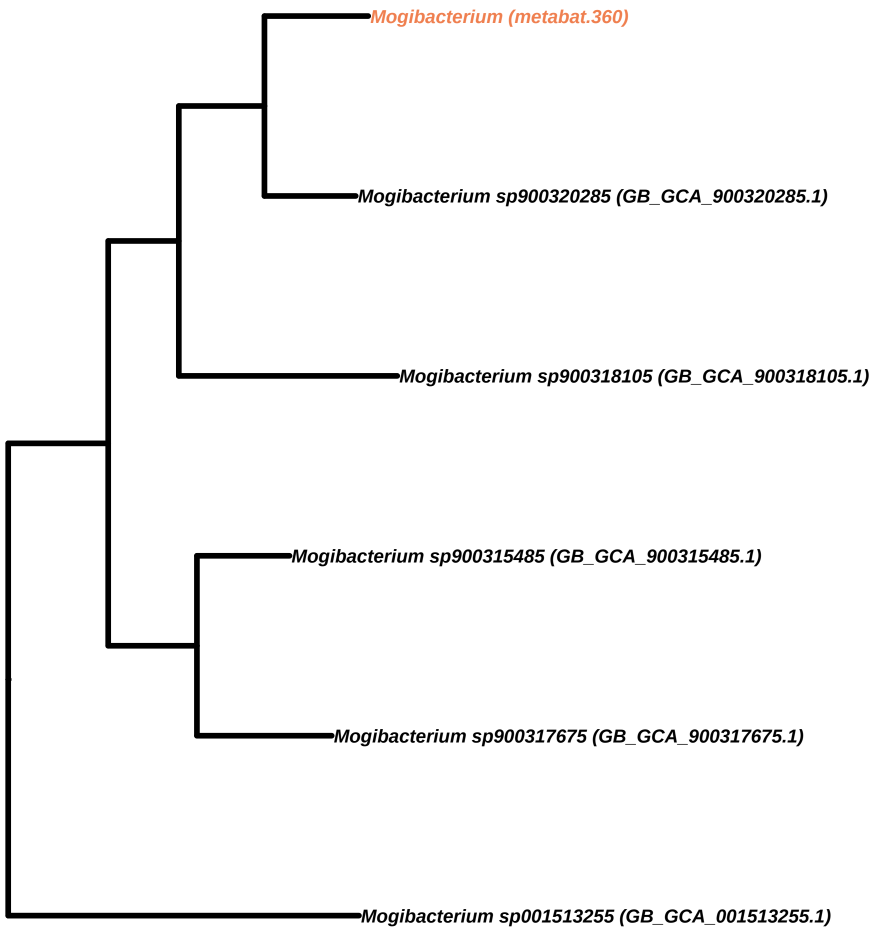
Mogibacterium (metabat.360)
Mogibacterium sp900320285 (GB_GCA_900320285.1)
Mogibacterium sp900318105 (GB_GCA_900318105.1)
Mogibacterium sp900315485 (GB_GCA_900315485.1)
Mogibacterium sp900317675 (GB_GCA_900317675.1)
Mogibacterium sp001513255 (GB_GCA_001513255.1)

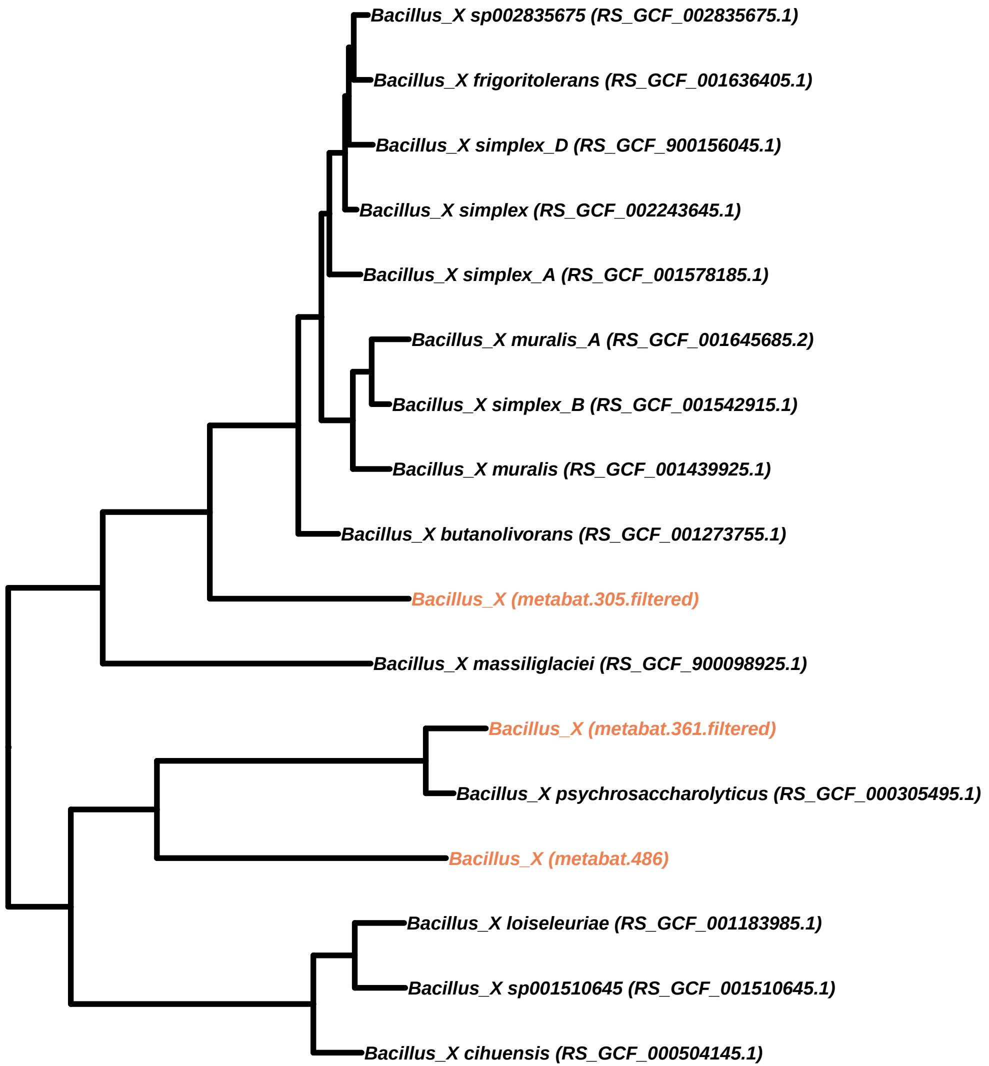
Bacillus_X sp002835675 (RS_GCF_002835675.1)
Bacillus_X frigoritolerans (RS_GCF_001636405.1)
Bacillus_X simplex_D (RS_GCF_900156045.1)
Bacillus_X simplex (RS_GCF_002243645.1)
Bacillus_X simplex_A (RS_GCF_001578185.1)
Bacillus_X muralis_A (RS_GCF_001645685.2)
Bacillus_X simplex_B (RS_GCF_001542915.1)
Bacillus_X muralis (RS_GCF_001439925.1)
Bacillus_X butanolivorans (RS_GCF_001273755.1)
Bacillus_X (metabat.305.filtered)
Bacillus_X massiliglaciei (RS_GCF_900098925.1)
Bacillus_X (metabat.361.filtered)
Bacillus_X psychrosaccharolyticus (RS_GCF_000305495.1)
Bacillus_X (metabat.486)
Bacillus_X loiseleuriae (RS_GCF_001183985.1)
Bacillus_X sp001510645 (RS_GCF_001510645.1)
Bacillus_X cihuensis (RS_GCF_000504145.1)

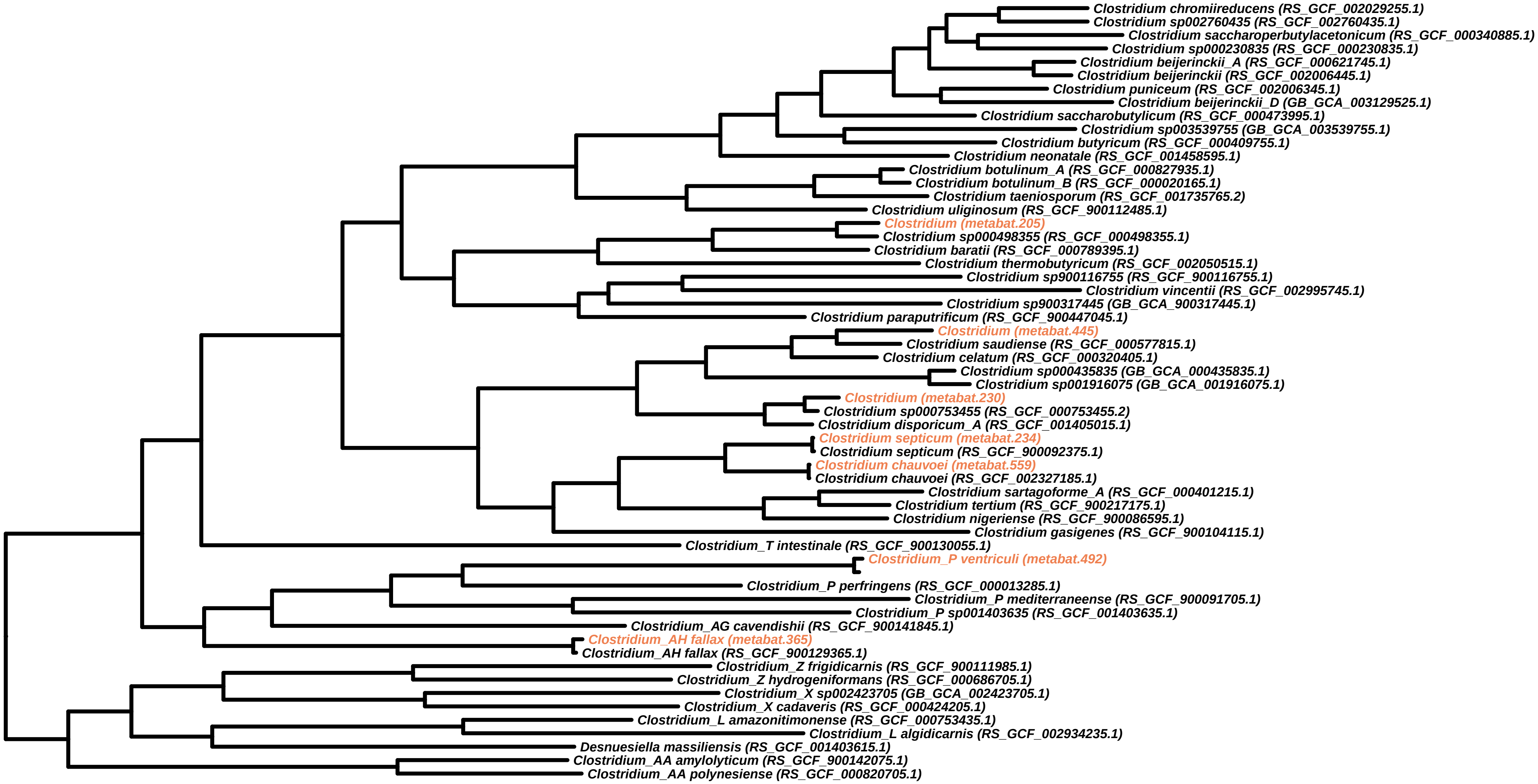
Clostridium chromiireducens (RS_GCF_002029255.1)
Clostridium sp002760435 (RS_GCF_002760435.1)
Clostridium saccharoperbutylacetonicum (RS_GCF_000340885.1)
Clostridium sp000230835 (RS_GCF_000230835.1)
Clostridium beijerinckii_A (RS_GCF_000621745.1)
Clostridium beijerinckii (RS_GCF_002006445.1)
Clostridium puniceum (RS_GCF_002006345.1)
Clostridium beijerinckii_D (GB_GCA_003129525.1)
Clostridium saccharobutylicum (RS_GCF_000473995.1)
Clostridium sp003539755 (GB_GCA_003539755.1)
Clostridium butyricum (RS_GCF_000409755.1)
Clostridium neonatale (RS_GCF_001458595.1)
Clostridium botulinum_A (RS_GCF_000827935.1)
Clostridium botulinum_B (RS_GCF_000020165.1)
Clostridium taeniosporum (RS_GCF_001735765.2)
Clostridium uliginosum (RS_GCF_900112485.1)
Clostridium (metabat.205)
Clostridium sp000498355 (RS_GCF_000498355.1)
Clostridium baratii (RS_GCF_000789395.1)
Clostridium thermobutyricum (RS_GCF_002050515.1)
Clostridium sp900116755 (RS_GCF_900116755.1)
Clostridium vincentii (RS_GCF_002995745.1)
Clostridium sp900317445 (GB_GCA_900317445.1)
Clostridium paraputrificum (RS_GCF_900447045.1)
Clostridium (metabat.445)
Clostridium saudiense (RS_GCF_000577815.1)
Clostridium celatum (RS_GCF_000320405.1)
Clostridium sp000435835 (GB_GCA_000435835.1)
Clostridium sp001916075 (GB_GCA_001916075.1)
Clostridium (metabat.230)
Clostridium sp000753455 (RS_GCF_000753455.2)
Clostridium disporicum_A (RS_GCF_001405015.1)
Clostridium septicum (metabat.234)
Clostridium septicum (RS_GCF_900092375.1)
Clostridium chauvoei (metabat.559)
Clostridium chauvoei (RS_GCF_002327185.1)
Clostridium sartagoforme_A (RS_GCF_000401215.1)
Clostridium tertium (RS_GCF_900217175.1)
Clostridium nigeriense (RS_GCF_900086595.1)
Clostridium gasigenes (RS_GCF_900104115.1)
Clostridium_T intestinale (RS_GCF_900130055.1)
Clostridium_P ventriculi (metabat.492)
Clostridium_P perfringens (RS_GCF_000013285.1)
Clostridium_P mediterraneense (RS_GCF_900091705.1)
Clostridium_P sp001403635 (RS_GCF_001403635.1)
Clostridium_AG cavendishii (RS_GCF_900141845.1)
Clostridium_AH fallax (metabat.365)
Clostridium_AH fallax (RS_GCF_900129365.1)
Clostridium_Z frigidicarnis (RS_GCF_900111985.1)
Clostridium_Z hydrogeniformans (RS_GCF_000686705.1)
Clostridium_X sp002423705 (GB_GCA_002423705.1)
Clostridium_X cadaveris (RS_GCF_000424205.1)
Clostridium_L amazonitimonense (RS_GCF_000753435.1)
Clostridium_L algidicarnis (RS_GCF_002934235.1)
Desnuesiella massiliensis (RS_GCF_001403615.1)
Clostridium_AA amylolyticum (RS_GCF_900142075.1)
Clostridium_AA polynesiense (RS_GCF_000820705.1)

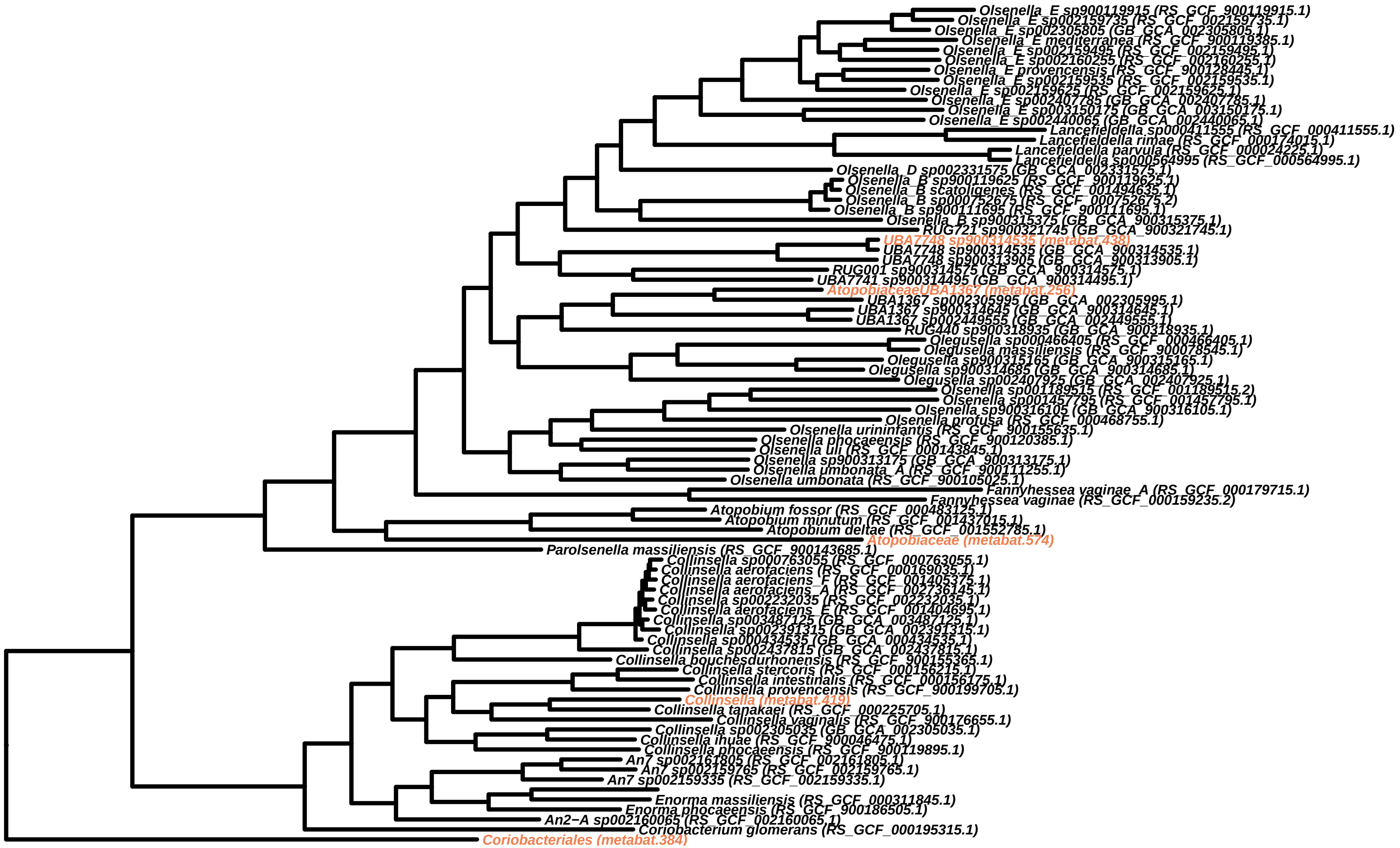
Olsenella_E sp900119915 (RS_GCF_900119915.1)
Olsenella_E sp002159735 (RS_GCF_002159735.1)
Olsenella_E sp002305805 (GB_GCA_002305805.1)
Olsenella_E mediterranea (RS_GCF_900119385.1)
Olsenella_E sp002159495 (RS_GCF_002159495.1)
Olsenella_E sp002160255 (RS_GCF_002160255.1)
Olsenella_E provencensis (RS_GCF_900128445.1)
Olsenella_E sp002159535 (RS_GCF_002159535.1)
Olsenella_E sp002159625 (RS_GCF_002159625.1)
Olsenella_E sp002407785 (GB_GCA_002407785.1)
Olsenella_E sp003150175 (GB_GCA_003150175.1)
Olsenella_E sp002440065 (GB_GCA_002440065.1)
Lancefieldella sp000411555 (RS_GCF_000411555.1)
Lancefieldella rimae (RS_GCF_000174015.1)
Lancefieldella parvula (RS_GCF_000024225.1)
Lancefieldella sp000564995 (RS_GCF_000564995.1)
Olsenella_D sp002331575 (GB_GCA_002331575.1)
Olsenella_B sp900119625 (RS_GCF_900119625.1)
Olsenella_B scatoligenes (RS_GCF_001494635.1)
Olsenella_B sp000752675 (RS_GCF_000752675.2)
Olsenella_B sp900111695 (RS_GCF_900111695.1)
Olsenella_B sp900315375 (GB_GCA_900315375.1)
RUG721 sp900321745 (GB_GCA_900321745.1)
UBA7748 sp900314535 (metabat.438)
UBA7748 sp900314535 (GB_GCA_900314535.1)
UBA7748 sp900313905 (GB_GCA_900313905.1)
RUG001 sp900314575 (GB_GCA_900314575.1)
UBA7741 sp900314495 (GB_GCA_900314495.1)
AtopobiaceaeUBA1367 (metabat.256)
UBA1367 sp002305995 (GB_GCA_002305995.1)
UBA1367 sp900314645 (GB_GCA_900314645.1)
UBA1367 sp002449555 (GB_GCA_002449555.1)
RUG440 sp900318935 (GB_GCA_900318935.1)
Olegusella sp000466405 (RS_GCF_000466405.1)
Olegusella massiliensis (RS_GCF_900078545.1)
Olegusella sp900315165 (GB_GCA_900315165.1)
Olegusella sp900314685 (GB_GCA_900314685.1)
Olegusella sp002407925 (GB_GCA_002407925.1)
Olsenella sp001189515 (RS_GCF_001189515.2)
Olsenella sp001457795 (RS_GCF_001457795.1)
Olsenella sp900316105 (GB_GCA_900316105.1)
Olsenella profusa (RS_GCF_000468755.1)
Olsenella urininfantis (RS_GCF_900155635.1)
Olsenella phocaeensis (RS_GCF_900120385.1)
Olsenella uli (RS_GCF_000143845.1)
Olsenella sp900313175 (GB_GCA_900313175.1)
Olsenella umbonata_A (RS_GCF_900111255.1)
Olsenella umbonata (RS_GCF_900105025.1)
Fannyhessea vaginae_A (RS_GCF_000179715.1)
Fannyhessea vaginae (RS_GCF_000159235.2)
Atopobium fossor (RS_GCF_000483125.1)
Atopobium minutum (RS_GCF_001437015.1)
Atopobium deltae (RS_GCF_001552785.1)
Atopobiaceae (metabat.574)
Parolsenella massiliensis (RS_GCF_900143685.1)
Collinsella sp000763055 (RS_GCF_000763055.1)
Collinsella aerofaciens (RS_GCF_000169035.1)
Collinsella aerofaciens_F (RS_GCF_001405375.1)
Collinsella aerofaciens_A (RS_GCF_002736145.1)
Collinsella sp002232035 (RS_GCF_002232035.1)
Collinsella aerofaciens_E (RS_GCF_001404695.1)
Collinsella sp003487125 (GB_GCA_003487125.1)
Collinsella sp002391315 (GB_GCA_002391315.1)
Collinsella sp000434535 (GB_GCA_000434535.1)
Collinsella sp002437815 (GB_GCA_002437815.1)
Collinsella bouchesdurhonensis (RS_GCF_900155365.1)
Collinsella stercoris (RS_GCF_000156215.1)
Collinsella intestinalis (RS_GCF_000156175.1)
Collinsella provencensis (RS_GCF_900199705.1)
Collinsella (metabat.419)
Collinsella tanakaei (RS_GCF_000225705.1)
Collinsella vaginalis (RS_GCF_900176655.1)
Collinsella sp002305035 (GB_GCA_002305035.1)
Collinsella ihuae (RS_GCF_900046475.1)
Collinsella phocaeensis (RS_GCF_900119895.1)
An7 sp002161805 (RS_GCF_002161805.1)
An7 sp002159765 (RS_GCF_002159765.1)
An7 sp002159335 (RS_GCF_002159335.1)
Enorma massiliensis (RS_GCF_000311845.1)
Enorma phocaeensis (RS_GCF_900186505.1)
An2−A sp002160065 (RS_GCF_002160065.1)
Coriobacterium glomerans (RS_GCF_000195315.1)
Coriobacteriales (metabat.384)

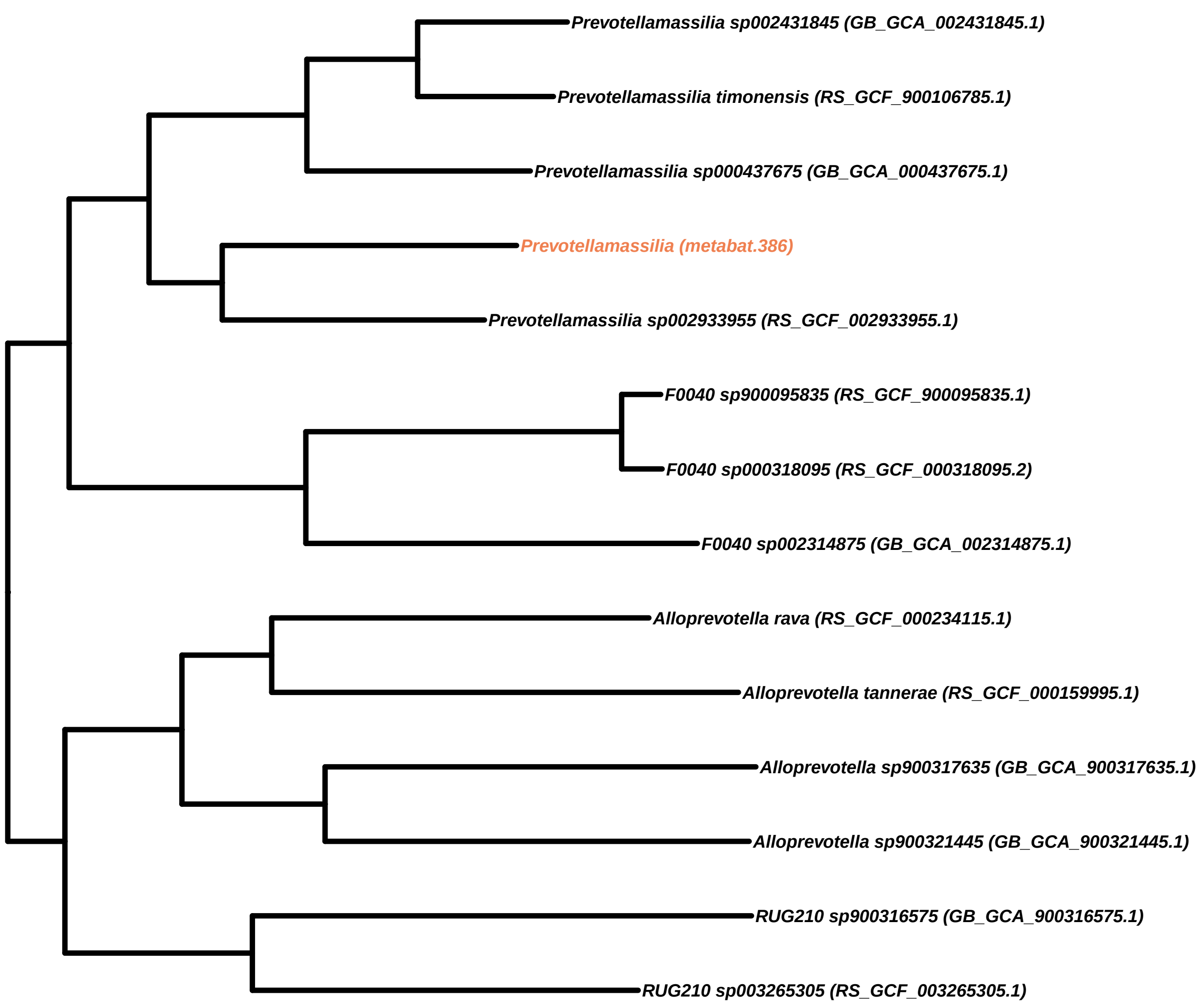

Prevotellamassilia sp002431845 (GB_GCA_002431845.1)
Prevotellamassilia timonensis (RS_GCF_900106785.1)
Prevotellamassilia sp000437675 (GB_GCA_000437675.1)
Prevotellamassilia (metabat.386)
Prevotellamassilia sp002933955 (RS_GCF_002933955.1)
F0040 sp900095835 (RS_GCF_900095835.1)
F0040 sp000318095 (RS_GCF_000318095.2)
F0040 sp002314875 (GB_GCA_002314875.1)
Alloprevotella rava (RS_GCF_000234115.1)
Alloprevotella tannerae (RS_GCF_000159995.1)
Alloprevotella sp900317635 (GB_GCA_900317635.1)
Alloprevotella sp900321445 (GB_GCA_900321445.1)
RUG210 sp900316575 (GB_GCA_900316575.1)
RUG210 sp003265305 (RS_GCF_003265305.1)

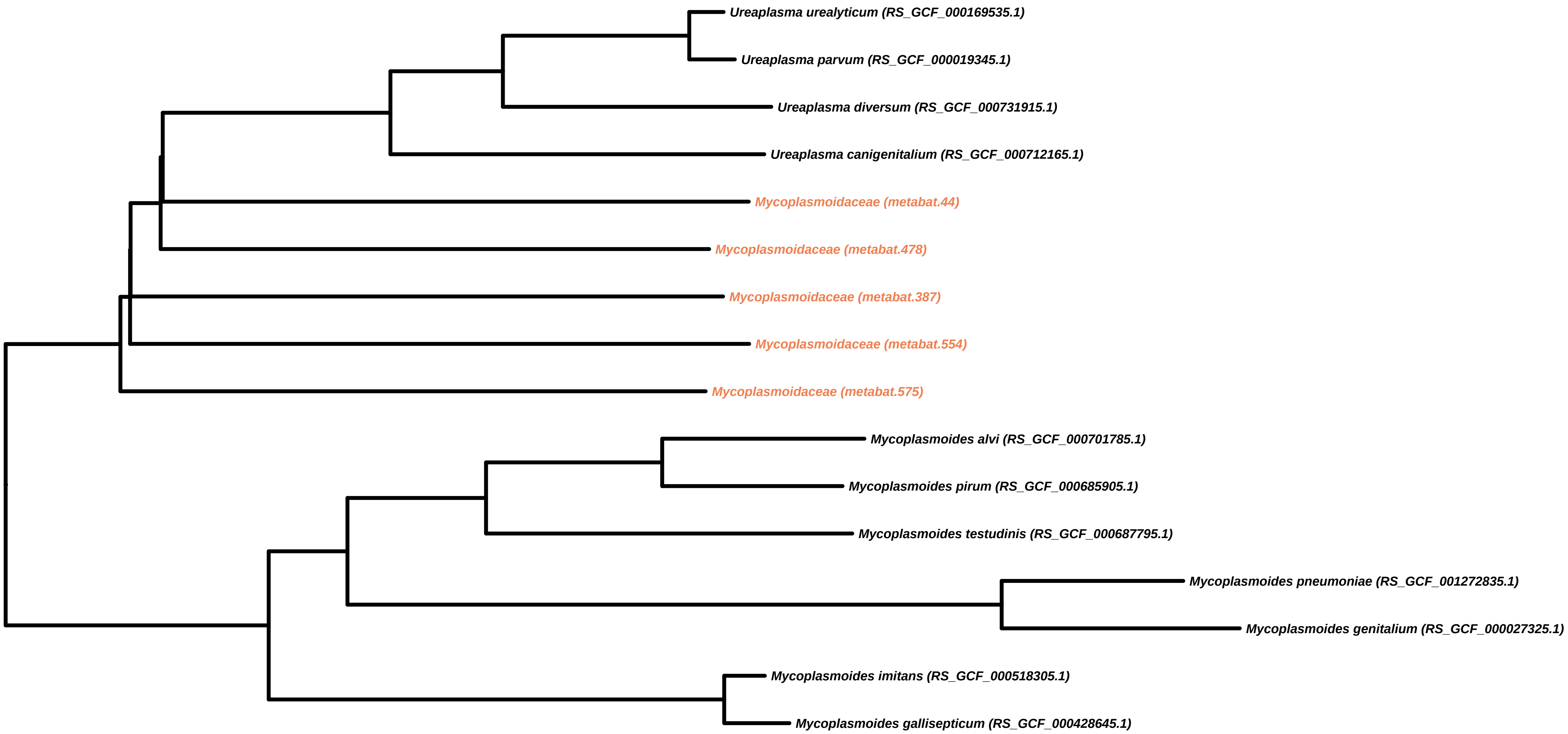

Ureaplasma urealyticum (RS_GCF_000169535.1)
Ureaplasma parvum (RS_GCF_000019345.1)
Ureaplasma diversum (RS_GCF_000731915.1)
Ureaplasma canigenitalium (RS_GCF_000712165.1)
Mycoplasmoidaceae (metabat.44)
Mycoplasmoidaceae (metabat.478)
Mycoplasmoidaceae (metabat.387)
Mycoplasmoidaceae (metabat.554)
Mycoplasmoidaceae (metabat.575)
Mycoplasmoides alvi (RS_GCF_000701785.1)
Mycoplasmoides pirum (RS_GCF_000685905.1)
Mycoplasmoides testudinis (RS_GCF_000687795.1)
Mycoplasmoides pneumoniae (RS_GCF_001272835.1)
Mycoplasmoides genitalium (RS_GCF_000027325.1)
Mycoplasmoides imitans (RS_GCF_000518305.1)
Mycoplasmoides gallisepticum (RS_GCF_000428645.1)

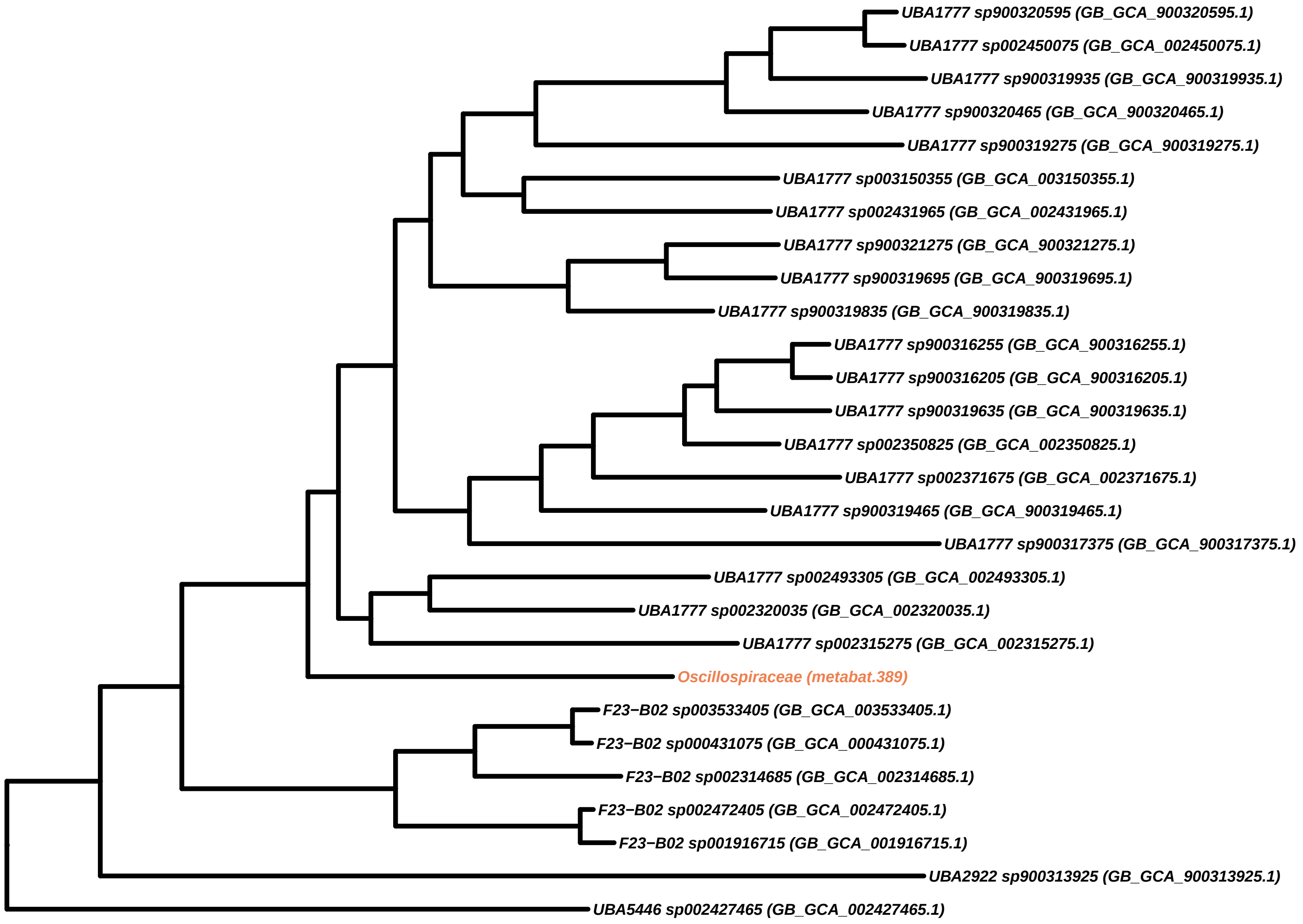
UBA1777 sp900320595 (GB_GCA_900320595.1)
UBA1777 sp002450075 (GB_GCA_002450075.1)
UBA1777 sp900319935 (GB_GCA_900319935.1)
UBA1777 sp900320465 (GB_GCA_900320465.1)
UBA1777 sp900319275 (GB_GCA_900319275.1)
UBA1777 sp003150355 (GB_GCA_003150355.1)
UBA1777 sp002431965 (GB_GCA_002431965.1)
UBA1777 sp900321275 (GB_GCA_900321275.1)
UBA1777 sp900319695 (GB_GCA_900319695.1)
UBA1777 sp900319835 (GB_GCA_900319835.1)
UBA1777 sp900316255 (GB_GCA_900316255.1)
UBA1777 sp900316205 (GB_GCA_900316205.1)
UBA1777 sp900319635 (GB_GCA_900319635.1)
UBA1777 sp002350825 (GB_GCA_002350825.1)
UBA1777 sp002371675 (GB_GCA_002371675.1)
UBA1777 sp900319465 (GB_GCA_900319465.1)
UBA1777 sp900317375 (GB_GCA_900317375.1)
UBA1777 sp002493305 (GB_GCA_002493305.1)
UBA1777 sp002320035 (GB_GCA_002320035.1)
UBA1777 sp002315275 (GB_GCA_002315275.1)
Oscillospiraceae (metabat.389)
F23−B02 sp003533405 (GB_GCA_003533405.1)
F23−B02 sp000431075 (GB_GCA_000431075.1)
F23−B02 sp002314685 (GB_GCA_002314685.1)
F23−B02 sp002472405 (GB_GCA_002472405.1)
F23−B02 sp001916715 (GB_GCA_001916715.1)
UBA2922 sp900313925 (GB_GCA_900313925.1)
UBA5446 sp002427465 (GB_GCA_002427465.1)

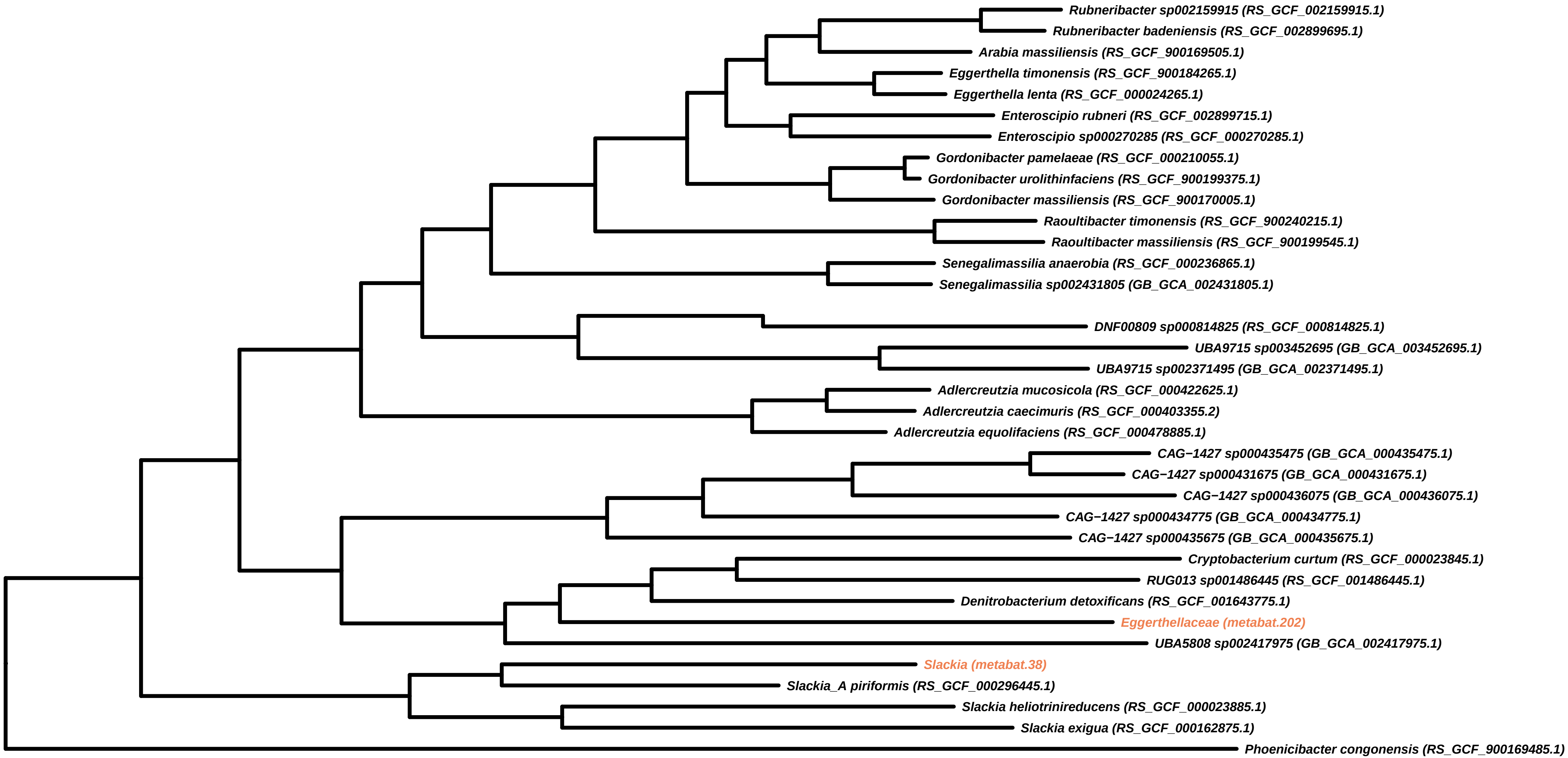
Rubneribacter sp002159915 (RS_GCF_002159915.1)
Rubneribacter badeniensis (RS_GCF_002899695.1)
Arabia massiliensis (RS_GCF_900169505.1)
Eggerthella timonensis (RS_GCF_900184265.1)
Eggerthella lenta (RS_GCF_000024265.1)
Enteroscipio rubneri (RS_GCF_002899715.1)
Enteroscipio sp000270285 (RS_GCF_000270285.1)
Gordonibacter pamelaeae (RS_GCF_000210055.1)
Gordonibacter urolithinfaciens (RS_GCF_900199375.1)
Gordonibacter massiliensis (RS_GCF_900170005.1)
Raoultibacter timonensis (RS_GCF_900240215.1)
Raoultibacter massiliensis (RS_GCF_900199545.1)
Senegalimassilia anaerobia (RS_GCF_000236865.1)
Senegalimassilia sp002431805 (GB_GCA_002431805.1)
DNF00809 sp000814825 (RS_GCF_000814825.1)
UBA9715 sp003452695 (GB_GCA_003452695.1)
UBA9715 sp002371495 (GB_GCA_002371495.1)
Adlercreutzia mucosicola (RS_GCF_000422625.1)
Adlercreutzia caecimuris (RS_GCF_000403355.2)
Adlercreutzia equolifaciens (RS_GCF_000478885.1)
CAG−1427 sp000435475 (GB_GCA_000435475.1)
CAG−1427 sp000431675 (GB_GCA_000431675.1)
CAG−1427 sp000436075 (GB_GCA_000436075.1)
CAG−1427 sp000434775 (GB_GCA_000434775.1)
CAG−1427 sp000435675 (GB_GCA_000435675.1)
Cryptobacterium curtum (RS_GCF_000023845.1)
RUG013 sp001486445 (RS_GCF_001486445.1)
Denitrobacterium detoxificans (RS_GCF_001643775.1)
Eggerthellaceae (metabat.202)
UBA5808 sp002417975 (GB_GCA_002417975.1)
Slackia (metabat.38)
Slackia_A piriformis (RS_GCF_000296445.1)
Slackia heliotrinireducens (RS_GCF_000023885.1)
Slackia exigua (RS_GCF_000162875.1)
Phoenicibacter congonensis (RS_GCF_900169485.1)

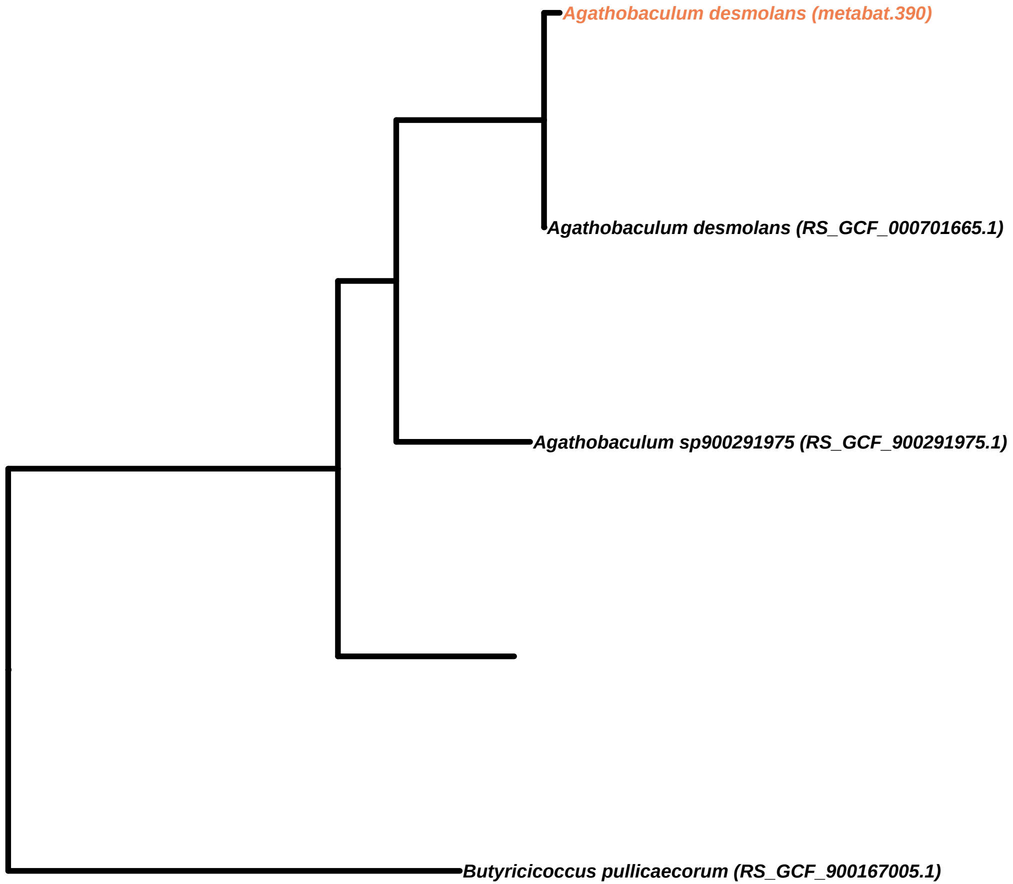

Agathobaculum desmolans (metabat.390)
Agathobaculum desmolans (RS_GCF_000701665.1)
Agathobaculum sp900291975 (RS_GCF_900291975.1)
Butyricicoccus pullicaecorum (RS_GCF_900167005.1)

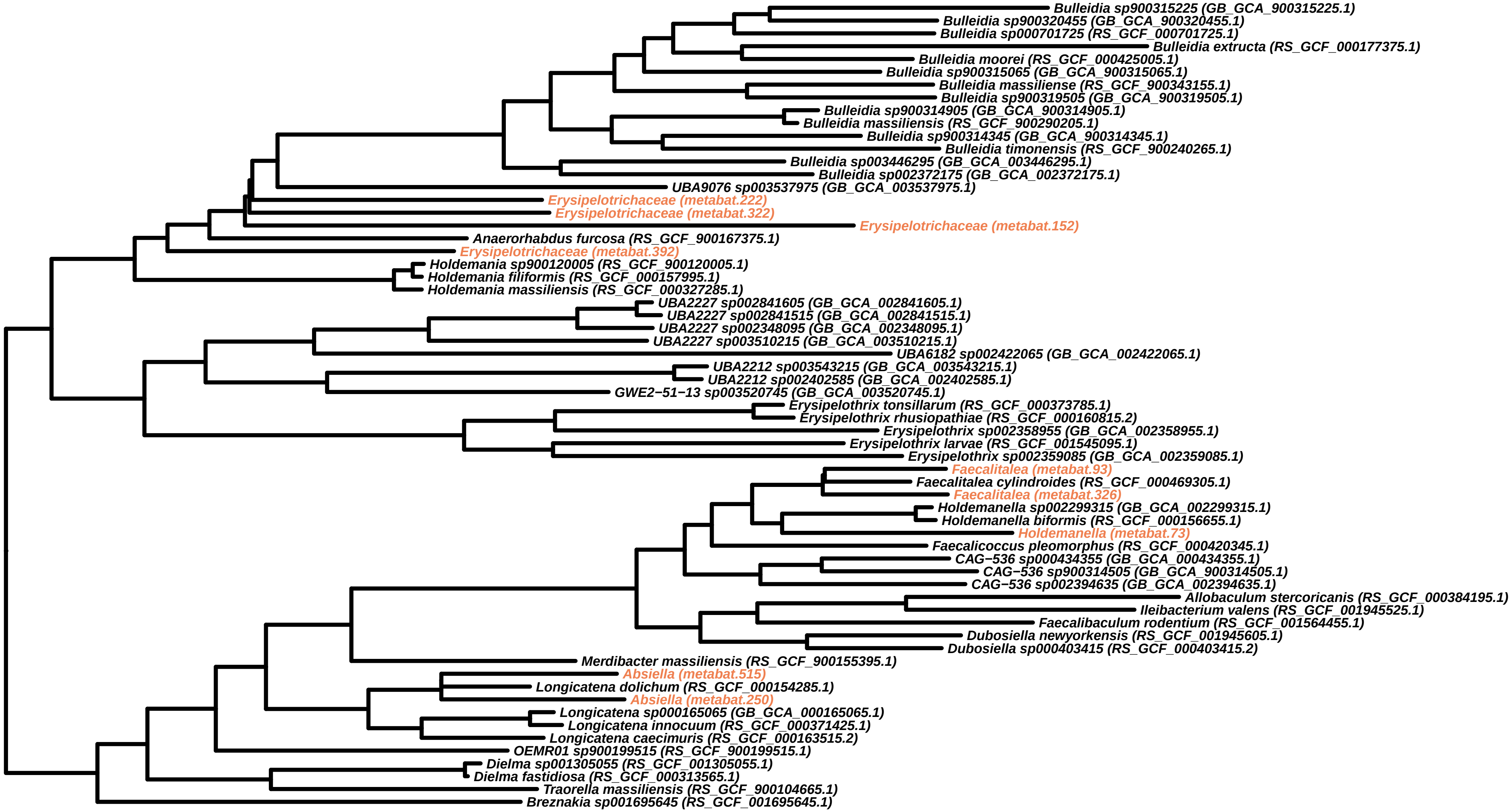
Bulleidia sp900315225 (GB_GCA_900315225.1)
Bulleidia sp900320455 (GB_GCA_900320455.1)
Bulleidia sp000701725 (RS_GCF_000701725.1)
Bulleidia extructa (RS_GCF_000177375.1)
Bulleidia moorei (RS_GCF_000425005.1)
Bulleidia sp900315065 (GB_GCA_900315065.1)
Bulleidia massiliense (RS_GCF_900343155.1)
Bulleidia sp900319505 (GB_GCA_900319505.1)
Bulleidia sp900314905 (GB_GCA_900314905.1)
Bulleidia massiliensis (RS_GCF_900290205.1)
Bulleidia sp900314345 (GB_GCA_900314345.1)
Bulleidia timonensis (RS_GCF_900240265.1)
Bulleidia sp003446295 (GB_GCA_003446295.1)
Bulleidia sp002372175 (GB_GCA_002372175.1)
UBA9076 sp003537975 (GB_GCA_003537975.1)
Erysipelotrichaceae (metabat.222)
Erysipelotrichaceae (metabat.322)
Erysipelotrichaceae (metabat.152)
Anaerorhabdus furcosa (RS_GCF_900167375.1)
Erysipelotrichaceae (metabat.392)
Holdemania sp900120005 (RS_GCF_900120005.1)
Holdemania filiformis (RS_GCF_000157995.1)
Holdemania massiliensis (RS_GCF_000327285.1)
UBA2227 sp002841605 (GB_GCA_002841605.1)
UBA2227 sp002841515 (GB_GCA_002841515.1)
UBA2227 sp002348095 (GB_GCA_002348095.1)
UBA2227 sp003510215 (GB_GCA_003510215.1)
UBA6182 sp002422065 (GB_GCA_002422065.1)
UBA2212 sp003543215 (GB_GCA_003543215.1)
UBA2212 sp002402585 (GB_GCA_002402585.1)
GWE2−51−13 sp003520745 (GB_GCA_003520745.1)
Erysipelothrix tonsillarum (RS_GCF_000373785.1)
Erysipelothrix rhusiopathiae (RS_GCF_000160815.2)
Erysipelothrix sp002358955 (GB_GCA_002358955.1)
Erysipelothrix larvae (RS_GCF_001545095.1)
Erysipelothrix sp002359085 (GB_GCA_002359085.1)
Faecalitalea (metabat.93)
Faecalitalea cylindroides (RS_GCF_000469305.1)
Faecalitalea (metabat.326)
Holdemanella sp002299315 (GB_GCA_002299315.1)
Holdemanella biformis (RS_GCF_000156655.1)
Holdemanella (metabat.73)
Faecalicoccus pleomorphus (RS_GCF_000420345.1)
CAG−536 sp000434355 (GB_GCA_000434355.1)
CAG−536 sp900314505 (GB_GCA_900314505.1)
CAG−536 sp002394635 (GB_GCA_002394635.1)
Allobaculum stercoricanis (RS_GCF_000384195.1)
Ileibacterium valens (RS_GCF_001945525.1)
Faecalibaculum rodentium (RS_GCF_001564455.1)
Dubosiella newyorkensis (RS_GCF_001945605.1)
Dubosiella sp000403415 (RS_GCF_000403415.2)
Merdibacter massiliensis (RS_GCF_900155395.1)
Absiella (metabat.515)
Longicatena dolichum (RS_GCF_000154285.1)
Absiella (metabat.250)
Longicatena sp000165065 (GB_GCA_000165065.1)
Longicatena innocuum (RS_GCF_000371425.1)
Longicatena caecimuris (RS_GCF_000163515.2)
OEMR01 sp900199515 (RS_GCF_900199515.1)
Dielma sp001305055 (RS_GCF_001305055.1)
Dielma fastidiosa (RS_GCF_000313565.1)
Traorella massiliensis (RS_GCF_900104665.1)
Breznakia sp001695645 (RS_GCF_001695645.1)

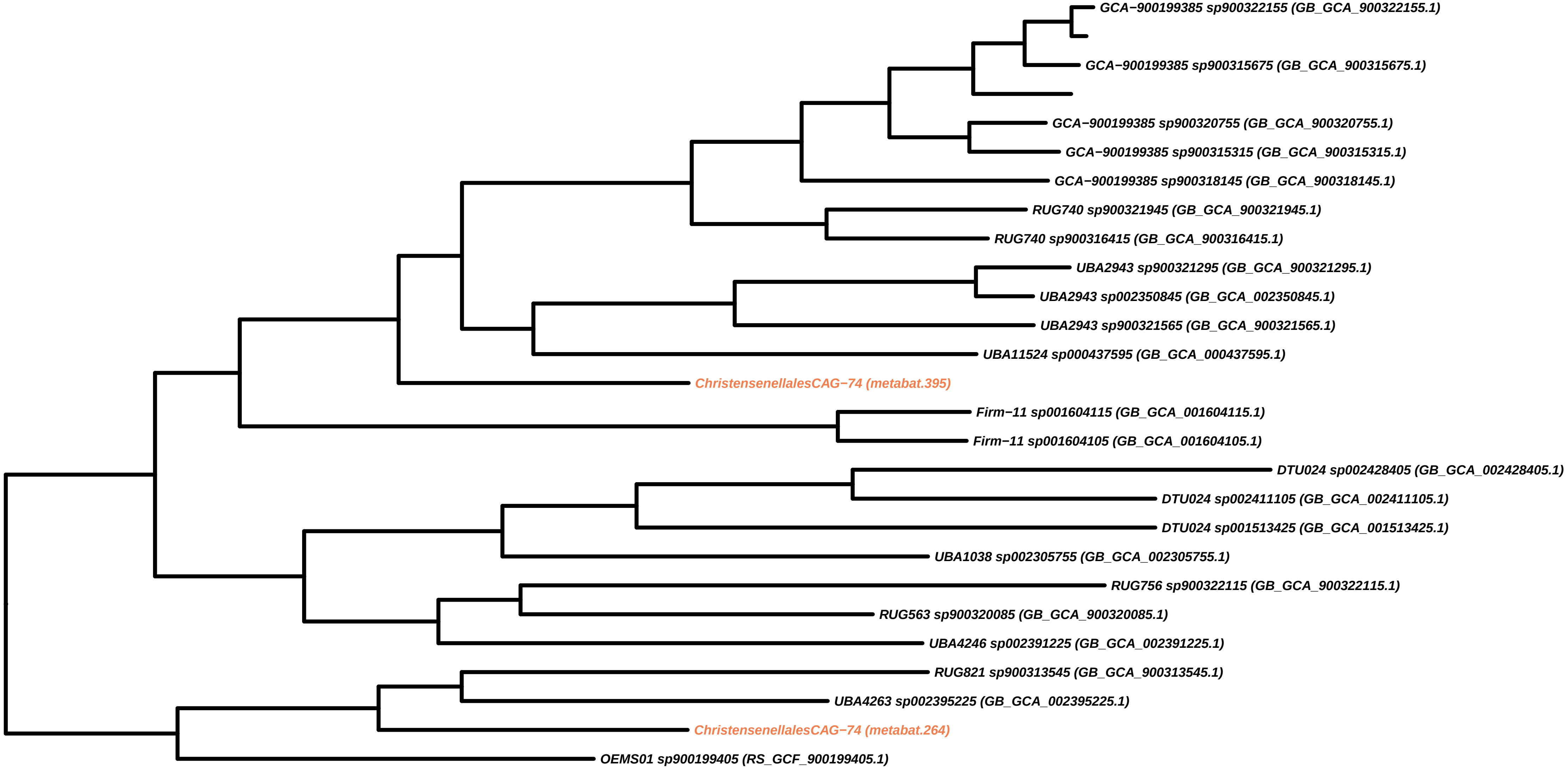
GCA−900199385 sp900322155 (GB_GCA_900322155.1)
GCA−900199385 sp900315675 (GB_GCA_900315675.1)
GCA−900199385 sp900320755 (GB_GCA_900320755.1)
GCA−900199385 sp900315315 (GB_GCA_900315315.1)
GCA−900199385 sp900318145 (GB_GCA_900318145.1)
RUG740 sp900321945 (GB_GCA_900321945.1)
RUG740 sp900316415 (GB_GCA_900316415.1)
UBA2943 sp900321295 (GB_GCA_900321295.1)
UBA2943 sp002350845 (GB_GCA_002350845.1)
UBA2943 sp900321565 (GB_GCA_900321565.1)
UBA11524 sp000437595 (GB_GCA_000437595.1)
ChristensenellalesCAG−74 (metabat.395)
Firm−11 sp001604115 (GB_GCA_001604115.1)
Firm−11 sp001604105 (GB_GCA_001604105.1)
DTU024 sp002428405 (GB_GCA_002428405.1)
DTU024 sp002411105 (GB_GCA_002411105.1)
DTU024 sp001513425 (GB_GCA_001513425.1)
UBA1038 sp002305755 (GB_GCA_002305755.1)
RUG756 sp900322115 (GB_GCA_900322115.1)
RUG563 sp900320085 (GB_GCA_900320085.1)
UBA4246 sp002391225 (GB_GCA_002391225.1)
RUG821 sp900313545 (GB_GCA_900313545.1)
UBA4263 sp002395225 (GB_GCA_002395225.1)
ChristensenellalesCAG−74 (metabat.264)
OEMS01 sp900199405 (RS_GCF_900199405.1)

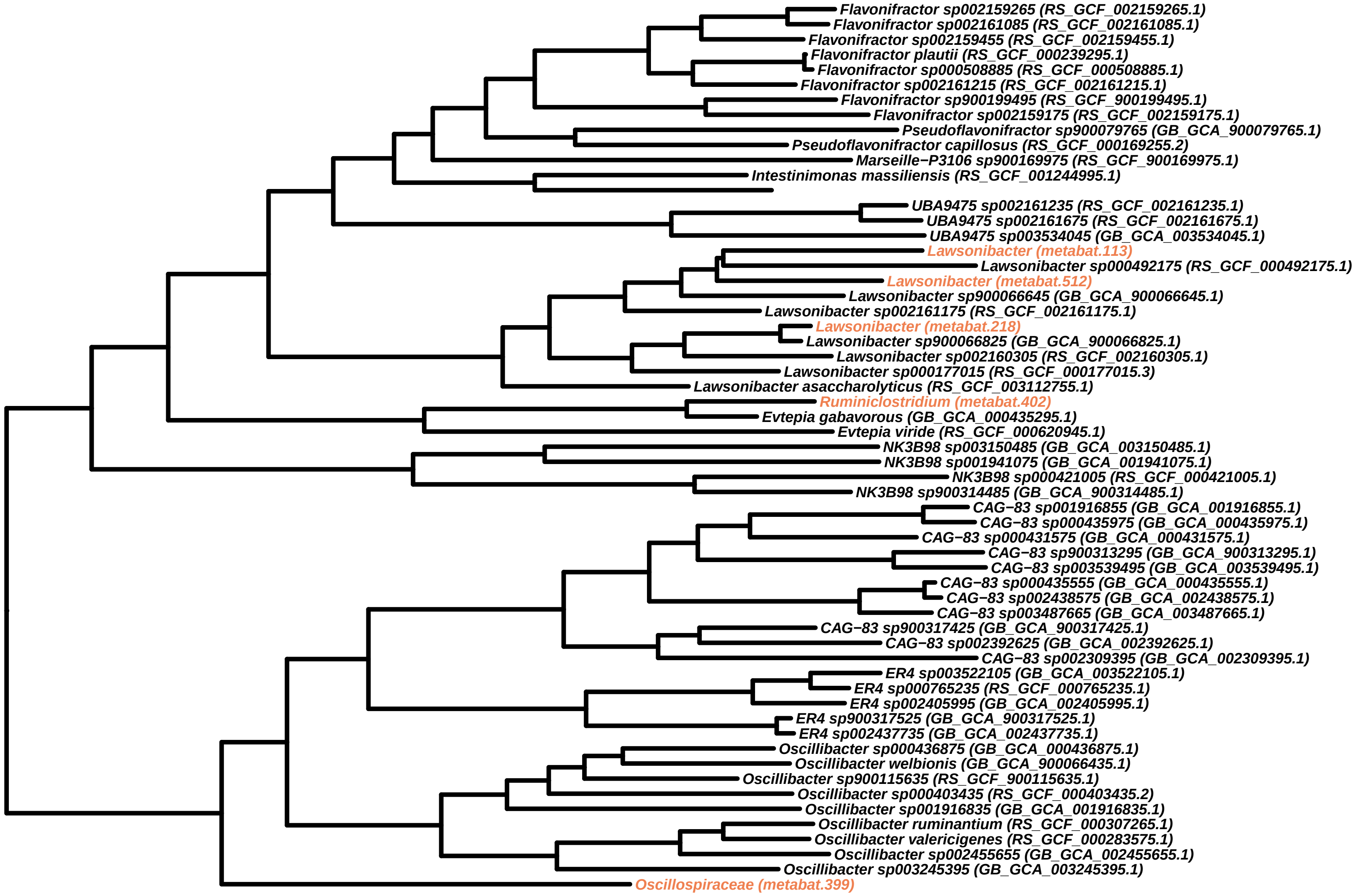
Flavonifractor sp002159265 (RS_GCF_002159265.1)
Flavonifractor sp002161085 (RS_GCF_002161085.1)
Flavonifractor sp002159455 (RS_GCF_002159455.1)
Flavonifractor plautii (RS_GCF_000239295.1)
Flavonifractor sp000508885 (RS_GCF_000508885.1)
Flavonifractor sp002161215 (RS_GCF_002161215.1)
Flavonifractor sp900199495 (RS_GCF_900199495.1)
Flavonifractor sp002159175 (RS_GCF_002159175.1)
Pseudoflavonifractor sp900079765 (GB_GCA_900079765.1)
Pseudoflavonifractor capillosus (RS_GCF_000169255.2)
Marseille−P3106 sp900169975 (RS_GCF_900169975.1)
Intestinimonas massiliensis (RS_GCF_001244995.1)
UBA9475 sp002161235 (RS_GCF_002161235.1)
UBA9475 sp002161675 (RS_GCF_002161675.1)
UBA9475 sp003534045 (GB_GCA_003534045.1)
Lawsonibacter (metabat.113)
Lawsonibacter sp000492175 (RS_GCF_000492175.1)
Lawsonibacter (metabat.512)
Lawsonibacter sp900066645 (GB_GCA_900066645.1)
Lawsonibacter sp002161175 (RS_GCF_002161175.1)
Lawsonibacter (metabat.218)
Lawsonibacter sp900066825 (GB_GCA_900066825.1)
Lawsonibacter sp002160305 (RS_GCF_002160305.1)
Lawsonibacter sp000177015 (RS_GCF_000177015.3)
Lawsonibacter asaccharolyticus (RS_GCF_003112755.1)
Ruminiclostridium (metabat.402)
Evtepia gabavorous (GB_GCA_000435295.1)
Evtepia viride (RS_GCF_000620945.1)
NK3B98 sp003150485 (GB_GCA_003150485.1)
NK3B98 sp001941075 (GB_GCA_001941075.1)
NK3B98 sp000421005 (RS_GCF_000421005.1)
NK3B98 sp900314485 (GB_GCA_900314485.1)
CAG−83 sp001916855 (GB_GCA_001916855.1)
CAG−83 sp000435975 (GB_GCA_000435975.1)
CAG−83 sp000431575 (GB_GCA_000431575.1)
CAG−83 sp900313295 (GB_GCA_900313295.1)
CAG−83 sp003539495 (GB_GCA_003539495.1)
CAG−83 sp000435555 (GB_GCA_000435555.1)
CAG−83 sp002438575 (GB_GCA_002438575.1)
CAG−83 sp003487665 (GB_GCA_003487665.1)
CAG−83 sp900317425 (GB_GCA_900317425.1)
CAG−83 sp002392625 (GB_GCA_002392625.1)
CAG−83 sp002309395 (GB_GCA_002309395.1)
ER4 sp003522105 (GB_GCA_003522105.1)
ER4 sp000765235 (RS_GCF_000765235.1)
ER4 sp002405995 (GB_GCA_002405995.1)
ER4 sp900317525 (GB_GCA_900317525.1)
ER4 sp002437735 (GB_GCA_002437735.1)
Oscillibacter sp000436875 (GB_GCA_000436875.1)
Oscillibacter welbionis (GB_GCA_900066435.1)
Oscillibacter sp900115635 (RS_GCF_900115635.1)
Oscillibacter sp000403435 (RS_GCF_000403435.2)
Oscillibacter sp001916835 (GB_GCA_001916835.1)
Oscillibacter ruminantium (RS_GCF_000307265.1)
Oscillibacter valericigenes (RS_GCF_000283575.1)
Oscillibacter sp002455655 (GB_GCA_002455655.1)
Oscillibacter sp003245395 (GB_GCA_003245395.1)
Oscillospiraceae (metabat.399)

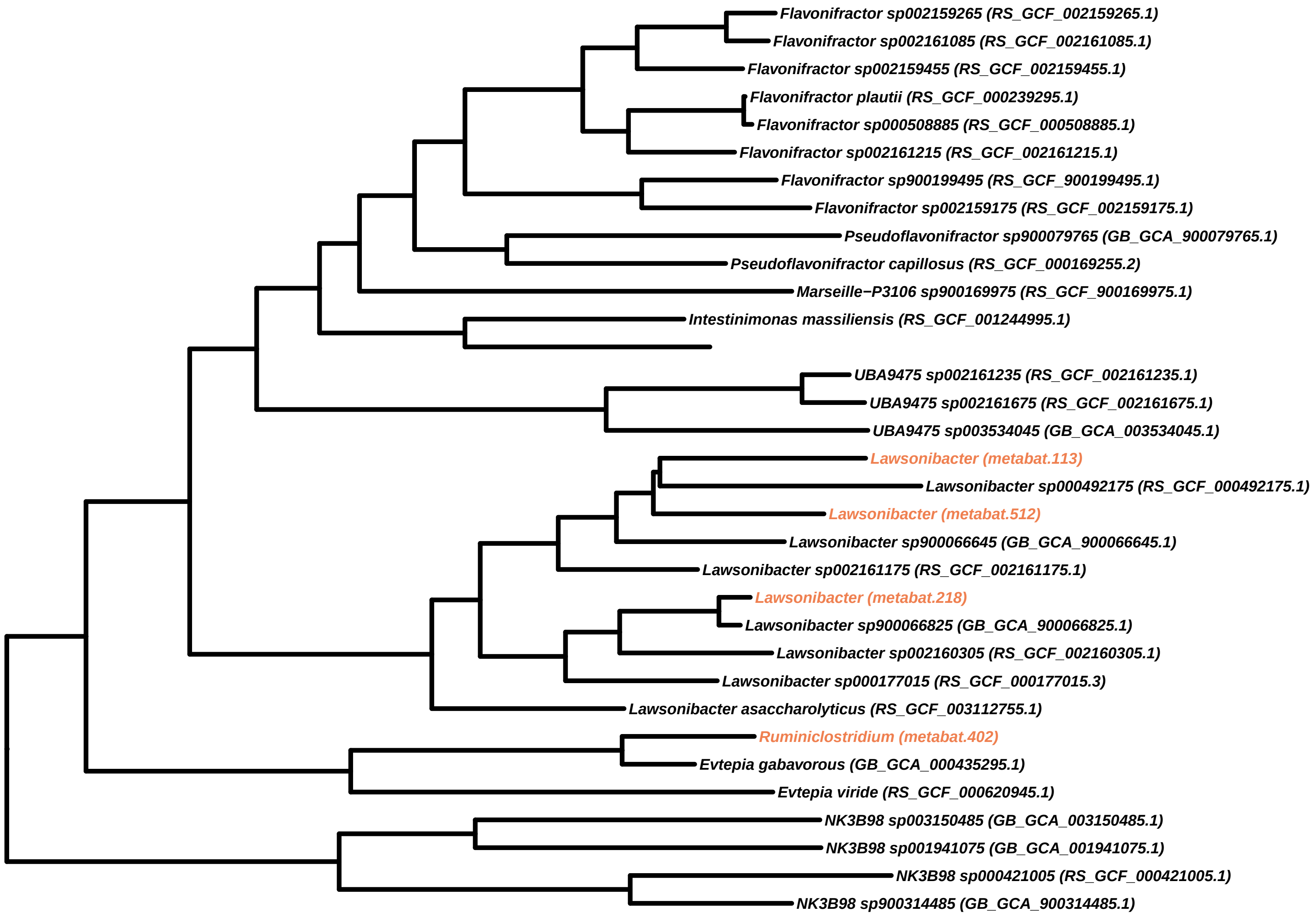
Flavonifractor sp002159265 (RS_GCF_002159265.1)
Flavonifractor sp002161085 (RS_GCF_002161085.1)
Flavonifractor sp002159455 (RS_GCF_002159455.1)
Flavonifractor plautii (RS_GCF_000239295.1)
Flavonifractor sp000508885 (RS_GCF_000508885.1)
Flavonifractor sp002161215 (RS_GCF_002161215.1)
Flavonifractor sp900199495 (RS_GCF_900199495.1)
Flavonifractor sp002159175 (RS_GCF_002159175.1)
Pseudoflavonifractor sp900079765 (GB_GCA_900079765.1)
Pseudoflavonifractor capillosus (RS_GCF_000169255.2)
Marseille−P3106 sp900169975 (RS_GCF_900169975.1)
Intestinimonas massiliensis (RS_GCF_001244995.1)
UBA9475 sp002161235 (RS_GCF_002161235.1)
UBA9475 sp002161675 (RS_GCF_002161675.1)
UBA9475 sp003534045 (GB_GCA_003534045.1)
Lawsonibacter (metabat.113)
Lawsonibacter sp000492175 (RS_GCF_000492175.1)
Lawsonibacter (metabat.512)
Lawsonibacter sp900066645 (GB_GCA_900066645.1)
Lawsonibacter sp002161175 (RS_GCF_002161175.1)
Lawsonibacter (metabat.218)
Lawsonibacter sp900066825 (GB_GCA_900066825.1)
Lawsonibacter sp002160305 (RS_GCF_002160305.1)
Lawsonibacter sp000177015 (RS_GCF_000177015.3)
Lawsonibacter asaccharolyticus (RS_GCF_003112755.1)
Ruminiclostridium (metabat.402)
Evtepia gabavorous (GB_GCA_000435295.1)
Evtepia viride (RS_GCF_000620945.1)
NK3B98 sp003150485 (GB_GCA_003150485.1)
NK3B98 sp001941075 (GB_GCA_001941075.1)
NK3B98 sp000421005 (RS_GCF_000421005.1)
NK3B98 sp900314485 (GB_GCA_900314485.1)

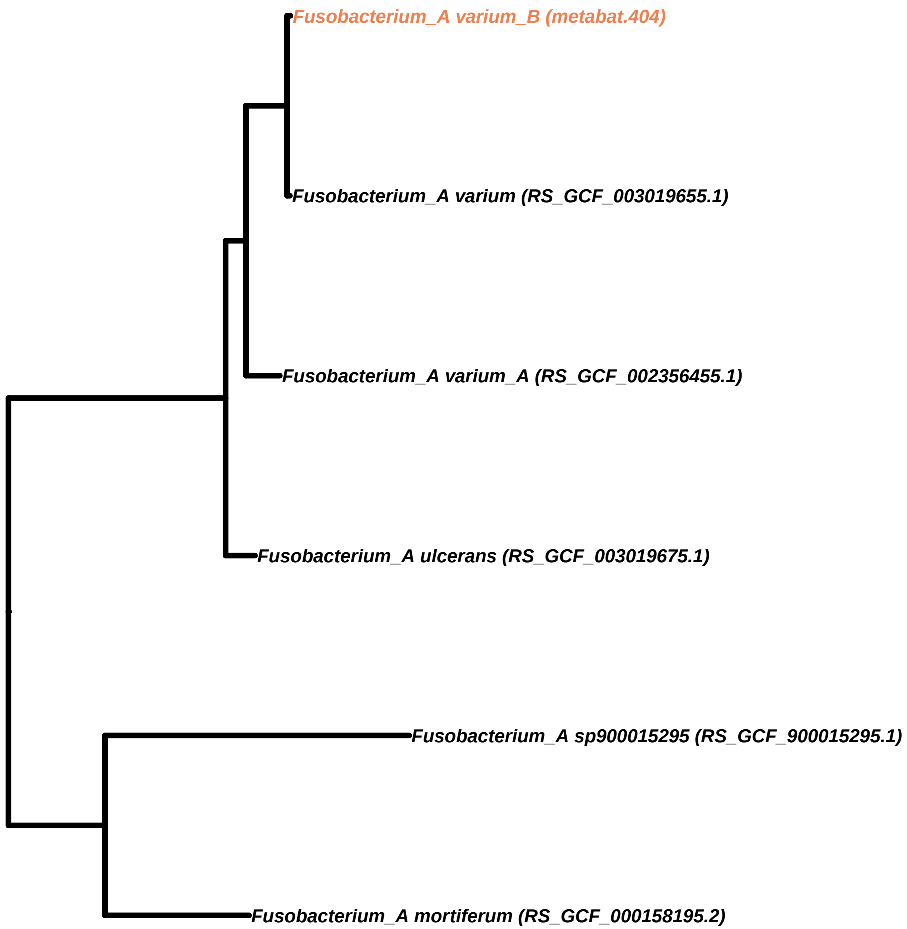
Fusobacterium_A varium_B (metabat.404)
Fusobacterium_A varium (RS_GCF_003019655.1)
Fusobacterium_A varium_A (RS_GCF_002356455.1)
Fusobacterium_A ulcerans (RS_GCF_003019675.1)
Fusobacterium_A sp900015295 (RS_GCF_900015295.1)
Fusobacterium_A mortiferum (RS_GCF_000158195.2)

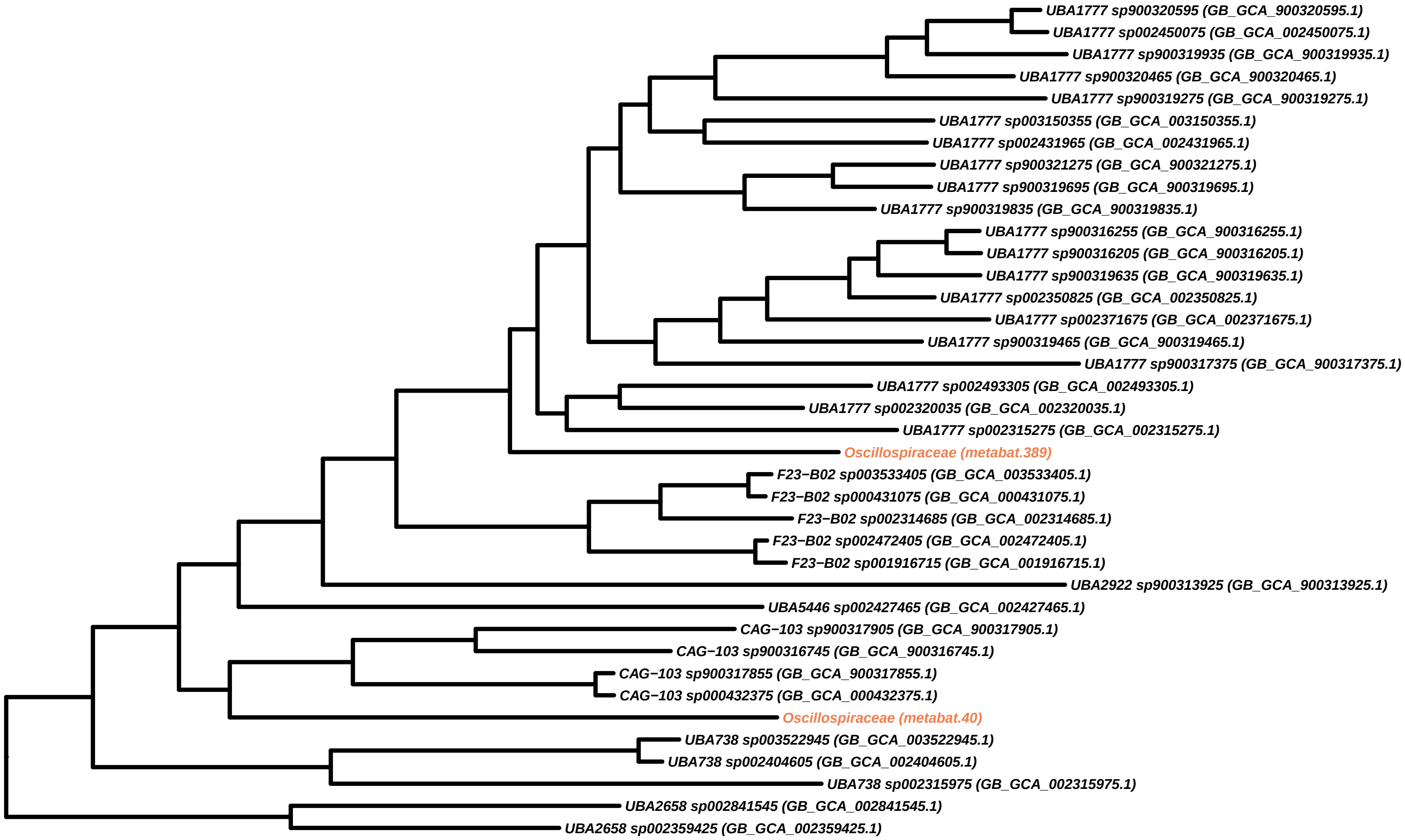
UBA1777 sp900320595 (GB_GCA_900320595.1)
UBA1777 sp002450075 (GB_GCA_002450075.1)
UBA1777 sp900319935 (GB_GCA_900319935.1)
UBA1777 sp900320465 (GB_GCA_900320465.1)
UBA1777 sp900319275 (GB_GCA_900319275.1)
UBA1777 sp003150355 (GB_GCA_003150355.1)
UBA1777 sp002431965 (GB_GCA_002431965.1)
UBA1777 sp900321275 (GB_GCA_900321275.1)
UBA1777 sp900319695 (GB_GCA_900319695.1)
UBA1777 sp900319835 (GB_GCA_900319835.1)
UBA1777 sp900316255 (GB_GCA_900316255.1)
UBA1777 sp900316205 (GB_GCA_900316205.1)
UBA1777 sp900319635 (GB_GCA_900319635.1)
UBA1777 sp002350825 (GB_GCA_002350825.1)
UBA1777 sp002371675 (GB_GCA_002371675.1)
UBA1777 sp900319465 (GB_GCA_900319465.1)
UBA1777 sp900317375 (GB_GCA_900317375.1)
UBA1777 sp002493305 (GB_GCA_002493305.1)
UBA1777 sp002320035 (GB_GCA_002320035.1)
UBA1777 sp002315275 (GB_GCA_002315275.1)
Oscillospiraceae (metabat.389)
F23−B02 sp003533405 (GB_GCA_003533405.1)
F23−B02 sp000431075 (GB_GCA_000431075.1)
F23−B02 sp002314685 (GB_GCA_002314685.1)
F23−B02 sp002472405 (GB_GCA_002472405.1)
F23−B02 sp001916715 (GB_GCA_001916715.1)
UBA2922 sp900313925 (GB_GCA_900313925.1)
UBA5446 sp002427465 (GB_GCA_002427465.1)
CAG−103 sp900317905 (GB_GCA_900317905.1)
CAG−103 sp900316745 (GB_GCA_900316745.1)
CAG−103 sp900317855 (GB_GCA_900317855.1)
CAG−103 sp000432375 (GB_GCA_000432375.1)
Oscillospiraceae (metabat.40)
UBA738 sp003522945 (GB_GCA_003522945.1)
UBA738 sp002404605 (GB_GCA_002404605.1)
UBA738 sp002315975 (GB_GCA_002315975.1)
UBA2658 sp002841545 (GB_GCA_002841545.1)
UBA2658 sp002359425 (GB_GCA_002359425.1)

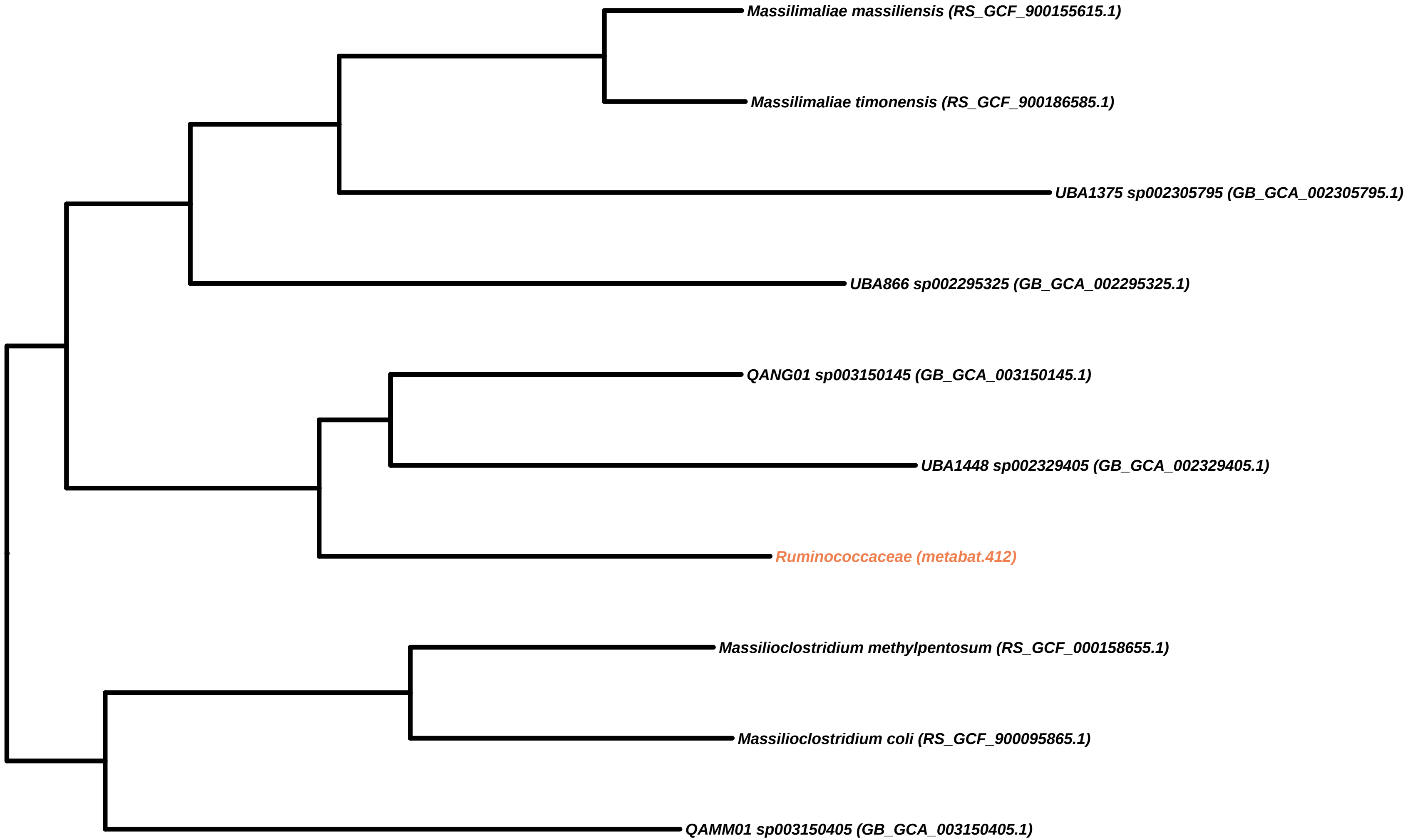
Massilimaliae massiliensis (RS_GCF_900155615.1)
Massilimaliae timonensis (RS_GCF_900186585.1)
UBA1375 sp002305795 (GB_GCA_002305795.1)
UBA866 sp002295325 (GB_GCA_002295325.1)
QANG01 sp003150145 (GB_GCA_003150145.1)
UBA1448 sp002329405 (GB_GCA_002329405.1)
Ruminococcaceae (metabat.412)
Massilioclostridium methylpentosum (RS_GCF_000158655.1)
Massilioclostridium coli (RS_GCF_900095865.1)
QAMM01 sp003150405 (GB_GCA_003150405.1)

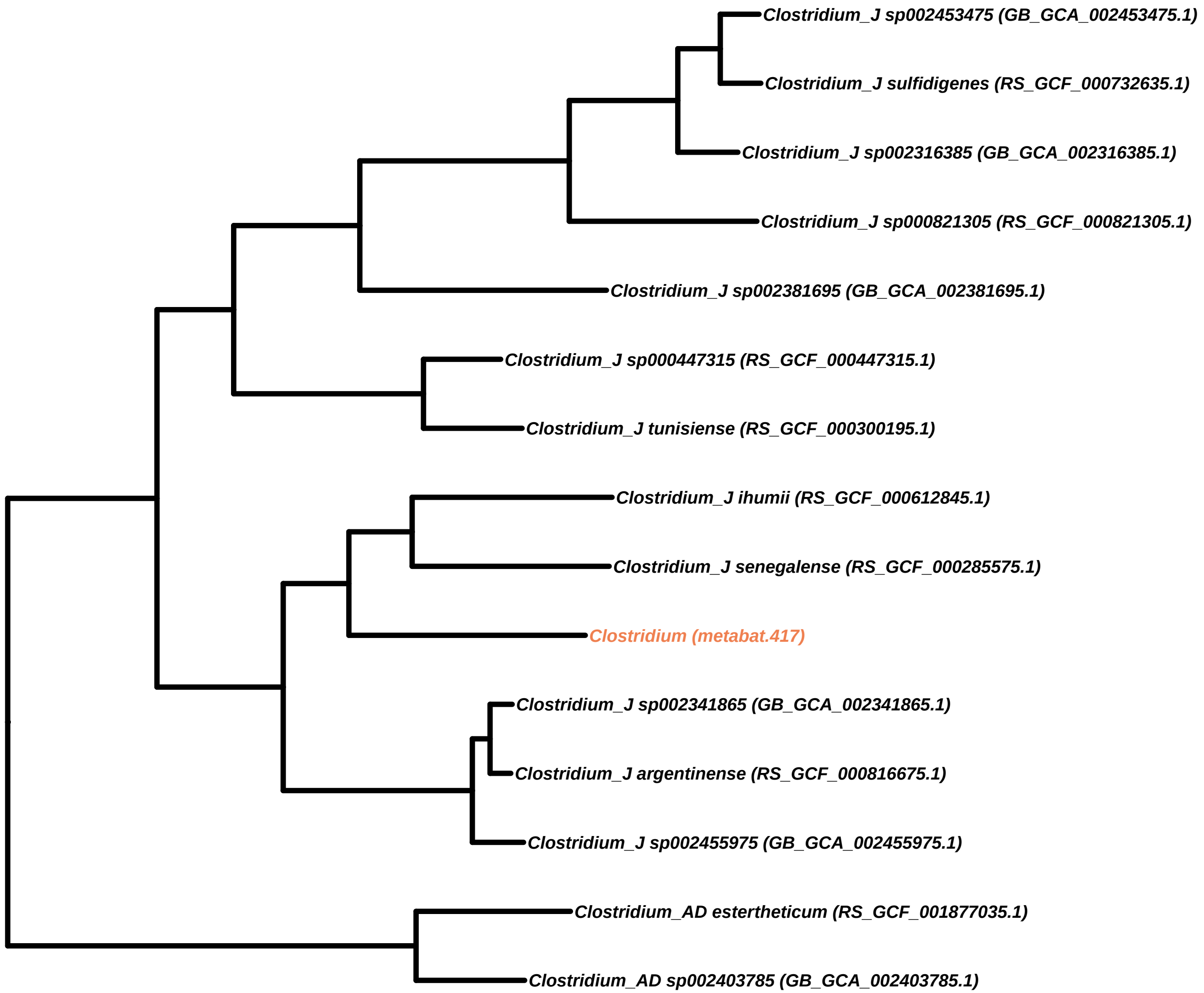

Clostridium_J sp002453475 (GB_GCA_002453475.1)
Clostridium_J sulfidigenes (RS_GCF_000732635.1)
Clostridium_J sp002316385 (GB_GCA_002316385.1)
Clostridium_J sp000821305 (RS_GCF_000821305.1)
Clostridium_J sp002381695 (GB_GCA_002381695.1)
Clostridium_J sp000447315 (RS_GCF_000447315.1)
Clostridium_J tunisiense (RS_GCF_000300195.1)
Clostridium_J ihumii (RS_GCF_000612845.1)
Clostridium_J senegalense (RS_GCF_000285575.1)
Clostridium (metabat.417)
Clostridium_J sp002341865 (GB_GCA_002341865.1)
Clostridium_J argentinense (RS_GCF_000816675.1)
Clostridium_J sp002455975 (GB_GCA_002455975.1)
Clostridium_AD estertheticum (RS_GCF_001877035.1)
Clostridium_AD sp002403785 (GB_GCA_002403785.1)

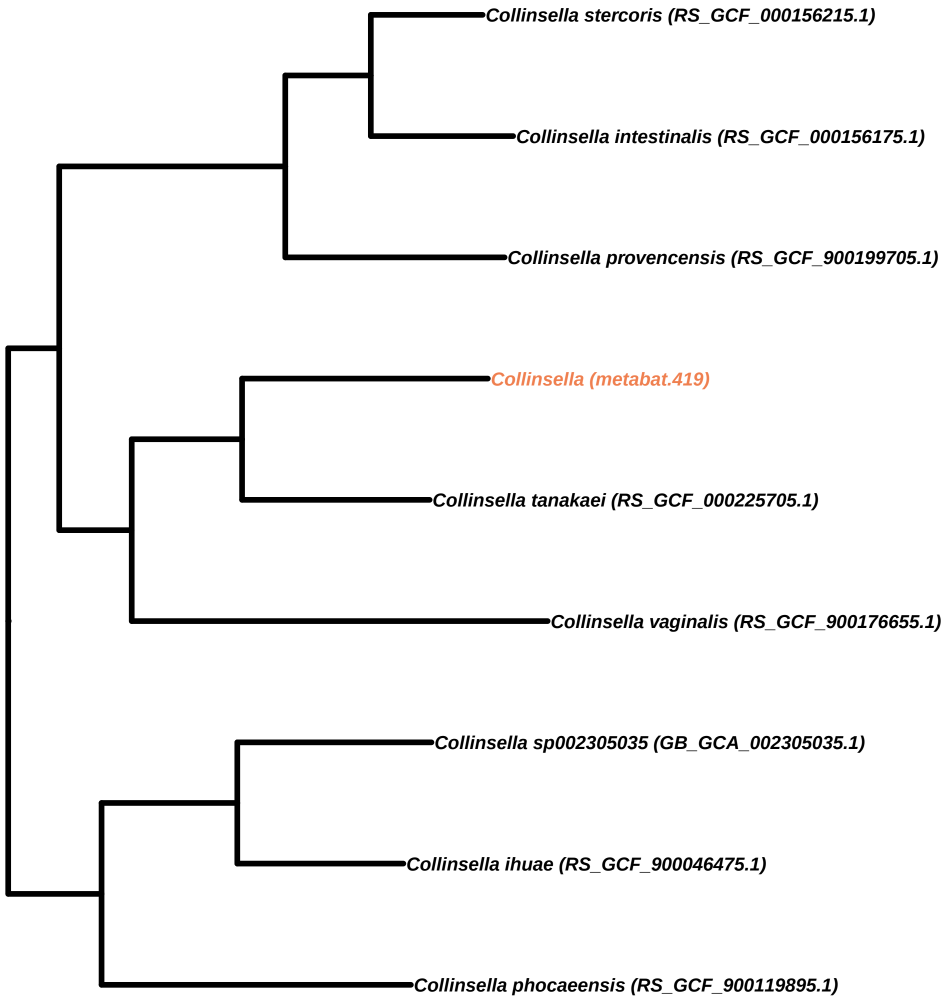
Collinsella stercoris (RS_GCF_000156215.1)
Collinsella intestinalis (RS_GCF_000156175.1)
Collinsella provencensis (RS_GCF_900199705.1)
Collinsella (metabat.419)
Collinsella tanakaei (RS_GCF_000225705.1)
Collinsella vaginalis (RS_GCF_900176655.1)
Collinsella sp002305035 (GB_GCA_002305035.1)
Collinsella ihuae (RS_GCF_900046475.1)
Collinsella phocaeensis (RS_GCF_900119895.1)

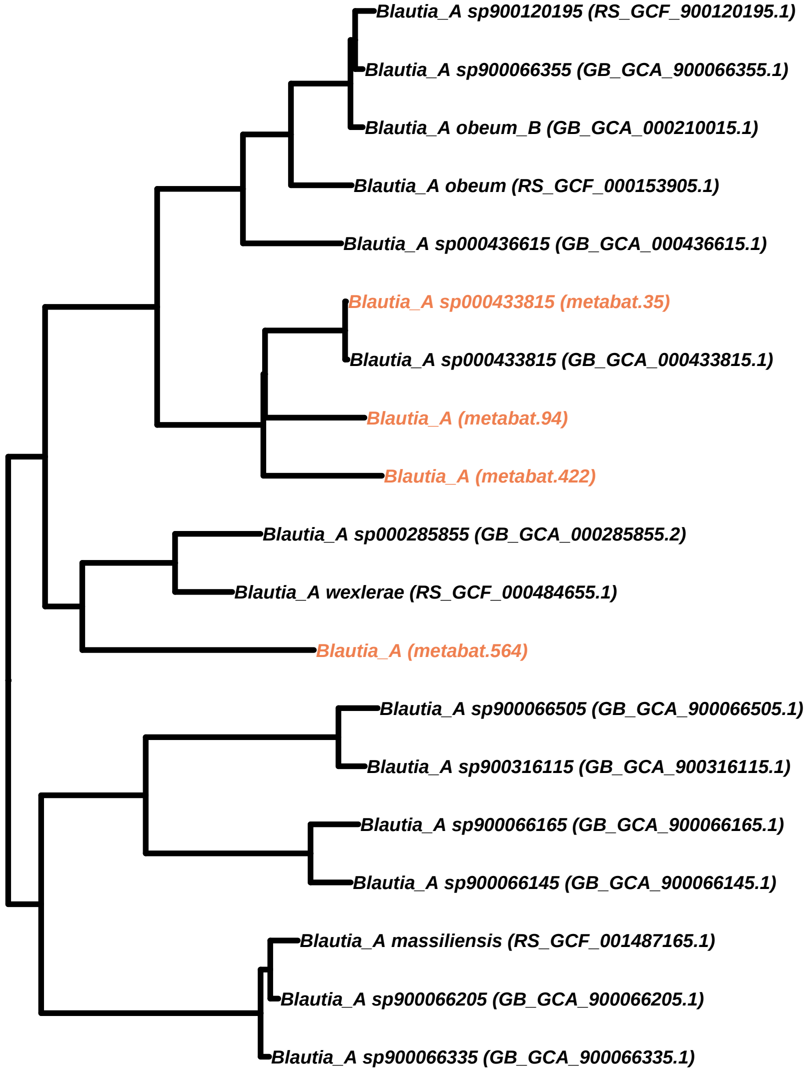
Blautia_A sp900120195 (RS_GCF_900120195.1)
Blautia_A sp900066355 (GB_GCA_900066355.1)
Blautia_A obeum_B (GB_GCA_000210015.1)
Blautia_A obeum (RS_GCF_000153905.1)
Blautia_A sp000436615 (GB_GCA_000436615.1)
Blautia_A sp000433815 (metabat.35)
Blautia_A sp000433815 (GB_GCA_000433815.1)
Blautia_A (metabat.94)
Blautia_A (metabat.422)
Blautia_A sp000285855 (GB_GCA_000285855.2)
Blautia_A wexlerae (RS_GCF_000484655.1)
Blautia_A (metabat.564)
Blautia_A sp900066505 (GB_GCA_900066505.1)
Blautia_A sp900316115 (GB_GCA_900316115.1)
Blautia_A sp900066165 (GB_GCA_900066165.1)
Blautia_A sp900066145 (GB_GCA_900066145.1)
Blautia_A massiliensis (RS_GCF_001487165.1)
Blautia_A sp900066205 (GB_GCA_900066205.1)
Blautia_A sp900066335 (GB_GCA_900066335.1)

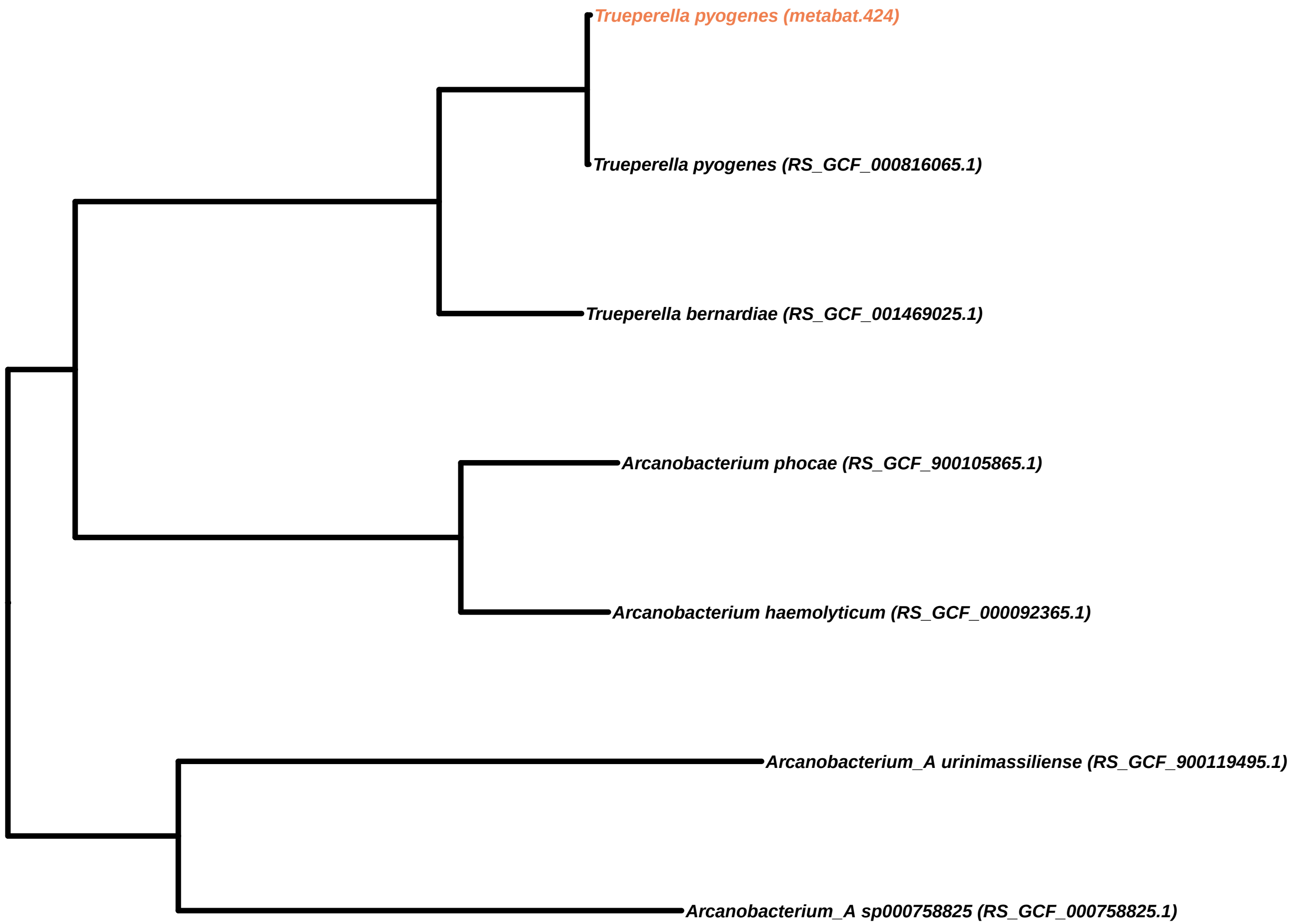
Trueperella pyogenes (metabat.424)
Trueperella pyogenes (RS_GCF_000816065.1)
Trueperella bernardiae (RS_GCF_001469025.1)
Arcanobacterium phocae (RS_GCF_900105865.1)
Arcanobacterium haemolyticum (RS_GCF_000092365.1)
Arcanobacterium_A urinimassiliense (RS_GCF_900119495.1)
Arcanobacterium_A sp000758825 (RS_GCF_000758825.1)

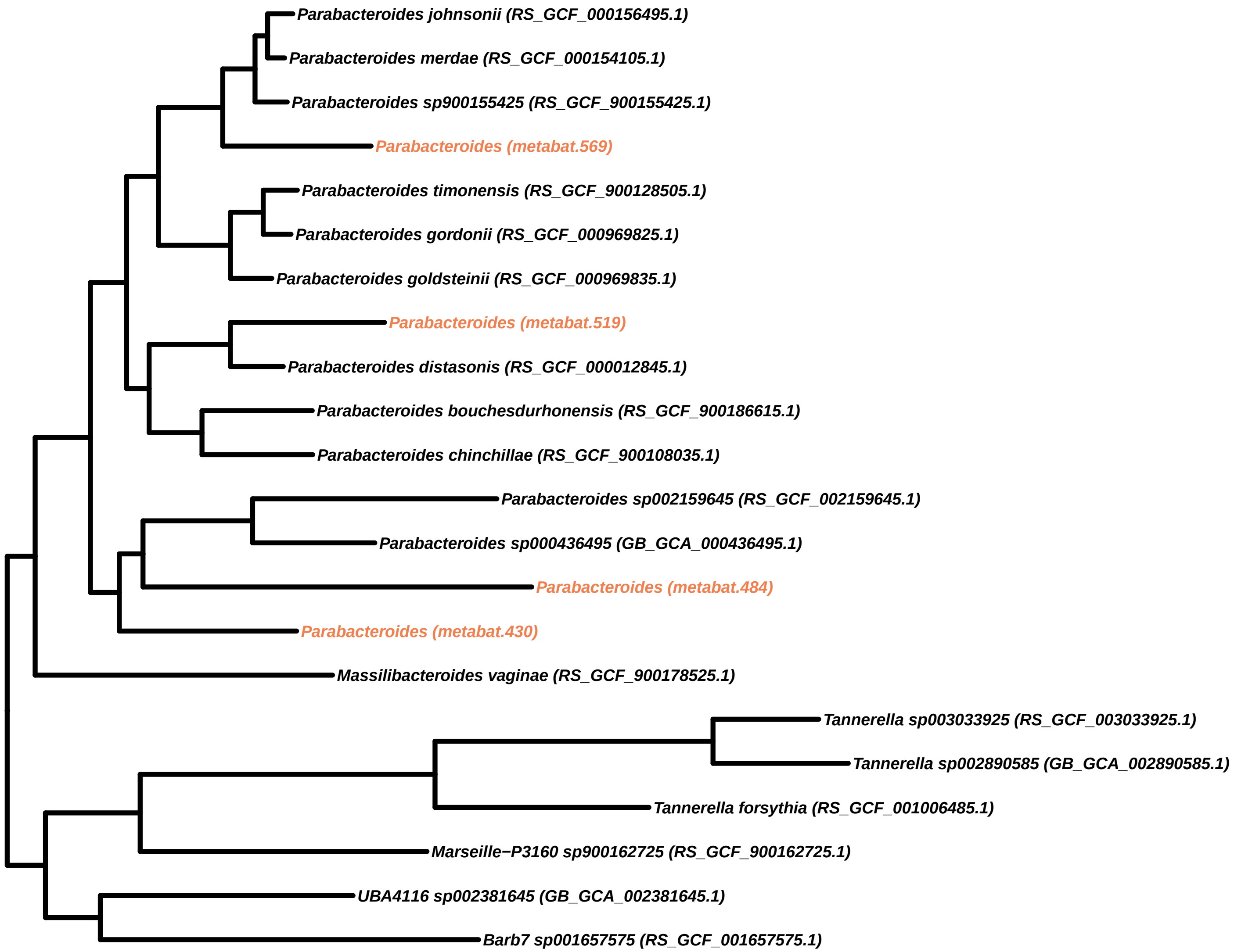
Parabacteroides johnsonii (RS_GCF_000156495.1)
Parabacteroides merdae (RS_GCF_000154105.1)
Parabacteroides sp900155425 (RS_GCF_900155425.1)
Parabacteroides (metabat.569)
Parabacteroides timonensis (RS_GCF_900128505.1)
Parabacteroides gordonii (RS_GCF_000969825.1)
Parabacteroides goldsteinii (RS_GCF_000969835.1)
Parabacteroides (metabat.519)
Parabacteroides distasonis (RS_GCF_000012845.1)
Parabacteroides bouchesdurhonensis (RS_GCF_900186615.1)
Parabacteroides chinchillae (RS_GCF_900108035.1)
Parabacteroides sp002159645 (RS_GCF_002159645.1)
Parabacteroides sp000436495 (GB_GCA_000436495.1)
Parabacteroides (metabat.484)
Parabacteroides (metabat.430)
Massilibacteroides vaginae (RS_GCF_900178525.1)
Tannerella sp003033925 (RS_GCF_003033925.1)
Tannerella sp002890585 (GB_GCA_002890585.1)
Tannerella forsythia (RS_GCF_001006485.1)
Marseille−P3160 sp900162725 (RS_GCF_900162725.1)
UBA4116 sp002381645 (GB_GCA_002381645.1)
Barb7 sp001657575 (RS_GCF_001657575.1)

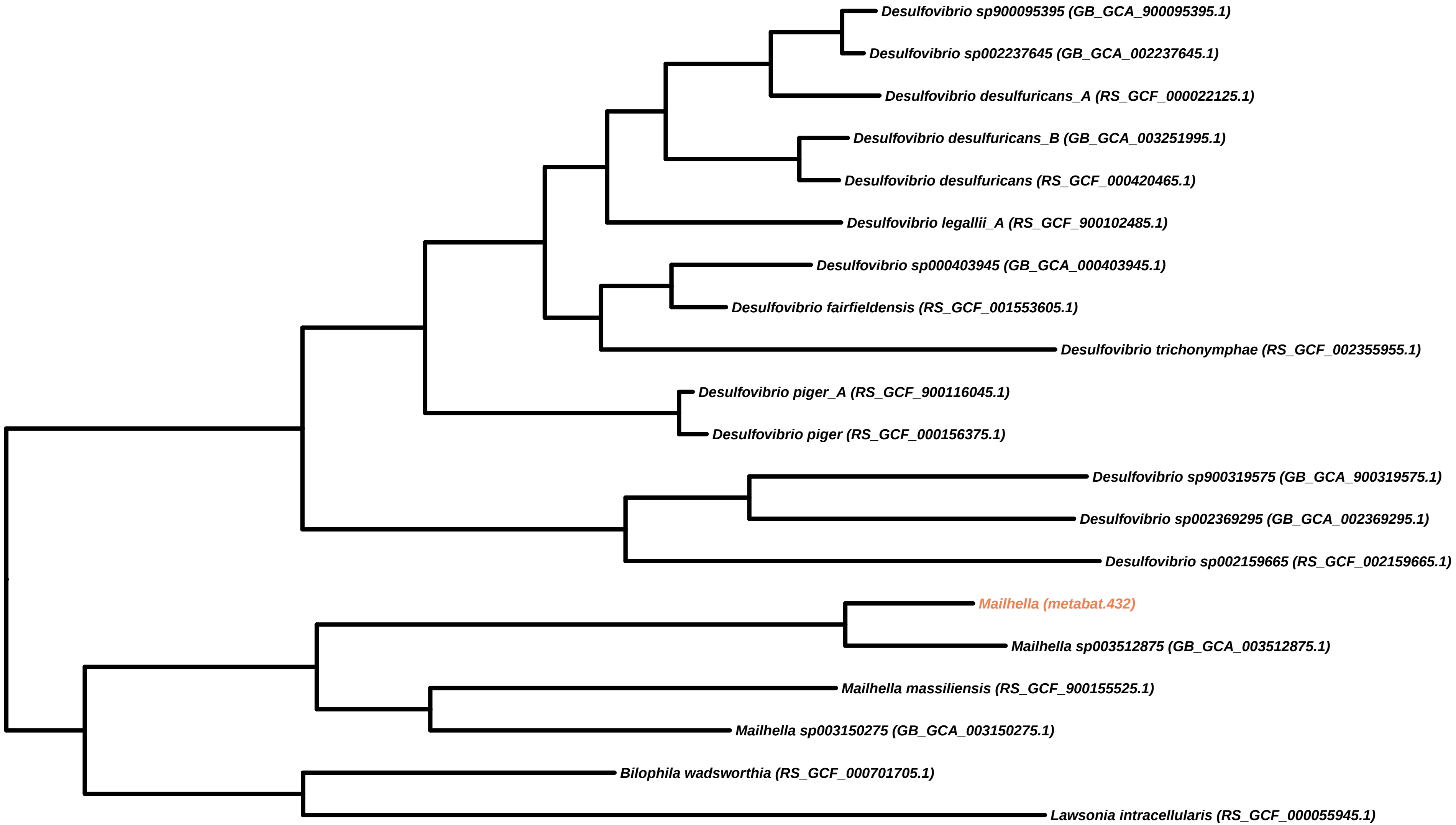
Desulfovibrio sp900095395 (GB_GCA_900095395.1)
Desulfovibrio sp002237645 (GB_GCA_002237645.1)
Desulfovibrio desulfuricans_A (RS_GCF_000022125.1)
Desulfovibrio desulfuricans_B (GB_GCA_003251995.1)
Desulfovibrio desulfuricans (RS_GCF_000420465.1)
Desulfovibrio legallii_A (RS_GCF_900102485.1)
Desulfovibrio sp000403945 (GB_GCA_000403945.1)
Desulfovibrio fairfieldensis (RS_GCF_001553605.1)
Desulfovibrio trichonymphae (RS_GCF_002355955.1)
Desulfovibrio piger_A (RS_GCF_900116045.1)
Desulfovibrio piger (RS_GCF_000156375.1)
Desulfovibrio sp900319575 (GB_GCA_900319575.1)
Desulfovibrio sp002369295 (GB_GCA_002369295.1)
Desulfovibrio sp002159665 (RS_GCF_002159665.1)
Mailhella (metabat.432)
Mailhella sp003512875 (GB_GCA_003512875.1)
Mailhella massiliensis (RS_GCF_900155525.1)
Mailhella sp003150275 (GB_GCA_003150275.1)
Bilophila wadsworthia (RS_GCF_000701705.1)
Lawsonia intracellularis (RS_GCF_000055945.1)

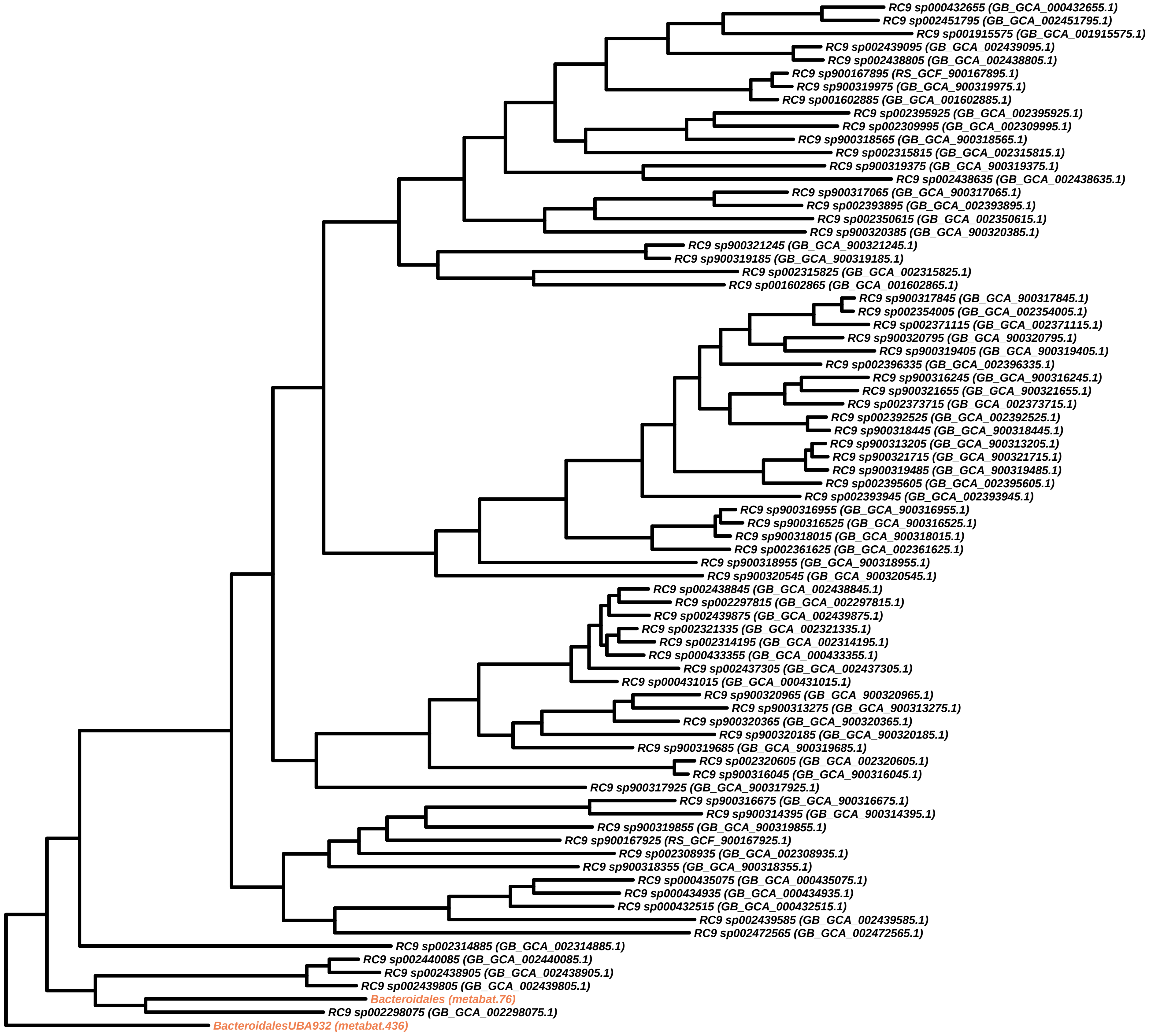
RC9 sp000432655 (GB_GCA_000432655.1)
RC9 sp002451795 (GB_GCA_002451795.1)
RC9 sp001915575 (GB_GCA_001915575.1)
RC9 sp002439095 (GB_GCA_002439095.1)
RC9 sp002438805 (GB_GCA_002438805.1)
RC9 sp900167895 (RS_GCF_900167895.1)
RC9 sp900319975 (GB_GCA_900319975.1)
RC9 sp001602885 (GB_GCA_001602885.1)
RC9 sp002395925 (GB_GCA_002395925.1)
RC9 sp002309995 (GB_GCA_002309995.1)
RC9 sp900318565 (GB_GCA_900318565.1)
RC9 sp002315815 (GB_GCA_002315815.1)
RC9 sp900319375 (GB_GCA_900319375.1)
RC9 sp002438635 (GB_GCA_002438635.1)
RC9 sp900317065 (GB_GCA_900317065.1)
RC9 sp002393895 (GB_GCA_002393895.1)
RC9 sp002350615 (GB_GCA_002350615.1)
RC9 sp900320385 (GB_GCA_900320385.1)
RC9 sp900321245 (GB_GCA_900321245.1)
RC9 sp900319185 (GB_GCA_900319185.1)
RC9 sp002315825 (GB_GCA_002315825.1)
RC9 sp001602865 (GB_GCA_001602865.1)
RC9 sp900317845 (GB_GCA_900317845.1)
RC9 sp002354005 (GB_GCA_002354005.1)
RC9 sp002371115 (GB_GCA_002371115.1)
RC9 sp900320795 (GB_GCA_900320795.1)
RC9 sp900319405 (GB_GCA_900319405.1)
RC9 sp002396335 (GB_GCA_002396335.1)
RC9 sp900316245 (GB_GCA_900316245.1)
RC9 sp900321655 (GB_GCA_900321655.1)
RC9 sp002373715 (GB_GCA_002373715.1)
RC9 sp002392525 (GB_GCA_002392525.1)
RC9 sp900318445 (GB_GCA_900318445.1)
RC9 sp900313205 (GB_GCA_900313205.1)
RC9 sp900321715 (GB_GCA_900321715.1)
RC9 sp900319485 (GB_GCA_900319485.1)
RC9 sp002395605 (GB_GCA_002395605.1)
RC9 sp002393945 (GB_GCA_002393945.1)
RC9 sp900316955 (GB_GCA_900316955.1)
RC9 sp900316525 (GB_GCA_900316525.1)
RC9 sp900318015 (GB_GCA_900318015.1)
RC9 sp002361625 (GB_GCA_002361625.1)
RC9 sp900318955 (GB_GCA_900318955.1)
RC9 sp900320545 (GB_GCA_900320545.1)
RC9 sp002438845 (GB_GCA_002438845.1)
RC9 sp002297815 (GB_GCA_002297815.1)
RC9 sp002439875 (GB_GCA_002439875.1)
RC9 sp002321335 (GB_GCA_002321335.1)
RC9 sp002314195 (GB_GCA_002314195.1)
RC9 sp000433355 (GB_GCA_000433355.1)
RC9 sp002437305 (GB_GCA_002437305.1)
RC9 sp000431015 (GB_GCA_000431015.1)
RC9 sp900320965 (GB_GCA_900320965.1)
RC9 sp900313275 (GB_GCA_900313275.1)
RC9 sp900320365 (GB_GCA_900320365.1)
RC9 sp900320185 (GB_GCA_900320185.1)
RC9 sp900319685 (GB_GCA_900319685.1)
RC9 sp002320605 (GB_GCA_002320605.1)
RC9 sp900316045 (GB_GCA_900316045.1)
RC9 sp900317925 (GB_GCA_900317925.1)
RC9 sp900316675 (GB_GCA_900316675.1)
RC9 sp900314395 (GB_GCA_900314395.1)
RC9 sp900319855 (GB_GCA_900319855.1)
RC9 sp900167925 (RS_GCF_900167925.1)
RC9 sp002308935 (GB_GCA_002308935.1)
RC9 sp900318355 (GB_GCA_900318355.1)
RC9 sp000435075 (GB_GCA_000435075.1)
RC9 sp000434935 (GB_GCA_000434935.1)
RC9 sp000432515 (GB_GCA_000432515.1)
RC9 sp002439585 (GB_GCA_002439585.1)
RC9 sp002472565 (GB_GCA_002472565.1)
RC9 sp002314885 (GB_GCA_002314885.1)
RC9 sp002440085 (GB_GCA_002440085.1)
RC9 sp002438905 (GB_GCA_002438905.1)
RC9 sp002439805 (GB_GCA_002439805.1)
Bacteroidales (metabat.76)
RC9 sp002298075 (GB_GCA_002298075.1)
BacteroidalesUBA932 (metabat.436)

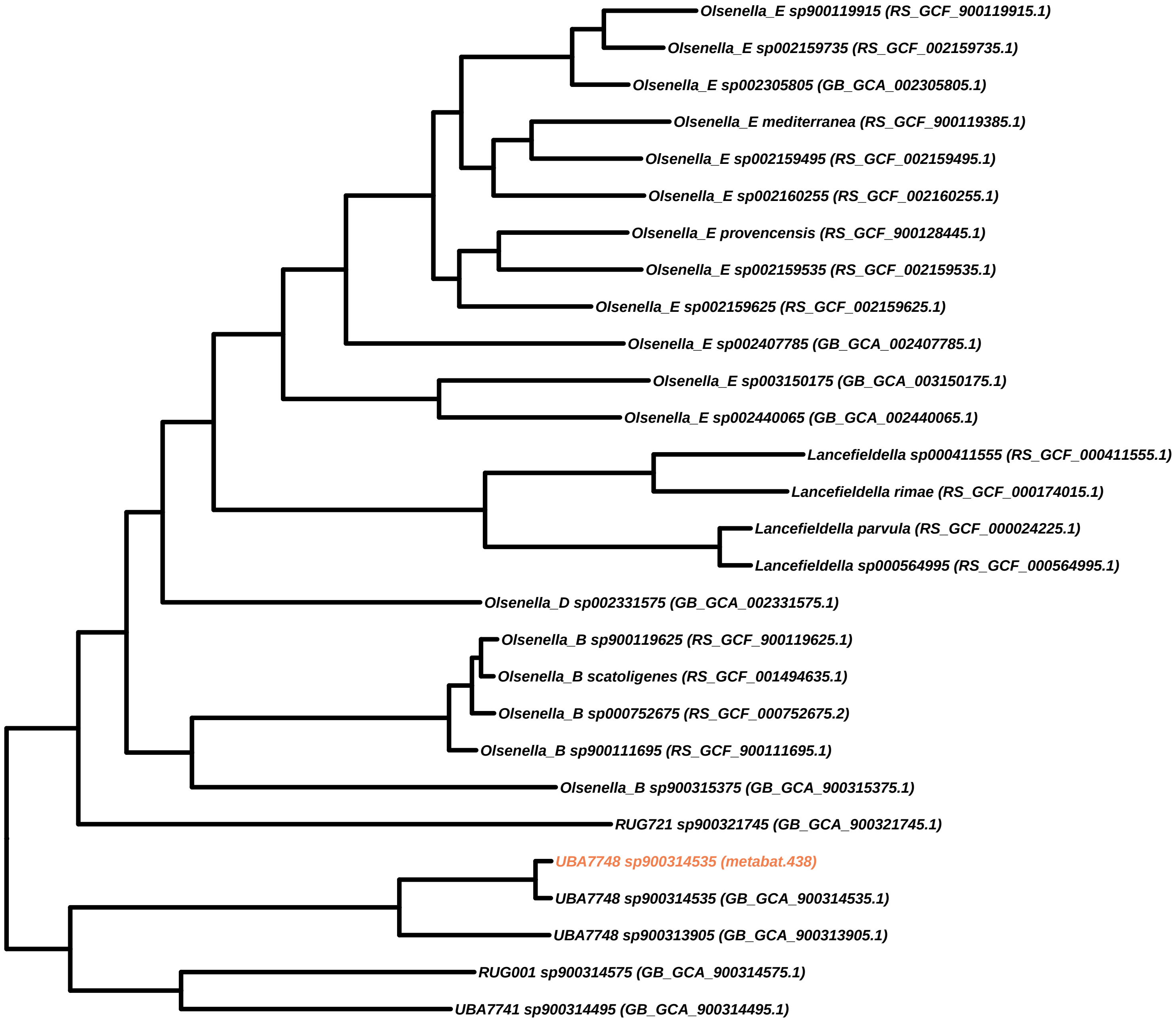
Olsenella_E sp900119915 (RS_GCF_900119915.1)
Olsenella_E sp002159735 (RS_GCF_002159735.1)
Olsenella_E sp002305805 (GB_GCA_002305805.1)
Olsenella_E mediterranea (RS_GCF_900119385.1)
Olsenella_E sp002159495 (RS_GCF_002159495.1)
Olsenella_E sp002160255 (RS_GCF_002160255.1)
Olsenella_E provencensis (RS_GCF_900128445.1)
Olsenella_E sp002159535 (RS_GCF_002159535.1)
Olsenella_E sp002159625 (RS_GCF_002159625.1)
Olsenella_E sp002407785 (GB_GCA_002407785.1)
Olsenella_E sp003150175 (GB_GCA_003150175.1)
Olsenella_E sp002440065 (GB_GCA_002440065.1)
Lancefieldella sp000411555 (RS_GCF_000411555.1)
Lancefieldella rimae (RS_GCF_000174015.1)
Lancefieldella parvula (RS_GCF_000024225.1)
Lancefieldella sp000564995 (RS_GCF_000564995.1)
Olsenella_D sp002331575 (GB_GCA_002331575.1)
Olsenella_B sp900119625 (RS_GCF_900119625.1)
Olsenella_B scatoligenes (RS_GCF_001494635.1)
Olsenella_B sp000752675 (RS_GCF_000752675.2)
Olsenella_B sp900111695 (RS_GCF_900111695.1)
Olsenella_B sp900315375 (GB_GCA_900315375.1)
RUG721 sp900321745 (GB_GCA_900321745.1)
UBA7748 sp900314535 (metabat.438)
UBA7748 sp900314535 (GB_GCA_900314535.1)
UBA7748 sp900313905 (GB_GCA_900313905.1)
RUG001 sp900314575 (GB_GCA_900314575.1)
UBA7741 sp900314495 (GB_GCA_900314495.1)

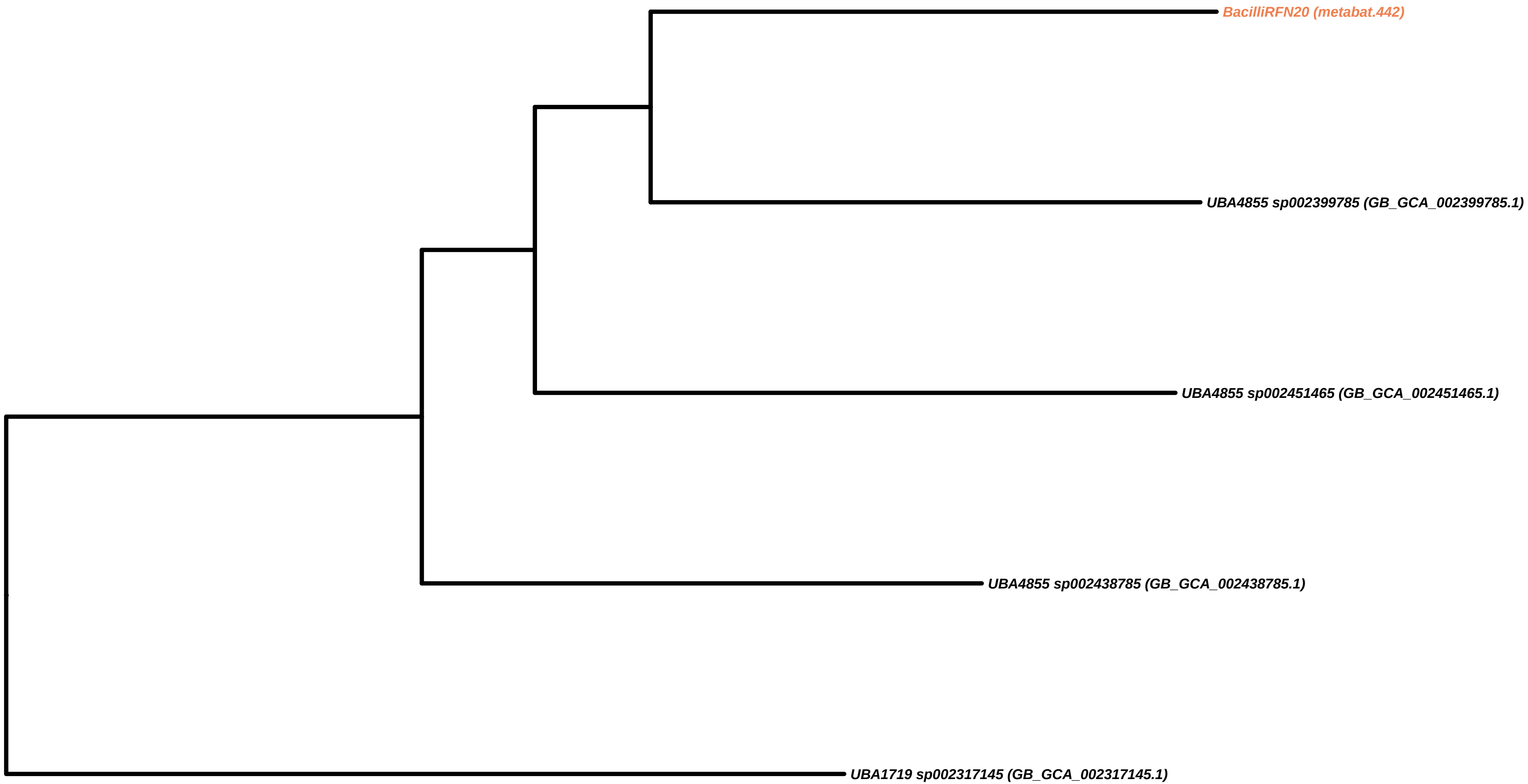
BacilliRFN20 (metabat.442)
UBA4855 sp002399785 (GB_GCA_002399785.1)
UBA4855 sp002451465 (GB_GCA_002451465.1)
UBA4855 sp002438785 (GB_GCA_002438785.1)
UBA1719 sp002317145 (GB_GCA_002317145.1)

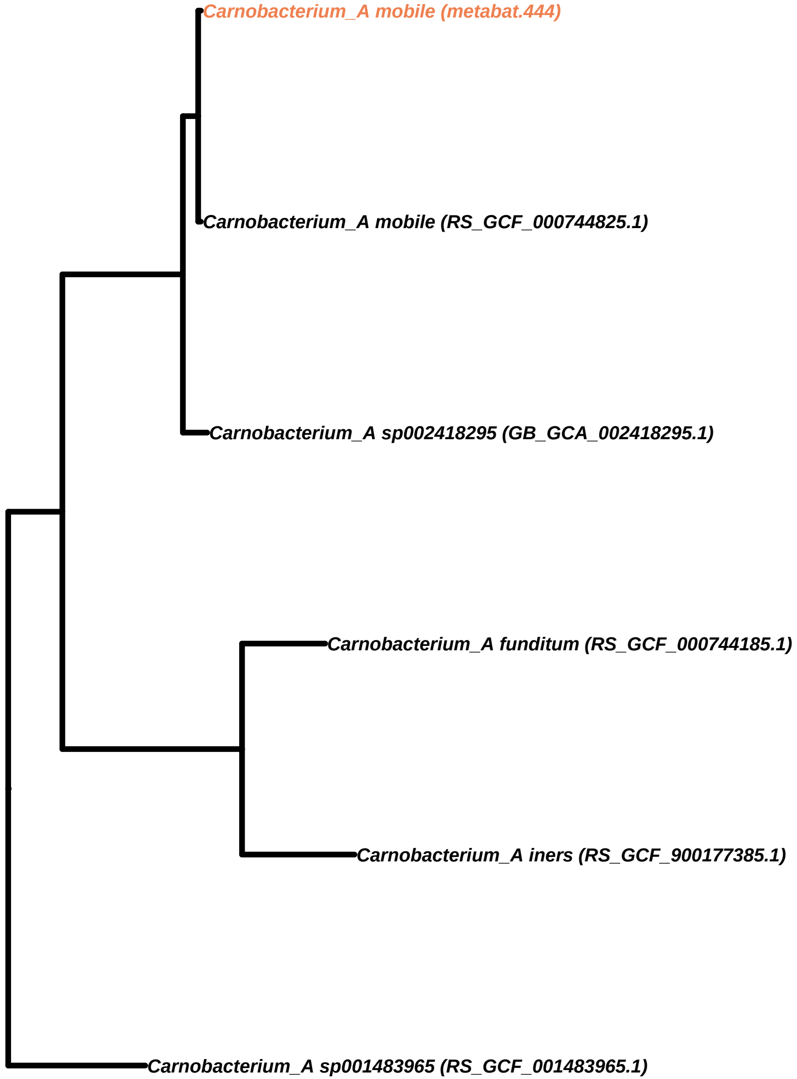
Carnobacterium_A mobile (metabat.444)
Carnobacterium_A mobile (RS_GCF_000744825.1)
Carnobacterium_A sp002418295 (GB_GCA_002418295.1)
Carnobacterium_A funditum (RS_GCF_000744185.1)
Carnobacterium_A iners (RS_GCF_900177385.1)
Carnobacterium_A sp001483965 (RS_GCF_001483965.1)

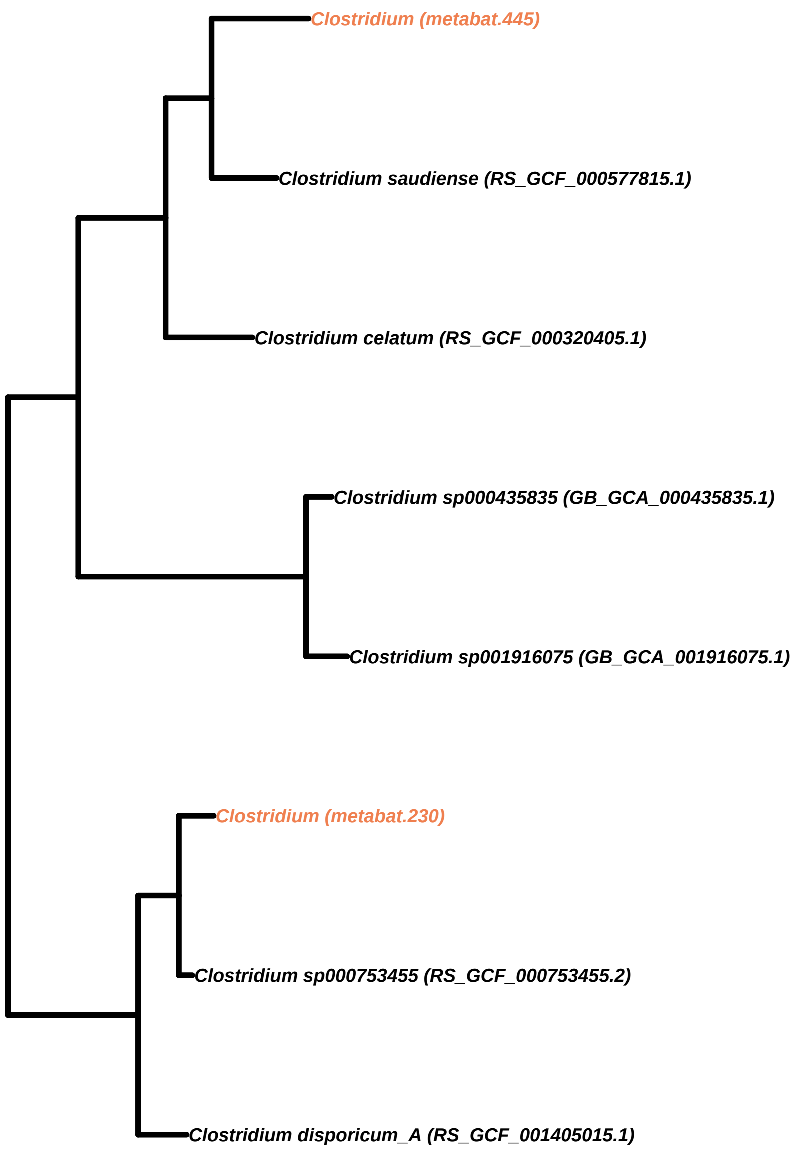

Clostridium (metabat.445)
Clostridium saudiense (RS_GCF_000577815.1)
Clostridium celatum (RS_GCF_000320405.1)
Clostridium sp000435835 (GB_GCA_000435835.1)
Clostridium sp001916075 (GB_GCA_001916075.1)
Clostridium (metabat.230)
Clostridium sp000753455 (RS_GCF_000753455.2)
Clostridium disporicum_A (RS_GCF_001405015.1)

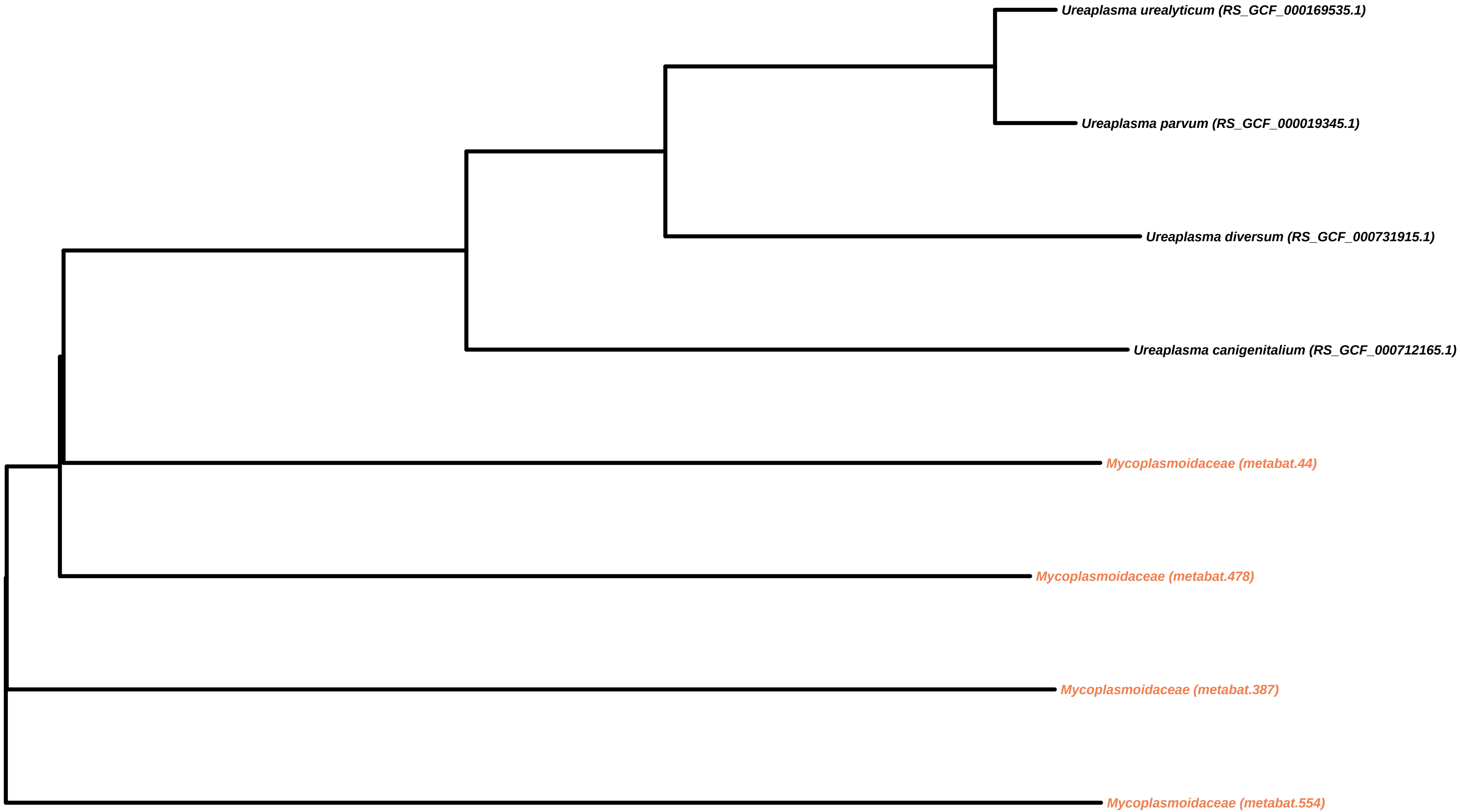
Ureaplasma urealyticum (RS_GCF_000169535.1)
Ureaplasma parvum (RS_GCF_000019345.1)
Ureaplasma diversum (RS_GCF_000731915.1)
Ureaplasma canigenitalium (RS_GCF_000712165.1)
Mycoplasmoidaceae (metabat.44)
Mycoplasmoidaceae (metabat.478)
Mycoplasmoidaceae (metabat.387)
Mycoplasmoidaceae (metabat.554)

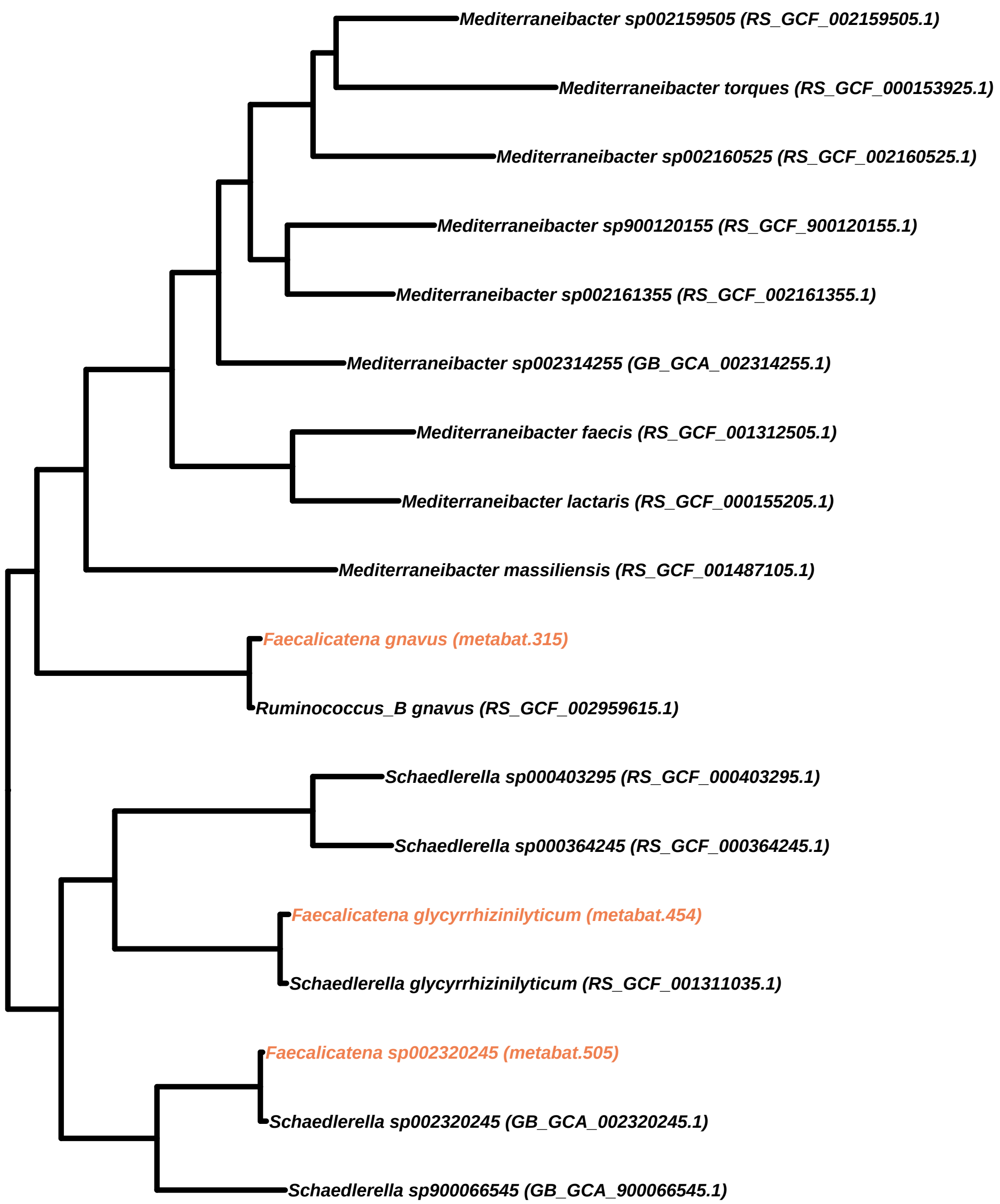
Mediterraneibacter sp002159505 (RS_GCF_002159505.1)
Mediterraneibacter torques (RS_GCF_000153925.1)
Mediterraneibacter sp002160525 (RS_GCF_002160525.1)
Mediterraneibacter sp900120155 (RS_GCF_900120155.1)
Mediterraneibacter sp002161355 (RS_GCF_002161355.1)
Mediterraneibacter sp002314255 (GB_GCA_002314255.1)
Mediterraneibacter faecis (RS_GCF_001312505.1)
Mediterraneibacter lactaris (RS_GCF_000155205.1)
Mediterraneibacter massiliensis (RS_GCF_001487105.1)
Faecalicatena gnavus (metabat.315)
Ruminococcus_B gnavus (RS_GCF_002959615.1)
Schaedlerella sp000403295 (RS_GCF_000403295.1)
Schaedlerella sp000364245 (RS_GCF_000364245.1)
Faecalicatena glycyrrhizinilyticum (metabat.454)
Schaedlerella glycyrrhizinilyticum (RS_GCF_001311035.1)
Faecalicatena sp002320245 (metabat.505)
Schaedlerella sp002320245 (GB_GCA_002320245.1)
Schaedlerella sp900066545 (GB_GCA_900066545.1)

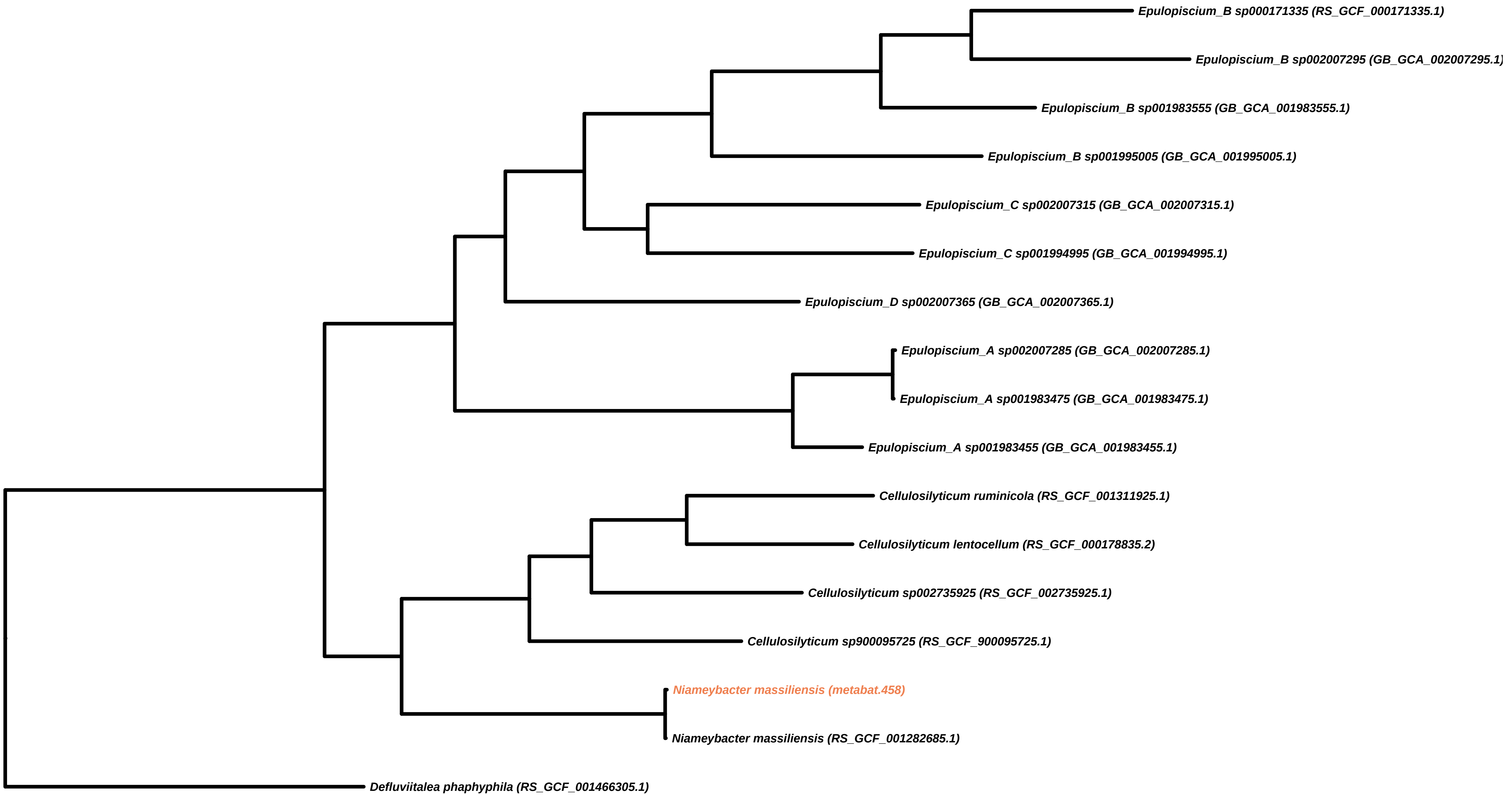
Epulopiscium_B sp000171335 (RS_GCF_000171335.1)
Epulopiscium_B sp002007295 (GB_GCA_002007295.1)
Epulopiscium_B sp001983555 (GB_GCA_001983555.1)
Epulopiscium_B sp001995005 (GB_GCA_001995005.1)
Epulopiscium_C sp002007315 (GB_GCA_002007315.1)
Epulopiscium_C sp001994995 (GB_GCA_001994995.1)
Epulopiscium_D sp002007365 (GB_GCA_002007365.1)
Epulopiscium_A sp002007285 (GB_GCA_002007285.1)
Epulopiscium_A sp001983475 (GB_GCA_001983475.1)
Epulopiscium_A sp001983455 (GB_GCA_001983455.1)
Cellulosilyticum ruminicola (RS_GCF_001311925.1)
Cellulosilyticum lentocellum (RS_GCF_000178835.2)
Cellulosilyticum sp002735925 (RS_GCF_002735925.1)
Cellulosilyticum sp900095725 (RS_GCF_900095725.1)
Niameybacter massiliensis (metabat.458)
Niameybacter massiliensis (RS_GCF_001282685.1)
Defluviitalea phaphyphila (RS_GCF_001466305.1)

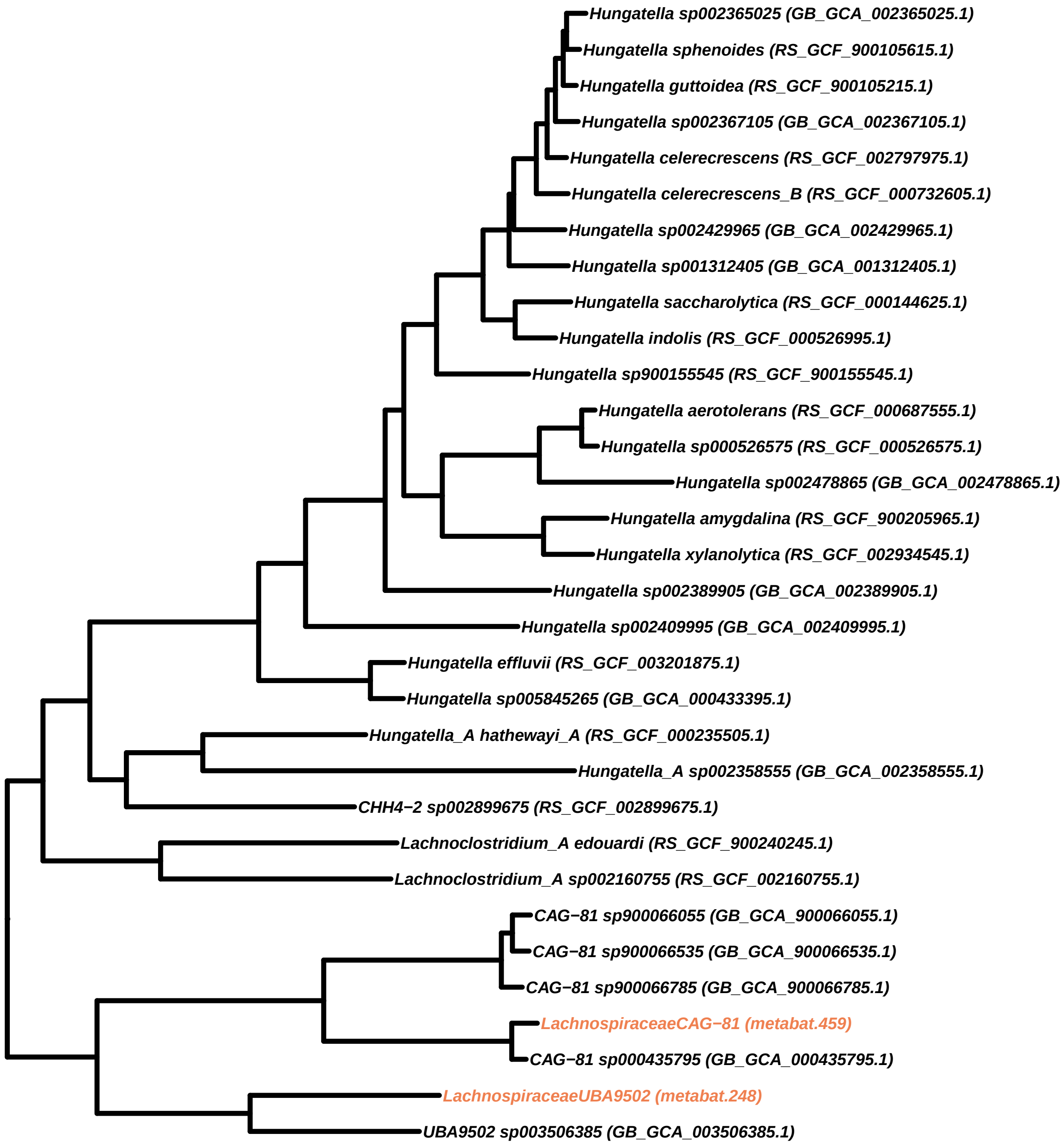
Hungatella sp002365025 (GB_GCA_002365025.1)
Hungatella sphenoides (RS_GCF_900105615.1)
Hungatella guttoidea (RS_GCF_900105215.1)
Hungatella sp002367105 (GB_GCA_002367105.1)
Hungatella celerecrescens (RS_GCF_002797975.1)
Hungatella celerecrescens_B (RS_GCF_000732605.1)
Hungatella sp002429965 (GB_GCA_002429965.1)
Hungatella sp001312405 (GB_GCA_001312405.1)
Hungatella saccharolytica (RS_GCF_000144625.1)
Hungatella indolis (RS_GCF_000526995.1)
Hungatella sp900155545 (RS_GCF_900155545.1)
Hungatella aerotolerans (RS_GCF_000687555.1)
Hungatella sp000526575 (RS_GCF_000526575.1)
Hungatella sp002478865 (GB_GCA_002478865.1)
Hungatella amygdalina (RS_GCF_900205965.1)
Hungatella xylanolytica (RS_GCF_002934545.1)
Hungatella sp002389905 (GB_GCA_002389905.1)
Hungatella sp002409995 (GB_GCA_002409995.1)
Hungatella effluvii (RS_GCF_003201875.1)
Hungatella sp005845265 (GB_GCA_000433395.1)
Hungatella_A hathewayi_A (RS_GCF_000235505.1)
Hungatella_A sp002358555 (GB_GCA_002358555.1)
CHH4−2 sp002899675 (RS_GCF_002899675.1)
Lachnoclostridium_A edouardi (RS_GCF_900240245.1)
Lachnoclostridium_A sp002160755 (RS_GCF_002160755.1)
CAG−81 sp900066055 (GB_GCA_900066055.1)
CAG−81 sp900066535 (GB_GCA_900066535.1)
CAG−81 sp900066785 (GB_GCA_900066785.1)
LachnospiraceaeCAG−81 (metabat.459)
CAG−81 sp000435795 (GB_GCA_000435795.1)
LachnospiraceaeUBA9502 (metabat.248)
UBA9502 sp003506385 (GB_GCA_003506385.1)

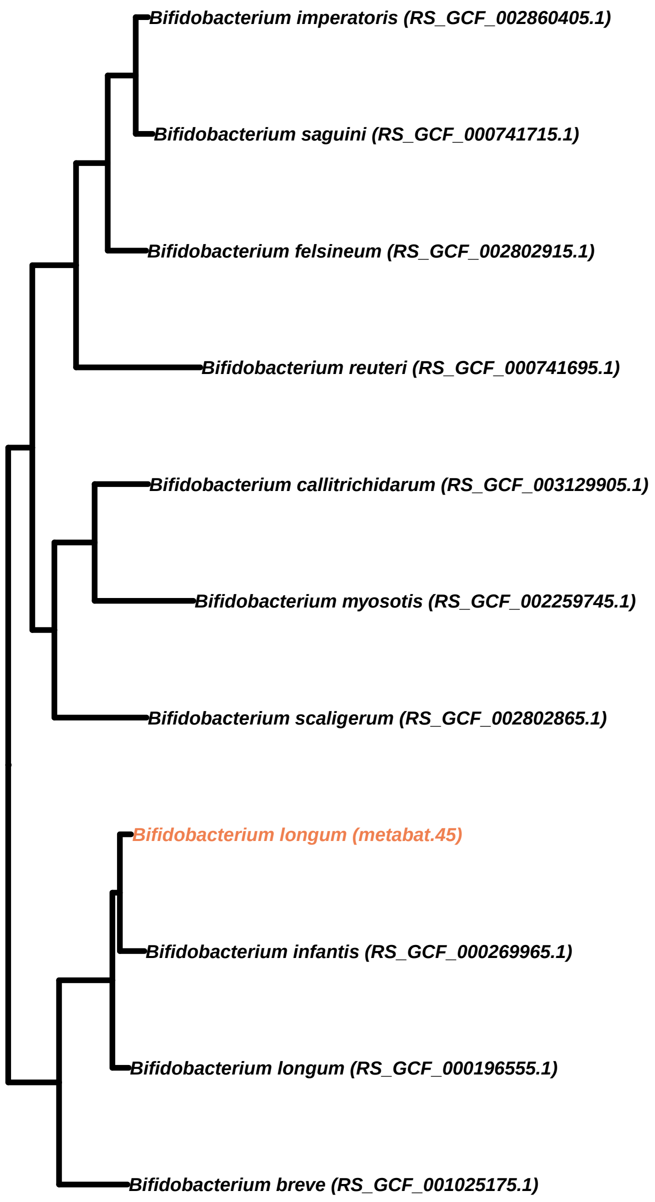
Bifidobacterium imperatoris (RS_GCF_002860405.1)
Bifidobacterium saguini (RS_GCF_000741715.1)
Bifidobacterium felsineum (RS_GCF_002802915.1)
Bifidobacterium reuteri (RS_GCF_000741695.1)
Bifidobacterium callitrichidarum (RS_GCF_003129905.1)
Bifidobacterium myosotis (RS_GCF_002259745.1)
Bifidobacterium scaligerum (RS_GCF_002802865.1)
Bifidobacterium longum (metabat.45)
Bifidobacterium infantis (RS_GCF_000269965.1)
Bifidobacterium longum (RS_GCF_000196555.1)
Bifidobacterium breve (RS_GCF_001025175.1)

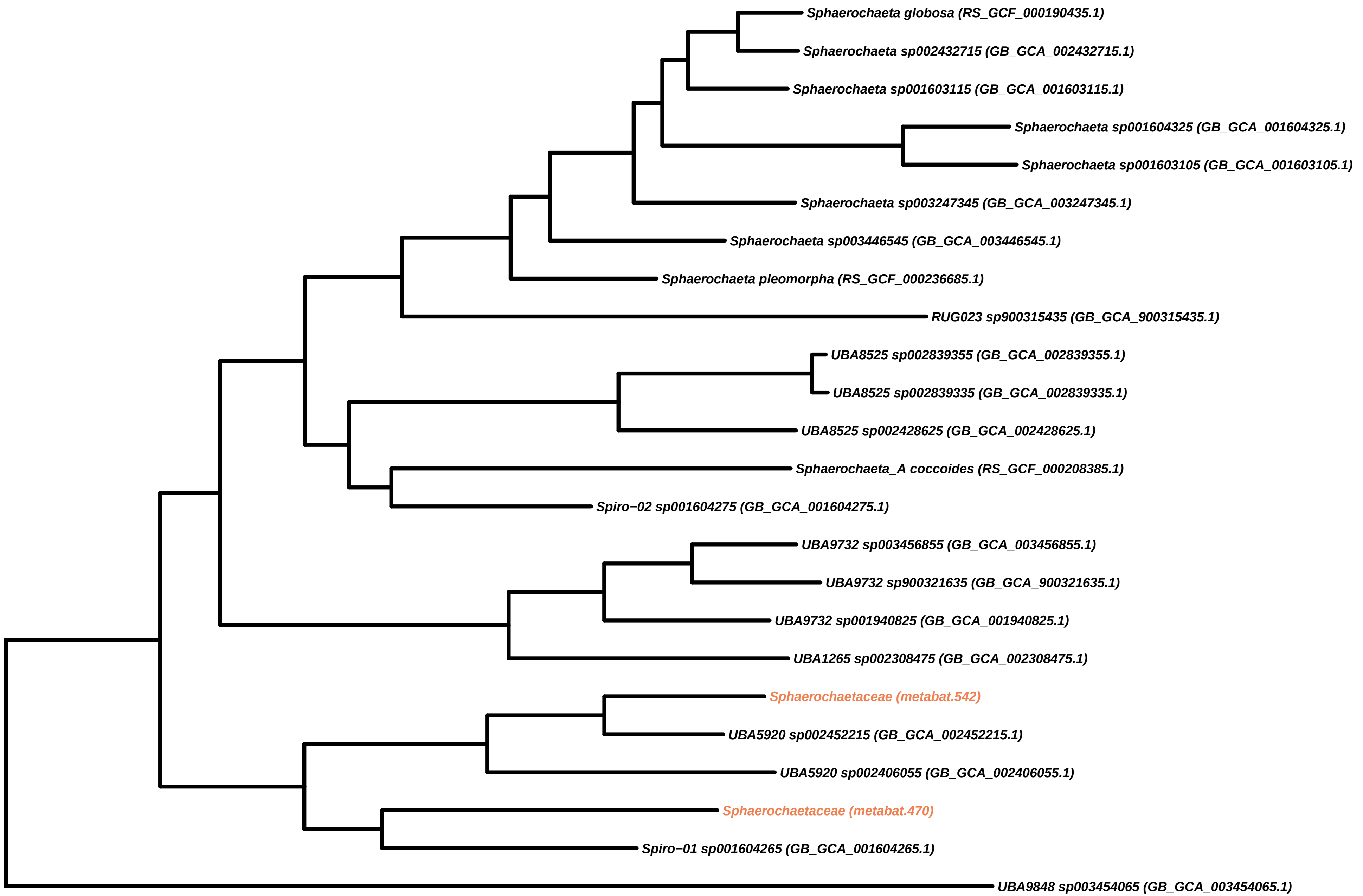
Sphaerochaeta globosa (RS_GCF_000190435.1)
Sphaerochaeta sp002432715 (GB_GCA_002432715.1)
Sphaerochaeta sp001603115 (GB_GCA_001603115.1)
Sphaerochaeta sp001604325 (GB_GCA_001604325.1)
Sphaerochaeta sp001603105 (GB_GCA_001603105.1)
Sphaerochaeta sp003247345 (GB_GCA_003247345.1)
Sphaerochaeta sp003446545 (GB_GCA_003446545.1)
Sphaerochaeta pleomorpha (RS_GCF_000236685.1)
RUG023 sp900315435 (GB_GCA_900315435.1)
UBA8525 sp002839355 (GB_GCA_002839355.1)
UBA8525 sp002839335 (GB_GCA_002839335.1)
UBA8525 sp002428625 (GB_GCA_002428625.1)
Sphaerochaeta_A coccoides (RS_GCF_000208385.1)
Spiro−02 sp001604275 (GB_GCA_001604275.1)
UBA9732 sp003456855 (GB_GCA_003456855.1)
UBA9732 sp900321635 (GB_GCA_900321635.1)
UBA9732 sp001940825 (GB_GCA_001940825.1)
UBA1265 sp002308475 (GB_GCA_002308475.1)
Sphaerochaetaceae (metabat.542)
UBA5920 sp002452215 (GB_GCA_002452215.1)
UBA5920 sp002406055 (GB_GCA_002406055.1)
Sphaerochaetaceae (metabat.470)
Spiro−01 sp001604265 (GB_GCA_001604265.1)
UBA9848 sp003454065 (GB_GCA_003454065.1)

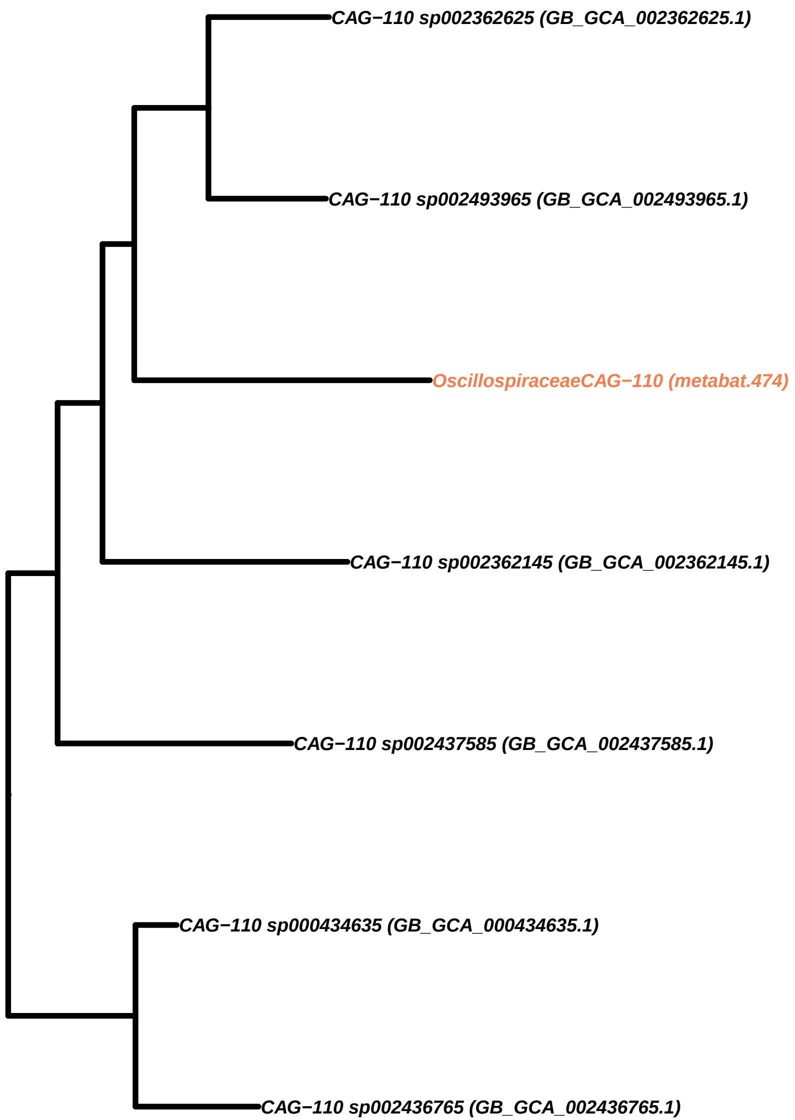
CAG−110 sp002362625 (GB_GCA_002362625.1)
CAG−110 sp002493965 (GB_GCA_002493965.1)
OscillospiraceaeCAG−110 (metabat.474)
CAG−110 sp002362145 (GB_GCA_002362145.1)
CAG−110 sp002437585 (GB_GCA_002437585.1)
CAG−110 sp000434635 (GB_GCA_000434635.1)
CAG−110 sp002436765 (GB_GCA_002436765.1)

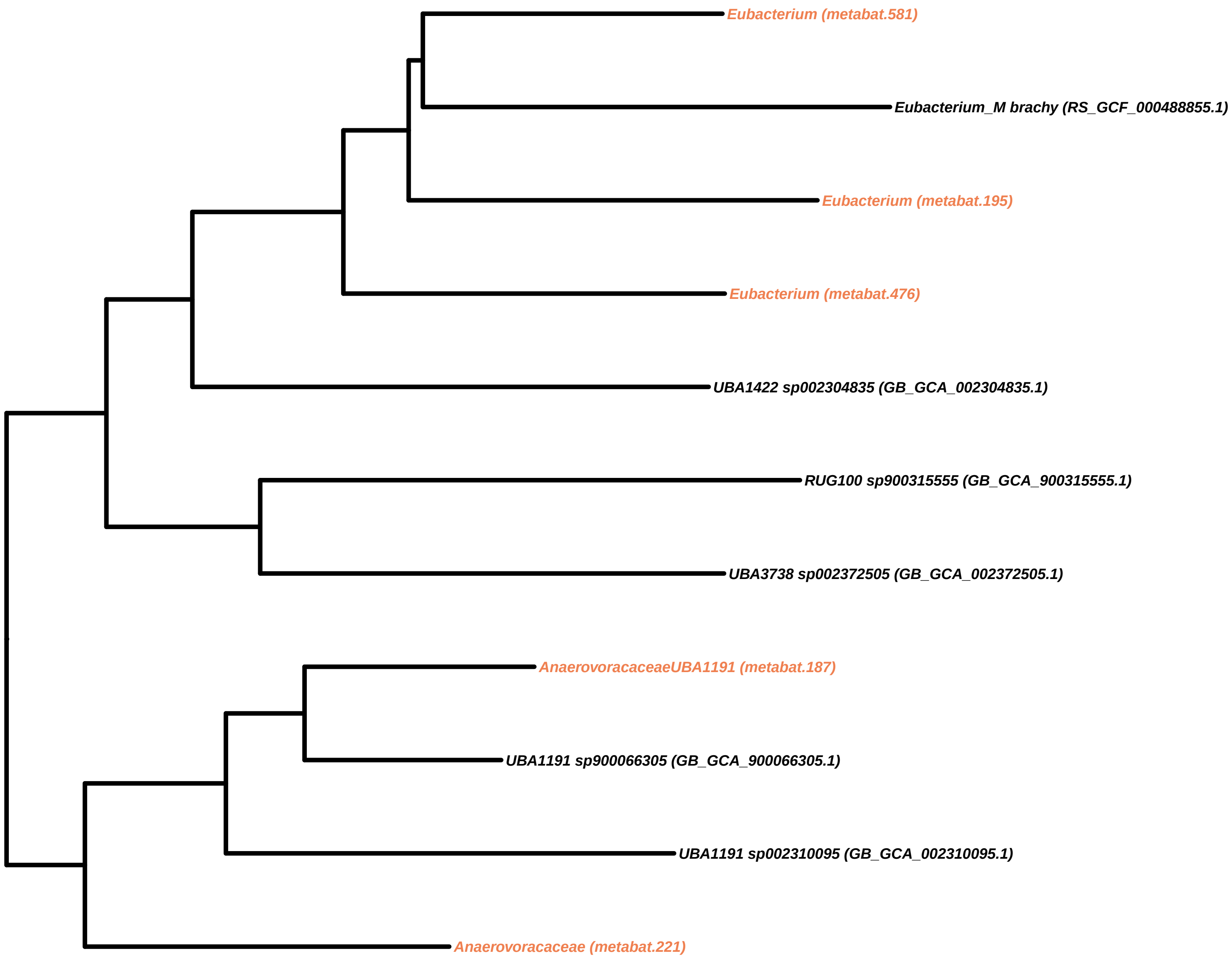

Eubacterium (metabat.581)
Eubacterium_M brachy (RS_GCF_000488855.1)
Eubacterium (metabat.195)
Eubacterium (metabat.476)
UBA1422 sp002304835 (GB_GCA_002304835.1)
RUG100 sp900315555 (GB_GCA_900315555.1)
UBA3738 sp002372505 (GB_GCA_002372505.1)
AnaerovoracaceaeUBA1191 (metabat.187)
UBA1191 sp900066305 (GB_GCA_900066305.1)
UBA1191 sp002310095 (GB_GCA_002310095.1)
Anaerovoracaceae (metabat.221)

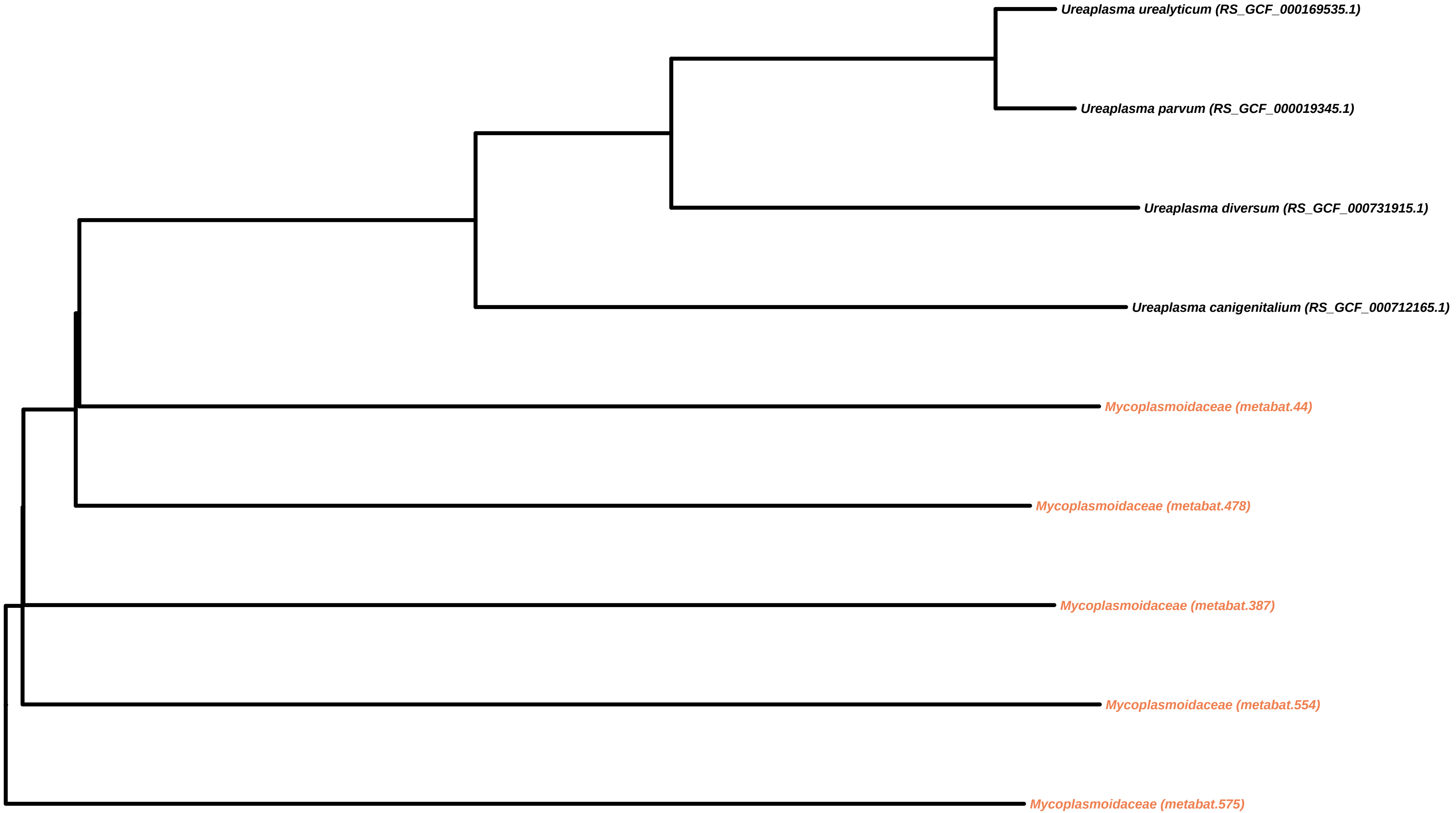
Ureaplasma urealyticum (RS_GCF_000169535.1)
Ureaplasma parvum (RS_GCF_000019345.1)
Ureaplasma diversum (RS_GCF_000731915.1)
Ureaplasma canigenitalium (RS_GCF_000712165.1)
Mycoplasmoidaceae (metabat.44)
Mycoplasmoidaceae (metabat.478)
Mycoplasmoidaceae (metabat.387)
Mycoplasmoidaceae (metabat.554)
Mycoplasmoidaceae (metabat.575)

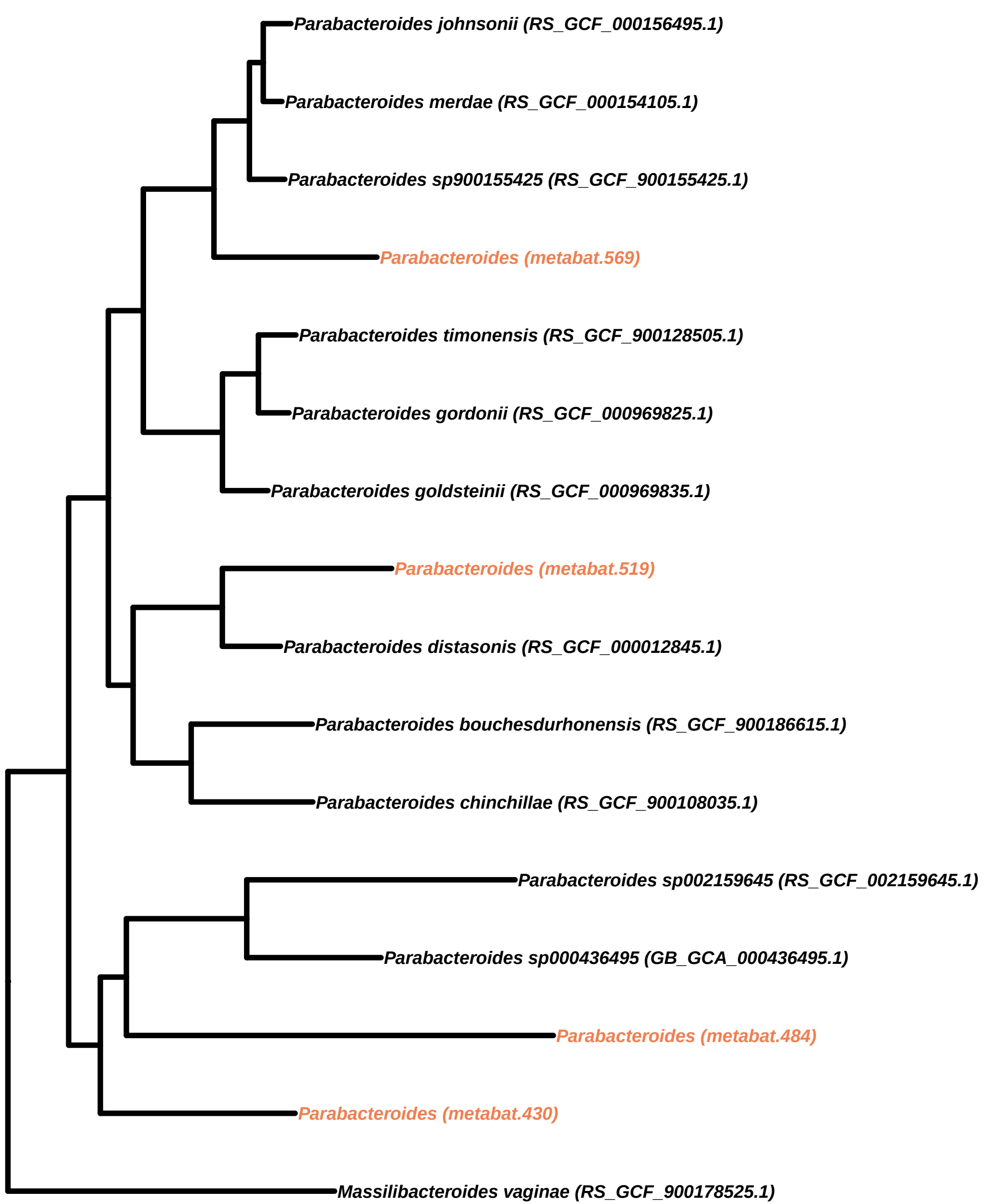
Parabacteroides johnsonii (RS_GCF_000156495.1)
Parabacteroides merdae (RS_GCF_000154105.1)
Parabacteroides sp900155425 (RS_GCF_900155425.1)
Parabacteroides (metabat.569)
Parabacteroides timonensis (RS_GCF_900128505.1)
Parabacteroides gordonii (RS_GCF_000969825.1)
Parabacteroides goldsteinii (RS_GCF_000969835.1)
Parabacteroides (metabat.519)
Parabacteroides distasonis (RS_GCF_000012845.1)
Parabacteroides bouchesdurhonensis (RS_GCF_900186615.1)
Parabacteroides chinchillae (RS_GCF_900108035.1)
Parabacteroides sp002159645 (RS_GCF_002159645.1)
Parabacteroides sp000436495 (GB_GCA_000436495.1)
Parabacteroides (metabat.484)
Parabacteroides (metabat.430)
Massilibacteroides vaginae (RS_GCF_900178525.1)

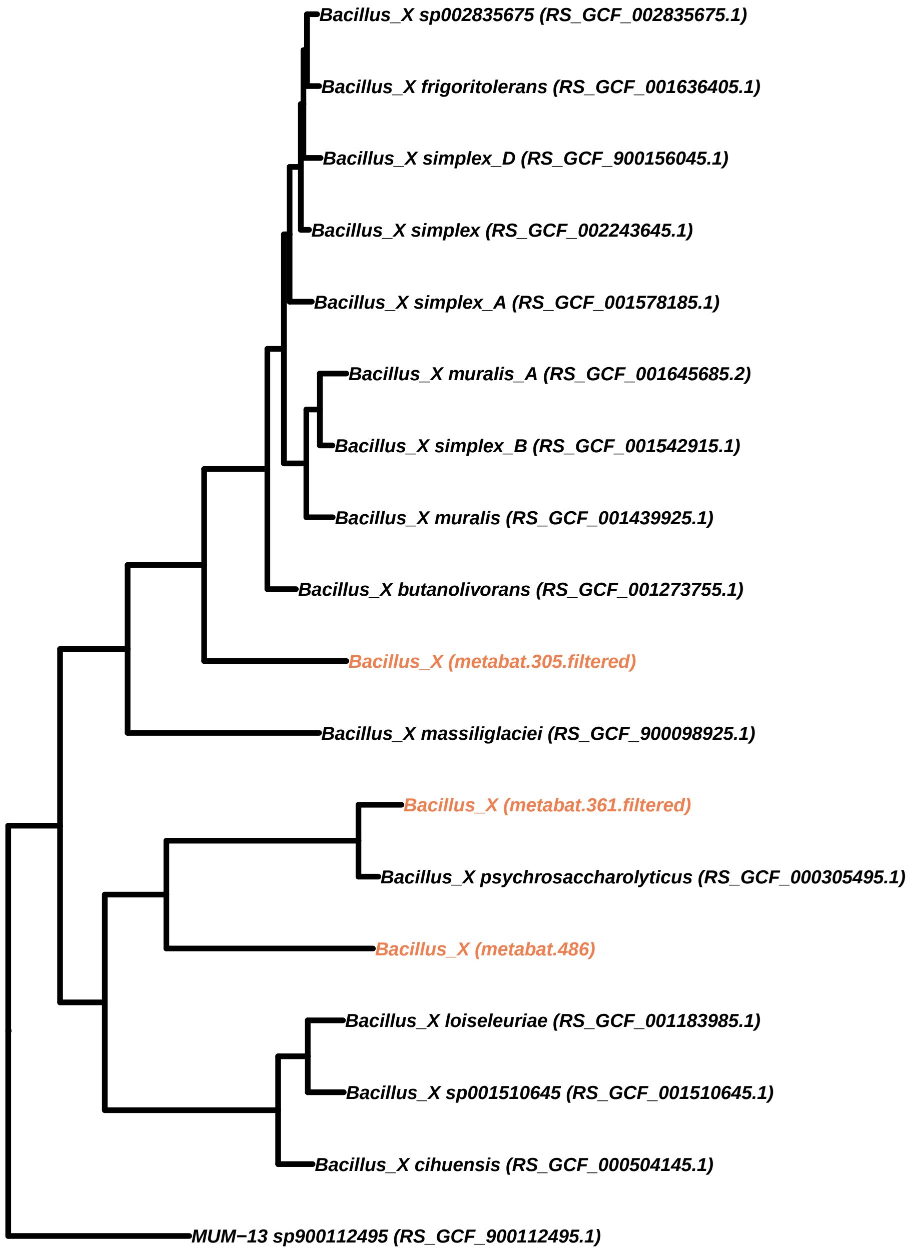
Bacillus_X sp002835675 (RS_GCF_002835675.1)
Bacillus_X frigoritolerans (RS_GCF_001636405.1)
Bacillus_X simplex_D (RS_GCF_900156045.1)
Bacillus_X simplex (RS_GCF_002243645.1)
Bacillus_X simplex_A (RS_GCF_001578185.1)
Bacillus_X muralis_A (RS_GCF_001645685.2)
Bacillus_X simplex_B (RS_GCF_001542915.1)
Bacillus_X muralis (RS_GCF_001439925.1)
Bacillus_X butanolivorans (RS_GCF_001273755.1)
Bacillus_X (metabat.305.filtered)
Bacillus_X massiliglaciei (RS_GCF_900098925.1)
Bacillus_X (metabat.361.filtered)
Bacillus_X psychrosaccharolyticus (RS_GCF_000305495.1)
Bacillus_X (metabat.486)
Bacillus_X loiseleuriae (RS_GCF_001183985.1)
Bacillus_X sp001510645 (RS_GCF_001510645.1)
Bacillus_X cihuensis (RS_GCF_000504145.1)
MUM−13 sp900112495 (RS_GCF_900112495.1)

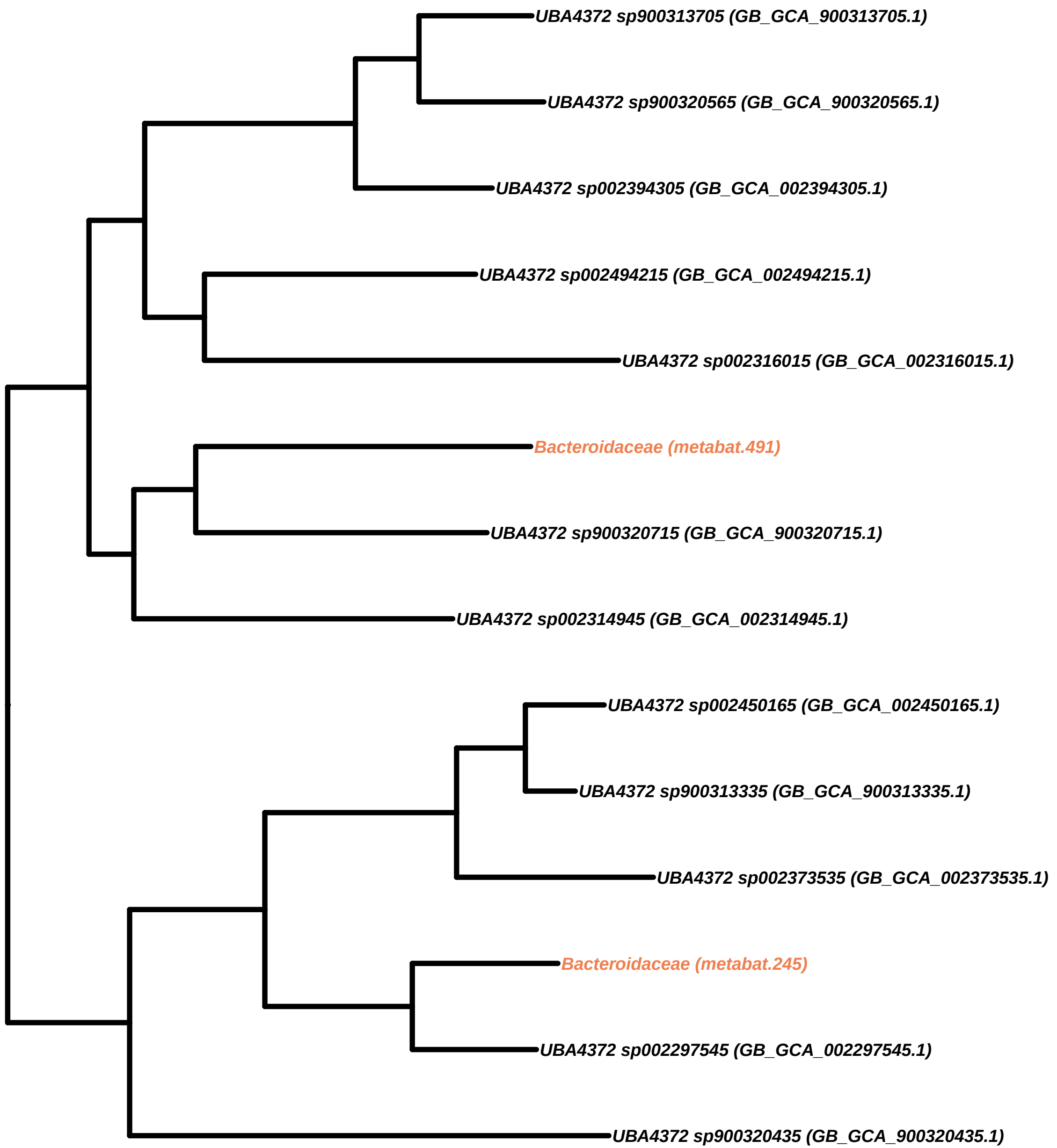
UBA4372 sp900313705 (GB_GCA_900313705.1)
UBA4372 sp900320565 (GB_GCA_900320565.1)
UBA4372 sp002394305 (GB_GCA_002394305.1)
UBA4372 sp002494215 (GB_GCA_002494215.1)
UBA4372 sp002316015 (GB_GCA_002316015.1)
Bacteroidaceae (metabat.491)
UBA4372 sp900320715 (GB_GCA_900320715.1)
UBA4372 sp002314945 (GB_GCA_002314945.1)
UBA4372 sp002450165 (GB_GCA_002450165.1)
UBA4372 sp900313335 (GB_GCA_900313335.1)
UBA4372 sp002373535 (GB_GCA_002373535.1)
Bacteroidaceae (metabat.245)
UBA4372 sp002297545 (GB_GCA_002297545.1)
UBA4372 sp900320435 (GB_GCA_900320435.1)

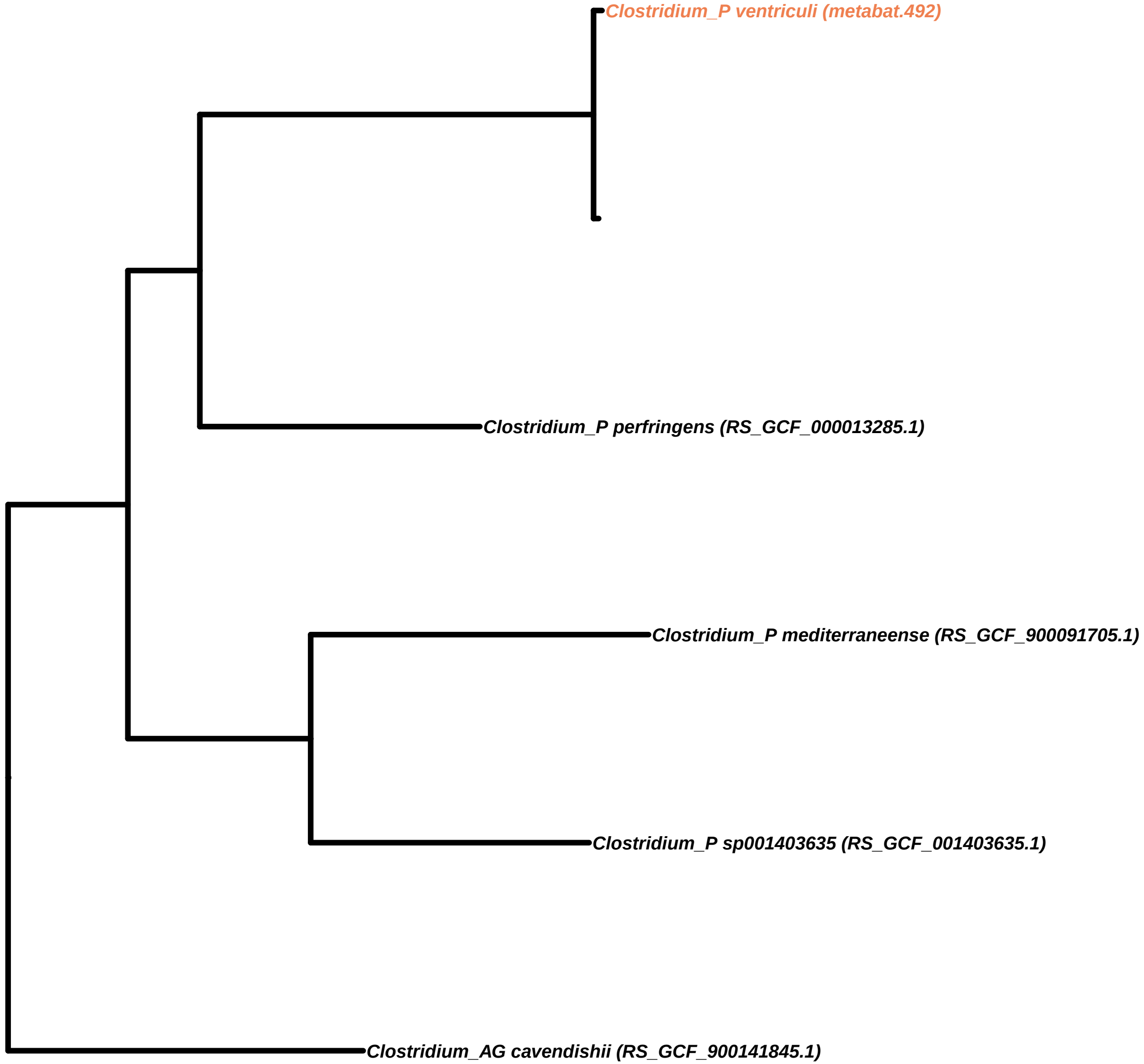

Clostridium_P ventriculi (metabat.492)
Clostridium_P perfringens (RS_GCF_000013285.1)
Clostridium_P mediterraneense (RS_GCF_900091705.1)
Clostridium_P sp001403635 (RS_GCF_001403635.1)
Clostridium_AG cavendishii (RS_GCF_900141845.1)

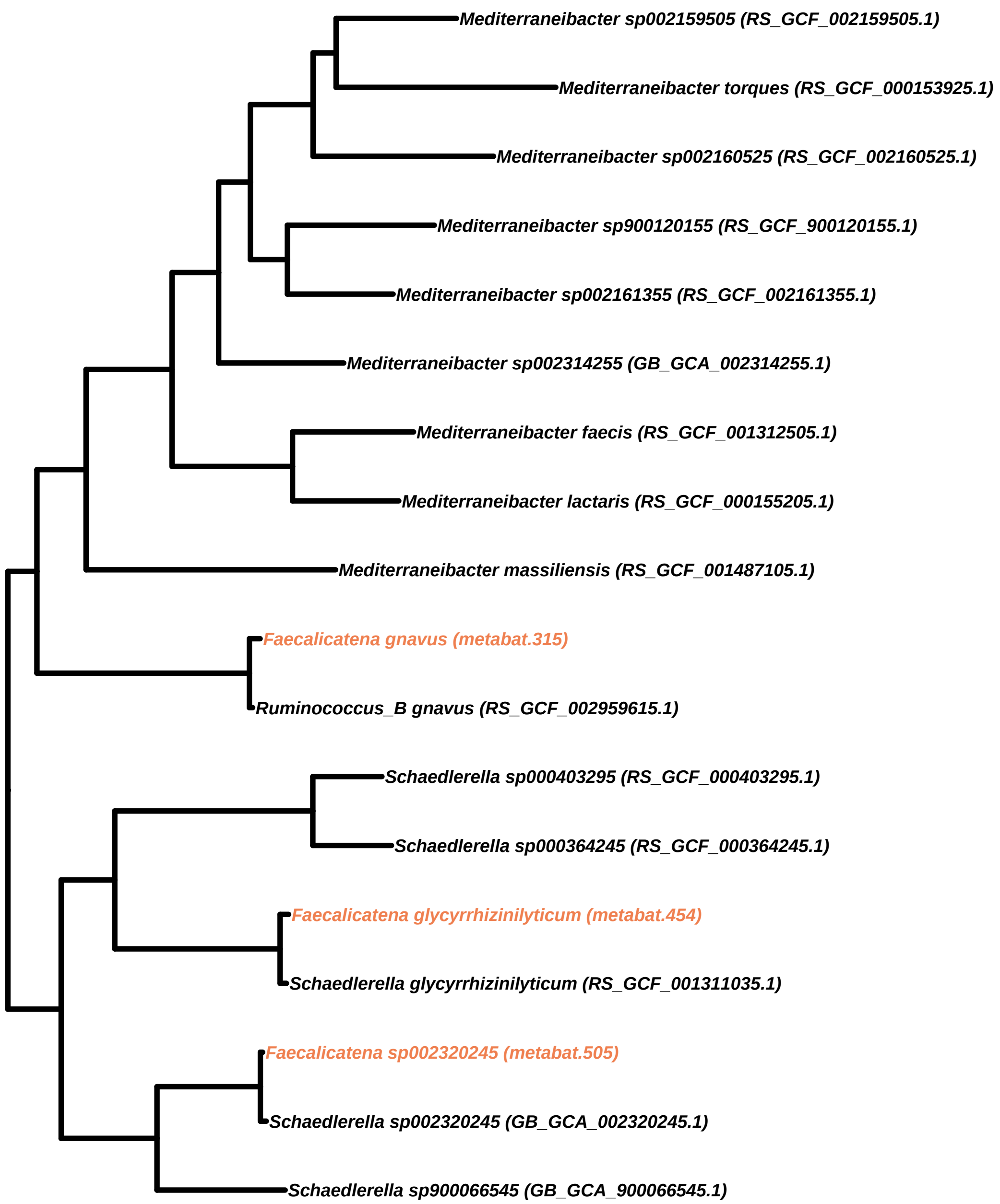

Mediterraneibacter sp002159505 (RS_GCF_002159505.1)
Mediterraneibacter torques (RS_GCF_000153925.1)
Mediterraneibacter sp002160525 (RS_GCF_002160525.1)
Mediterraneibacter sp900120155 (RS_GCF_900120155.1)
Mediterraneibacter sp002161355 (RS_GCF_002161355.1)
Mediterraneibacter sp002314255 (GB_GCA_002314255.1)
Mediterraneibacter faecis (RS_GCF_001312505.1)
Mediterraneibacter lactaris (RS_GCF_000155205.1)
Mediterraneibacter massiliensis (RS_GCF_001487105.1)
Faecalicatena gnavus (metabat.315)
Ruminococcus_B gnavus (RS_GCF_002959615.1)
Schaedlerella sp000403295 (RS_GCF_000403295.1)
Schaedlerella sp000364245 (RS_GCF_000364245.1)
Faecalicatena glycyrrhizinilyticum (metabat.454)
Schaedlerella glycyrrhizinilyticum (RS_GCF_001311035.1)
Faecalicatena sp002320245 (metabat.505)
Schaedlerella sp002320245 (GB_GCA_002320245.1)
Schaedlerella sp900066545 (GB_GCA_900066545.1)

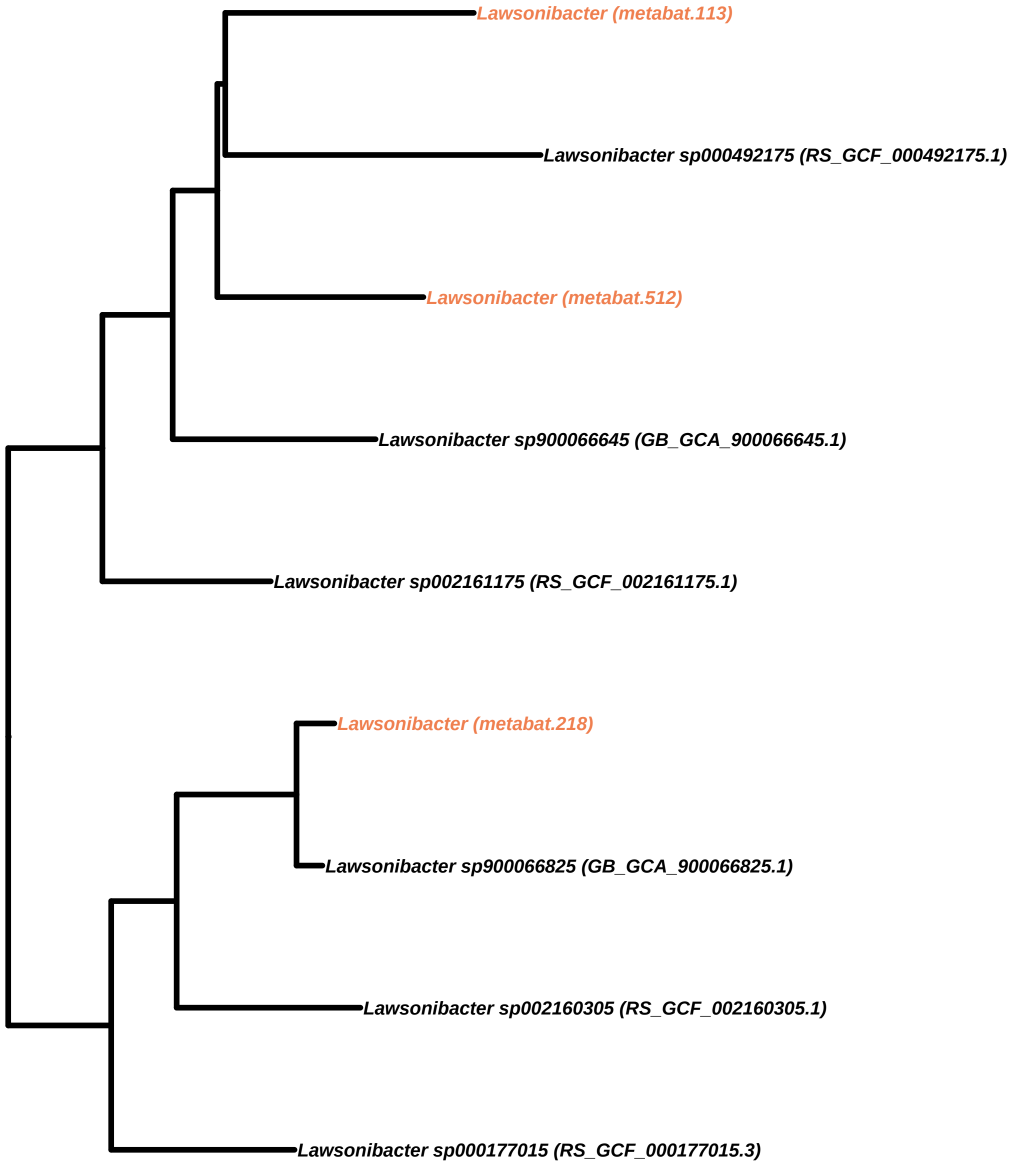

Lawsonibacter (metabat.113)
Lawsonibacter sp000492175 (RS_GCF_000492175.1)
Lawsonibacter (metabat.512)
Lawsonibacter sp900066645 (GB_GCA_900066645.1)
Lawsonibacter sp002161175 (RS_GCF_002161175.1)
Lawsonibacter (metabat.218)
Lawsonibacter sp900066825 (GB_GCA_900066825.1)
Lawsonibacter sp002160305 (RS_GCF_002160305.1)
Lawsonibacter sp000177015 (RS_GCF_000177015.3)

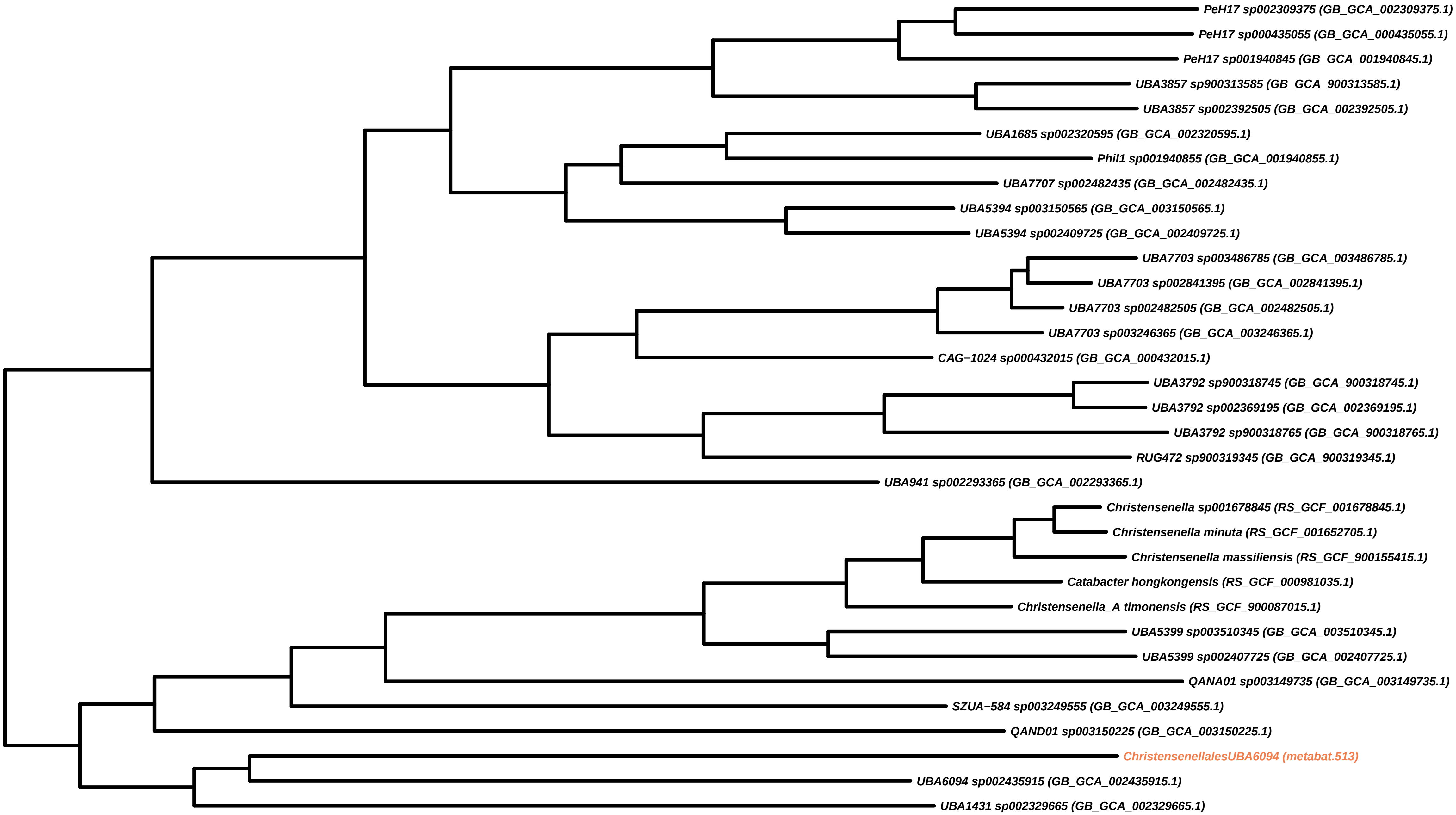
PeH17 sp002309375 (GB_GCA_002309375.1)
PeH17 sp000435055 (GB_GCA_000435055.1)
PeH17 sp001940845 (GB_GCA_001940845.1)
UBA3857 sp900313585 (GB_GCA_900313585.1)
UBA3857 sp002392505 (GB_GCA_002392505.1)
UBA1685 sp002320595 (GB_GCA_002320595.1)
Phil1 sp001940855 (GB_GCA_001940855.1)
UBA7707 sp002482435 (GB_GCA_002482435.1)
UBA5394 sp003150565 (GB_GCA_003150565.1)
UBA5394 sp002409725 (GB_GCA_002409725.1)
UBA7703 sp003486785 (GB_GCA_003486785.1)
UBA7703 sp002841395 (GB_GCA_002841395.1)
UBA7703 sp002482505 (GB_GCA_002482505.1)
UBA7703 sp003246365 (GB_GCA_003246365.1)
CAG−1024 sp000432015 (GB_GCA_000432015.1)
UBA3792 sp900318745 (GB_GCA_900318745.1)
UBA3792 sp002369195 (GB_GCA_002369195.1)
UBA3792 sp900318765 (GB_GCA_900318765.1)
RUG472 sp900319345 (GB_GCA_900319345.1)
UBA941 sp002293365 (GB_GCA_002293365.1)
Christensenella sp001678845 (RS_GCF_001678845.1)
Christensenella minuta (RS_GCF_001652705.1)
Christensenella massiliensis (RS_GCF_900155415.1)
Catabacter hongkongensis (RS_GCF_000981035.1)
Christensenella_A timonensis (RS_GCF_900087015.1)
UBA5399 sp003510345 (GB_GCA_003510345.1)
UBA5399 sp002407725 (GB_GCA_002407725.1)
QANA01 sp003149735 (GB_GCA_003149735.1)
SZUA−584 sp003249555 (GB_GCA_003249555.1)
QAND01 sp003150225 (GB_GCA_003150225.1)
ChristensenellalesUBA6094 (metabat.513)
UBA6094 sp002435915 (GB_GCA_002435915.1)
UBA1431 sp002329665 (GB_GCA_002329665.1)

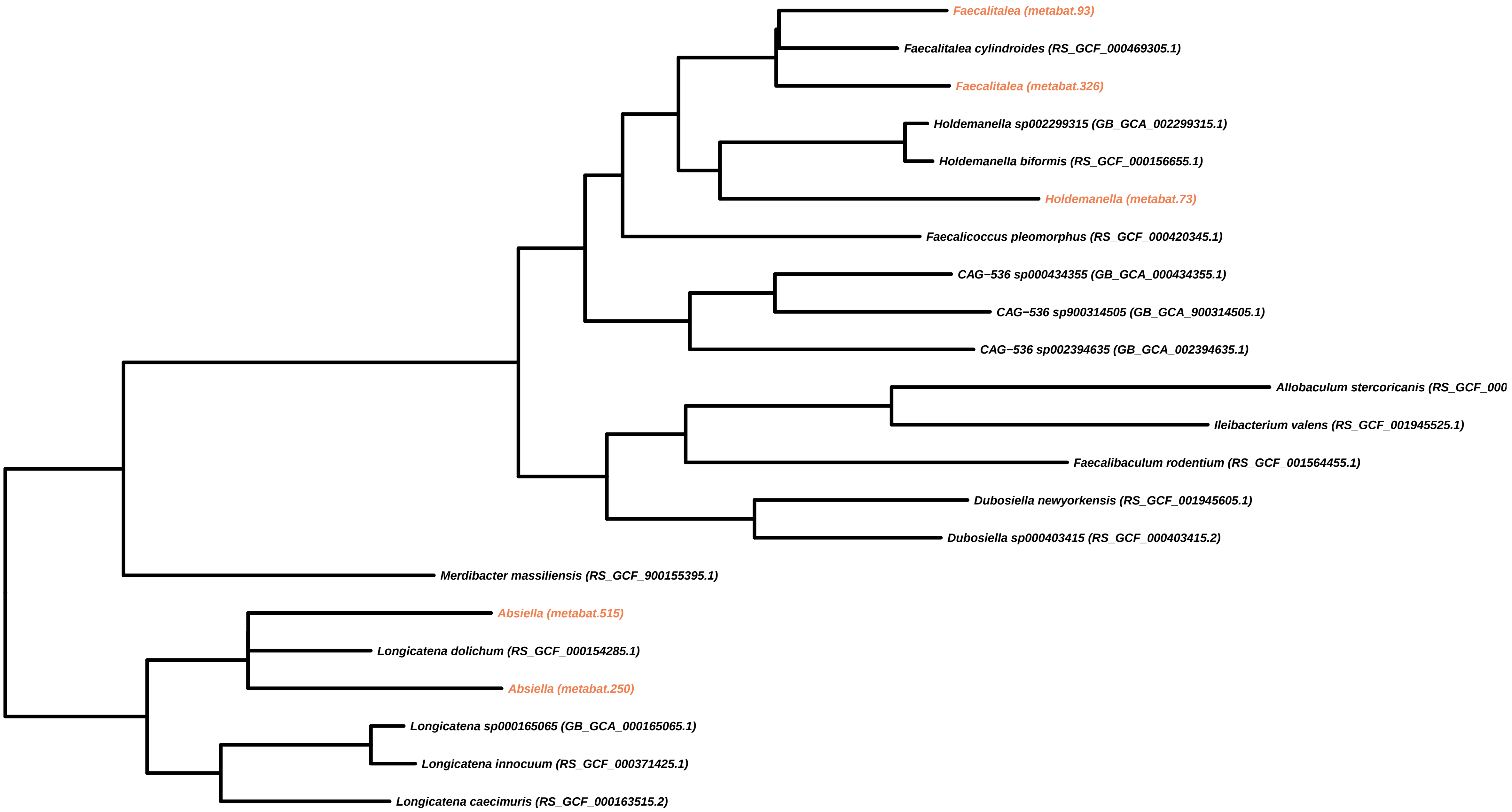
Faecalitalea (metabat.93)
Faecalitalea cylindroides (RS_GCF_000469305.1)
Faecalitalea (metabat.326)
Holdemanella sp002299315 (GB_GCA_002299315.1)
Holdemanella biformis (RS_GCF_000156655.1)
Holdemanella (metabat.73)
Faecalicoccus pleomorphus (RS_GCF_000420345.1)
CAG−536 sp000434355 (GB_GCA_000434355.1)
CAG−536 sp900314505 (GB_GCA_900314505.1)
CAG−536 sp002394635 (GB_GCA_002394635.1)
Allobaculum stercoricanis (RS_GCF_000
Ileibacterium valens (RS_GCF_001945525.1)
Faecalibaculum rodentium (RS_GCF_001564455.1)
Dubosiella newyorkensis (RS_GCF_001945605.1)
Dubosiella sp000403415 (RS_GCF_000403415.2)
Merdibacter massiliensis (RS_GCF_900155395.1)
Absiella (metabat.515)
Longicatena dolichum (RS_GCF_000154285.1)
Absiella (metabat.250)
Longicatena sp000165065 (GB_GCA_000165065.1)
Longicatena innocuum (RS_GCF_000371425.1)
Longicatena caecimuris (RS_GCF_000163515.2)

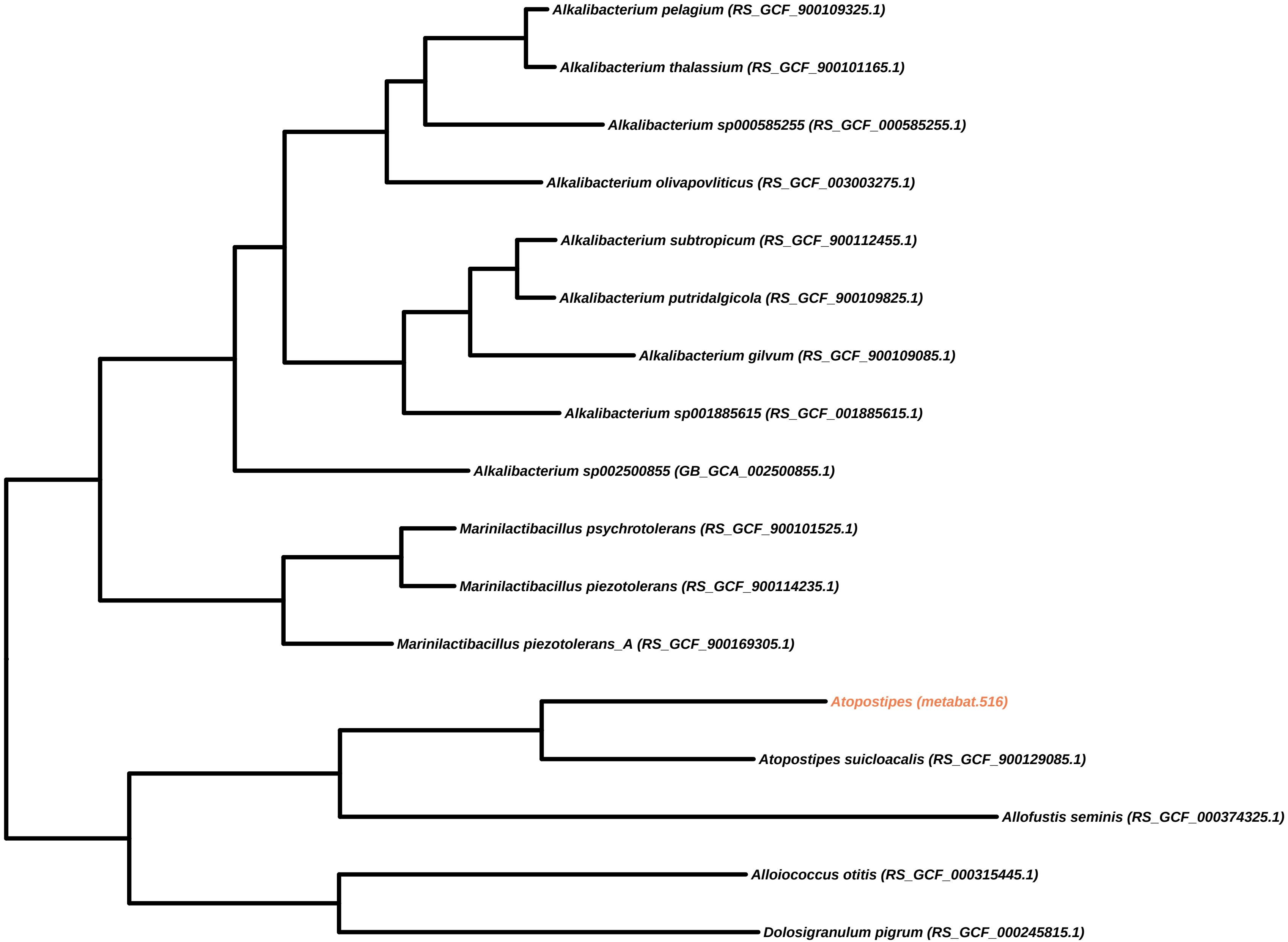

Alkalibacterium pelagium (RS_GCF_900109325.1)
Alkalibacterium thalassium (RS_GCF_900101165.1)
Alkalibacterium sp000585255 (RS_GCF_000585255.1)
Alkalibacterium olivapovliticus (RS_GCF_003003275.1)
Alkalibacterium subtropicum (RS_GCF_900112455.1)
Alkalibacterium putridalgicola (RS_GCF_900109825.1)
Alkalibacterium gilvum (RS_GCF_900109085.1)
Alkalibacterium sp001885615 (RS_GCF_001885615.1)
Alkalibacterium sp002500855 (GB_GCA_002500855.1)
Marinilactibacillus psychrotolerans (RS_GCF_900101525.1)
Marinilactibacillus piezotolerans (RS_GCF_900114235.1)
Marinilactibacillus piezotolerans_A (RS_GCF_900169305.1)
Atopostipes (metabat.516)
Atopostipes suicloacalis (RS_GCF_900129085.1)
Allofustis seminis (RS_GCF_000374325.1)
Alloiococcus otitis (RS_GCF_000315445.1)
Dolosigranulum pigrum (RS_GCF_000245815.1)

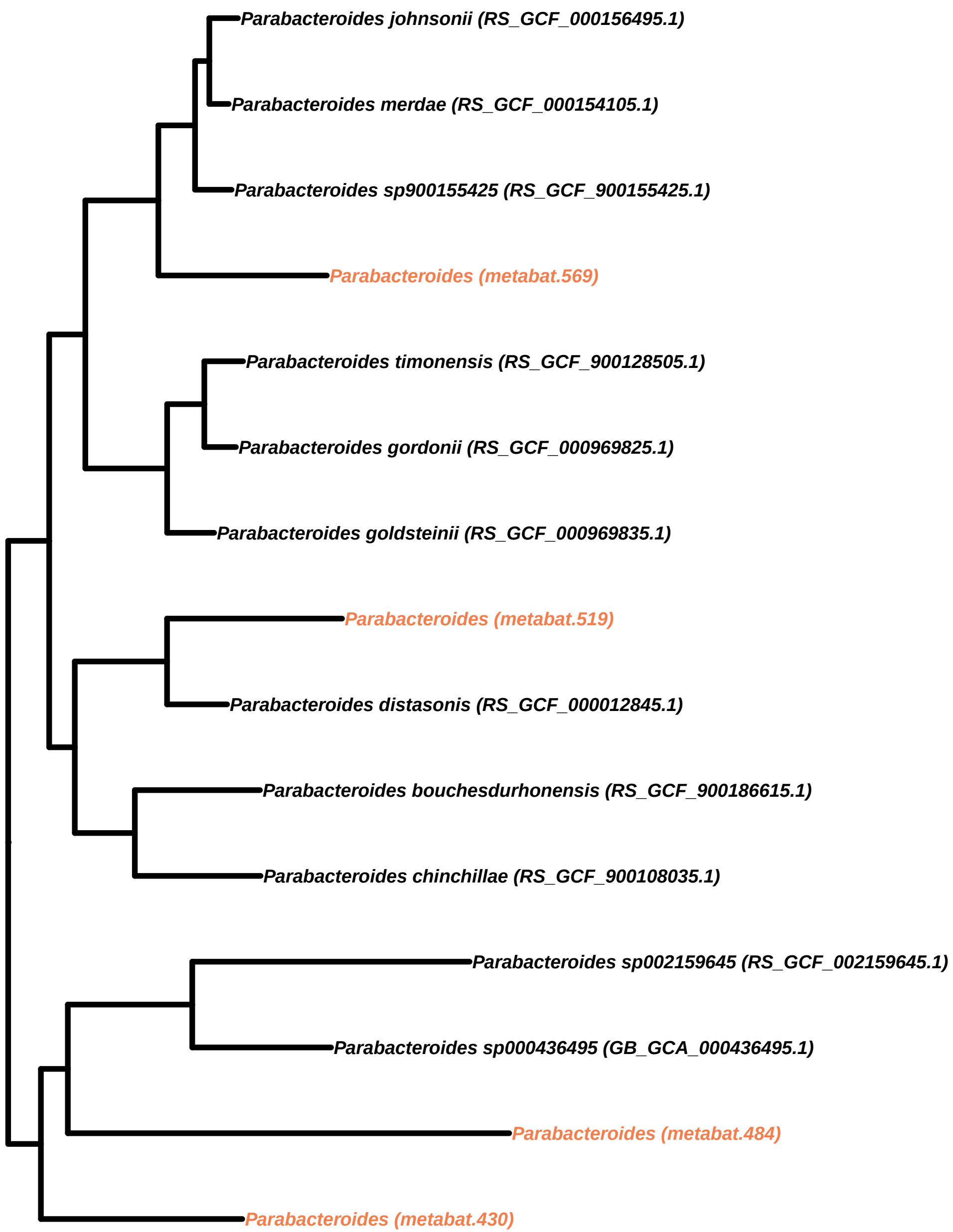
Parabacteroides johnsonii (RS_GCF_000156495.1)
Parabacteroides merdae (RS_GCF_000154105.1)
Parabacteroides sp900155425 (RS_GCF_900155425.1)
Parabacteroides (metabat.569)
Parabacteroides timonensis (RS_GCF_900128505.1)
Parabacteroides gordonii (RS_GCF_000969825.1)
Parabacteroides goldsteinii (RS_GCF_000969835.1)
Parabacteroides (metabat.519)
Parabacteroides distasonis (RS_GCF_000012845.1)
Parabacteroides bouchesdurhonensis (RS_GCF_900186615.1)
Parabacteroides chinchillae (RS_GCF_900108035.1)
Parabacteroides sp002159645 (RS_GCF_002159645.1)
Parabacteroides sp000436495 (GB_GCA_000436495.1)
Parabacteroides (metabat.484)
Parabacteroides (metabat.430)

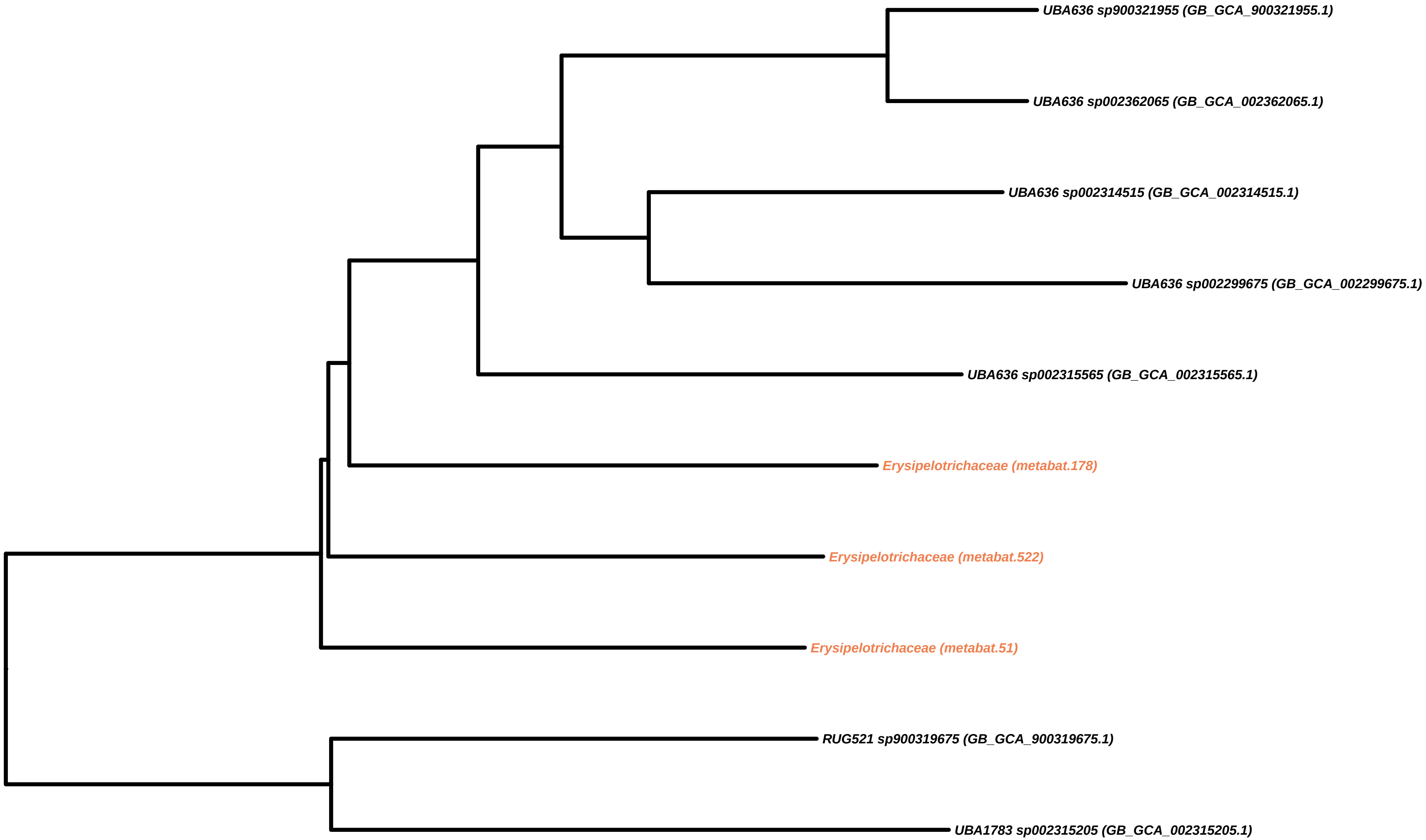
UBA636 sp900321955 (GB_GCA_900321955.1)
UBA636 sp002362065 (GB_GCA_002362065.1)
UBA636 sp002314515 (GB_GCA_002314515.1)
UBA636 sp002299675 (GB_GCA_002299675.1)
UBA636 sp002315565 (GB_GCA_002315565.1)
Erysipelotrichaceae (metabat.178)
Erysipelotrichaceae (metabat.522)
Erysipelotrichaceae (metabat.51)
RUG521 sp900319675 (GB_GCA_900319675.1)
UBA1783 sp002315205 (GB_GCA_002315205.1)

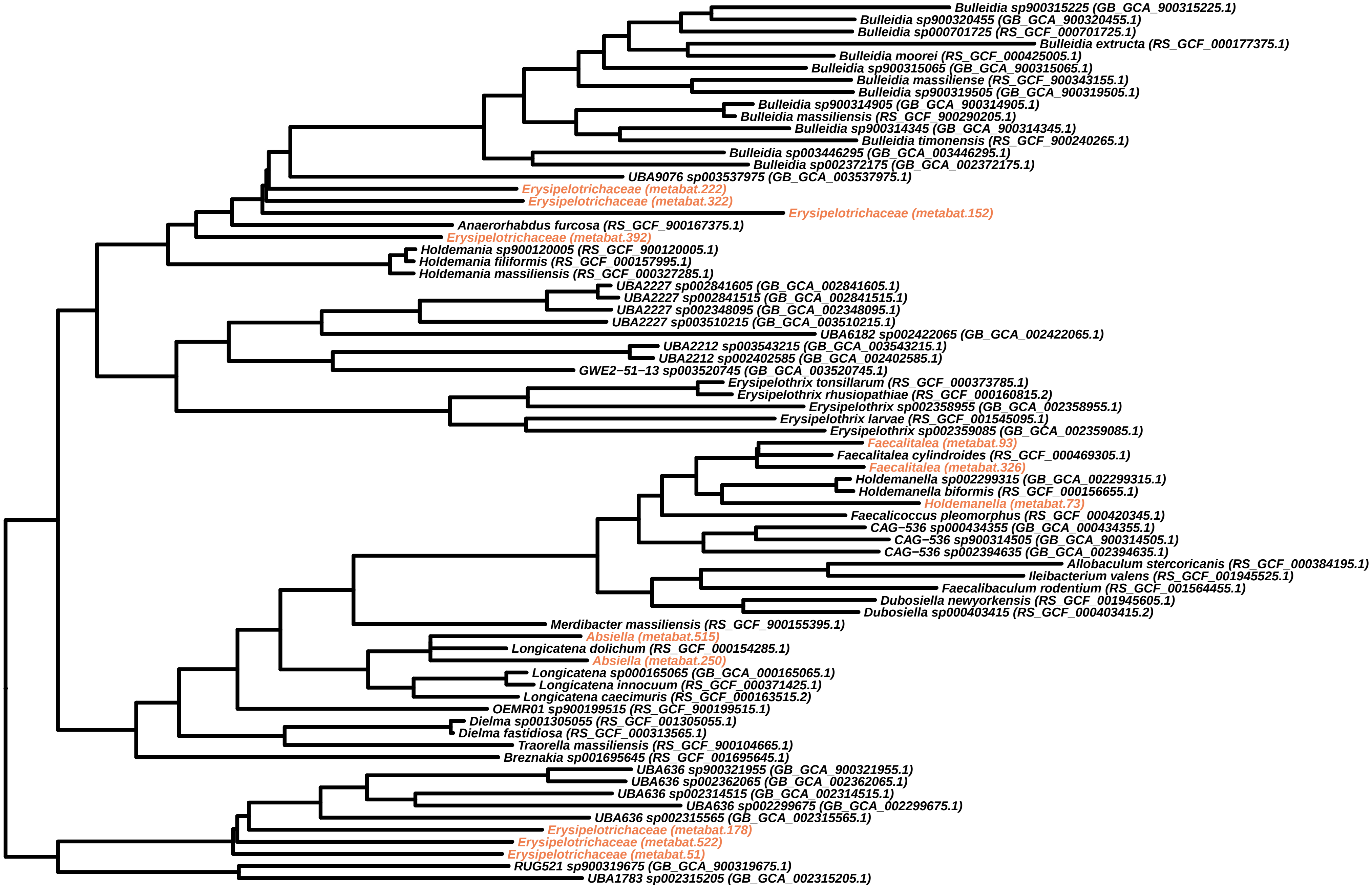
Bulleidia sp900315225 (GB_GCA_900315225.1)
Bulleidia sp900320455 (GB_GCA_900320455.1)
Bulleidia sp000701725 (RS_GCF_000701725.1)
Bulleidia extructa (RS_GCF_000177375.1)
Bulleidia moorei (RS_GCF_000425005.1)
Bulleidia sp900315065 (GB_GCA_900315065.1)
Bulleidia massiliense (RS_GCF_900343155.1)
Bulleidia sp900319505 (GB_GCA_900319505.1)
Bulleidia sp900314905 (GB_GCA_900314905.1)
Bulleidia massiliensis (RS_GCF_900290205.1)
Bulleidia sp900314345 (GB_GCA_900314345.1)
Bulleidia timonensis (RS_GCF_900240265.1)
Bulleidia sp003446295 (GB_GCA_003446295.1)
Bulleidia sp002372175 (GB_GCA_002372175.1)
UBA9076 sp003537975 (GB_GCA_003537975.1)
Erysipelotrichaceae (metabat.222)
Erysipelotrichaceae (metabat.322)
Erysipelotrichaceae (metabat.152)
Anaerorhabdus furcosa (RS_GCF_900167375.1)
Erysipelotrichaceae (metabat.392)
Holdemania sp900120005 (RS_GCF_900120005.1)
Holdemania filiformis (RS_GCF_000157995.1)
Holdemania massiliensis (RS_GCF_000327285.1)
UBA2227 sp002841605 (GB_GCA_002841605.1)
UBA2227 sp002841515 (GB_GCA_002841515.1)
UBA2227 sp002348095 (GB_GCA_002348095.1)
UBA2227 sp003510215 (GB_GCA_003510215.1)
UBA6182 sp002422065 (GB_GCA_002422065.1)
UBA2212 sp003543215 (GB_GCA_003543215.1)
UBA2212 sp002402585 (GB_GCA_002402585.1)
GWE2−51−13 sp003520745 (GB_GCA_003520745.1)
Erysipelothrix tonsillarum (RS_GCF_000373785.1)
Erysipelothrix rhusiopathiae (RS_GCF_000160815.2)
Erysipelothrix sp002358955 (GB_GCA_002358955.1)
Erysipelothrix larvae (RS_GCF_001545095.1)
Erysipelothrix sp002359085 (GB_GCA_002359085.1)
Faecalitalea (metabat.93)
Faecalitalea cylindroides (RS_GCF_000469305.1)
Faecalitalea (metabat.326)
Holdemanella sp002299315 (GB_GCA_002299315.1)
Holdemanella biformis (RS_GCF_000156655.1)
Holdemanella (metabat.73)
Faecalicoccus pleomorphus (RS_GCF_000420345.1)
CAG−536 sp000434355 (GB_GCA_000434355.1)
CAG−536 sp900314505 (GB_GCA_900314505.1)
CAG−536 sp002394635 (GB_GCA_002394635.1)
Allobaculum stercoricanis (RS_GCF_000384195.1)
Ileibacterium valens (RS_GCF_001945525.1)
Faecalibaculum rodentium (RS_GCF_001564455.1)
Dubosiella newyorkensis (RS_GCF_001945605.1)
Dubosiella sp000403415 (RS_GCF_000403415.2)
Merdibacter massiliensis (RS_GCF_900155395.1)
Absiella (metabat.515)
Longicatena dolichum (RS_GCF_000154285.1)
Absiella (metabat.250)
Longicatena sp000165065 (GB_GCA_000165065.1)
Longicatena innocuum (RS_GCF_000371425.1)
Longicatena caecimuris (RS_GCF_000163515.2)
OEMR01 sp900199515 (RS_GCF_900199515.1)
Dielma sp001305055 (RS_GCF_001305055.1)
Dielma fastidiosa (RS_GCF_000313565.1)
Traorella massiliensis (RS_GCF_900104665.1)
Breznakia sp001695645 (RS_GCF_001695645.1)
UBA636 sp900321955 (GB_GCA_900321955.1)
UBA636 sp002362065 (GB_GCA_002362065.1)
UBA636 sp002314515 (GB_GCA_002314515.1)
UBA636 sp002299675 (GB_GCA_002299675.1)
UBA636 sp002315565 (GB_GCA_002315565.1)
Erysipelotrichaceae (metabat.178)
Erysipelotrichaceae (metabat.522)
Erysipelotrichaceae (metabat.51)
RUG521 sp900319675 (GB_GCA_900319675.1)
UBA1783 sp002315205 (GB_GCA_002315205.1)

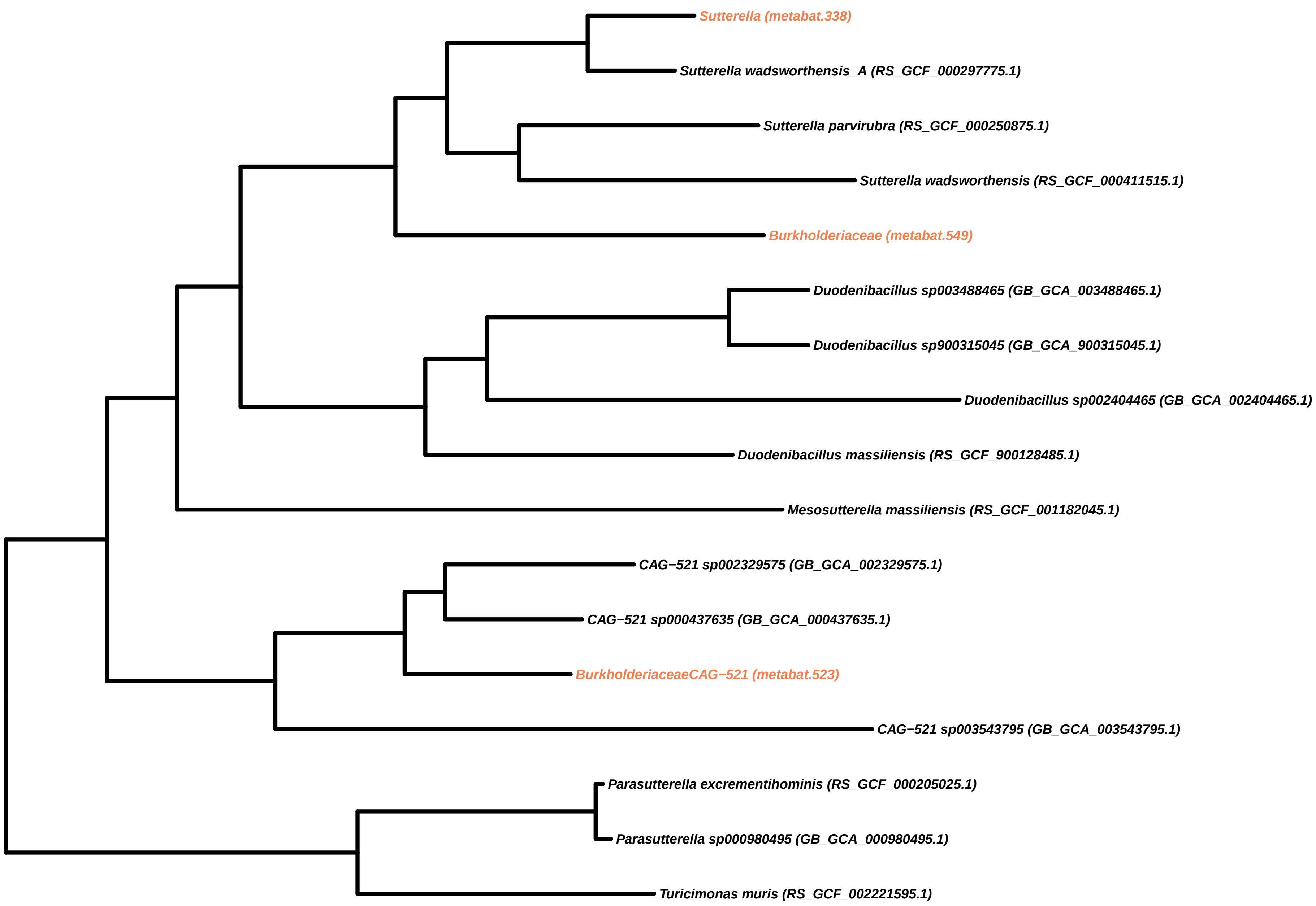
Sutterella (metabat.338)
Sutterella wadsworthensis_A (RS_GCF_000297775.1)
Sutterella parvirubra (RS_GCF_000250875.1)
Sutterella wadsworthensis (RS_GCF_000411515.1)
Burkholderiaceae (metabat.549)
Duodenibacillus sp003488465 (GB_GCA_003488465.1)
Duodenibacillus sp900315045 (GB_GCA_900315045.1)
Duodenibacillus sp002404465 (GB_GCA_002404465.1)
Duodenibacillus massiliensis (RS_GCF_900128485.1)
Mesosutterella massiliensis (RS_GCF_001182045.1)
CAG−521 sp002329575 (GB_GCA_002329575.1)
CAG−521 sp000437635 (GB_GCA_000437635.1)
BurkholderiaceaeCAG−521 (metabat.523)
CAG−521 sp003543795 (GB_GCA_003543795.1)
Parasutterella excrementihominis (RS_GCF_000205025.1)
Parasutterella sp000980495 (GB_GCA_000980495.1)
Turicimonas muris (RS_GCF_002221595.1)

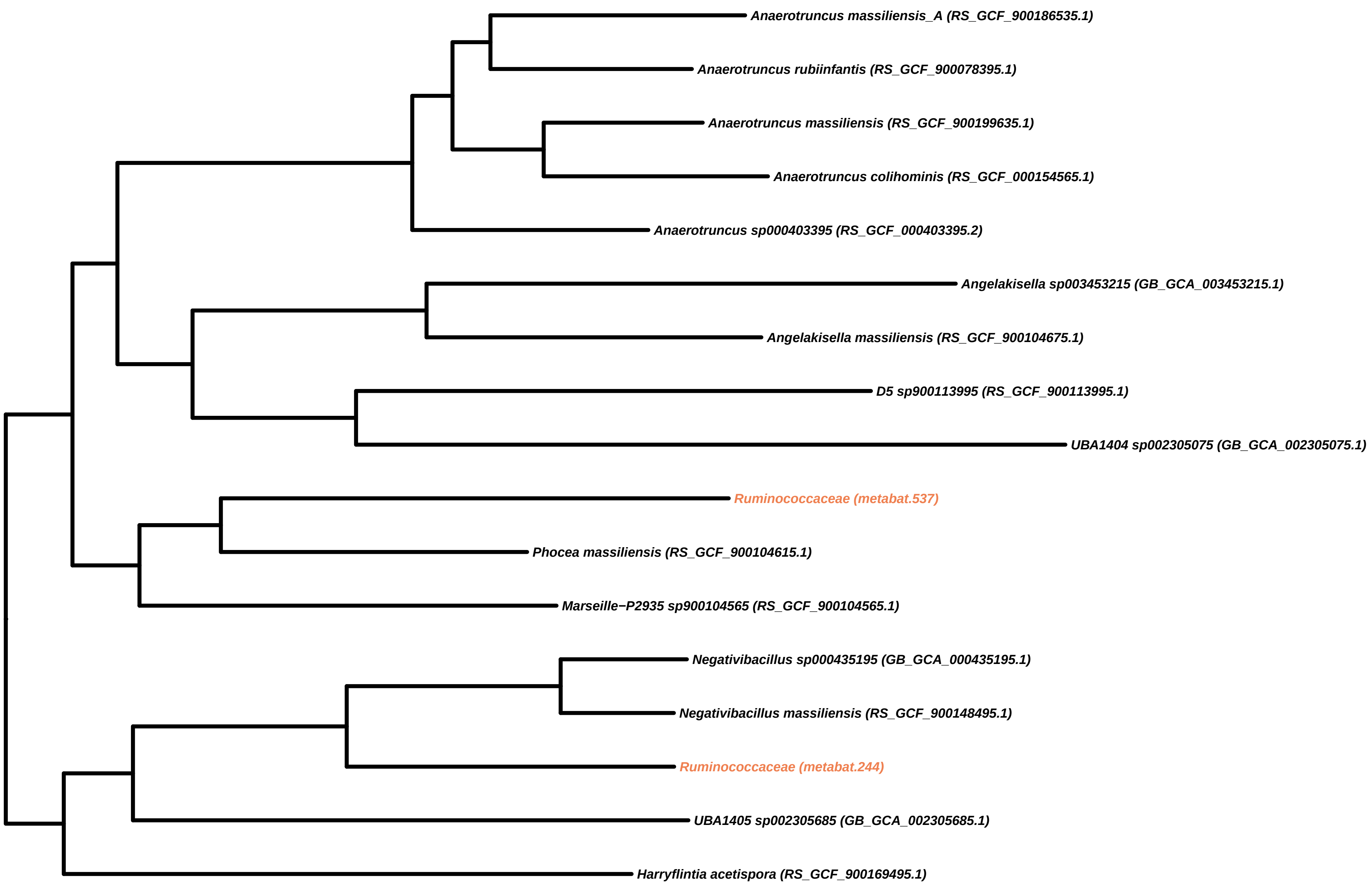
Anaerotruncus massiliensis_A (RS_GCF_900186535.1)
Anaerotruncus rubiinfantis (RS_GCF_900078395.1)
Anaerotruncus massiliensis (RS_GCF_900199635.1)
Anaerotruncus colihominis (RS_GCF_000154565.1)
Anaerotruncus sp000403395 (RS_GCF_000403395.2)
Angelakisella sp003453215 (GB_GCA_003453215.1)
Angelakisella massiliensis (RS_GCF_900104675.1)
D5 sp900113995 (RS_GCF_900113995.1)
UBA1404 sp002305075 (GB_GCA_002305075.1)
Ruminococcaceae (metabat.537)
Phocea massiliensis (RS_GCF_900104615.1)
Marseille−P2935 sp900104565 (RS_GCF_900104565.1)
Negativibacillus sp000435195 (GB_GCA_000435195.1)
Negativibacillus massiliensis (RS_GCF_900148495.1)
Ruminococcaceae (metabat.244)
UBA1405 sp002305685 (GB_GCA_002305685.1)
Harryflintia acetispora (RS_GCF_900169495.1)

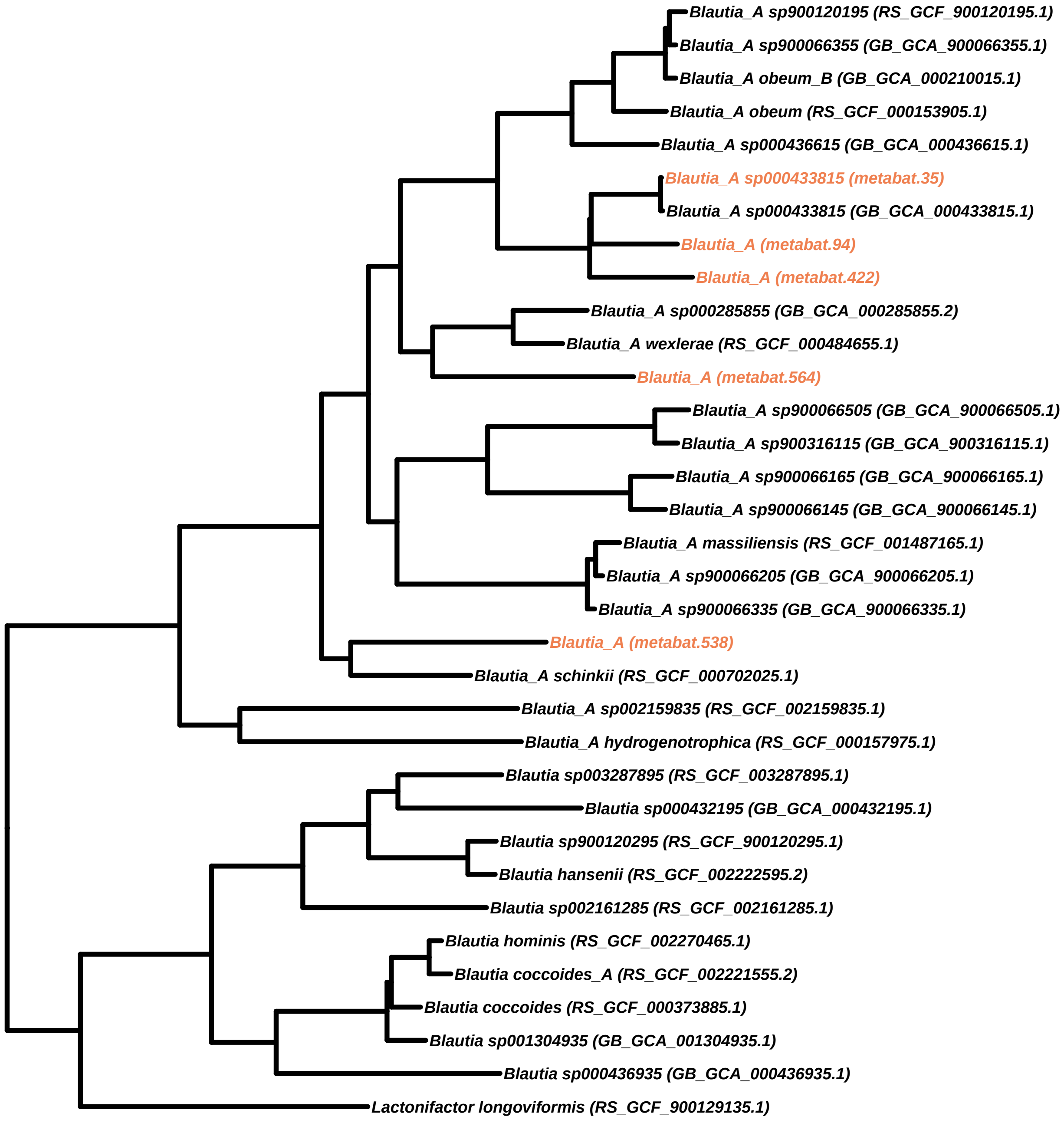
Blautia_A sp900120195 (RS_GCF_900120195.1)
Blautia_A sp900066355 (GB_GCA_900066355.1)
Blautia_A obeum_B (GB_GCA_000210015.1)
Blautia_A obeum (RS_GCF_000153905.1)
Blautia_A sp000436615 (GB_GCA_000436615.1)
Blautia_A sp000433815 (metabat.35)
Blautia_A sp000433815 (GB_GCA_000433815.1)
Blautia_A (metabat.94)
Blautia_A (metabat.422)
Blautia_A sp000285855 (GB_GCA_000285855.2)
Blautia_A wexlerae (RS_GCF_000484655.1)
Blautia_A (metabat.564)
Blautia_A sp900066505 (GB_GCA_900066505.1)
Blautia_A sp900316115 (GB_GCA_900316115.1)
Blautia_A sp900066165 (GB_GCA_900066165.1)
Blautia_A sp900066145 (GB_GCA_900066145.1)
Blautia_A massiliensis (RS_GCF_001487165.1)
Blautia_A sp900066205 (GB_GCA_900066205.1)
Blautia_A sp900066335 (GB_GCA_900066335.1)
Blautia_A (metabat.538)
Blautia_A schinkii (RS_GCF_000702025.1)
Blautia_A sp002159835 (RS_GCF_002159835.1)
Blautia_A hydrogenotrophica (RS_GCF_000157975.1)
Blautia sp003287895 (RS_GCF_003287895.1)
Blautia sp000432195 (GB_GCA_000432195.1)
Blautia sp900120295 (RS_GCF_900120295.1)
Blautia hansenii (RS_GCF_002222595.2)
Blautia sp002161285 (RS_GCF_002161285.1)
Blautia hominis (RS_GCF_002270465.1)
Blautia coccoides_A (RS_GCF_002221555.2)
Blautia coccoides (RS_GCF_000373885.1)
Blautia sp001304935 (GB_GCA_001304935.1)
Blautia sp000436935 (GB_GCA_000436935.1)
Lactonifactor longoviformis (RS_GCF_900129135.1)

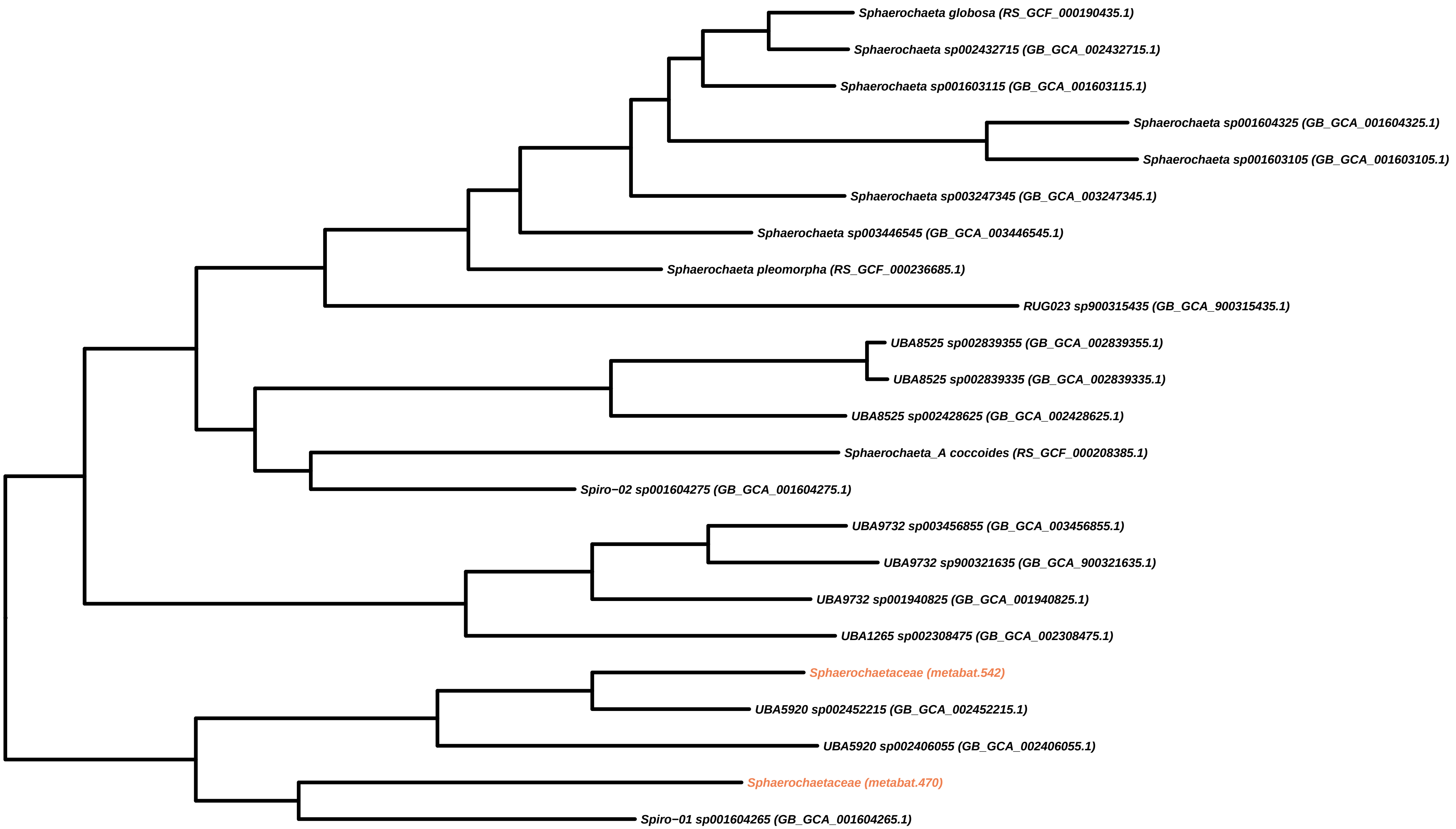
Sphaerochaeta globosa (RS_GCF_000190435.1)
Sphaerochaeta sp002432715 (GB_GCA_002432715.1)
Sphaerochaeta sp001603115 (GB_GCA_001603115.1)
Sphaerochaeta sp001604325 (GB_GCA_001604325.1)
Sphaerochaeta sp001603105 (GB_GCA_001603105.1)
Sphaerochaeta sp003247345 (GB_GCA_003247345.1)
Sphaerochaeta sp003446545 (GB_GCA_003446545.1)
Sphaerochaeta pleomorpha (RS_GCF_000236685.1)
RUG023 sp900315435 (GB_GCA_900315435.1)
UBA8525 sp002839355 (GB_GCA_002839355.1)
UBA8525 sp002839335 (GB_GCA_002839335.1)
UBA8525 sp002428625 (GB_GCA_002428625.1)
Sphaerochaeta_A coccoides (RS_GCF_000208385.1)
Spiro−02 sp001604275 (GB_GCA_001604275.1)
UBA9732 sp003456855 (GB_GCA_003456855.1)
UBA9732 sp900321635 (GB_GCA_900321635.1)
UBA9732 sp001940825 (GB_GCA_001940825.1)
UBA1265 sp002308475 (GB_GCA_002308475.1)
Sphaerochaetaceae (metabat.542)
UBA5920 sp002452215 (GB_GCA_002452215.1)
UBA5920 sp002406055 (GB_GCA_002406055.1)
Sphaerochaetaceae (metabat.470)
Spiro−01 sp001604265 (GB_GCA_001604265.1)

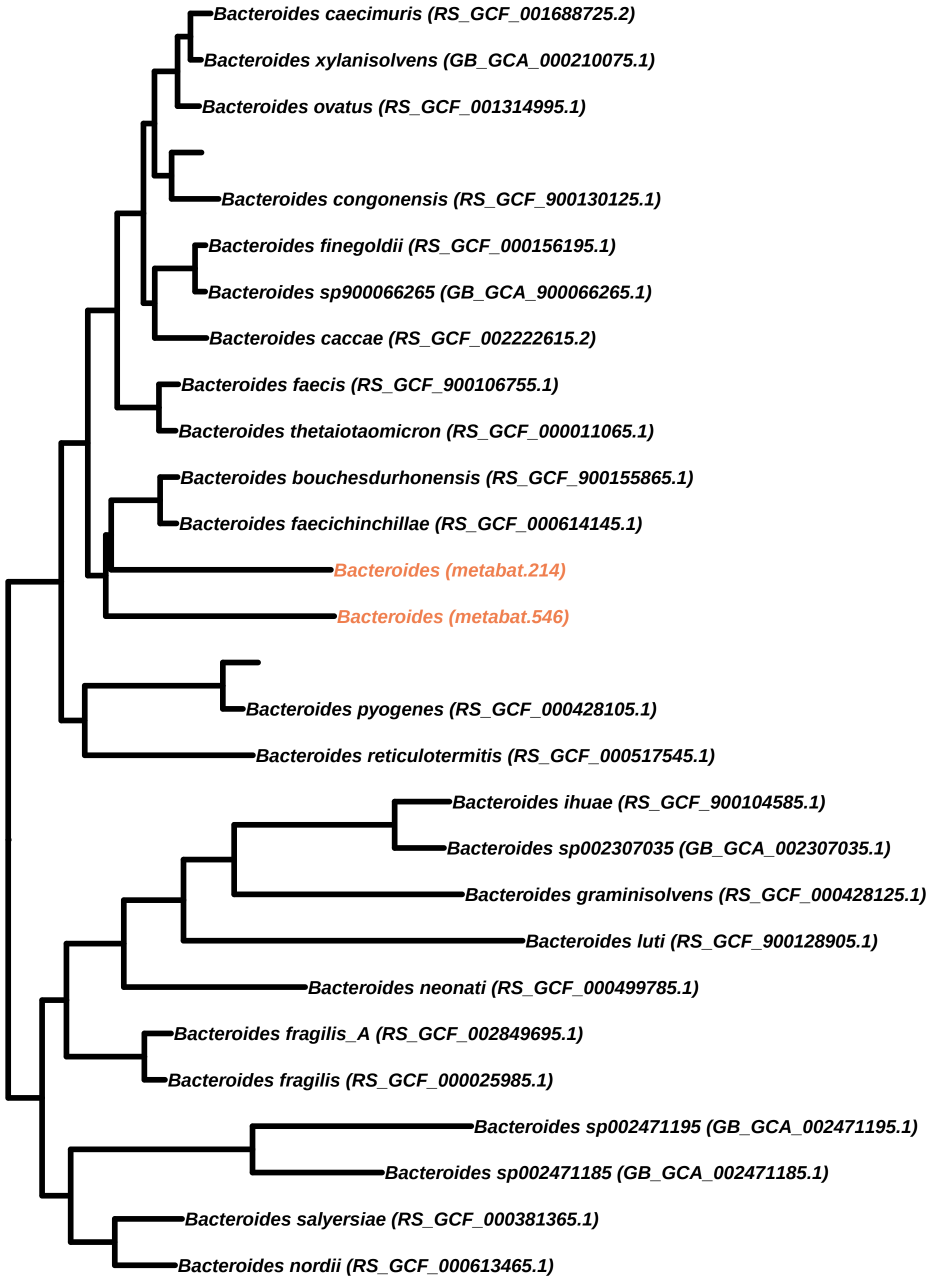
Bacteroides caecimuris (RS_GCF_001688725.2)
Bacteroides xylanisolvens (GB_GCA_000210075.1)
Bacteroides ovatus (RS_GCF_001314995.1)
Bacteroides congonensis (RS_GCF_900130125.1)
Bacteroides finegoldii (RS_GCF_000156195.1)
Bacteroides sp900066265 (GB_GCA_900066265.1)
Bacteroides caccae (RS_GCF_002222615.2)
Bacteroides faecis (RS_GCF_900106755.1)
Bacteroides thetaiotaomicron (RS_GCF_000011065.1)
Bacteroides bouchesdurhonensis (RS_GCF_900155865.1)
Bacteroides faecichinchillae (RS_GCF_000614145.1)
Bacteroides (metabat.214)
Bacteroides (metabat.546)
Bacteroides pyogenes (RS_GCF_000428105.1)
Bacteroides reticulotermitis (RS_GCF_000517545.1)
Bacteroides ihuae (RS_GCF_900104585.1)
Bacteroides sp002307035 (GB_GCA_002307035.1)
Bacteroides graminisolvens (RS_GCF_000428125.1)
Bacteroides luti (RS_GCF_900128905.1)
Bacteroides neonati (RS_GCF_000499785.1)
Bacteroides fragilis_A (RS_GCF_002849695.1)
Bacteroides fragilis (RS_GCF_000025985.1)
Bacteroides sp002471195 (GB_GCA_002471195.1)
Bacteroides sp002471185 (GB_GCA_002471185.1)
Bacteroides salyersiae (RS_GCF_000381365.1)
Bacteroides nordii (RS_GCF_000613465.1)

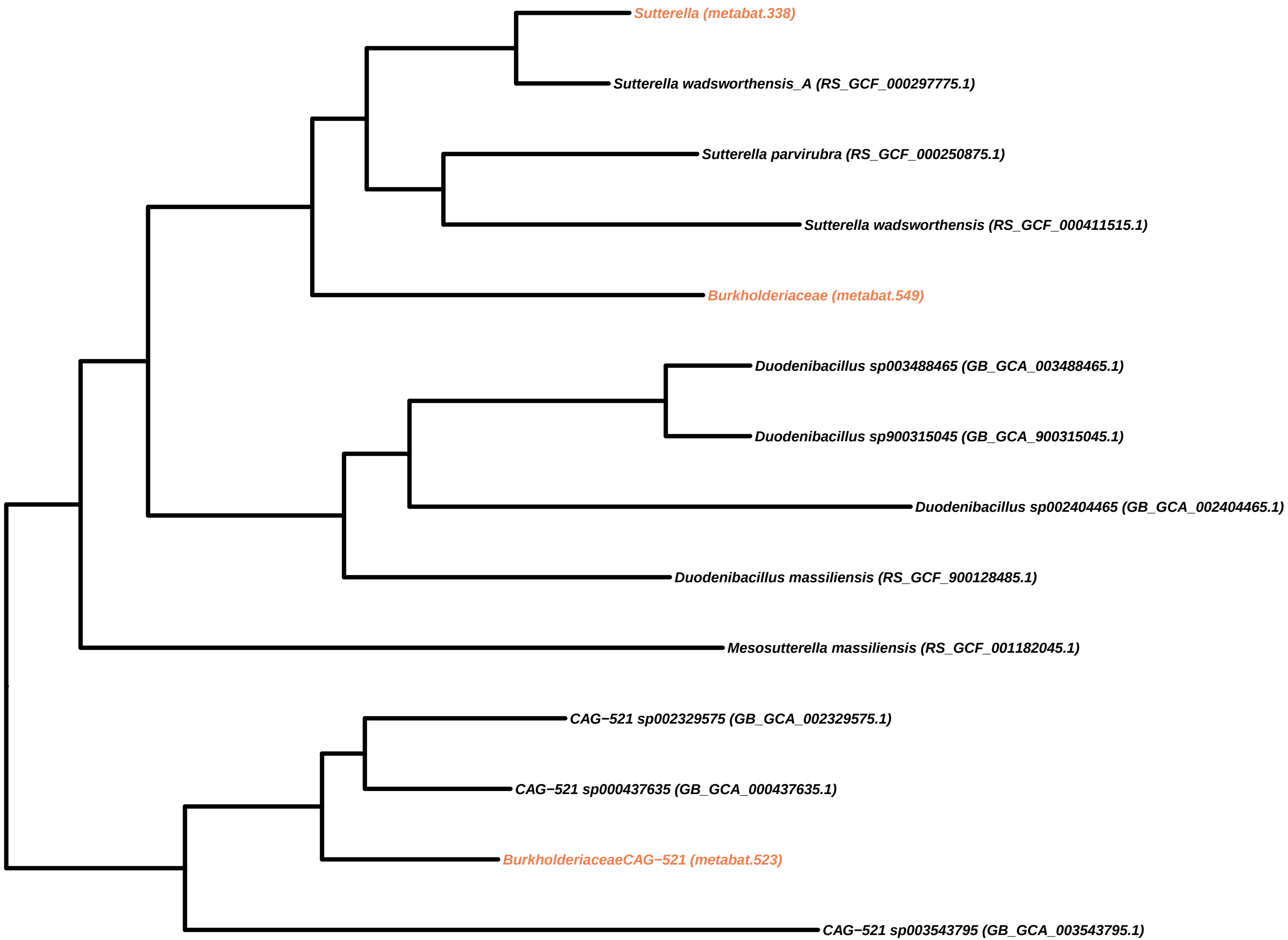

Sutterella (metabat.338)
Sutterella wadsworthensis_A (RS_GCF_000297775.1)
Sutterella parvirubra (RS_GCF_000250875.1)
Sutterella wadsworthensis (RS_GCF_000411515.1)
Burkholderiaceae (metabat.549)
Duodenibacillus sp003488465 (GB_GCA_003488465.1)
Duodenibacillus sp900315045 (GB_GCA_900315045.1)
Duodenibacillus sp002404465 (GB_GCA_002404465.1)
Duodenibacillus massiliensis (RS_GCF_900128485.1)
Mesosutterella massiliensis (RS_GCF_001182045.1)
CAG−521 sp002329575 (GB_GCA_002329575.1)
CAG−521 sp000437635 (GB_GCA_000437635.1)
BurkholderiaceaeCAG−521 (metabat.523)
CAG−521 sp003543795 (GB_GCA_003543795.1)

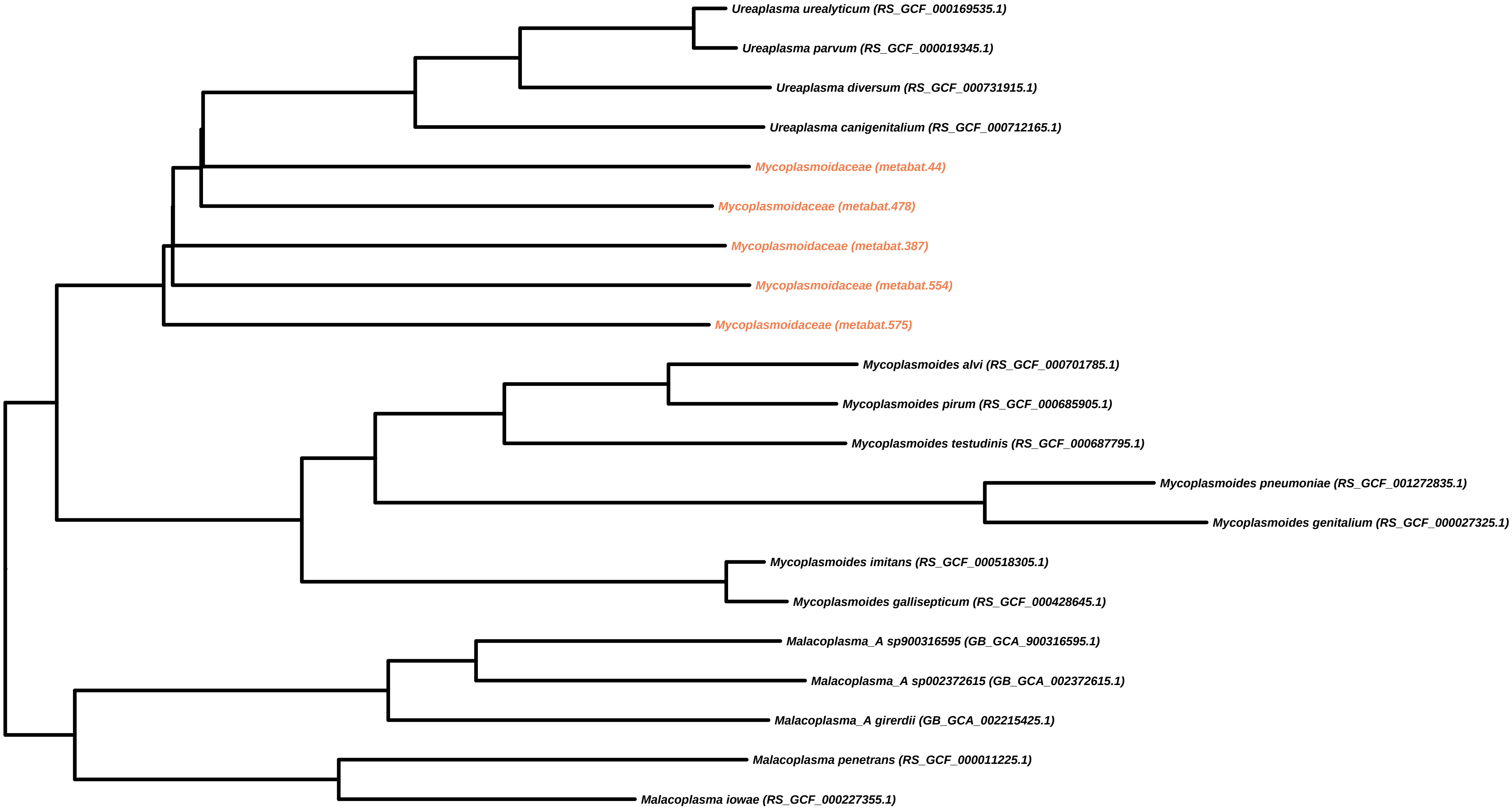
Ureaplasma urealyticum (RS_GCF_000169535.1)
Ureaplasma parvum (RS_GCF_000019345.1)
Ureaplasma diversum (RS_GCF_000731915.1)
Ureaplasma canigenitalium (RS_GCF_000712165.1)
Mycoplasmoidaceae (metabat.44)
Mycoplasmoidaceae (metabat.478)
Mycoplasmoidaceae (metabat.387)
Mycoplasmoidaceae (metabat.554)
Mycoplasmoidaceae (metabat.575)
Mycoplasmoides alvi (RS_GCF_000701785.1)
Mycoplasmoides pirum (RS_GCF_000685905.1)
Mycoplasmoides testudinis (RS_GCF_000687795.1)
Mycoplasmoides pneumoniae (RS_GCF_001272835.1)
Mycoplasmoides genitalium (RS_GCF_000027325.1)
Mycoplasmoides imitans (RS_GCF_000518305.1)
Mycoplasmoides gallisepticum (RS_GCF_000428645.1)
Malacoplasma_A sp900316595 (GB_GCA_900316595.1)
Malacoplasma_A sp002372615 (GB_GCA_002372615.1)
Malacoplasma_A girerdii (GB_GCA_002215425.1)
Malacoplasma penetrans (RS_GCF_000011225.1)
Malacoplasma iowae (RS_GCF_000227355.1)

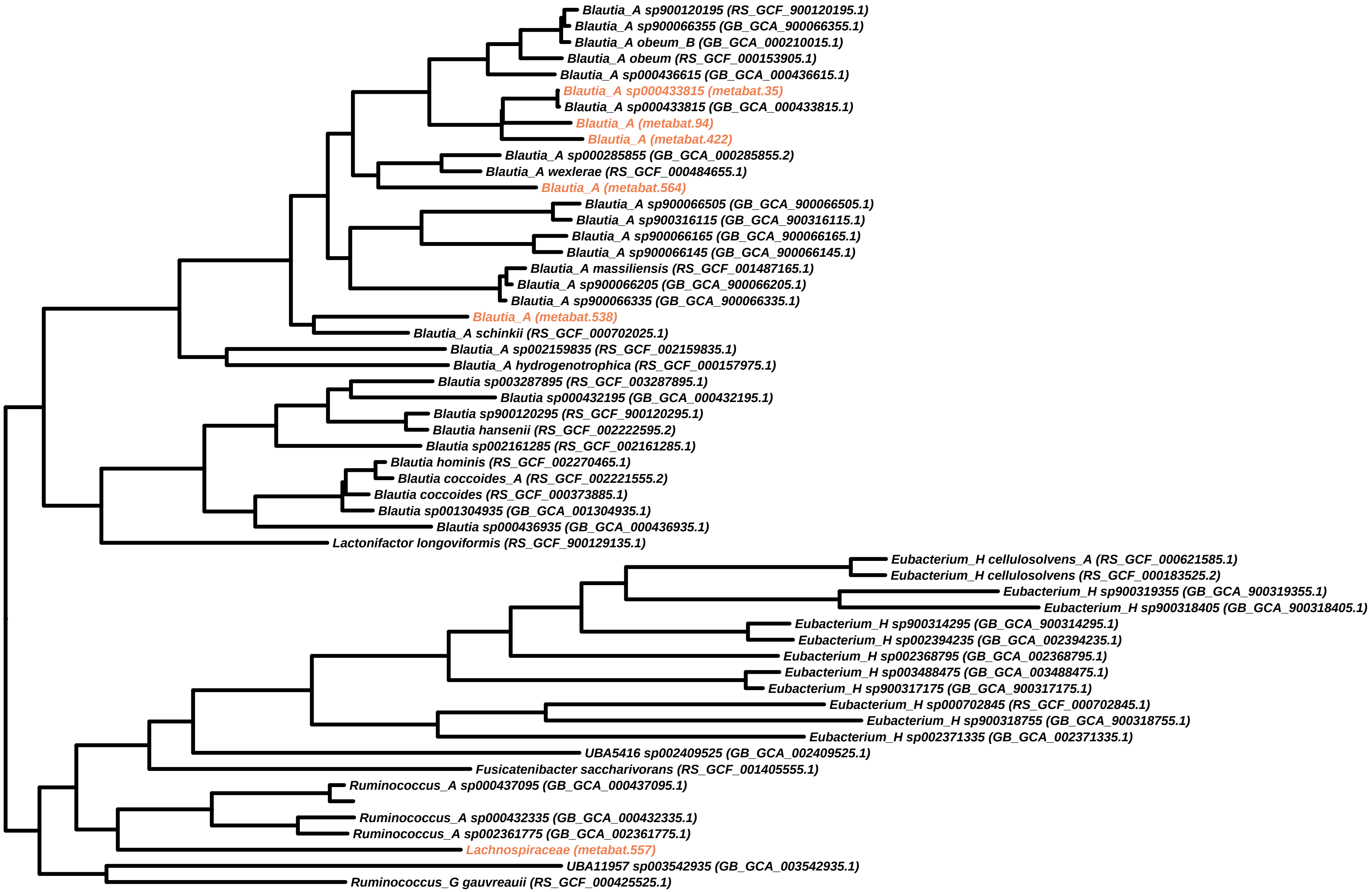

Blautia_A sp900120195 (RS_GCF_900120195.1)
Blautia_A sp900066355 (GB_GCA_900066355.1)
Blautia_A obeum_B (GB_GCA_000210015.1)
Blautia_A obeum (RS_GCF_000153905.1)
Blautia_A sp000436615 (GB_GCA_000436615.1)
Blautia_A sp000433815 (metabat.35)
Blautia_A sp000433815 (GB_GCA_000433815.1)
Blautia_A (metabat.94)
Blautia_A (metabat.422)
Blautia_A sp000285855 (GB_GCA_000285855.2)
Blautia_A wexlerae (RS_GCF_000484655.1)
Blautia_A (metabat.564)
Blautia_A sp900066505 (GB_GCA_900066505.1)
Blautia_A sp900316115 (GB_GCA_900316115.1)
Blautia_A sp900066165 (GB_GCA_900066165.1)
Blautia_A sp900066145 (GB_GCA_900066145.1)
Blautia_A massiliensis (RS_GCF_001487165.1)
Blautia_A sp900066205 (GB_GCA_900066205.1)
Blautia_A sp900066335 (GB_GCA_900066335.1)
Blautia_A (metabat.538)
Blautia_A schinkii (RS_GCF_000702025.1)
Blautia_A sp002159835 (RS_GCF_002159835.1)
Blautia_A hydrogenotrophica (RS_GCF_000157975.1)
Blautia sp003287895 (RS_GCF_003287895.1)
Blautia sp000432195 (GB_GCA_000432195.1)
Blautia sp900120295 (RS_GCF_900120295.1)
Blautia hansenii (RS_GCF_002222595.2)
Blautia sp002161285 (RS_GCF_002161285.1)
Blautia hominis (RS_GCF_002270465.1)
Blautia coccoides_A (RS_GCF_002221555.2)
Blautia coccoides (RS_GCF_000373885.1)
Blautia sp001304935 (GB_GCA_001304935.1)
Blautia sp000436935 (GB_GCA_000436935.1)
Lactonifactor longoviformis (RS_GCF_900129135.1)
Eubacterium_H cellulosolvens_A (RS_GCF_000621585.1)
Eubacterium_H cellulosolvens (RS_GCF_000183525.2)
Eubacterium_H sp900319355 (GB_GCA_900319355.1)
Eubacterium_H sp900318405 (GB_GCA_900318405.1)
Eubacterium_H sp900314295 (GB_GCA_900314295.1)
Eubacterium_H sp002394235 (GB_GCA_002394235.1)
Eubacterium_H sp002368795 (GB_GCA_002368795.1)
Eubacterium_H sp003488475 (GB_GCA_003488475.1)
Eubacterium_H sp900317175 (GB_GCA_900317175.1)
Eubacterium_H sp000702845 (RS_GCF_000702845.1)
Eubacterium_H sp900318755 (GB_GCA_900318755.1)
Eubacterium_H sp002371335 (GB_GCA_002371335.1)
UBA5416 sp002409525 (GB_GCA_002409525.1)
Fusicatenibacter saccharivorans (RS_GCF_001405555.1)
Ruminococcus_A sp000437095 (GB_GCA_000437095.1)
Ruminococcus_A sp000432335 (GB_GCA_000432335.1)
Ruminococcus_A sp002361775 (GB_GCA_002361775.1)
Lachnospiraceae (metabat.557)
UBA11957 sp003542935 (GB_GCA_003542935.1)
Ruminococcus_G gauvreauii (RS_GCF_000425525.1)

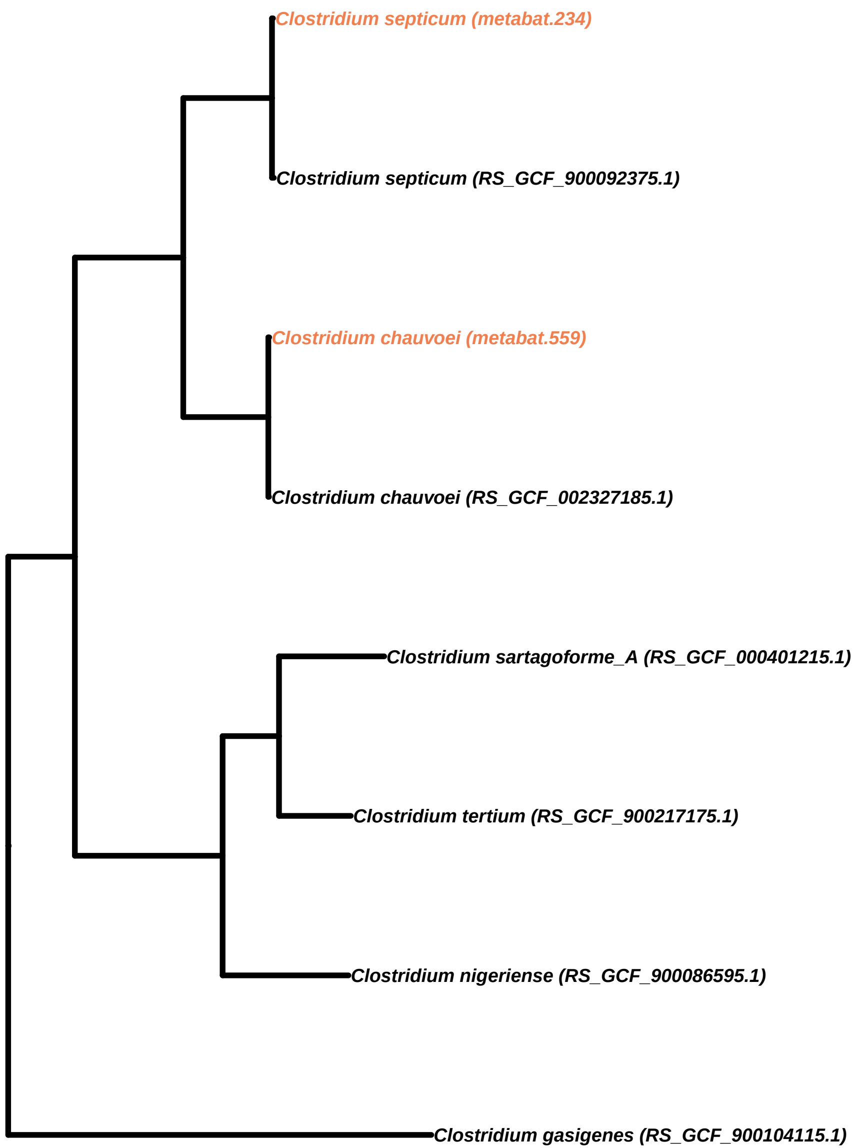

Clostridium septicum (metabat.234)
Clostridium septicum (RS_GCF_900092375.1)
Clostridium chauvoei (metabat.559)
Clostridium chauvoei (RS_GCF_002327185.1)
Clostridium sartagoforme_A (RS_GCF_000401215.1)
Clostridium tertium (RS_GCF_900217175.1)
Clostridium nigeriense (RS_GCF_900086595.1)
Clostridium gasigenes (RS_GCF_900104115.1)

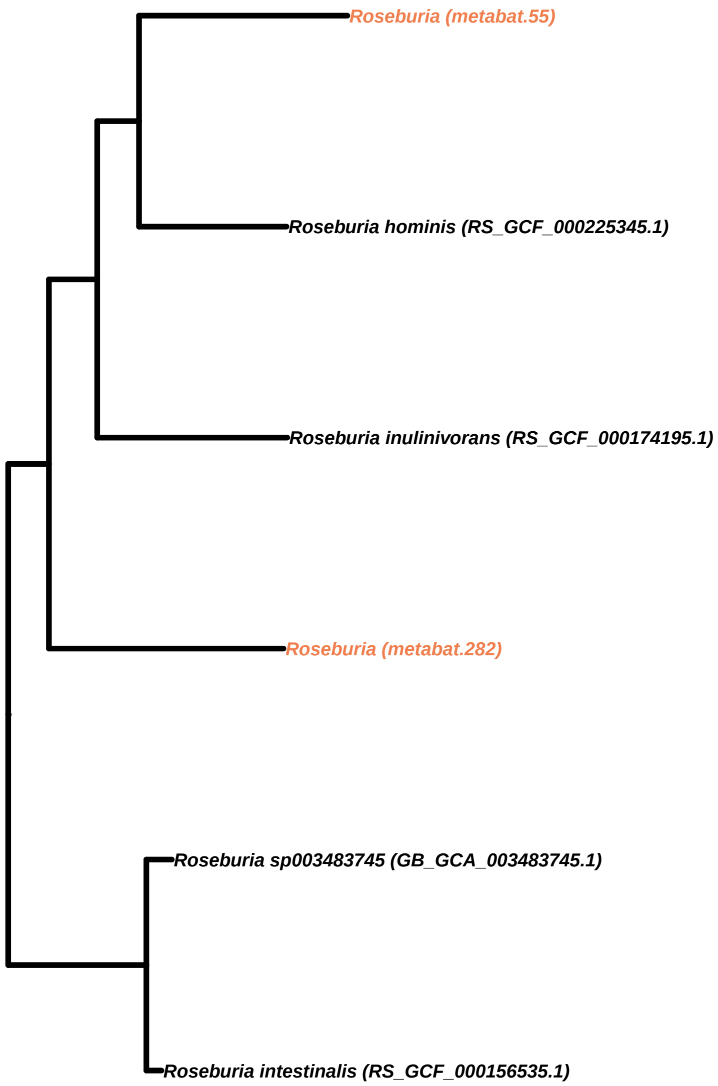
Roseburia (metabat.55)
Roseburia hominis (RS_GCF_000225345.1)
Roseburia inulinivorans (RS_GCF_000174195.1)
Roseburia (metabat.282)
Roseburia sp003483745 (GB_GCA_003483745.1)
Roseburia intestinalis (RS_GCF_000156535.1)

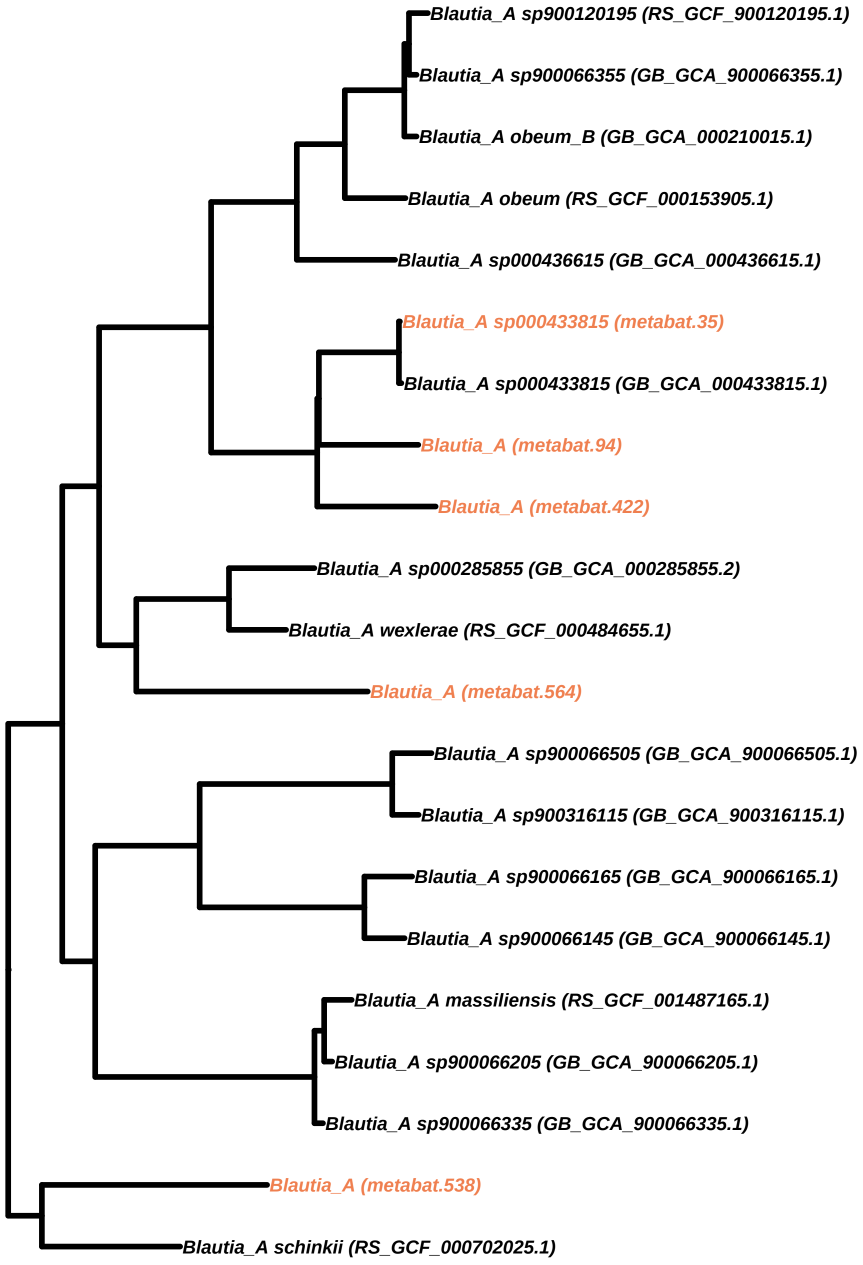
Blautia_A sp900120195 (RS_GCF_900120195.1)
Blautia_A sp900066355 (GB_GCA_900066355.1)
Blautia_A obeum_B (GB_GCA_000210015.1)
Blautia_A obeum (RS_GCF_000153905.1)
Blautia_A sp000436615 (GB_GCA_000436615.1)
Blautia_A sp000433815 (metabat.35)
Blautia_A sp000433815 (GB_GCA_000433815.1)
Blautia_A (metabat.94)
Blautia_A (metabat.422)
Blautia_A sp000285855 (GB_GCA_000285855.2)
Blautia_A wexlerae (RS_GCF_000484655.1)
Blautia_A (metabat.564)
Blautia_A sp900066505 (GB_GCA_900066505.1)
Blautia_A sp900316115 (GB_GCA_900316115.1)
Blautia_A sp900066165 (GB_GCA_900066165.1)
Blautia_A sp900066145 (GB_GCA_900066145.1)
Blautia_A massiliensis (RS_GCF_001487165.1)
Blautia_A sp900066205 (GB_GCA_900066205.1)
Blautia_A sp900066335 (GB_GCA_900066335.1)
Blautia_A (metabat.538)
Blautia_A schinkii (RS_GCF_000702025.1)

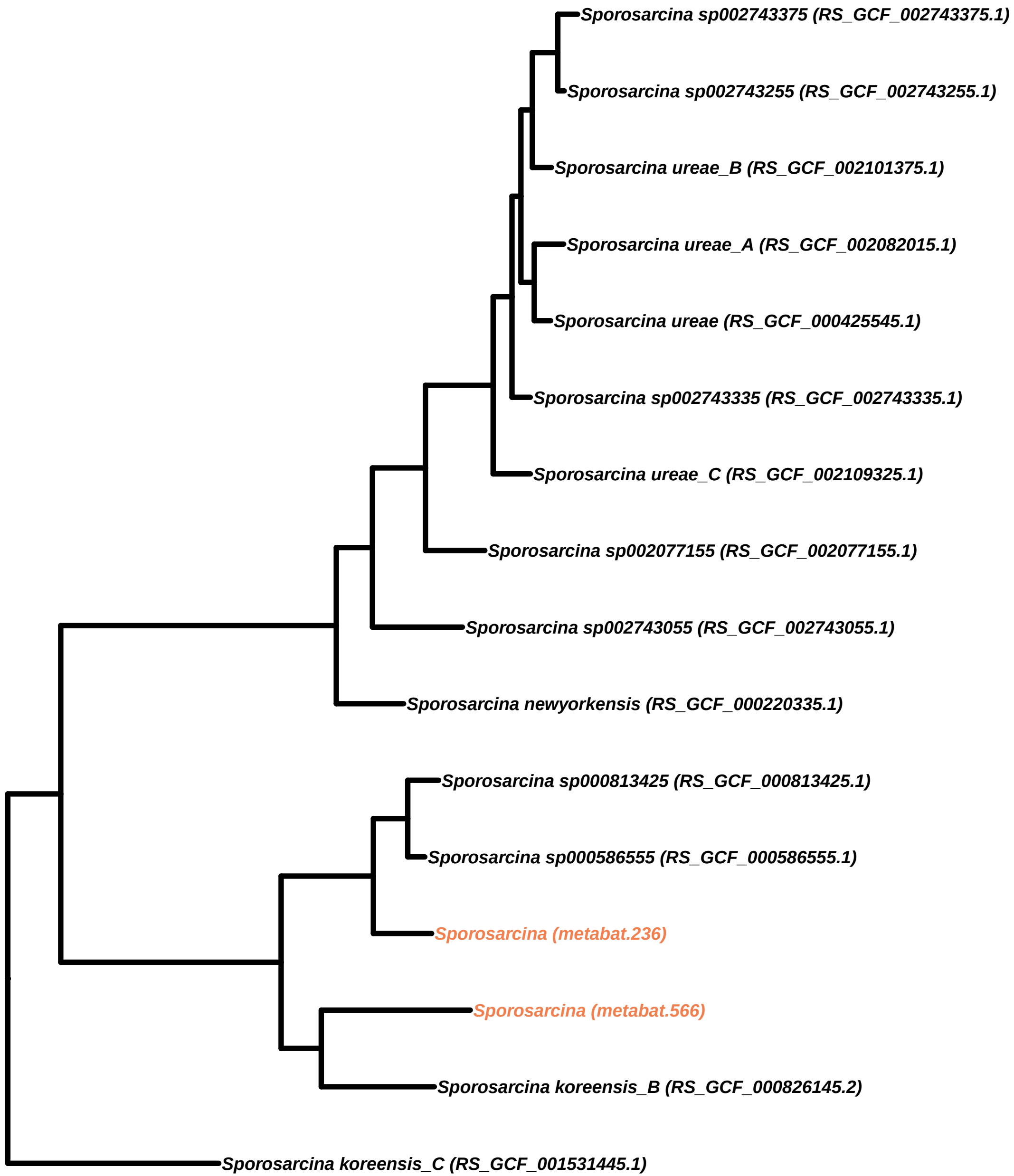
Sporosarcina sp002743375 (RS_GCF_002743375.1)
Sporosarcina sp002743255 (RS_GCF_002743255.1)
Sporosarcina ureae_B (RS_GCF_002101375.1)
Sporosarcina ureae_A (RS_GCF_002082015.1)
Sporosarcina ureae (RS_GCF_000425545.1)
Sporosarcina sp002743335 (RS_GCF_002743335.1)
Sporosarcina ureae_C (RS_GCF_002109325.1)
Sporosarcina sp002077155 (RS_GCF_002077155.1)
Sporosarcina sp002743055 (RS_GCF_002743055.1)
Sporosarcina newyorkensis (RS_GCF_000220335.1)
Sporosarcina sp000813425 (RS_GCF_000813425.1)
Sporosarcina sp000586555 (RS_GCF_000586555.1)
Sporosarcina (metabat.236)
Sporosarcina (metabat.566)
Sporosarcina koreensis_B (RS_GCF_000826145.2)
Sporosarcina koreensis_C (RS_GCF_001531445.1)

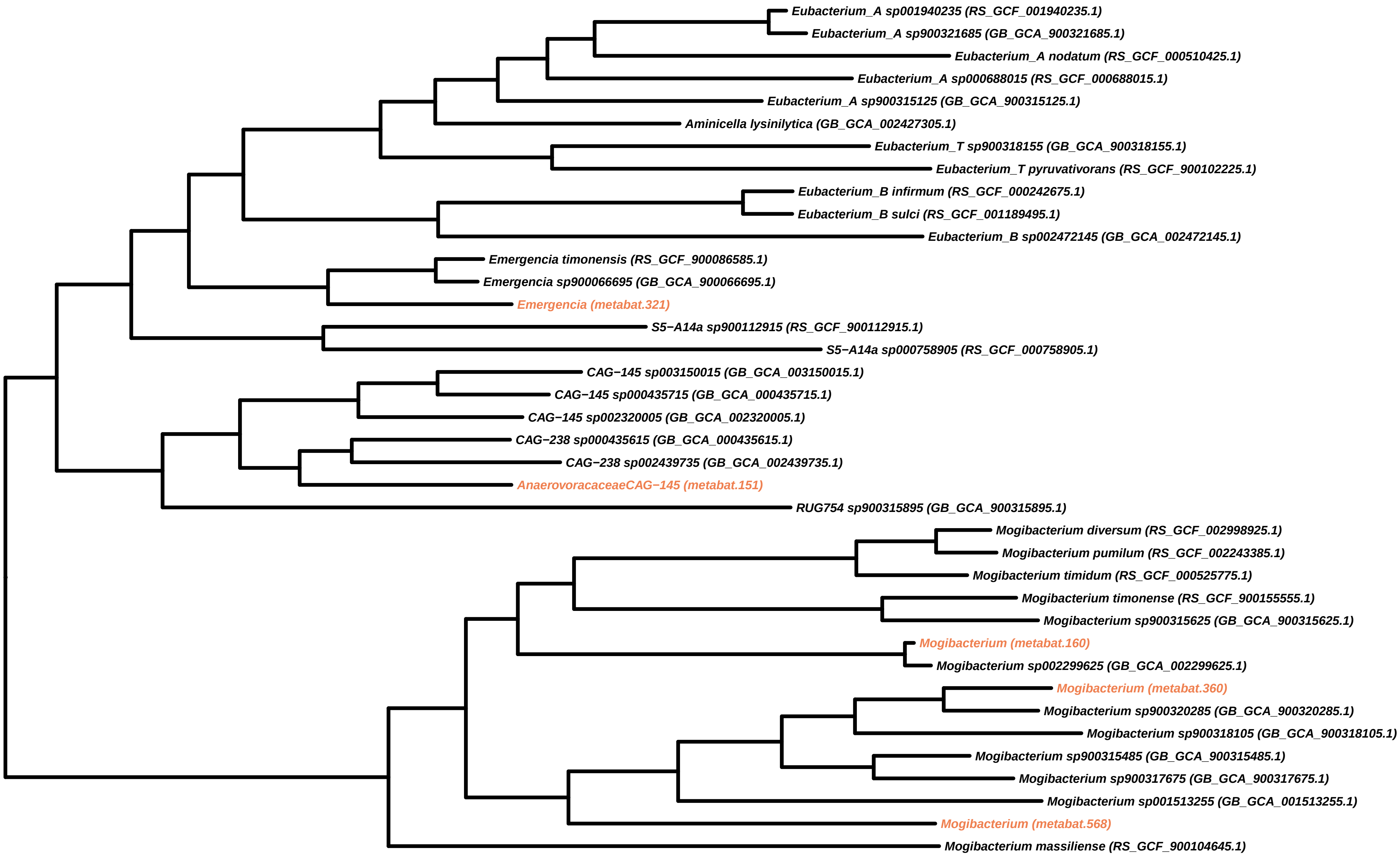
Eubacterium_A sp001940235 (RS_GCF_001940235.1)
Eubacterium_A sp900321685 (GB_GCA_900321685.1)
Eubacterium_A nodatum (RS_GCF_000510425.1)
Eubacterium_A sp000688015 (RS_GCF_000688015.1)
Eubacterium_A sp900315125 (GB_GCA_900315125.1)
Aminicella lysinilytica (GB_GCA_002427305.1)
Eubacterium_T sp900318155 (GB_GCA_900318155.1)
Eubacterium_T pyruvativorans (RS_GCF_900102225.1)
Eubacterium_B infirmum (RS_GCF_000242675.1)
Eubacterium_B sulci (RS_GCF_001189495.1)
Eubacterium_B sp002472145 (GB_GCA_002472145.1)
Emergencia timonensis (RS_GCF_900086585.1)
Emergencia sp900066695 (GB_GCA_900066695.1)
Emergencia (metabat.321)
S5−A14a sp900112915 (RS_GCF_900112915.1)
S5−A14a sp000758905 (RS_GCF_000758905.1)
CAG−145 sp003150015 (GB_GCA_003150015.1)
CAG−145 sp000435715 (GB_GCA_000435715.1)
CAG−145 sp002320005 (GB_GCA_002320005.1)
CAG−238 sp000435615 (GB_GCA_000435615.1)
CAG−238 sp002439735 (GB_GCA_002439735.1)
AnaerovoracaceaeCAG−145 (metabat.151)
RUG754 sp900315895 (GB_GCA_900315895.1)
Mogibacterium diversum (RS_GCF_002998925.1)
Mogibacterium pumilum (RS_GCF_002243385.1)
Mogibacterium timidum (RS_GCF_000525775.1)
Mogibacterium timonense (RS_GCF_900155555.1)
Mogibacterium sp900315625 (GB_GCA_900315625.1)
Mogibacterium (metabat.160)
Mogibacterium sp002299625 (GB_GCA_002299625.1)
Mogibacterium (metabat.360)
Mogibacterium sp900320285 (GB_GCA_900320285.1)
Mogibacterium sp900318105 (GB_GCA_900318105.1)
Mogibacterium sp900315485 (GB_GCA_900315485.1)
Mogibacterium sp900317675 (GB_GCA_900317675.1)
Mogibacterium sp001513255 (GB_GCA_001513255.1)
Mogibacterium (metabat.568)
Mogibacterium massiliense (RS_GCF_900104645.1)

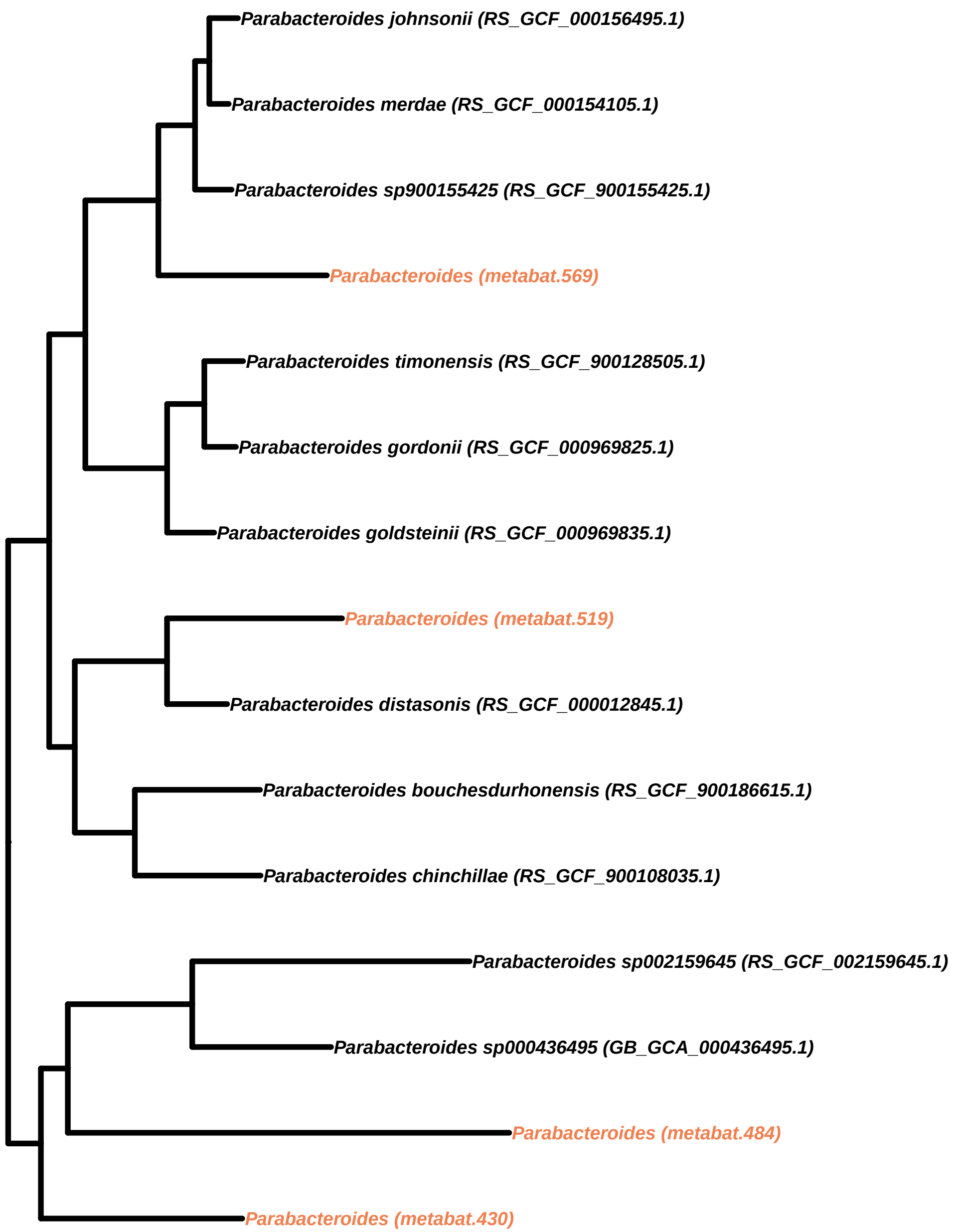
Parabacteroides johnsonii (RS_GCF_000156495.1)
Parabacteroides merdae (RS_GCF_000154105.1)
Parabacteroides sp900155425 (RS_GCF_900155425.1)
Parabacteroides (metabat.569)
Parabacteroides timonensis (RS_GCF_900128505.1)
Parabacteroides gordonii (RS_GCF_000969825.1)
Parabacteroides goldsteinii (RS_GCF_000969835.1)
Parabacteroides (metabat.519)
Parabacteroides distasonis (RS_GCF_000012845.1)
Parabacteroides bouchesdurhonensis (RS_GCF_900186615.1)
Parabacteroides chinchillae (RS_GCF_900108035.1)
Parabacteroides sp002159645 (RS_GCF_002159645.1)
Parabacteroides sp000436495 (GB_GCA_000436495.1)
Parabacteroides (metabat.484)
Parabacteroides (metabat.430)

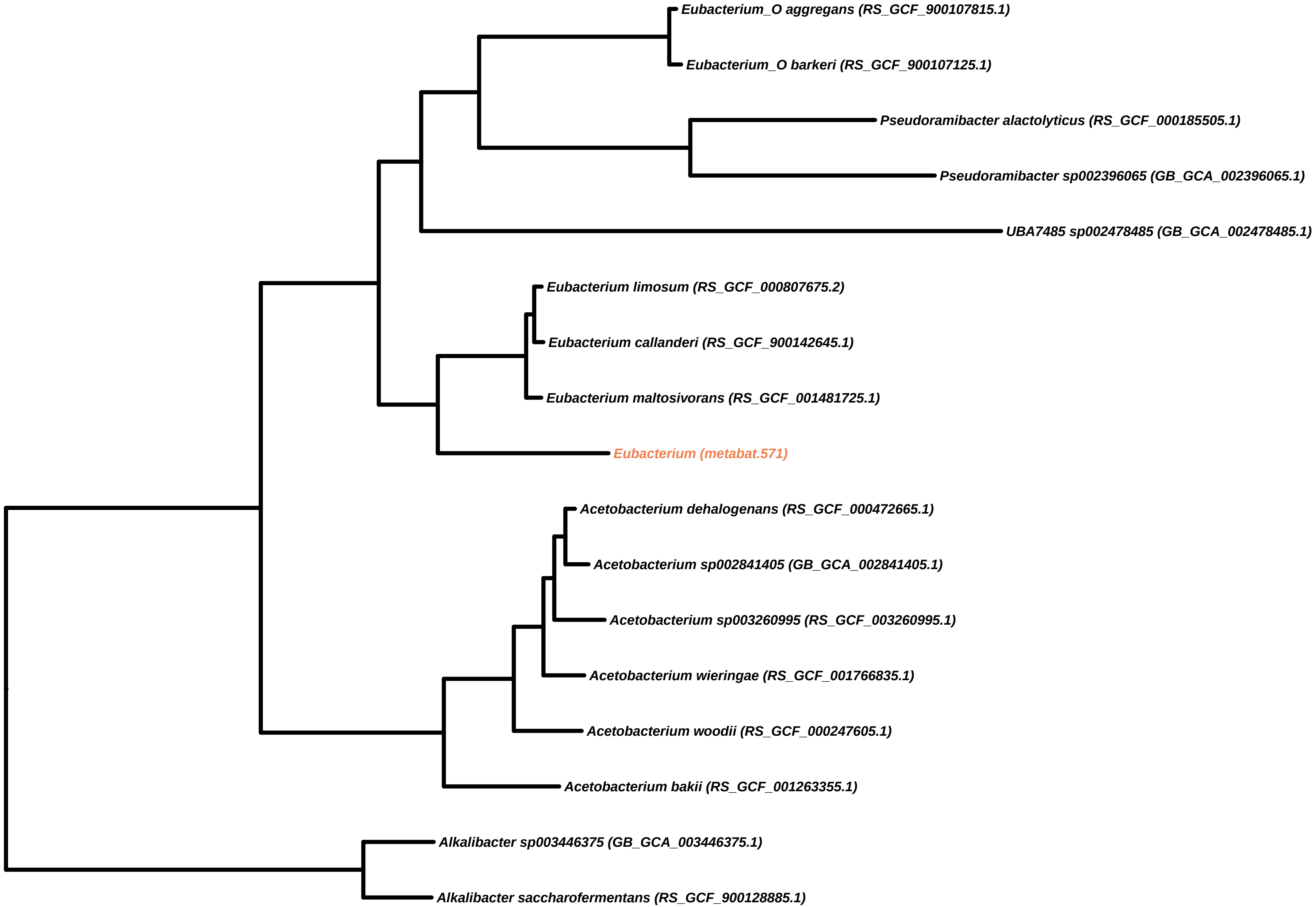
Eubacterium_O aggregans (RS_GCF_900107815.1)
Eubacterium_O barkeri (RS_GCF_900107125.1)
Pseudoramibacter alactolyticus (RS_GCF_000185505.1)
Pseudoramibacter sp002396065 (GB_GCA_002396065.1)
UBA7485 sp002478485 (GB_GCA_002478485.1)
Eubacterium limosum (RS_GCF_000807675.2)
Eubacterium callanderi (RS_GCF_900142645.1)
Eubacterium maltosivorans (RS_GCF_001481725.1)
Eubacterium (metabat.571)
Acetobacterium dehalogenans (RS_GCF_000472665.1)
Acetobacterium sp002841405 (GB_GCA_002841405.1)
Acetobacterium sp003260995 (RS_GCF_003260995.1)
Acetobacterium wieringae (RS_GCF_001766835.1)
Acetobacterium woodii (RS_GCF_000247605.1)
Acetobacterium bakii (RS_GCF_001263355.1)
Alkalibacter sp003446375 (GB_GCA_003446375.1)
Alkalibacter saccharofermentans (RS_GCF_900128885.1)

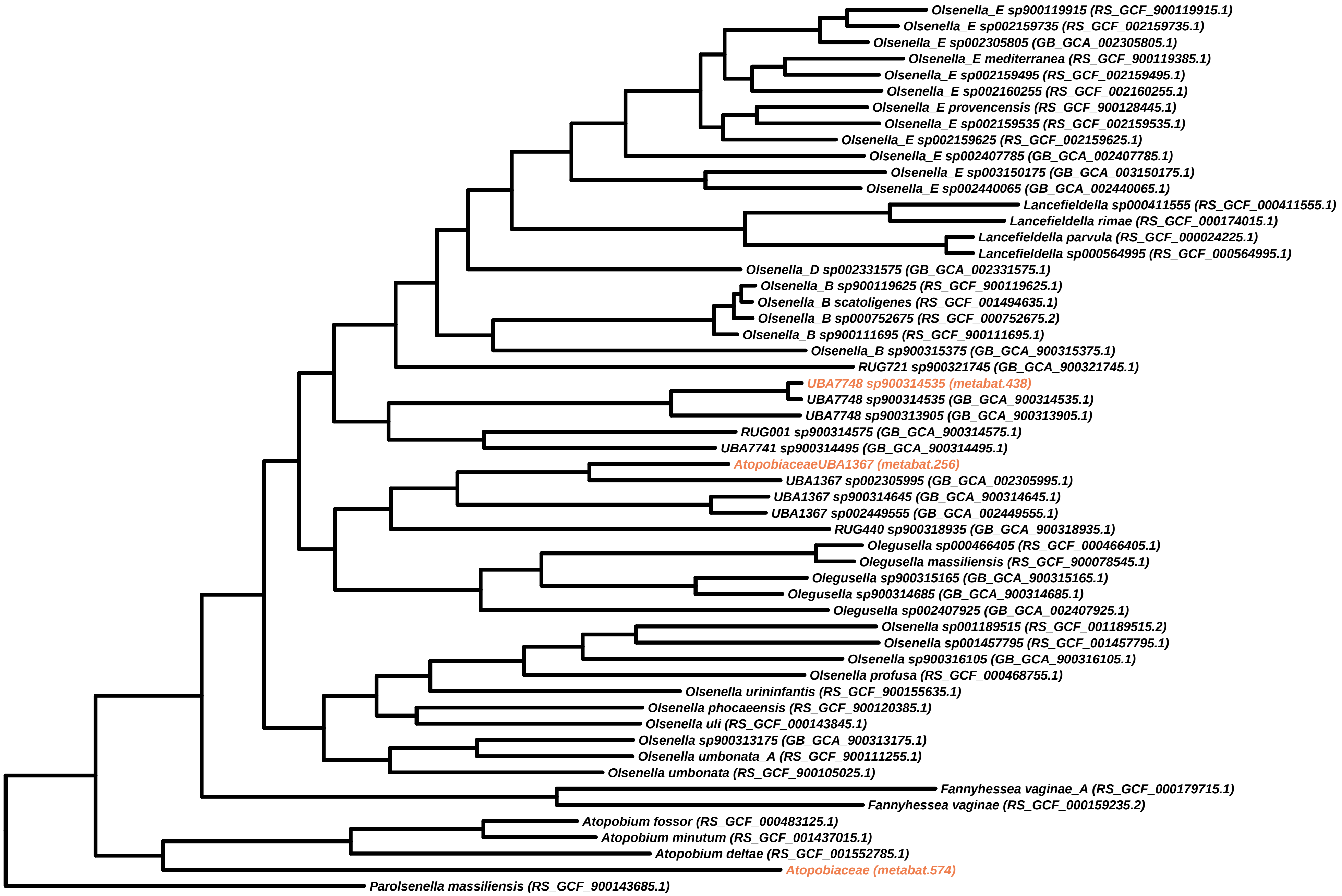
Olsenella_E sp900119915 (RS_GCF_900119915.1)
Olsenella_E sp002159735 (RS_GCF_002159735.1)
Olsenella_E sp002305805 (GB_GCA_002305805.1)
Olsenella_E mediterranea (RS_GCF_900119385.1)
Olsenella_E sp002159495 (RS_GCF_002159495.1)
Olsenella_E sp002160255 (RS_GCF_002160255.1)
Olsenella_E provencensis (RS_GCF_900128445.1)
Olsenella_E sp002159535 (RS_GCF_002159535.1)
Olsenella_E sp002159625 (RS_GCF_002159625.1)
Olsenella_E sp002407785 (GB_GCA_002407785.1)
Olsenella_E sp003150175 (GB_GCA_003150175.1)
Olsenella_E sp002440065 (GB_GCA_002440065.1)
Lancefieldella sp000411555 (RS_GCF_000411555.1)
Lancefieldella rimae (RS_GCF_000174015.1)
Lancefieldella parvula (RS_GCF_000024225.1)
Lancefieldella sp000564995 (RS_GCF_000564995.1)
Olsenella_D sp002331575 (GB_GCA_002331575.1)
Olsenella_B sp900119625 (RS_GCF_900119625.1)
Olsenella_B scatoligenes (RS_GCF_001494635.1)
Olsenella_B sp000752675 (RS_GCF_000752675.2)
Olsenella_B sp900111695 (RS_GCF_900111695.1)
Olsenella_B sp900315375 (GB_GCA_900315375.1)
RUG721 sp900321745 (GB_GCA_900321745.1)
UBA7748 sp900314535 (metabat.438)
UBA7748 sp900314535 (GB_GCA_900314535.1)
UBA7748 sp900313905 (GB_GCA_900313905.1)
RUG001 sp900314575 (GB_GCA_900314575.1)
UBA7741 sp900314495 (GB_GCA_900314495.1)
AtopobiaceaeUBA1367 (metabat.256)
UBA1367 sp002305995 (GB_GCA_002305995.1)
UBA1367 sp900314645 (GB_GCA_900314645.1)
UBA1367 sp002449555 (GB_GCA_002449555.1)
RUG440 sp900318935 (GB_GCA_900318935.1)
Olegusella sp000466405 (RS_GCF_000466405.1)
Olegusella massiliensis (RS_GCF_900078545.1)
Olegusella sp900315165 (GB_GCA_900315165.1)
Olegusella sp900314685 (GB_GCA_900314685.1)
Olegusella sp002407925 (GB_GCA_002407925.1)
Olsenella sp001189515 (RS_GCF_001189515.2)
Olsenella sp001457795 (RS_GCF_001457795.1)
Olsenella sp900316105 (GB_GCA_900316105.1)
Olsenella profusa (RS_GCF_000468755.1)
Olsenella urininfantis (RS_GCF_900155635.1)
Olsenella phocaeensis (RS_GCF_900120385.1)
Olsenella uli (RS_GCF_000143845.1)
Olsenella sp900313175 (GB_GCA_900313175.1)
Olsenella umbonata_A (RS_GCF_900111255.1)
Olsenella umbonata (RS_GCF_900105025.1)
Fannyhessea vaginae_A (RS_GCF_000179715.1)
Fannyhessea vaginae (RS_GCF_000159235.2)
Atopobium fossor (RS_GCF_000483125.1)
Atopobium minutum (RS_GCF_001437015.1)
Atopobium deltae (RS_GCF_001552785.1)
Atopobiaceae (metabat.574)
Parolsenella massiliensis (RS_GCF_900143685.1)

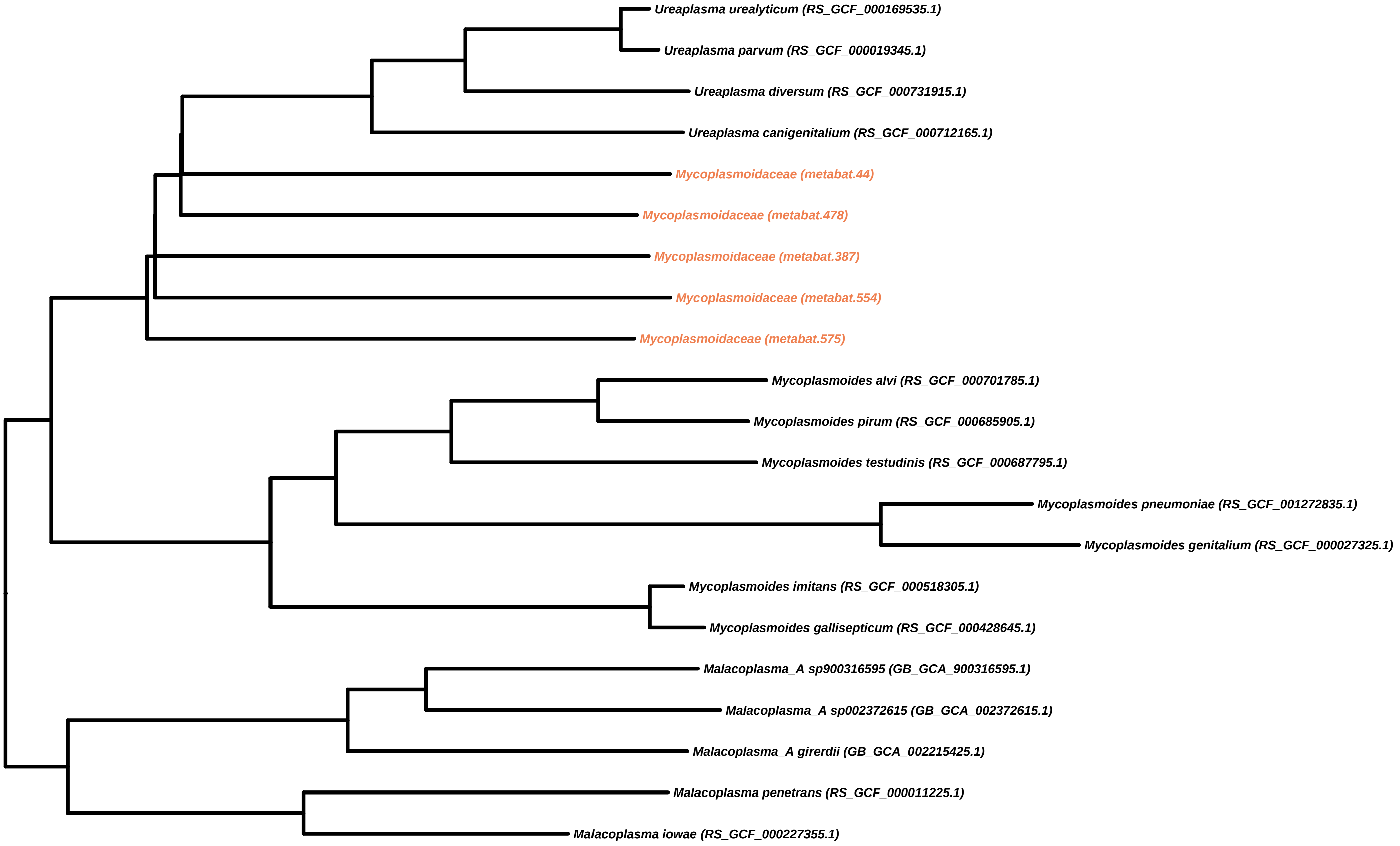

Ureaplasma urealyticum (RS_GCF_000169535.1)
Ureaplasma parvum (RS_GCF_000019345.1)
Ureaplasma diversum (RS_GCF_000731915.1)
Ureaplasma canigenitalium (RS_GCF_000712165.1)
Mycoplasmoidaceae (metabat.44)
Mycoplasmoidaceae (metabat.478)
Mycoplasmoidaceae (metabat.387)
Mycoplasmoidaceae (metabat.554)
Mycoplasmoidaceae (metabat.575)
Mycoplasmoides alvi (RS_GCF_000701785.1)
Mycoplasmoides pirum (RS_GCF_000685905.1)
Mycoplasmoides testudinis (RS_GCF_000687795.1)
Mycoplasmoides pneumoniae (RS_GCF_001272835.1)
Mycoplasmoides genitalium (RS_GCF_000027325.1)
Mycoplasmoides imitans (RS_GCF_000518305.1)
Mycoplasmoides gallisepticum (RS_GCF_000428645.1)
Malacoplasma_A sp900316595 (GB_GCA_900316595.1)
Malacoplasma_A sp002372615 (GB_GCA_002372615.1)
Malacoplasma_A girerdii (GB_GCA_002215425.1)
Malacoplasma penetrans (RS_GCF_000011225.1)
Malacoplasma iowae (RS_GCF_000227355.1)

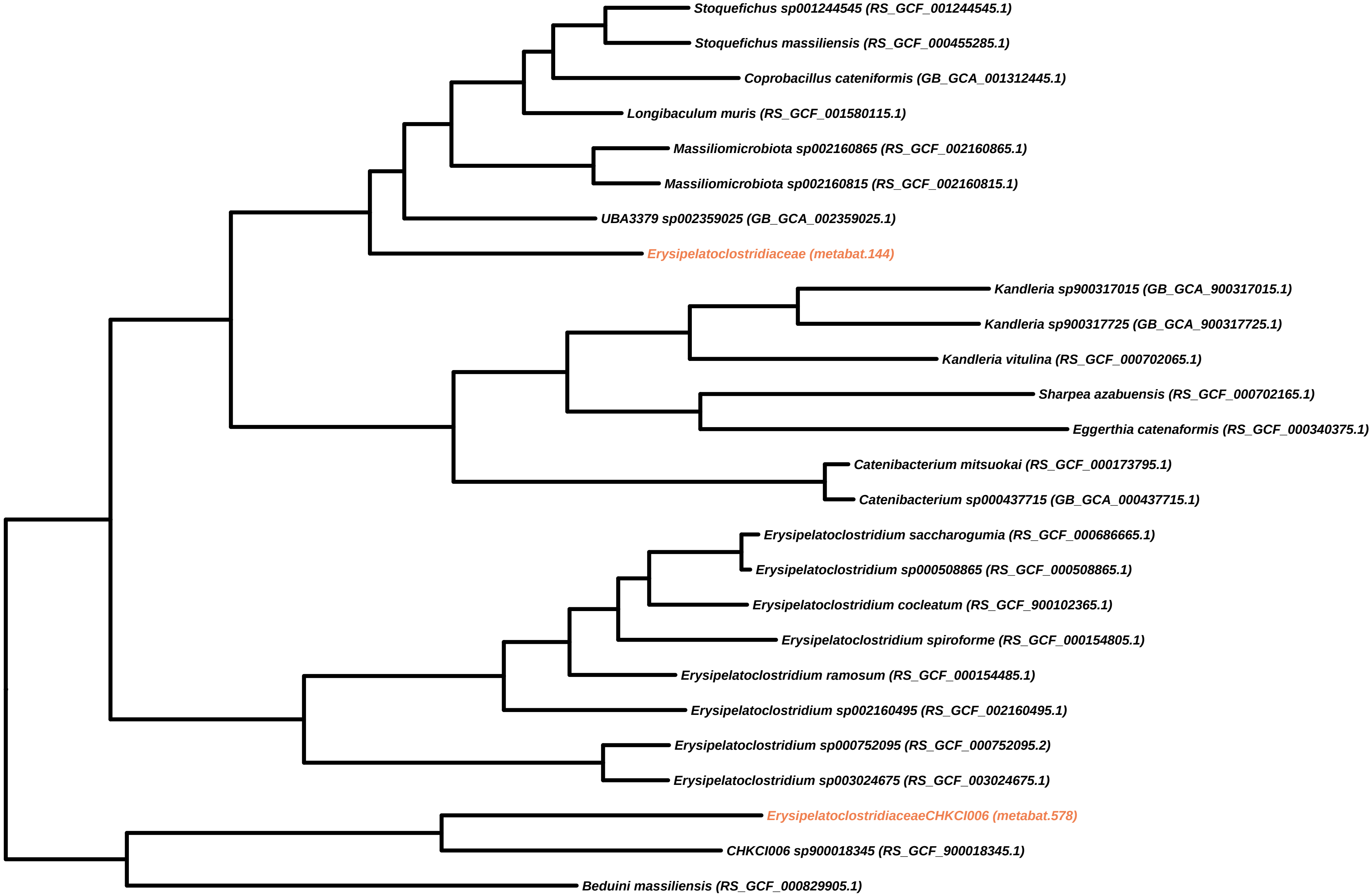

Stoquefichus sp001244545 (RS_GCF_001244545.1)
Stoquefichus massiliensis (RS_GCF_000455285.1)
Coprobacillus cateniformis (GB_GCA_001312445.1)
Longibaculum muris (RS_GCF_001580115.1)
Massiliomicrobiota sp002160865 (RS_GCF_002160865.1)
Massiliomicrobiota sp002160815 (RS_GCF_002160815.1)
UBA3379 sp002359025 (GB_GCA_002359025.1)
Erysipelatoclostridiaceae (metabat.144)
Kandleria sp900317015 (GB_GCA_900317015.1)
Kandleria sp900317725 (GB_GCA_900317725.1)
Kandleria vitulina (RS_GCF_000702065.1)
Sharpea azabuensis (RS_GCF_000702165.1)
Eggerthia catenaformis (RS_GCF_000340375.1)
Catenibacterium mitsuokai (RS_GCF_000173795.1)
Catenibacterium sp000437715 (GB_GCA_000437715.1)
Erysipelatoclostridium saccharogumia (RS_GCF_000686665.1)
Erysipelatoclostridium sp000508865 (RS_GCF_000508865.1)
Erysipelatoclostridium cocleatum (RS_GCF_900102365.1)
Erysipelatoclostridium spiroforme (RS_GCF_000154805.1)
Erysipelatoclostridium ramosum (RS_GCF_000154485.1)
Erysipelatoclostridium sp002160495 (RS_GCF_002160495.1)
Erysipelatoclostridium sp000752095 (RS_GCF_000752095.2)
Erysipelatoclostridium sp003024675 (RS_GCF_003024675.1)
ErysipelatoclostridiaceaeCHKCI006 (metabat.578)
CHKCI006 sp900018345 (RS_GCF_900018345.1)
Beduini massiliensis (RS_GCF_000829905.1)

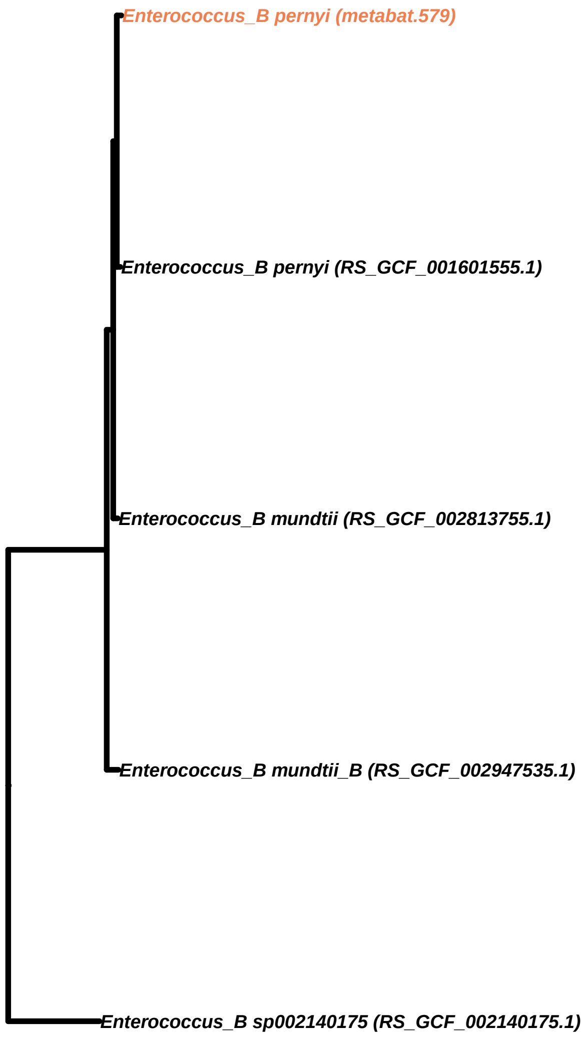
Enterococcus_B pernyi (metabat.579)
Enterococcus_B pernyi (RS_GCF_001601555.1)
Enterococcus_B mundtii (RS_GCF_002813755.1)
Enterococcus_B mundtii_B (RS_GCF_002947535.1)
Enterococcus_B sp002140175 (RS_GCF_002140175.1)

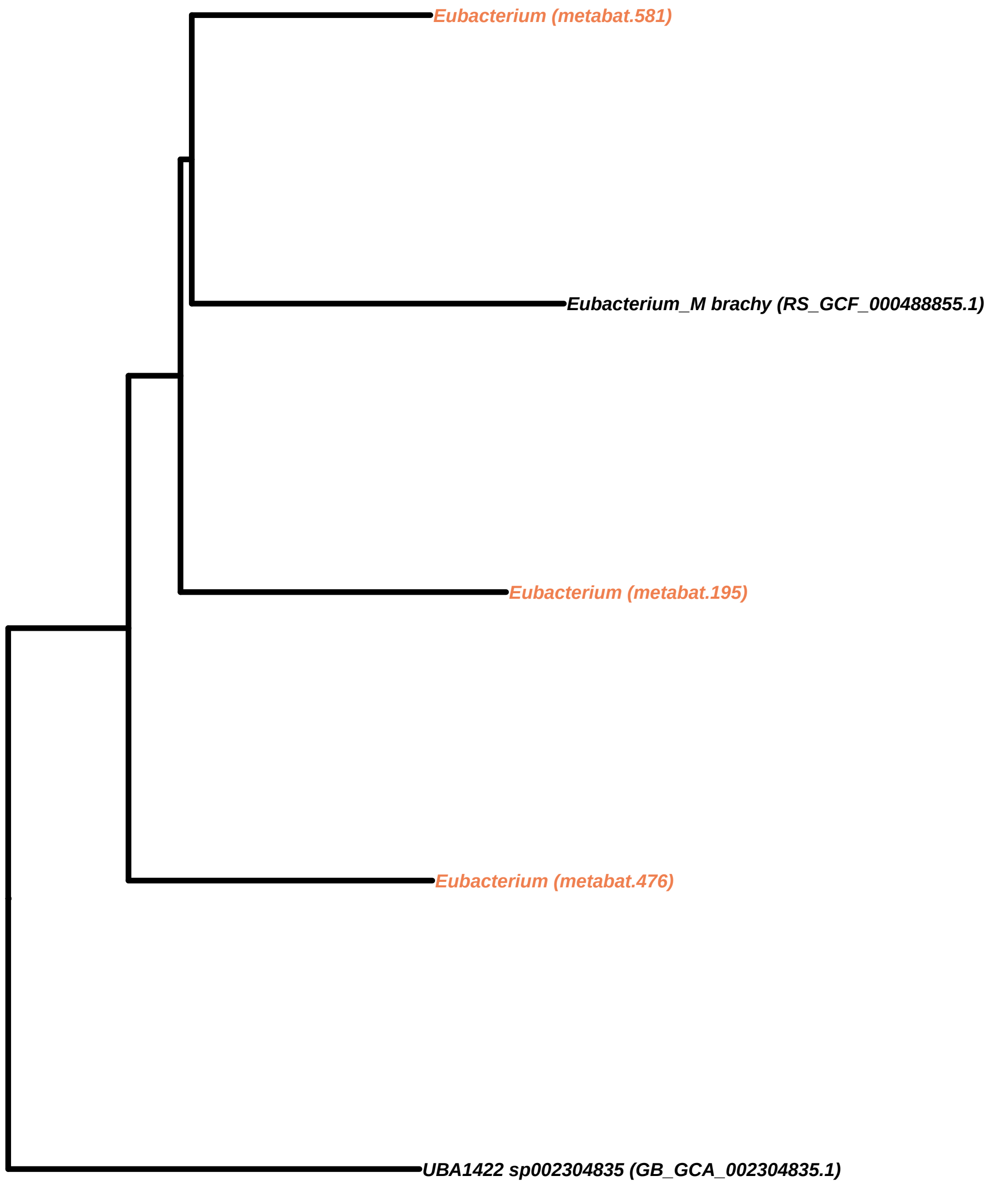
Eubacterium (metabat.581)
Eubacterium_M brachy (RS_GCF_000488855.1)
Eubacterium (metabat.195)
Eubacterium (metabat.476)
UBA1422 sp002304835 (GB_GCA_002304835.1)

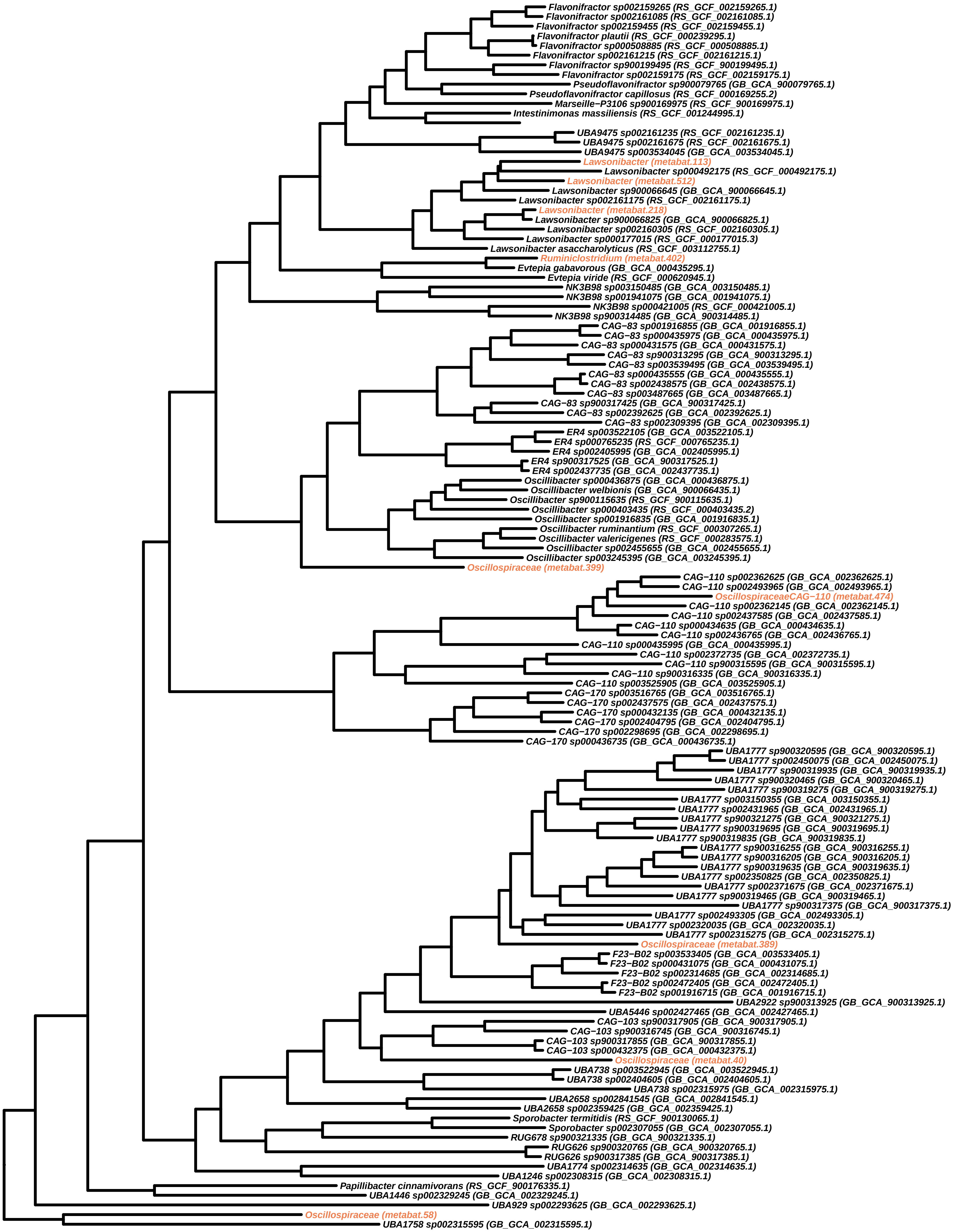

Flavonifractor sp002159265 (RS_GCF_002159265.1)
Flavonifractor sp002161085 (RS_GCF_002161085.1)
Flavonifractor sp002159455 (RS_GCF_002159455.1)
Flavonifractor plautii (RS_GCF_000239295.1)
Flavonifractor sp000508885 (RS_GCF_000508885.1)
Flavonifractor sp002161215 (RS_GCF_002161215.1)
Flavonifractor sp900199495 (RS_GCF_900199495.1)
Flavonifractor sp002159175 (RS_GCF_002159175.1)
Pseudoflavonifractor sp900079765 (GB_GCA_900079765.1)
Pseudoflavonifractor capillosus (RS_GCF_000169255.2)
Marseille−P3106 sp900169975 (RS_GCF_900169975.1)
Intestinimonas massiliensis (RS_GCF_001244995.1)
UBA9475 sp002161235 (RS_GCF_002161235.1)
UBA9475 sp002161675 (RS_GCF_002161675.1)
UBA9475 sp003534045 (GB_GCA_003534045.1)
Lawsonibacter (metabat.113)
Lawsonibacter sp000492175 (RS_GCF_000492175.1)
Lawsonibacter (metabat.512)
Lawsonibacter sp900066645 (GB_GCA_900066645.1)
Lawsonibacter sp002161175 (RS_GCF_002161175.1)
Lawsonibacter (metabat.218)
Lawsonibacter sp900066825 (GB_GCA_900066825.1)
Lawsonibacter sp002160305 (RS_GCF_002160305.1)
Lawsonibacter sp000177015 (RS_GCF_000177015.3)
Lawsonibacter asaccharolyticus (RS_GCF_003112755.1)
Ruminiclostridium (metabat.402)
Evtepia gabavorous (GB_GCA_000435295.1)
Evtepia viride (RS_GCF_000620945.1)
NK3B98 sp003150485 (GB_GCA_003150485.1)
NK3B98 sp001941075 (GB_GCA_001941075.1)
NK3B98 sp000421005 (RS_GCF_000421005.1)
NK3B98 sp900314485 (GB_GCA_900314485.1)
CAG−83 sp001916855 (GB_GCA_001916855.1)
CAG−83 sp000435975 (GB_GCA_000435975.1)
CAG−83 sp000431575 (GB_GCA_000431575.1)
CAG−83 sp900313295 (GB_GCA_900313295.1)
CAG−83 sp003539495 (GB_GCA_003539495.1)
CAG−83 sp000435555 (GB_GCA_000435555.1)
CAG−83 sp002438575 (GB_GCA_002438575.1)
CAG−83 sp003487665 (GB_GCA_003487665.1)
CAG−83 sp900317425 (GB_GCA_900317425.1)
CAG−83 sp002392625 (GB_GCA_002392625.1)
CAG−83 sp002309395 (GB_GCA_002309395.1)
ER4 sp003522105 (GB_GCA_003522105.1)
ER4 sp000765235 (RS_GCF_000765235.1)
ER4 sp002405995 (GB_GCA_002405995.1)
ER4 sp900317525 (GB_GCA_900317525.1)
ER4 sp002437735 (GB_GCA_002437735.1)
Oscillibacter sp000436875 (GB_GCA_000436875.1)
Oscillibacter welbionis (GB_GCA_900066435.1)
Oscillibacter sp900115635 (RS_GCF_900115635.1)
Oscillibacter sp000403435 (RS_GCF_000403435.2)
Oscillibacter sp001916835 (GB_GCA_001916835.1)
Oscillibacter ruminantium (RS_GCF_000307265.1)
Oscillibacter valericigenes (RS_GCF_000283575.1)
Oscillibacter sp002455655 (GB_GCA_002455655.1)
Oscillibacter sp003245395 (GB_GCA_003245395.1)
Oscillospiraceae (metabat.399)
CAG−110 sp002362625 (GB_GCA_002362625.1)
CAG−110 sp002493965 (GB_GCA_002493965.1)
OscillospiraceaeCAG−110 (metabat.474)
CAG−110 sp002362145 (GB_GCA_002362145.1)
CAG−110 sp002437585 (GB_GCA_002437585.1)
CAG−110 sp000434635 (GB_GCA_000434635.1)
CAG−110 sp002436765 (GB_GCA_002436765.1)
CAG−110 sp000435995 (GB_GCA_000435995.1)
CAG−110 sp002372735 (GB_GCA_002372735.1)
CAG−110 sp900315595 (GB_GCA_900315595.1)
CAG−110 sp900316335 (GB_GCA_900316335.1)
CAG−110 sp003525905 (GB_GCA_003525905.1)
CAG−170 sp003516765 (GB_GCA_003516765.1)
CAG−170 sp002437575 (GB_GCA_002437575.1)
CAG−170 sp000432135 (GB_GCA_000432135.1)
CAG−170 sp002404795 (GB_GCA_002404795.1)
CAG−170 sp002298695 (GB_GCA_002298695.1)
CAG−170 sp000436735 (GB_GCA_000436735.1)
UBA1777 sp900320595 (GB_GCA_900320595.1)
UBA1777 sp002450075 (GB_GCA_002450075.1)
UBA1777 sp900319935 (GB_GCA_900319935.1)
UBA1777 sp900320465 (GB_GCA_900320465.1)
UBA1777 sp900319275 (GB_GCA_900319275.1)
UBA1777 sp003150355 (GB_GCA_003150355.1)
UBA1777 sp002431965 (GB_GCA_002431965.1)
UBA1777 sp900321275 (GB_GCA_900321275.1)
UBA1777 sp900319695 (GB_GCA_900319695.1)
UBA1777 sp900319835 (GB_GCA_900319835.1)
UBA1777 sp900316255 (GB_GCA_900316255.1)
UBA1777 sp900316205 (GB_GCA_900316205.1)
UBA1777 sp900319635 (GB_GCA_900319635.1)
UBA1777 sp002350825 (GB_GCA_002350825.1)
UBA1777 sp002371675 (GB_GCA_002371675.1)
UBA1777 sp900319465 (GB_GCA_900319465.1)
UBA1777 sp900317375 (GB_GCA_900317375.1)
UBA1777 sp002493305 (GB_GCA_002493305.1)
UBA1777 sp002320035 (GB_GCA_002320035.1)
UBA1777 sp002315275 (GB_GCA_002315275.1)
Oscillospiraceae (metabat.389)
F23−B02 sp003533405 (GB_GCA_003533405.1)
F23−B02 sp000431075 (GB_GCA_000431075.1)
F23−B02 sp002314685 (GB_GCA_002314685.1)
F23−B02 sp002472405 (GB_GCA_002472405.1)
F23−B02 sp001916715 (GB_GCA_001916715.1)
UBA2922 sp900313925 (GB_GCA_900313925.1)
UBA5446 sp002427465 (GB_GCA_002427465.1)
CAG−103 sp900317905 (GB_GCA_900317905.1)
CAG−103 sp900316745 (GB_GCA_900316745.1)
CAG−103 sp900317855 (GB_GCA_900317855.1)
CAG−103 sp000432375 (GB_GCA_000432375.1)
Oscillospiraceae (metabat.40)
UBA738 sp003522945 (GB_GCA_003522945.1)
UBA738 sp002404605 (GB_GCA_002404605.1)
UBA738 sp002315975 (GB_GCA_002315975.1)
UBA2658 sp002841545 (GB_GCA_002841545.1)
UBA2658 sp002359425 (GB_GCA_002359425.1)
Sporobacter termitidis (RS_GCF_900130065.1)
Sporobacter sp002307055 (GB_GCA_002307055.1)
RUG678 sp900321335 (GB_GCA_900321335.1)
RUG626 sp900320765 (GB_GCA_900320765.1)
RUG626 sp900317385 (GB_GCA_900317385.1)
UBA1774 sp002314635 (GB_GCA_002314635.1)
UBA1246 sp002308315 (GB_GCA_002308315.1)
Papillibacter cinnamivorans (RS_GCF_900176335.1)
UBA1446 sp002329245 (GB_GCA_002329245.1)
UBA929 sp002293625 (GB_GCA_002293625.1)
Oscillospiraceae (metabat.58)
UBA1758 sp002315595 (GB_GCA_002315595.1)

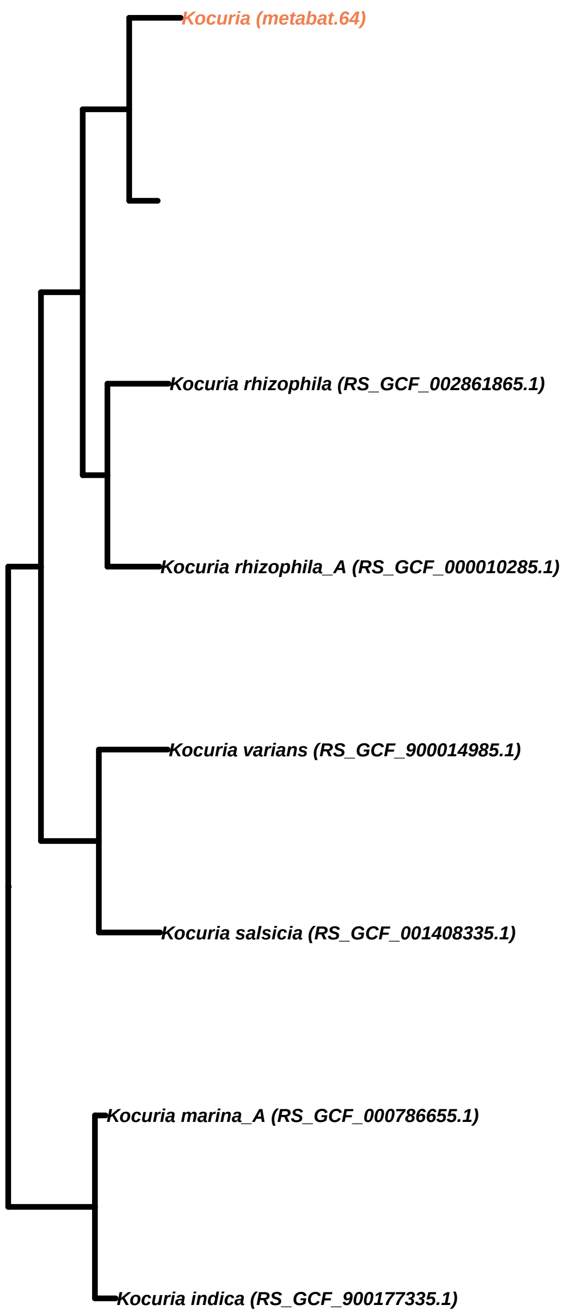
Kocuria (metabat.64)
Kocuria rhizophila (RS_GCF_002861865.1)
Kocuria rhizophila_A (RS_GCF_000010285.1)
Kocuria varians (RS_GCF_900014985.1)
Kocuria salsicia (RS_GCF_001408335.1)
Kocuria marina_A (RS_GCF_000786655.1)
Kocuria indica (RS_GCF_900177335.1)

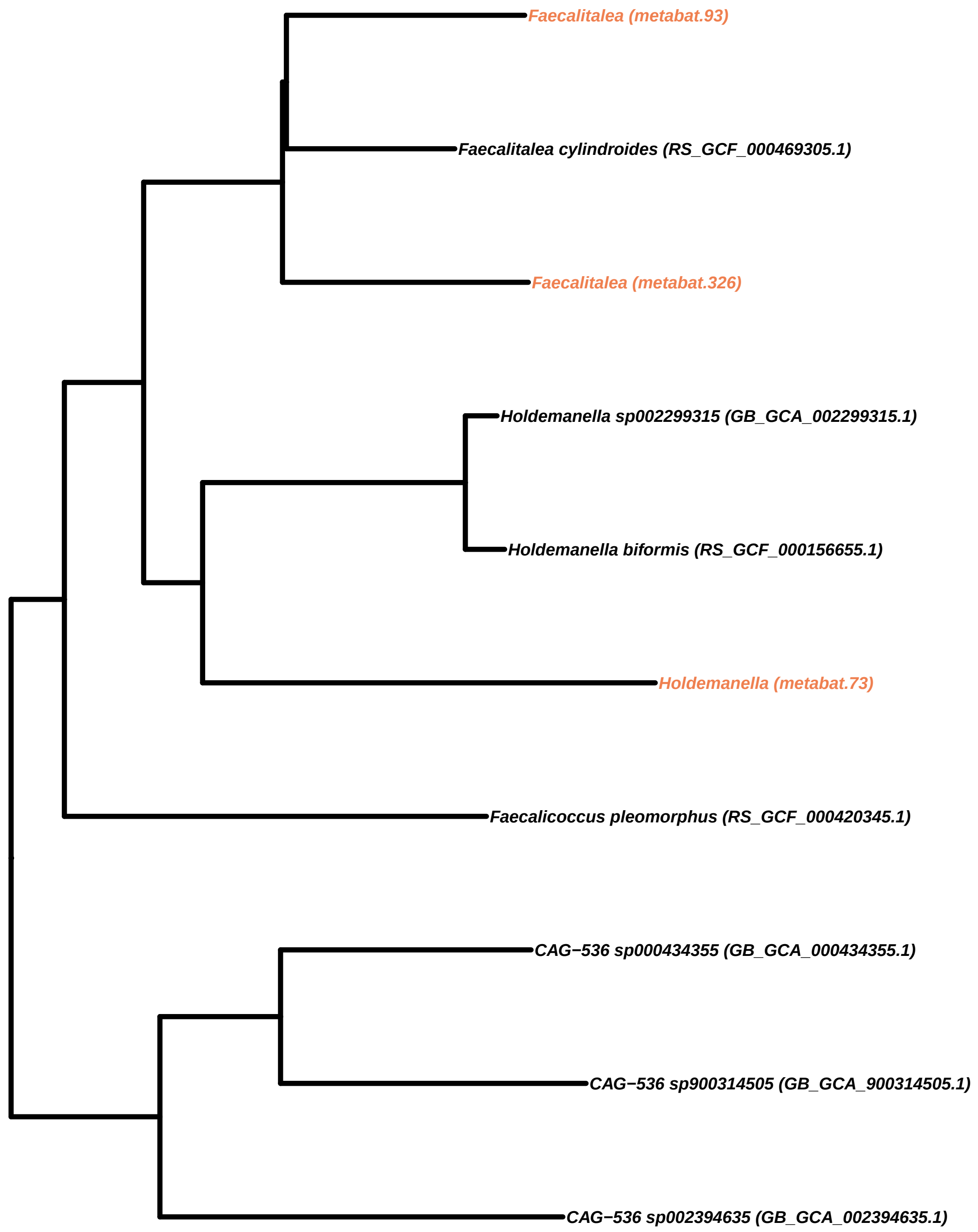
Faecalitalea (metabat.93)
Faecalitalea cylindroides (RS_GCF_000469305.1)
Faecalitalea (metabat.326)
Holdemanella sp002299315 (GB_GCA_002299315.1)
Holdemanella biformis (RS_GCF_000156655.1)
Holdemanella (metabat.73)
Faecalicoccus pleomorphus (RS_GCF_000420345.1)
CAG−536 sp000434355 (GB_GCA_000434355.1)
CAG−536 sp900314505 (GB_GCA_900314505.1)
CAG−536 sp002394635 (GB_GCA_002394635.1)

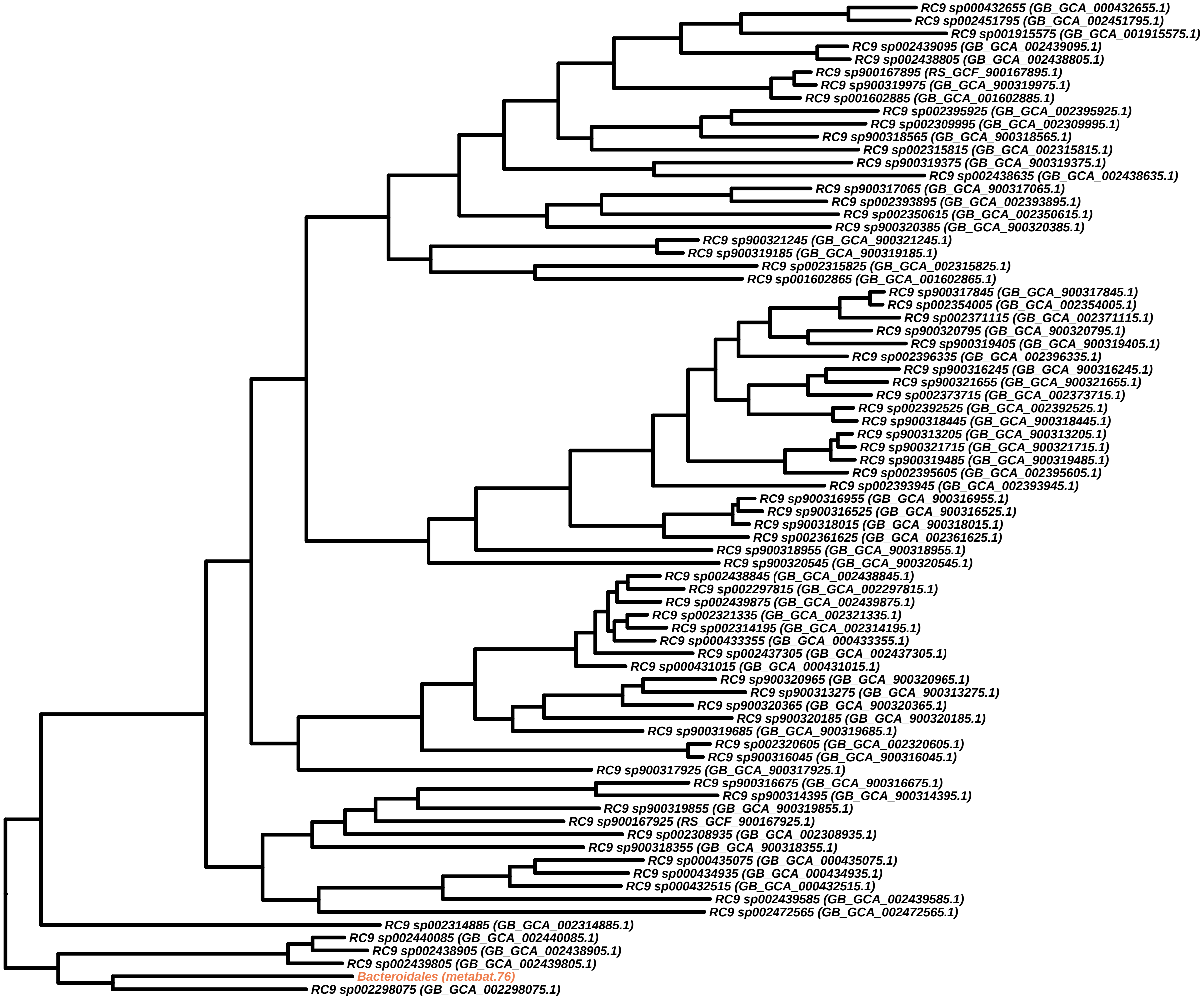
RC9 sp000432655 (GB_GCA_000432655.1)
RC9 sp002451795 (GB_GCA_002451795.1)
RC9 sp001915575 (GB_GCA_001915575.1)
RC9 sp002439095 (GB_GCA_002439095.1)
RC9 sp002438805 (GB_GCA_002438805.1)
RC9 sp900167895 (RS_GCF_900167895.1)
RC9 sp900319975 (GB_GCA_900319975.1)
RC9 sp001602885 (GB_GCA_001602885.1)
RC9 sp002395925 (GB_GCA_002395925.1)
RC9 sp002309995 (GB_GCA_002309995.1)
RC9 sp900318565 (GB_GCA_900318565.1)
RC9 sp002315815 (GB_GCA_002315815.1)
RC9 sp900319375 (GB_GCA_900319375.1)
RC9 sp002438635 (GB_GCA_002438635.1)
RC9 sp900317065 (GB_GCA_900317065.1)
RC9 sp002393895 (GB_GCA_002393895.1)
RC9 sp002350615 (GB_GCA_002350615.1)
RC9 sp900320385 (GB_GCA_900320385.1)
RC9 sp900321245 (GB_GCA_900321245.1)
RC9 sp900319185 (GB_GCA_900319185.1)
RC9 sp002315825 (GB_GCA_002315825.1)
RC9 sp001602865 (GB_GCA_001602865.1)
RC9 sp900317845 (GB_GCA_900317845.1)
RC9 sp002354005 (GB_GCA_002354005.1)
RC9 sp002371115 (GB_GCA_002371115.1)
RC9 sp900320795 (GB_GCA_900320795.1)
RC9 sp900319405 (GB_GCA_900319405.1)
RC9 sp002396335 (GB_GCA_002396335.1)
RC9 sp900316245 (GB_GCA_900316245.1)
RC9 sp900321655 (GB_GCA_900321655.1)
RC9 sp002373715 (GB_GCA_002373715.1)
RC9 sp002392525 (GB_GCA_002392525.1)
RC9 sp900318445 (GB_GCA_900318445.1)
RC9 sp900313205 (GB_GCA_900313205.1)
RC9 sp900321715 (GB_GCA_900321715.1)
RC9 sp900319485 (GB_GCA_900319485.1)
RC9 sp002395605 (GB_GCA_002395605.1)
RC9 sp002393945 (GB_GCA_002393945.1)
RC9 sp900316955 (GB_GCA_900316955.1)
RC9 sp900316525 (GB_GCA_900316525.1)
RC9 sp900318015 (GB_GCA_900318015.1)
RC9 sp002361625 (GB_GCA_002361625.1)
RC9 sp900318955 (GB_GCA_900318955.1)
RC9 sp900320545 (GB_GCA_900320545.1)
RC9 sp002438845 (GB_GCA_002438845.1)
RC9 sp002297815 (GB_GCA_002297815.1)
RC9 sp002439875 (GB_GCA_002439875.1)
RC9 sp002321335 (GB_GCA_002321335.1)
RC9 sp002314195 (GB_GCA_002314195.1)
RC9 sp000433355 (GB_GCA_000433355.1)
RC9 sp002437305 (GB_GCA_002437305.1)
RC9 sp000431015 (GB_GCA_000431015.1)
RC9 sp900320965 (GB_GCA_900320965.1)
RC9 sp900313275 (GB_GCA_900313275.1)
RC9 sp900320365 (GB_GCA_900320365.1)
RC9 sp900320185 (GB_GCA_900320185.1)
RC9 sp900319685 (GB_GCA_900319685.1)
RC9 sp002320605 (GB_GCA_002320605.1)
RC9 sp900316045 (GB_GCA_900316045.1)
RC9 sp900317925 (GB_GCA_900317925.1)
RC9 sp900316675 (GB_GCA_900316675.1)
RC9 sp900314395 (GB_GCA_900314395.1)
RC9 sp900319855 (GB_GCA_900319855.1)
RC9 sp900167925 (RS_GCF_900167925.1)
RC9 sp002308935 (GB_GCA_002308935.1)
RC9 sp900318355 (GB_GCA_900318355.1)
RC9 sp000435075 (GB_GCA_000435075.1)
RC9 sp000434935 (GB_GCA_000434935.1)
RC9 sp000432515 (GB_GCA_000432515.1)
RC9 sp002439585 (GB_GCA_002439585.1)
RC9 sp002472565 (GB_GCA_002472565.1)
RC9 sp002314885 (GB_GCA_002314885.1)
RC9 sp002440085 (GB_GCA_002440085.1)
RC9 sp002438905 (GB_GCA_002438905.1)
RC9 sp002439805 (GB_GCA_002439805.1)
Bacteroidales (metabat.76)
RC9 sp002298075 (GB_GCA_002298075.1)

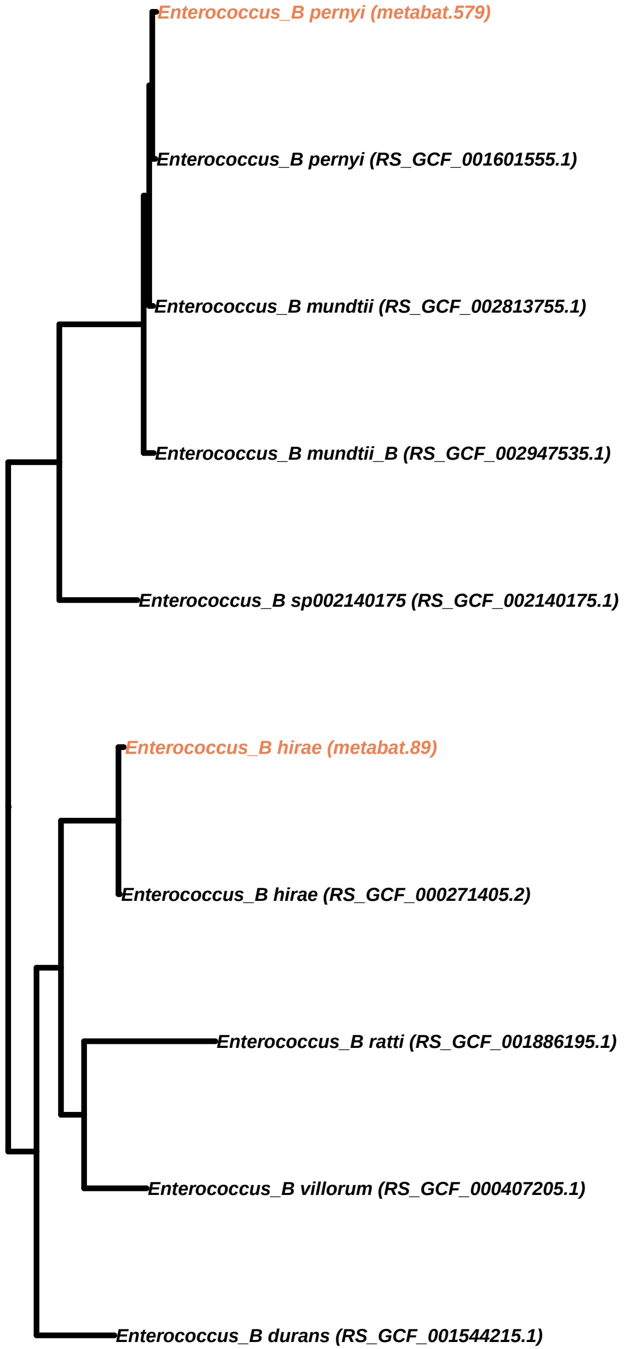

Enterococcus_B pernyi (metabat.579)
Enterococcus_B pernyi (RS_GCF_001601555.1)
Enterococcus_B mundtii (RS_GCF_002813755.1)
Enterococcus_B mundtii_B (RS_GCF_002947535.1)
Enterococcus_B sp002140175 (RS_GCF_002140175.1)
Enterococcus_B hirae (metabat.89)
Enterococcus_B hirae (RS_GCF_000271405.2)
Enterococcus_B ratti (RS_GCF_001886195.1)
Enterococcus_B villorum (RS_GCF_000407205.1)
Enterococcus_B durans (RS_GCF_001544215.1)

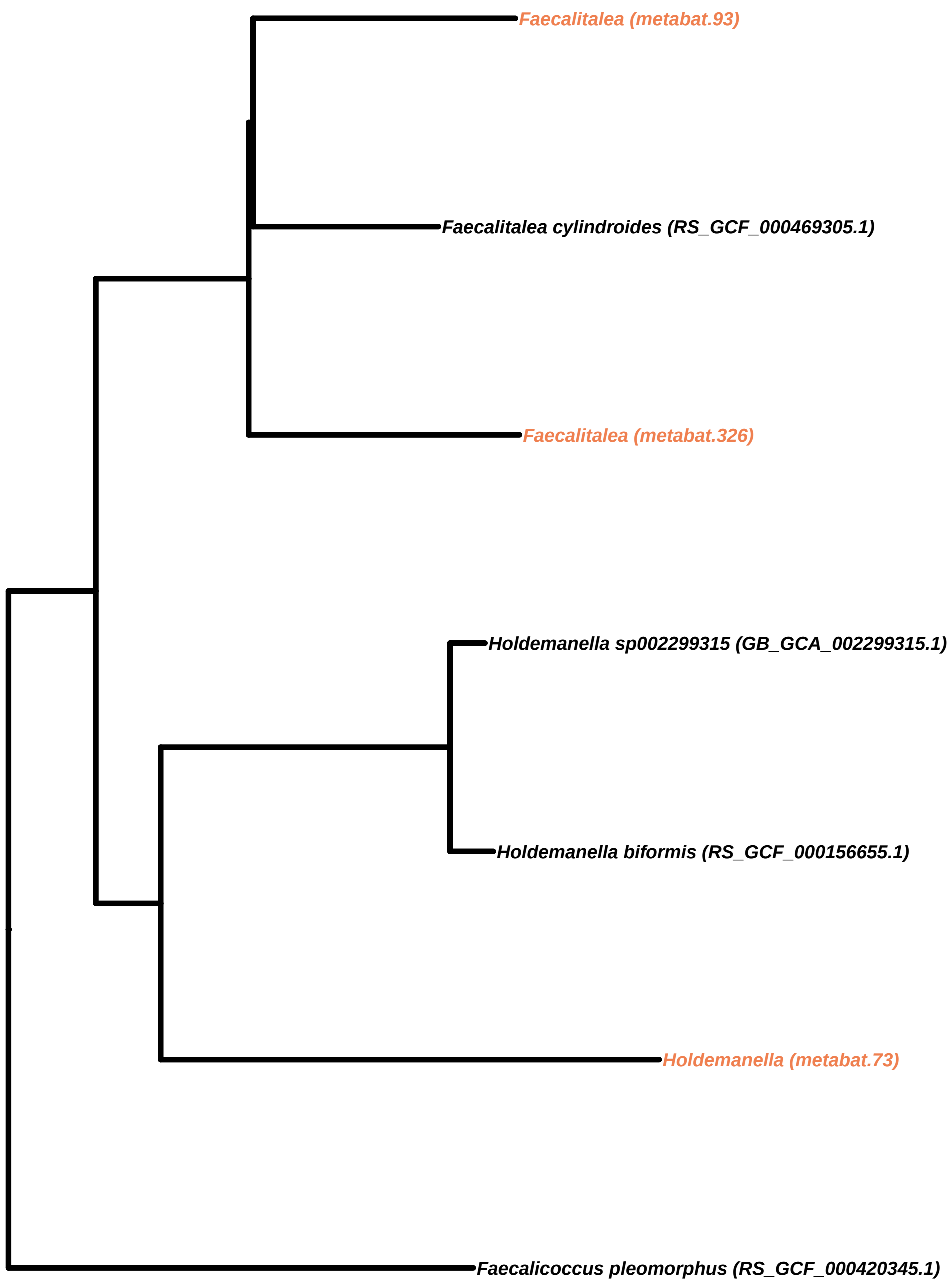

Faecalitalea (metabat.93)
Faecalitalea cylindroides (RS_GCF_000469305.1)
Faecalitalea (metabat.326)
Holdemanella sp002299315 (GB_GCA_002299315.1)
Holdemanella biformis (RS_GCF_000156655.1)
Holdemanella (metabat.73)
Faecalicoccus pleomorphus (RS_GCF_000420345.1)

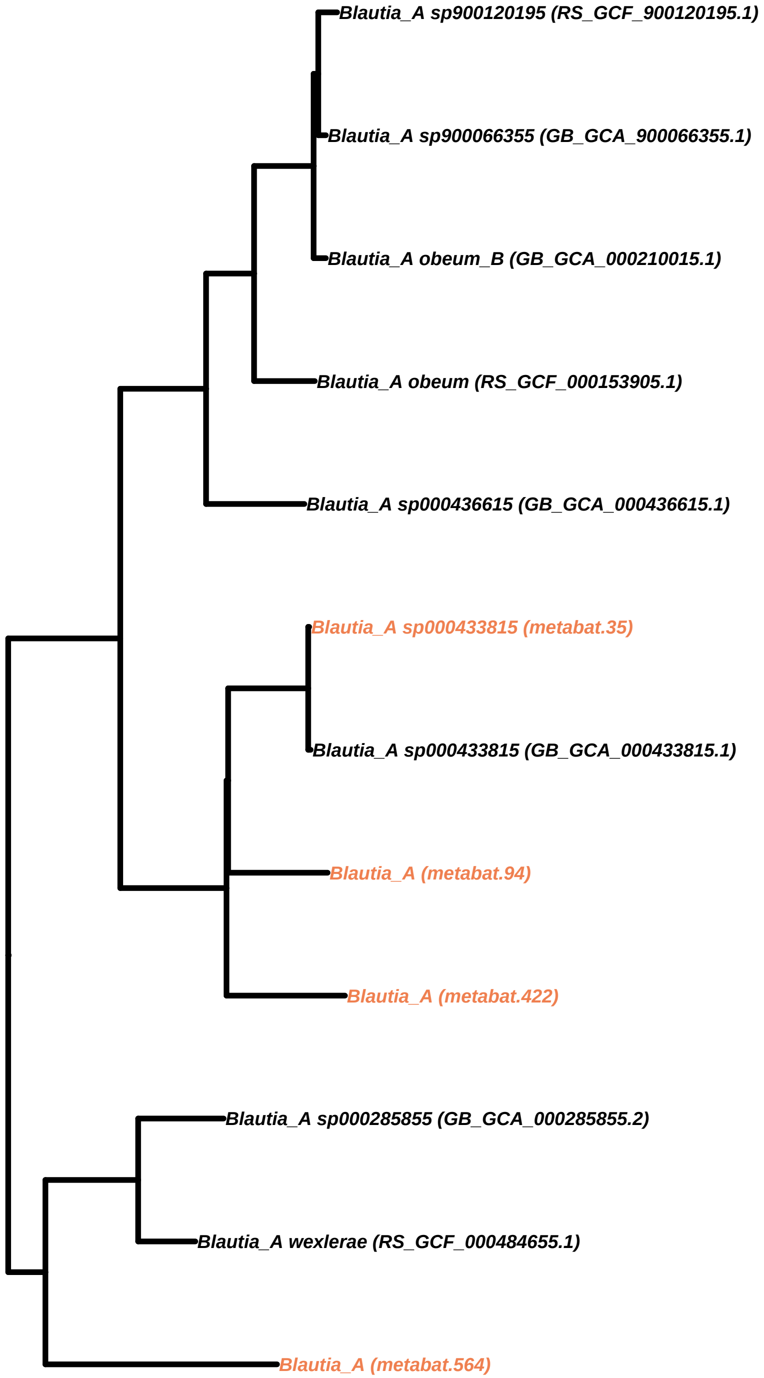
Blautia_A sp900120195 (RS_GCF_900120195.1)
Blautia_A sp900066355 (GB_GCA_900066355.1)
Blautia_A obeum_B (GB_GCA_000210015.1)
Blautia_A obeum (RS_GCF_000153905.1)
Blautia_A sp000436615 (GB_GCA_000436615.1)
Blautia_A sp000433815 (metabat.35)
Blautia_A sp000433815 (GB_GCA_000433815.1)
Blautia_A (metabat.94)
Blautia_A (metabat.422)
Blautia_A sp000285855 (GB_GCA_000285855.2)
Blautia_A wexlerae (RS_GCF_000484655.1)
Blautia_A (metabat.564)

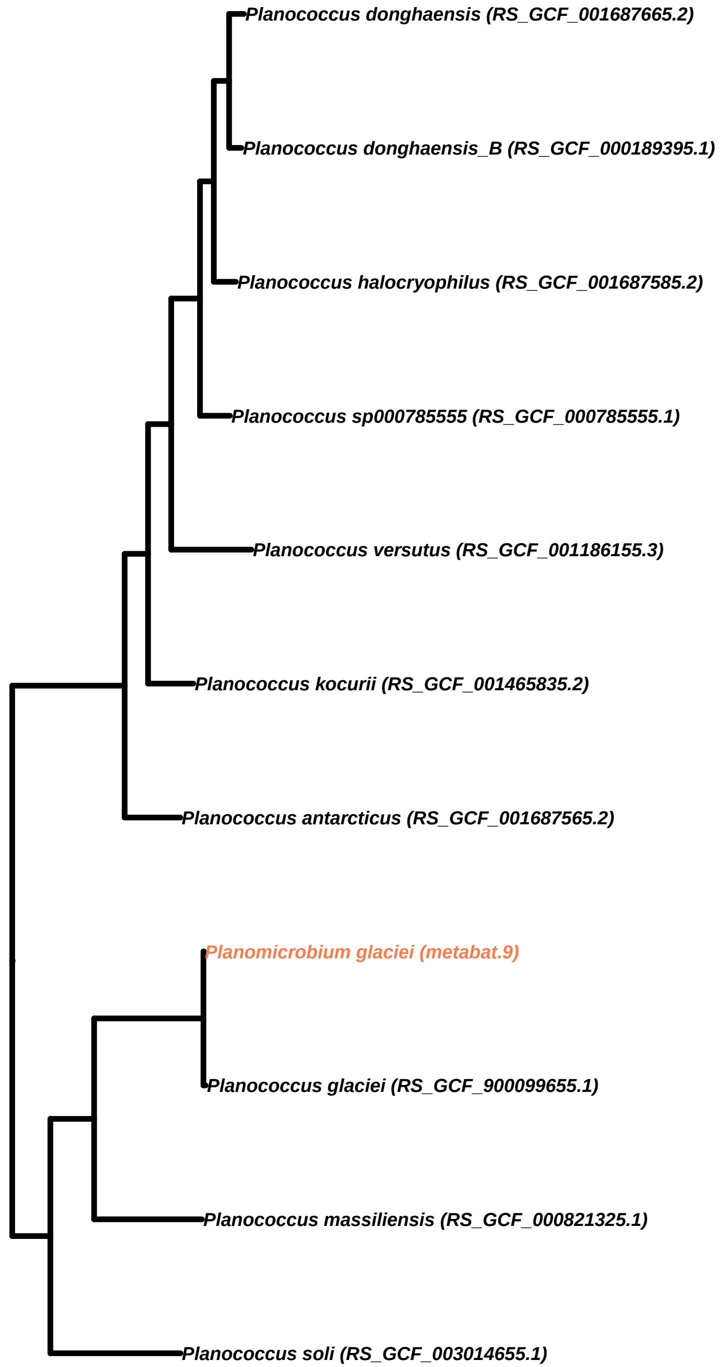
Planococcus donghaensis (RS_GCF_001687665.2)
Planococcus donghaensis_B (RS_GCF_000189395.1)
Planococcus halocryophilus (RS_GCF_001687585.2)
Planococcus sp000785555 (RS_GCF_000785555.1)
Planococcus versutus (RS_GCF_001186155.3)
Planococcus kocurii (RS_GCF_001465835.2)
Planococcus antarcticus (RS_GCF_001687565.2)
Planomicrobium glaciei (metabat.9)
Planococcus glaciei (RS_GCF_900099655.1)
Planococcus massiliensis (RS_GCF_000821325.1)
Planococcus soli (RS_GCF_003014655.1)
